# Investigation of the underlying hub genes and molexular pathogensis in gastric cancer by integrated bioinformatic analyses

**DOI:** 10.1101/2020.12.20.423656

**Authors:** Basavaraj Vastrad, Chanabasayya Vastrad

## Abstract

The high mortality rate of gastric cancer (GC) is in part due to the absence of initial disclosure of its biomarkers. The recognition of important genes associated in GC is therefore recommended to advance clinical prognosis, diagnosis and and treatment outcomes. The current investigation used the microarray dataset GSE113255 RNA seq data from the Gene Expression Omnibus database to diagnose differentially expressed genes (DEGs). Pathway and gene ontology enrichment analyses were performed, and a protein protein interaction network, modules, target genes - miRNA regulatory network and target genes - TF regulatory network were constructed and analyzed. Finally, validation of hub genes was performed. The 1008 DEGs identified consisted of 505 up regulated genes and 503 down regulated genes. The pathways and GO functions of the up and down regulated genes were mainly enriched in pyrimidine deoxyribonucleosides degradation, extracellular structure organization, allopregnanolone biosynthesis and digestion. FN1, PLK1, ANLN, MCM7, MCM2, EEF1A2, PTGER3, CKB, ERBB4 and PRKAA2 were identified as the most important genes of GC, and validated by TCGA database, The Human Protein Atlas database, receiver operating characteristic curve (ROC) analysis and RT-PCR. Bioinformatics analysis might be useful method to explore the molecular pathogensis of GC. In addition, FN1, PLK1, ANLN, MCM7, MCM2, EEF1A2, PTGER3, CKB, ERBB4 and PRKAA2 might be the most important genes of GC.

## Introduction

Gastric cancer (GC) presents the major cause of cancer deaths in the past few decades and has been a major public health problem [1]. GC is the fifth most frequently detected cancer in worldwide [2]. Since GC absence reliable clinical and biochemical features in the initial stages, the majority of patients are only definitively identified with GC at a leading stage. Although some routine treatments such as surgery and chemotherapy can be used, the recurrence rate is still as high [3]. Thus, investigation on GC is important, especially on its molecular pathogenesis and the identification of biomarkers that are useful for early detection, risk stratification, determination of appropriate intervention, prognostication and identification of novel therapeutic targets.

Investigation has exposed key pathways and interactions between gene modfications, and tumorigenesis and tumor advancement of various types of cancers [4]. Pathways such as Wnt pathways [5], PI3K and MAPK signaling pathways [6], Notch1 signal pathway [7], STAT and ERK2 pathways [8] and Src/FAK pathway [9] were responsible for pathogenesis of GC. Genes such as IL- 10 and TNF-A [10], p53 and TGF-,RII [11], IL-17A and IL-17F [12], PFKFB-3 and PFKFB-4 [13], and Runx3 and CHFR [14] were liable for pathogenesis of GC. Thus, a complete understanding of the transcriptional modification may add to the advancement of preventive, diagnostic and therapeutic strategies for GC.

The prognosis and diagnosis of GC mainly depends on the extent of disease. With the fast advancement of gene or RNA sequencing technology, GEO, ArrayExpress and TCGA have been playing enhancing key roles in bioinformatics analysis [15–17]. These databases provide sequencing data for research of novel functional genes and pathways and for analyzing the effect of these genes on prognosis, diagnosis and therapeutic.

The current investigation aimed to improve the understanding of the molecular basis of GC and to identify novel prognostic factors for GC. The GSE113255 RNA seq dataset was downloaded from the Gene Expression Omnibus (GEO) database, in order to determine the up regulated and down regulated differentially expressed genes (DEGs) in the tumor tissues of patients with GC. Several prognosis and diagnosis related genes and pathways of GC were obtained through, pathway and gene ontology (GO) enrichment analysis, protein– protein interaction (PPI) network analysis, module analysis, target genes - miRNA regulatory network analysis and target genes – TF regulatory network analysis. To verify the prognostic roles of hub genes in GC, the GC dataset from The Cancer Genome Atlas (TCGA) was used to perform survival analysis, expression analysis, stage analysis, mutation analysis, immune histochemical (ICH) analysis and immune infiltration analysis. Secondly, the current investigation confirmed the selected findings using receiver operating characteristic (ROC) analysis), with the aim of prognostic and diagnostic value of hub genes. Finally, the most key expressed hub genes were selected and conducted for preliminary validation by real-time PCR (RT-PCR). In summery we believe that it is feasible to identify new genes linked with GC with the explained method, providing key intuition into the GC at the molecular level that guided the GC advancement and progression to full blown stages.

## Materials and methods

### Microarray data

RNA-seq Data relevant to GC was obtained from the GEO database and the GSE113255 (https://www.ncbi.nlm.nih.gov/geo/query/acc.cgi?acc=GSE113255) dataset was selected [18]. This dataset was established on a GPL18573 Illumina NextSeq 500 (Homo sapiens) platform. The GSE113255 dataset contains 107 diffuse type gastric cancer tissues samples and 10 normal gastric tissues samples.

### Data pre-processing and identification of DEGs in GC

Microarray data preprocessing is associated with quality assessment, quality control, background correction and normalization. Quantile method was used to perform background correction, normalization and probe summarization [19]. The empirical Bayesian method in limma package [20] was used to perform the differential analysis, |log2 fold change (FC)| >1.5 for up regulated genes, |log2 fold change (FC)| < −1.13 for down regulated genes and adjusted P<0.05 were considered as statistically significant. Since dataset was diagnosed to have a larger sample size of both GC and healthy tissues, heat maps and volcano plots were further established from the RNA-seq data of this dataset in order to visualize DEGs.

### Pathway enrichment analysis of DEGs

BIOCYC (https://biocyc.org/) [21], Kyoto Encyclopedia of Genes and Genomes (KEGG) (http://www.genome.jp/kegg/pathway.html) [22], Pathway Interaction Database (PID) (https://wiki.nci.nih.gov/pages/viewpage.action?pageId=315491760) [23], REACTOME (https://reactome.org/) [24], GenMAPP (http://www.genmapp.org/) [25], MSigDB C2 BIOCARTA (http://software.broadinstitute.org/gsea/msigdb/collections.jsp) [26], PantherDB (http://www.pantherdb.org/) [27], Pathway Ontology (http://www.obofoundry.org/ontology/pw.html) [28] and Small Molecule Pathway Database (SMPDB) (http://smpdb.ca/) [29] are a databases collection which can be used to analyze genomes, biological pathways, diseases, chemical substances and drugs. ToppGene (ToppFun) (https://toppgene.cchmc.org/enrichment.jsp) [30] was used for annotation of the pathway results. For this analyses, p<0 .05 was considered to indicate a statistically significant difference.

### Gene ontology (GO) enrichment analysis of DEGs

ToppGene (ToppFun) (https://toppgene.cchmc.org/enrichment.jsp) [30] was used to perform GO enrichment analysis. GO (http://www.geneontology.org/) enrichment analysis is extensively used to annotate specific genes and gene products for high throughput genome and transcriptome data [31]. In the current investigation, GO enrichment analysis was implemented to predict the probable functions of the DEGs based on biological process (BP), molecular function (MF) and cellular component (CC). p<0.05 was set as the cutoff criterion.

### PPI network construction and module analysis

We evaluated the protein–protein interaction (PPI) data using the Human Integrated Protein-Protein Interaction rEference (HIPPIE) (http://cbdm.uni-mainz.de/hippie/) database [32], which integrates with various PPI data bases such as IntAct (https://www.ebi.ac.uk/intact/) [33], BioGRID (https://thebiogrid.org/) [34], HPRD (http://www.hprd.org/) [35], MINT (https://mint.bio.uniroma2.it/) [36], BIND (http://download.baderlab.org/BINDTranslation/) [37], MIPS (http://mips.helmholtz-muenchen.de/proj/ppi/) [38] and DIP (http://dip.doe-mbi.ucla.edu/dip/Main.cgi) [39]. Cytoscape software (http://www.cytoscape.org/) [40] was applied to visualize the protein interaction network relationships. With the plugin Network analyzer the gene lists of the top ranked in node degree [41], betweenness centrality [42], stress centrality [43], closeness centrality [44] and clustering coefficient [45] were obtained. PEWCC1 (http://apps.cytoscape.org/apps/PEWCC1), a plug-in used to produce the best results for considerate correlation levels, was subsequently utilized to identify modules in the network [46].

### Target genes - miRNA regulatory network construction

Relevant miRNA targets were predicted using miRNet database (available online: (https://www.mirnet.ca/) [47], which is a comprehensive atlas of predicted and validated target - miRNA interactions. The potential targets of miRNA were identified by 10 miRNA databases such as TarBase (http://diana.imis.athena-innovation.gr/DianaTools/index.php?r=tarbase/index) [48], miRTarBase (http://mirtarbase.mbc.nctu.edu.tw/php/download.php) [49], miRecords (http://miRecords.umn.edu/miRecords) [50], miR2Disease (http://www.mir2disease.org/) [51], HMDD (http://www.cuilab.cn/hmdd) [52], PhenomiR (http://mips.helmholtz-muenchen.de/phenomir/) [53], SM2miR (http://bioinfo.hrbmu.edu.cn/SM2miR/) [54], PharmacomiR (http://www.pharmaco-mir.org/) [55], EpimiR (http://bioinfo.hrbmu.edu.cn/EpimiR/) [56] and starBase (http://starbase.sysu.edu.cn/) [57]. The integrated regulatory networks were then visualized by Cytoscape (http://www.cytoscape.org/) [40].

### Target genes - TF regulatory network construction

Relevant TF targets were predicted using (https://www.networkanalyst.ca/) [58], which is a comprehensive atlas of predicted and validated target - TF interactions. The potential targets of TF were identified by TF database ChEA (http://amp.pharm.mssm.edu/lib/chea.jsp) [59]. The integrated regulatory networks were then visualized by Cytoscape (http://www.cytoscape.org/) [40].

### Validations of hub genes

The probability of survival and importance was determined using the UALCAN (http://ualcan.path.uab.edu/analysis.html) [60] database. UALCAN is a newly created online interactive web server which implements users to examine the RNA sequencing expression data of tumors/normal tissues or samples from The Cancer Genome Atlas (TCGA), based on a criterion processing pipeline. The overall survival analyses of hub gene were performed in UALCAN. The RNA expression level of hub genes between GC samples and normal control samples was visualized by UALCAN. The RNA expression level of hub genes between different stages of GC samples compared to normal control samples was visualized by UALCAN. The cBio Cancer Genomics Portal (cBioPortal) (http://www.cbioportal.org) [61] is a key tool for online analysis, visualization, and examination of cancer genomics data. cBioPortal, which is the database used in this investigation, scrutinize genetic modifications of prognostic and diagnostic hub genes in TCGA GC patients. The expression levels of hub genes in cancerous and non cancerous tissue were validated using the Human Protein Atlas (HPA) database (http://www.proteinatlas.org/) [62]. To calculate whether established hub genes had key diagnostic and prognostic values for GC, the receiver operating characteristic (ROC) analyses was accomplished using the R “pROC” package [63]. The area under the ROC curve (AUC) of hub gene was determined. The diagnostic and prognostic accuracy of hub genes for GC was calculated with AUC value. Herein, when AUC value was larger than 0.8, the hub gene could differentiate GC from normal control. Hub genes were further validated by real time polymer chain reaction (RT-PCR). According to the manufacturer’s instruction, total RNA of gastric tissues in both GC groups and controls were extracted using the TRI Reagent® (Sigma, USA) and reversely transcribed to complementary DNA using the FastQuant RT kit (with gDNase; Tiangen Biotech Co., Ltd.). RT-PCR was completed by using QuantStudio 7 Flex real-time PCR system (Thermo Fisher Scientific, Waltham, MA, USA). The RT-PCR amplification reaction protocol was as follows: A total of 40 cycles at 95°C for 5 sec and 60°C for 10 sec. Relative quantification was achieved by using the comparative 2^ΔΔCq^ method [64] with β-actin as the reference gene. The sequences of all primer pairs are given in Table 1. Immune infiltration analysis was performed for hub genes using the TIMER online analysis tool (https://cistrome.shinyapps.io/timer/) [65], which incorporates the prognostic and diagnostic data from the RNA-Seq expression profiling database from The Cancer Genome Atlas (TCGA). Immune infiltration analysis was assigned to check the immune infiltrates (B cells, CD4+ T cells, CD8+ T cells, neutrophils, macrophages, and dendritic cells) across GC.

**Table 1.**
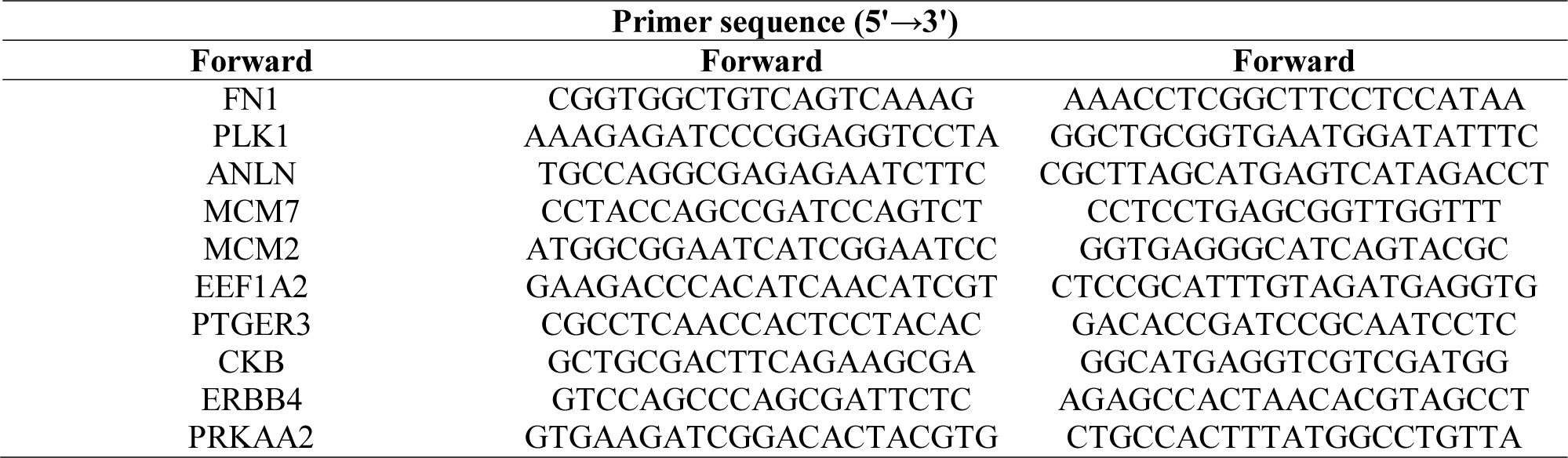
Primers used for quantitative PCR

## Results

### Data pre processing and identification of DEGs in GC

NCBI-GEO is a free database of gene expression profiles and RNA-seq, from which GC and normal control RNA-seq data of GSE113255 were obtained. The expression values for all the genes from the 107 GC samples and 10 normal control samples were normalized using the Quantile method, and values with an unchanged position in the boxplot were used for consequent analysis, as this can be used as a intermediary for normalization (Fig. 1A and Fig. 1B). Using P < 0.05, |log2 fold change (FC)| >1.5 for up regulated genes, |log2 fold change (FC)| < - 1.13 for down regulated genes as the cutoff criteria, after integrated bioinformatics analysis. A total of 1008 DEGs were identified in the active group, including 505 up regulated genes and 503 down regulated genes and are given in Table 2. Fig. 2 and Fig. 3 shows the heat maps of those up and down regulated genes, and reveals that the up and down regulated genes can be easily distinguished from each of the samples. The distribution patterns of expressed genes in GSE113255 data is displayed in Fig. 4, respectively. Red dots in the volcano plots mean significantly up regulated genes, while green dots mean significantly down regulated genes.

**Fig. 1.**
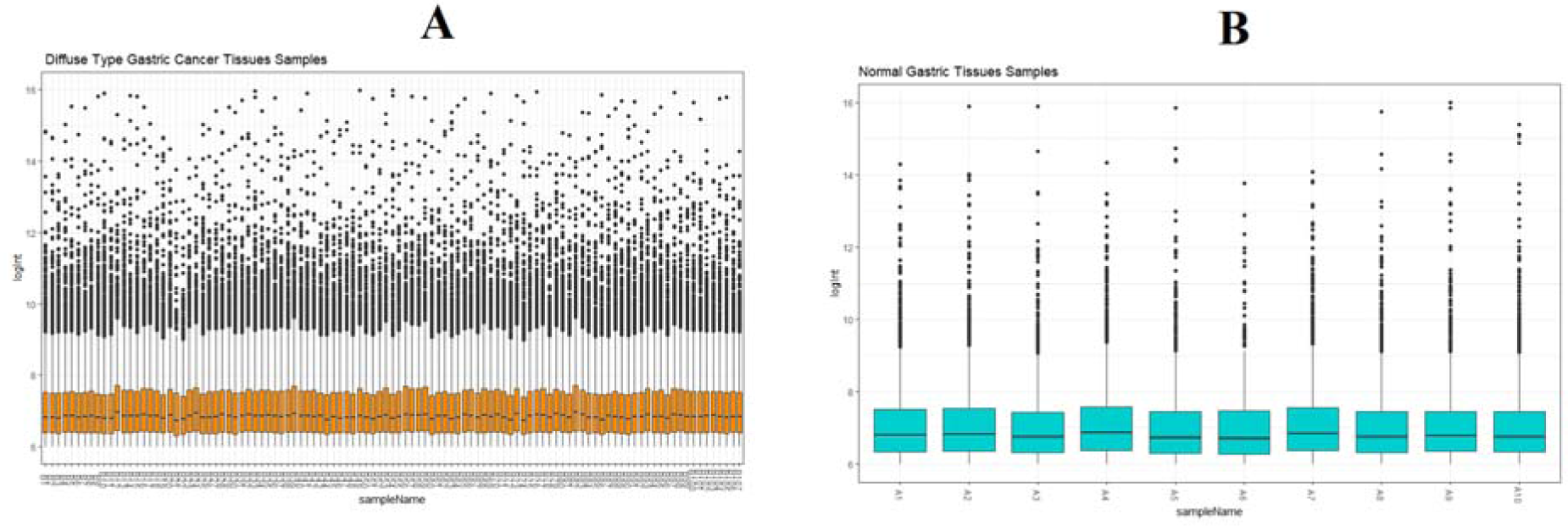
Box plots of the normalized RNA-seq data. (A) 107 GC samples (B) 10 normal gastric tissue samples. Horizontal axis represents the sample symbol and the vertical axis represents the gene expression values. The black line in the box plot represents the median value of gene expression. (A1 – A10 = Normal gastric tissue samples (blue color box); B1 – B107 = GC samples (brown color box))

**Fig. 2.**
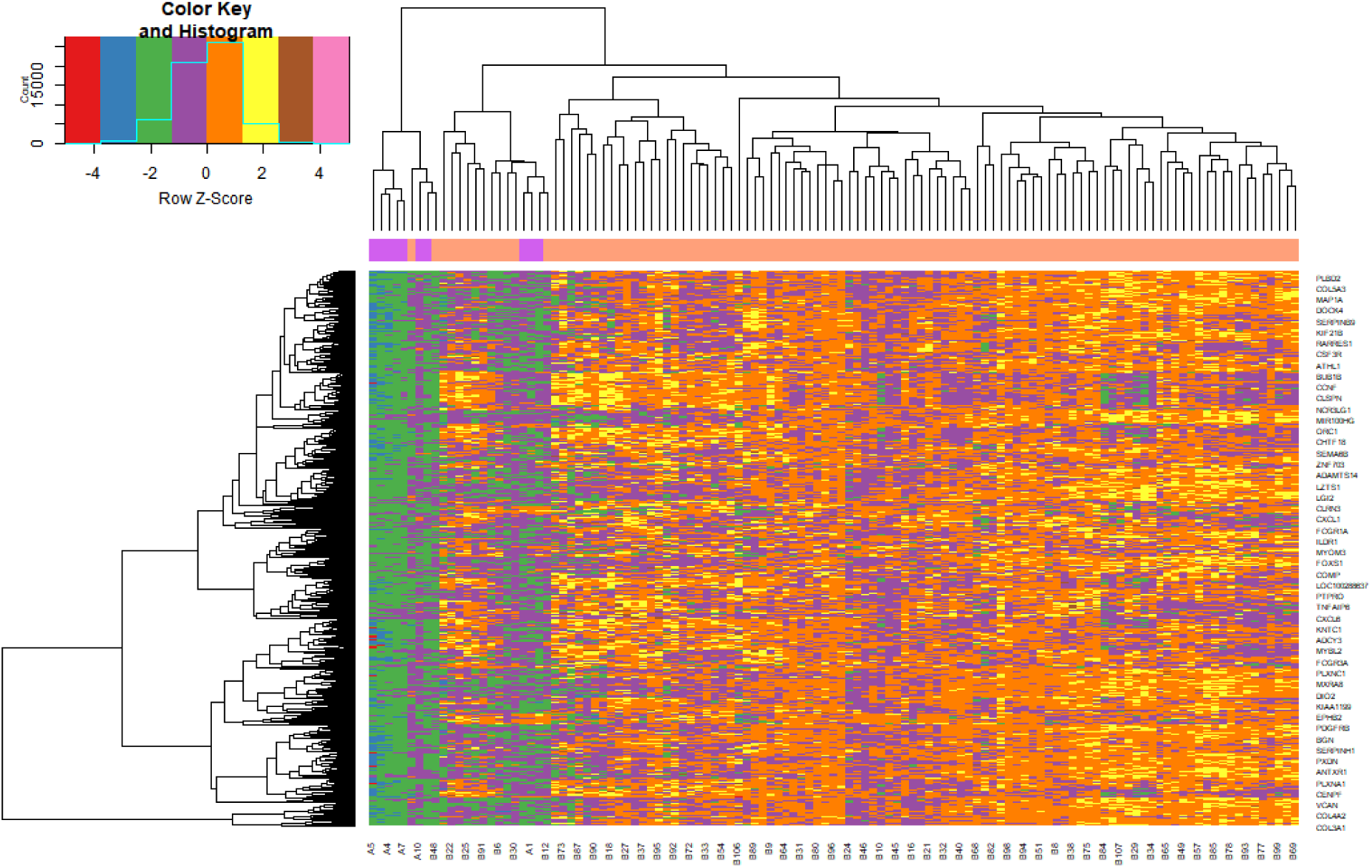
Heat map of up regulated differentially expressed genes. Legend on the top left indicate log fold change of genes. (A1 – A10 = Normal gastric tissue samples; B1 – B107 = GC samples)

**Fig. 3.**
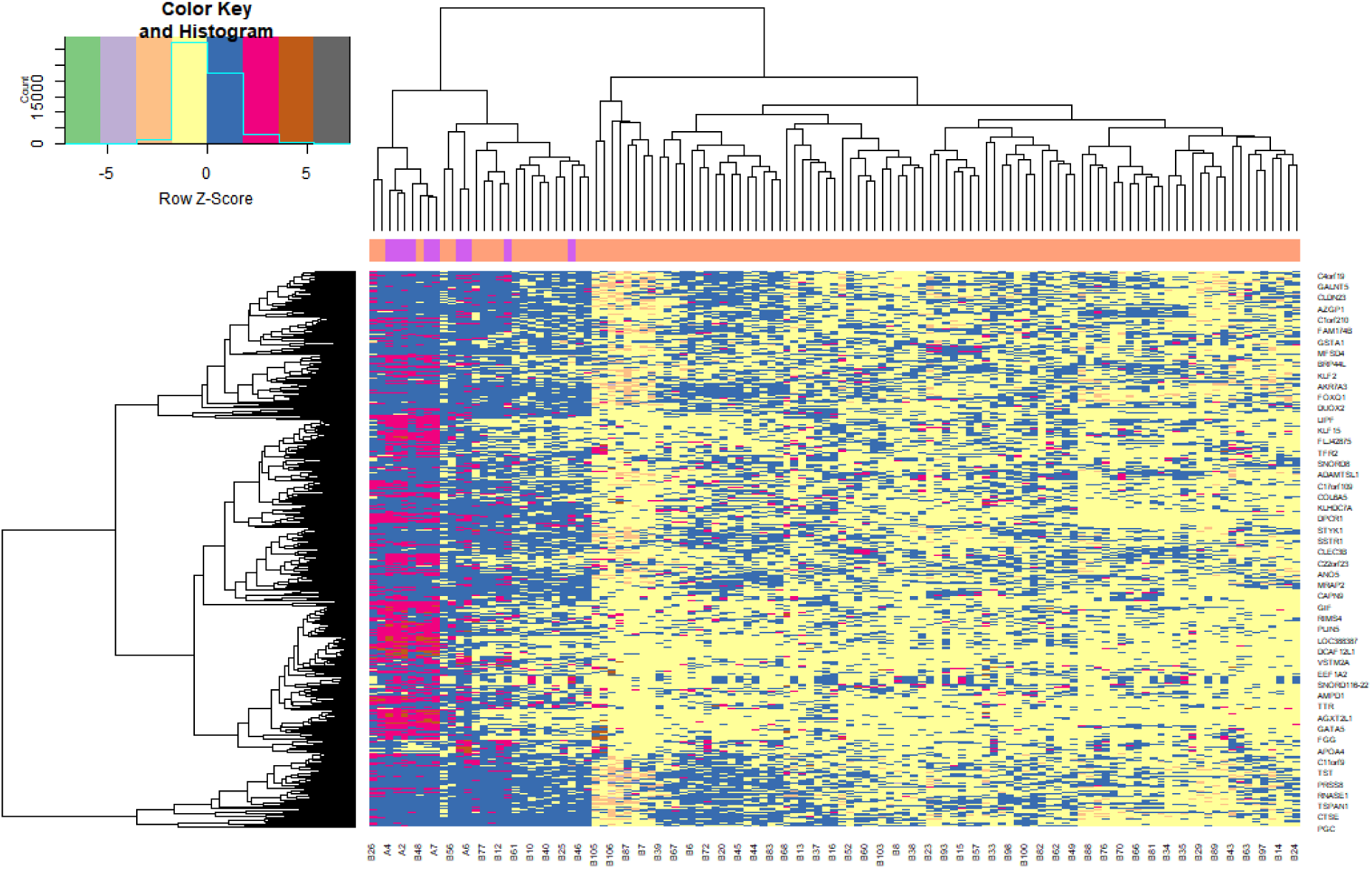
Heat map of down regulated differentially expressed genes. Legend on the top left indicate log fold change of genes. (A1 – A10 = Normal gastric tissue samples; B1 – B107 = GC samples)

**Fig. 4.**
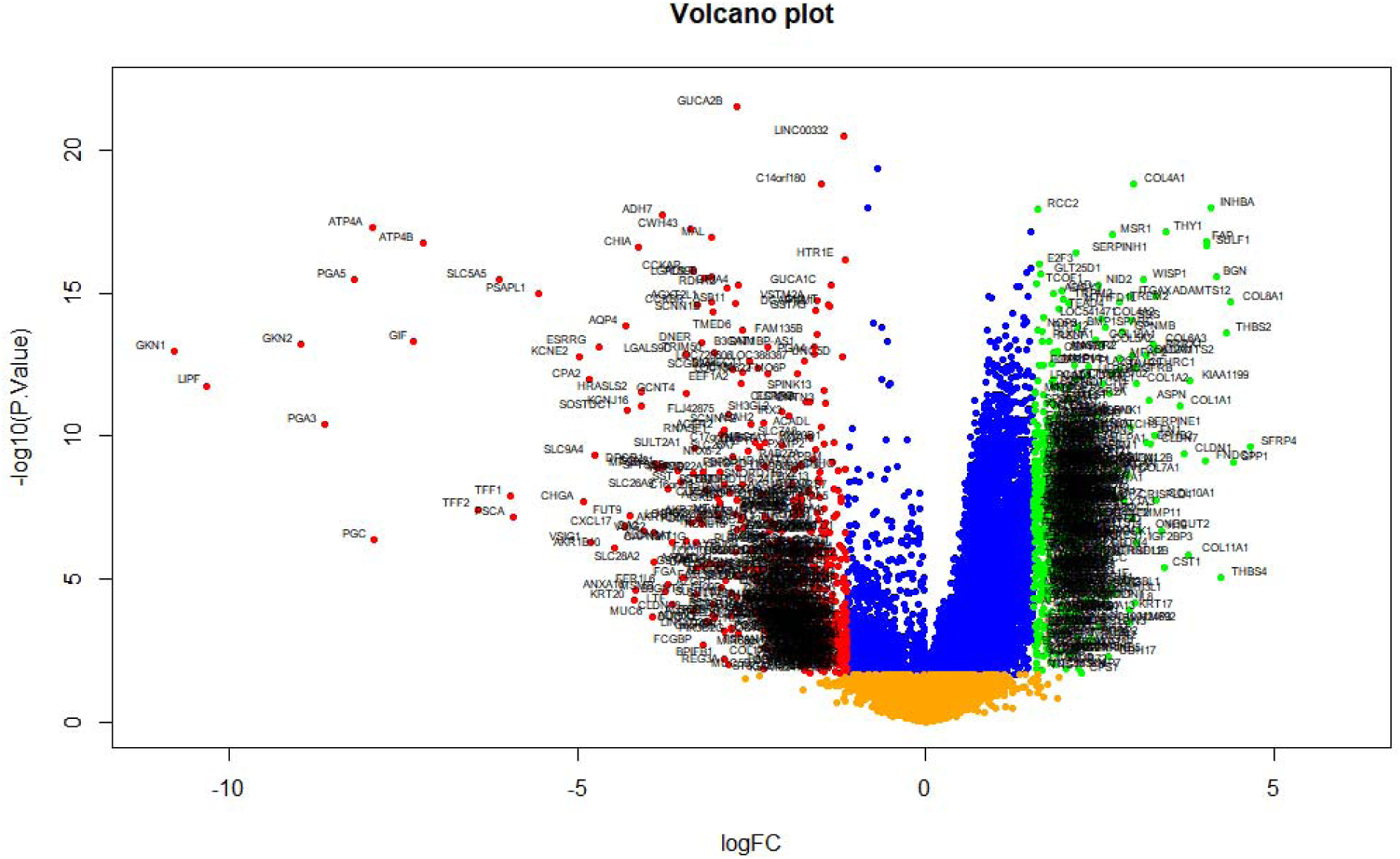
Volcano plot of differentially expressed genes. Genes with a significant change of more than two-fold were selected.

**Table 2.**
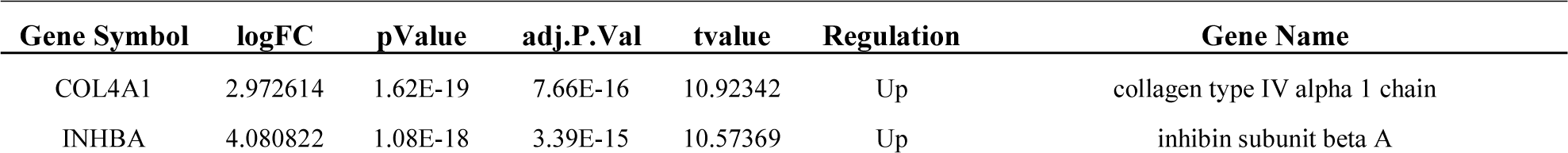
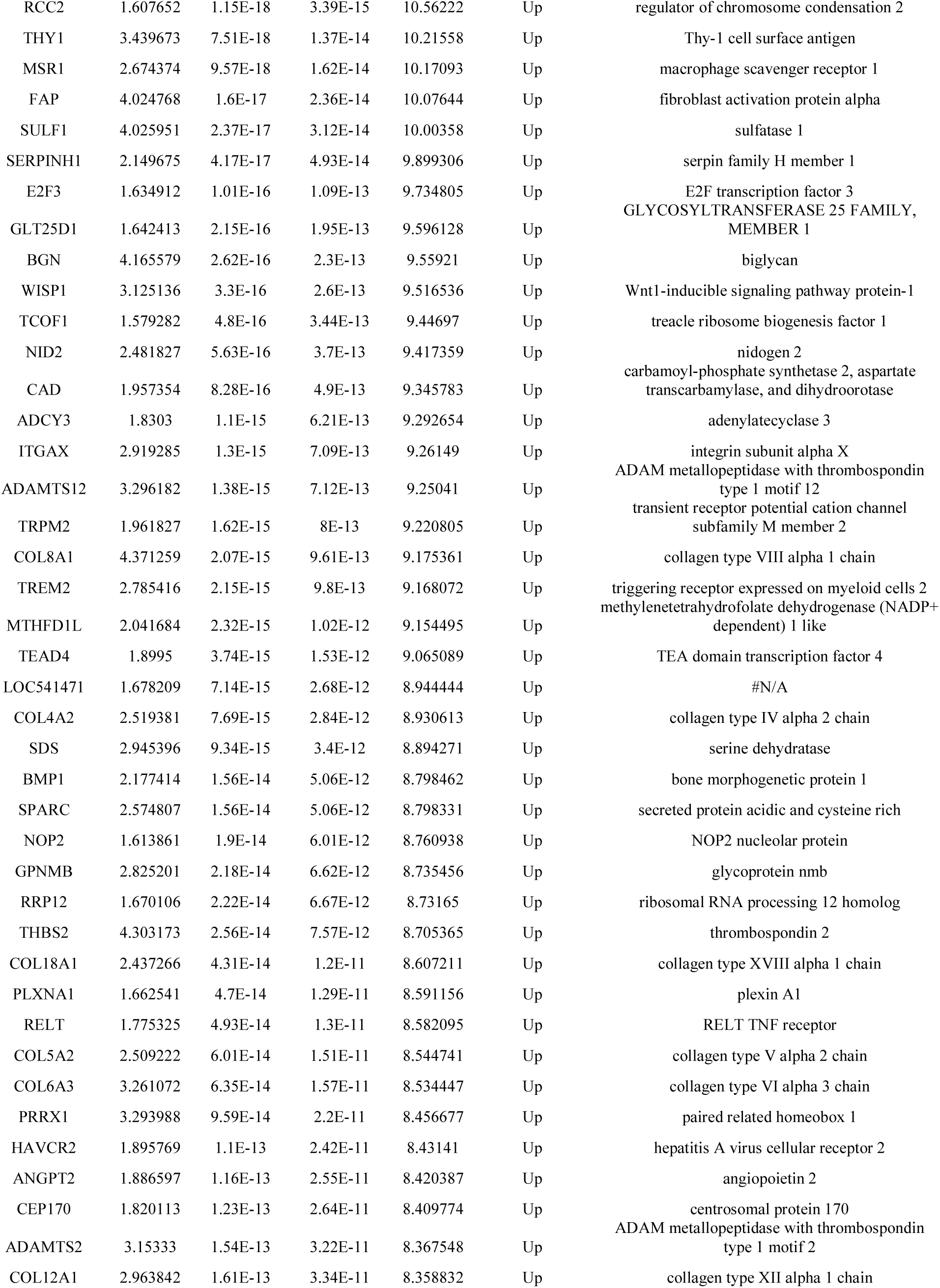
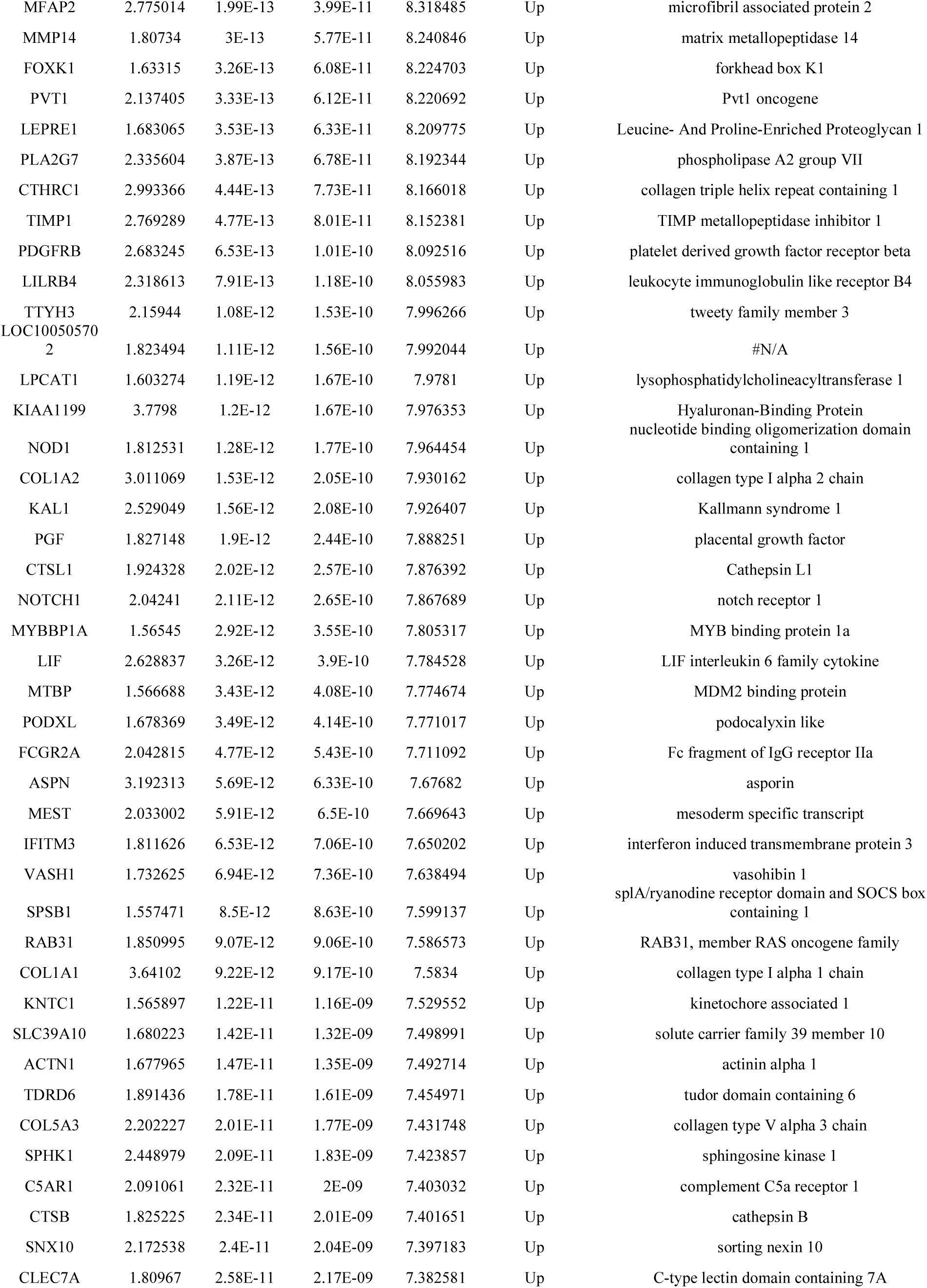
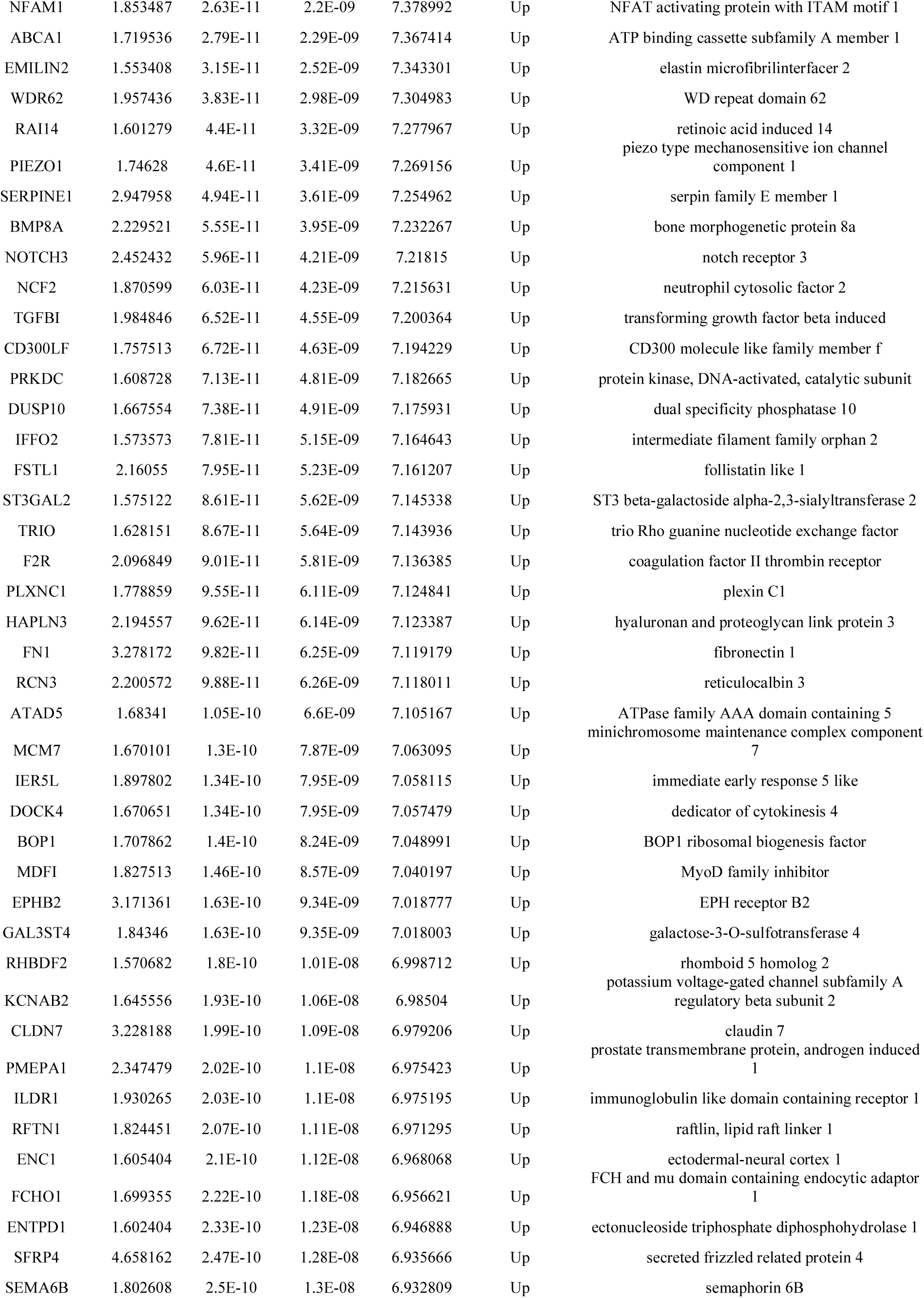
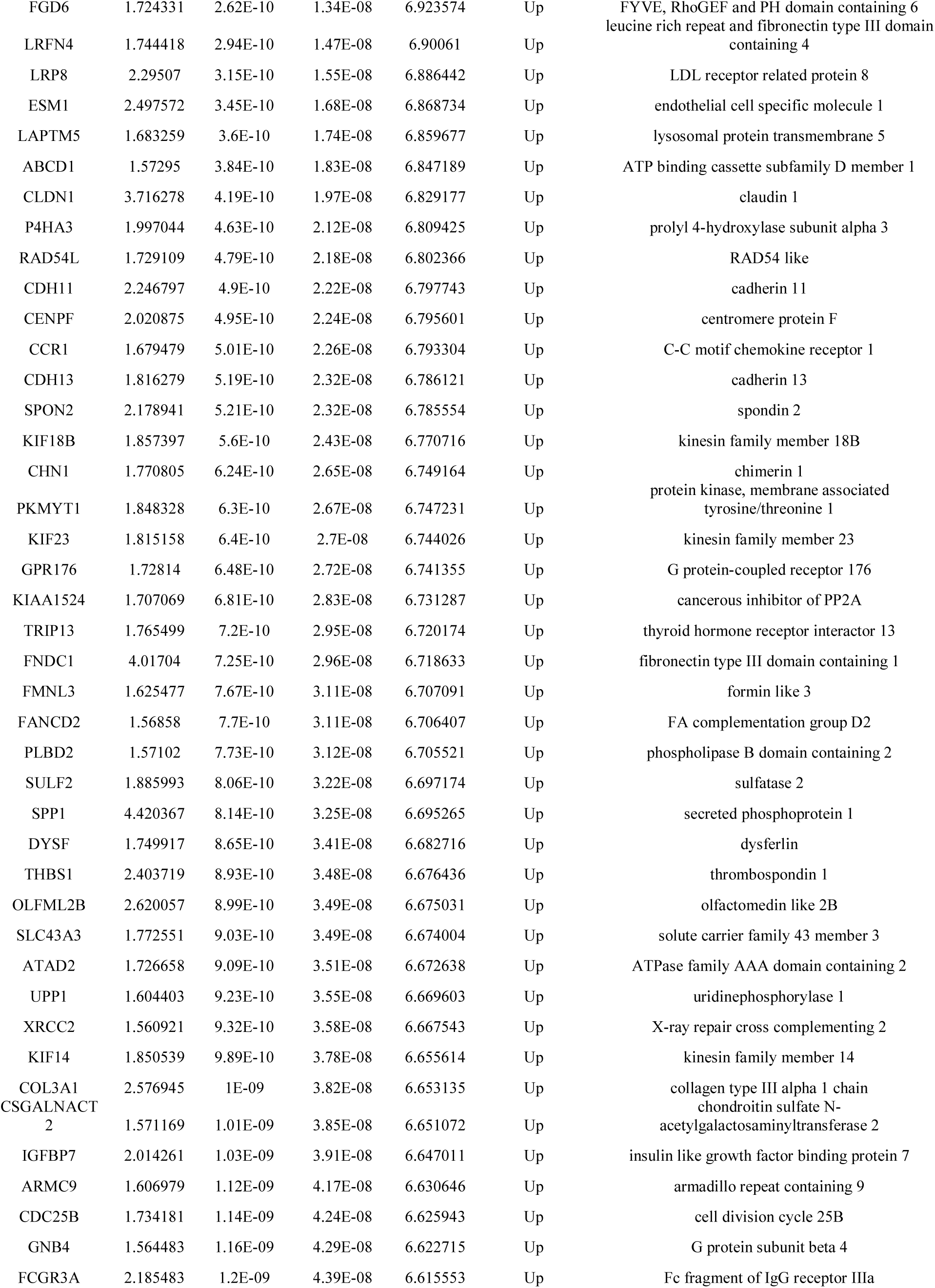
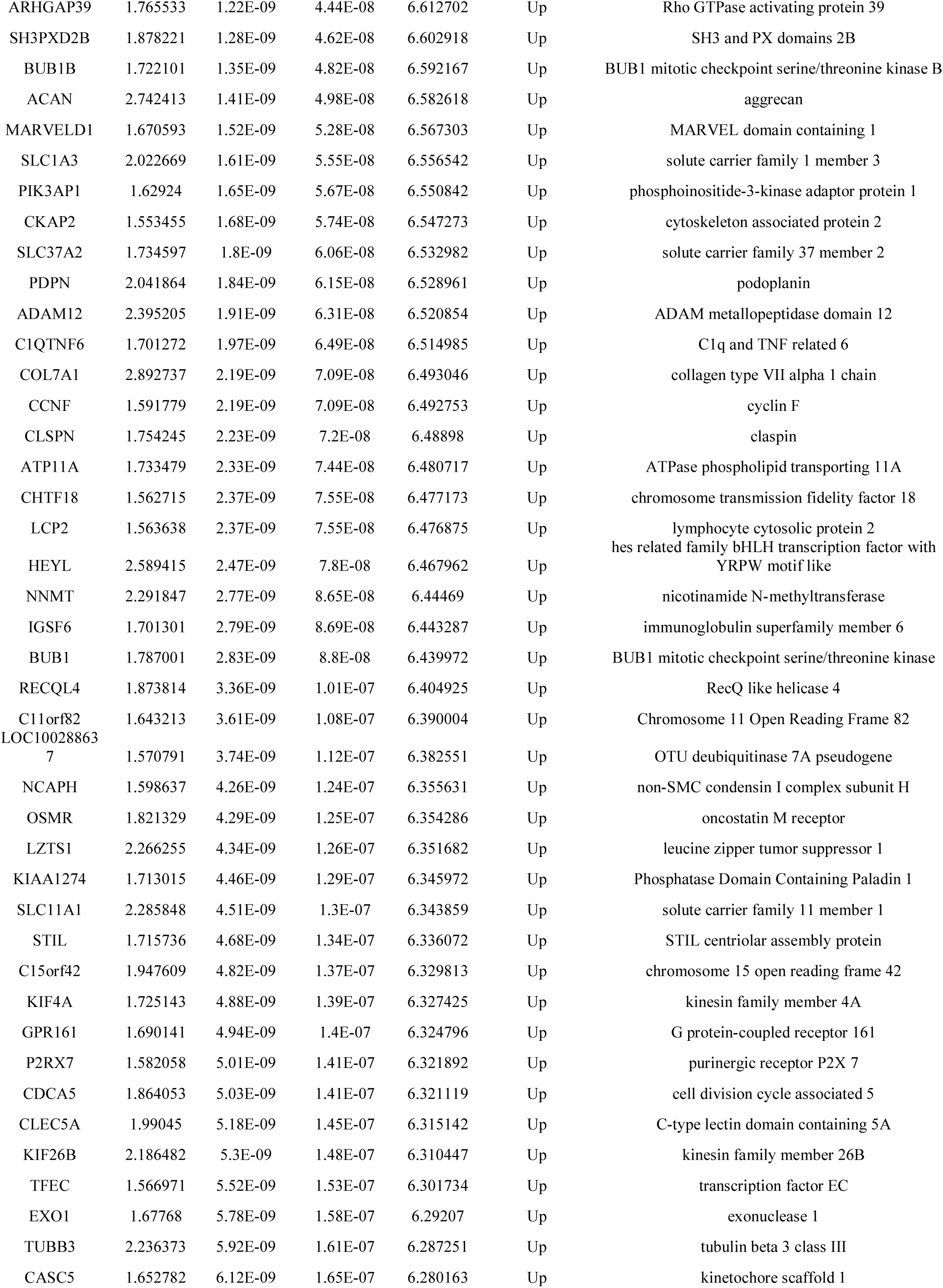
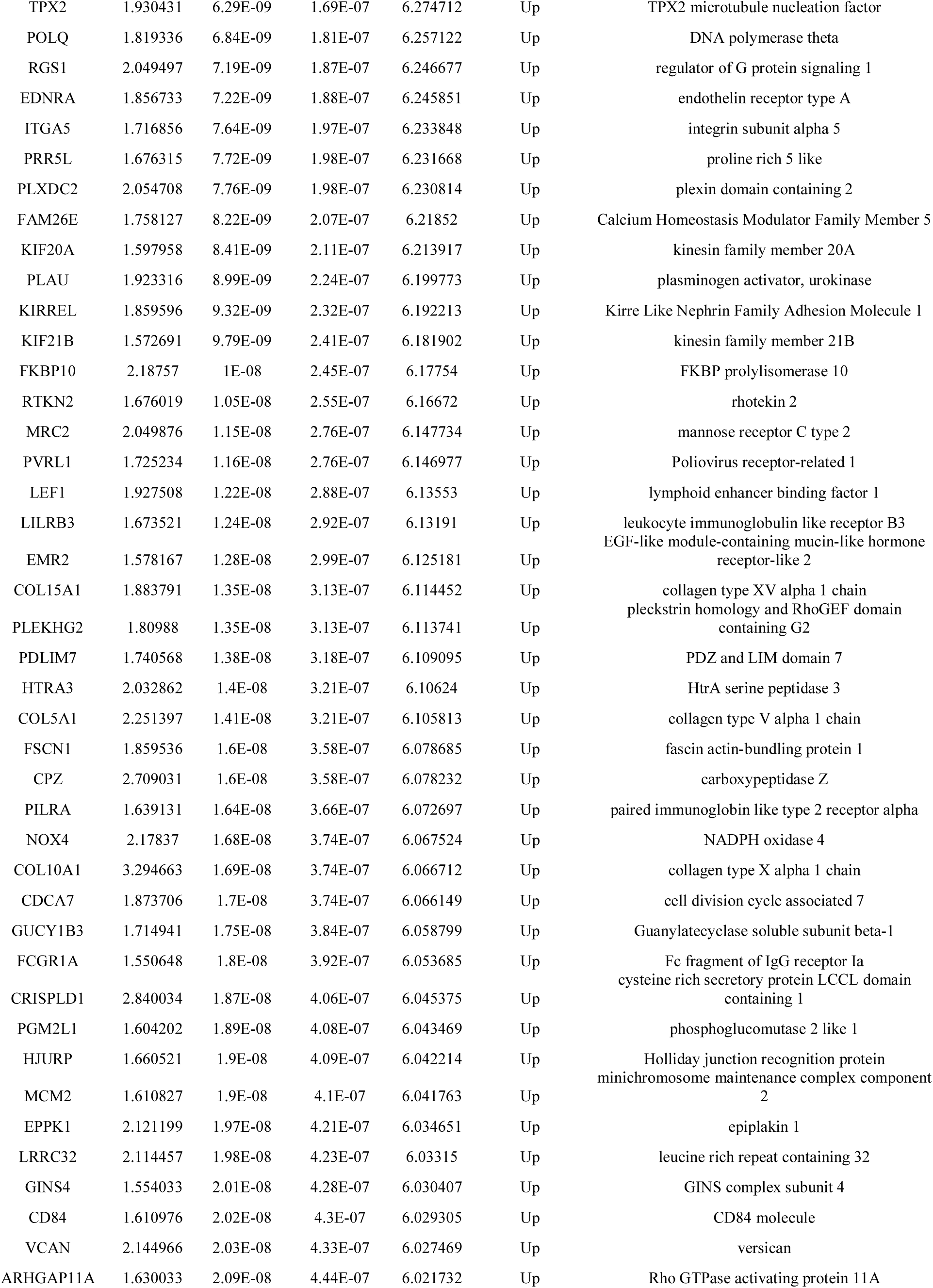
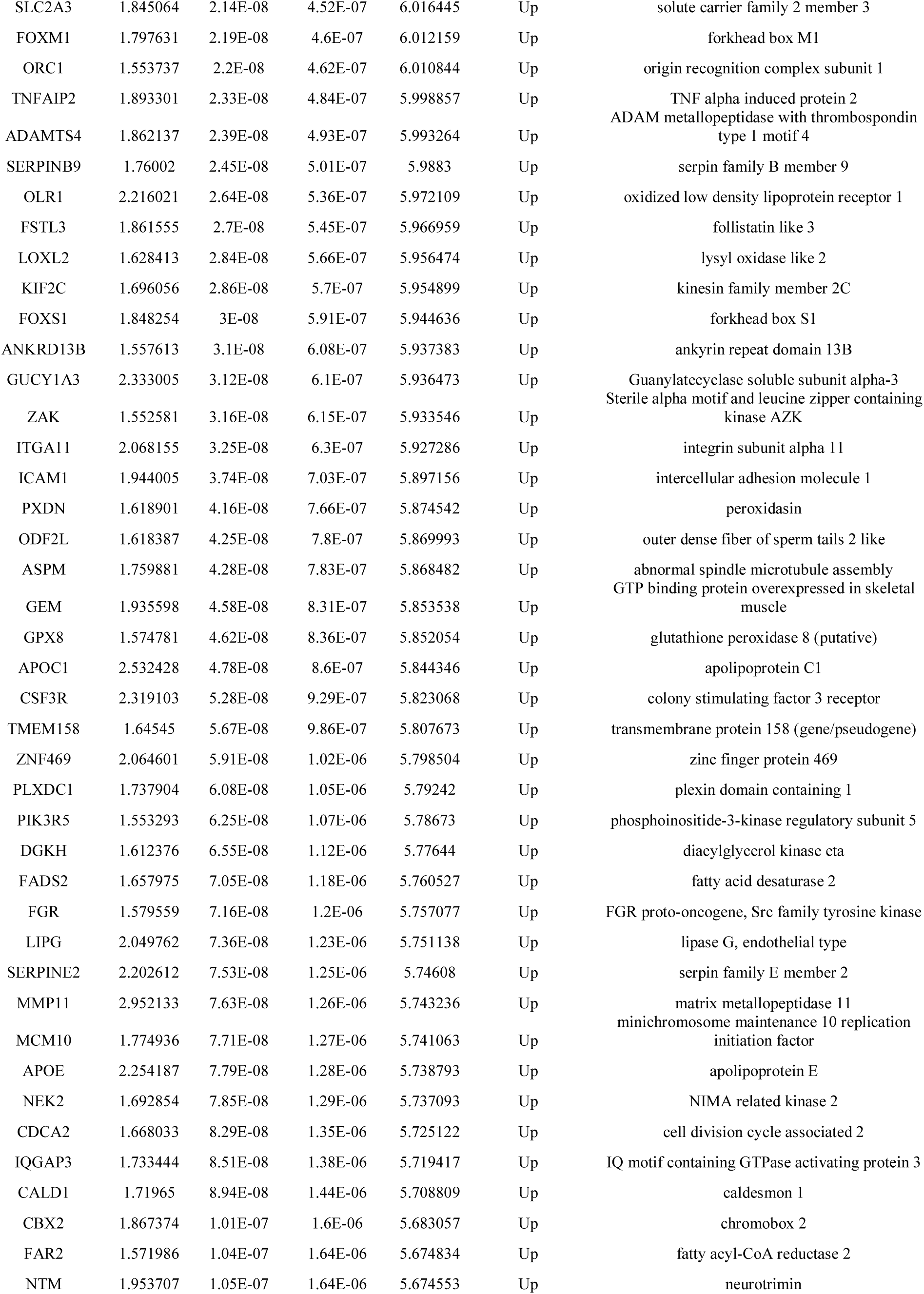
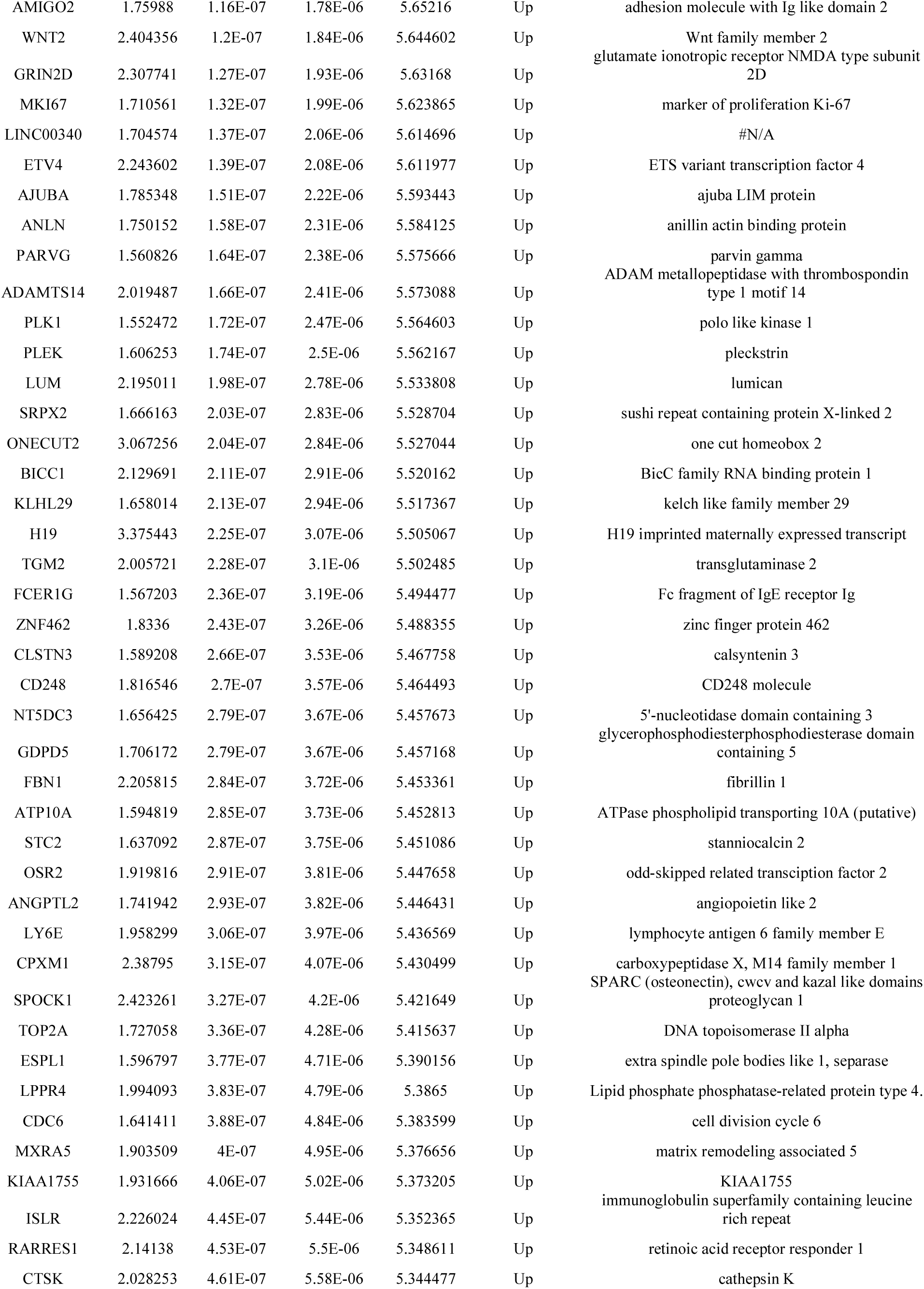
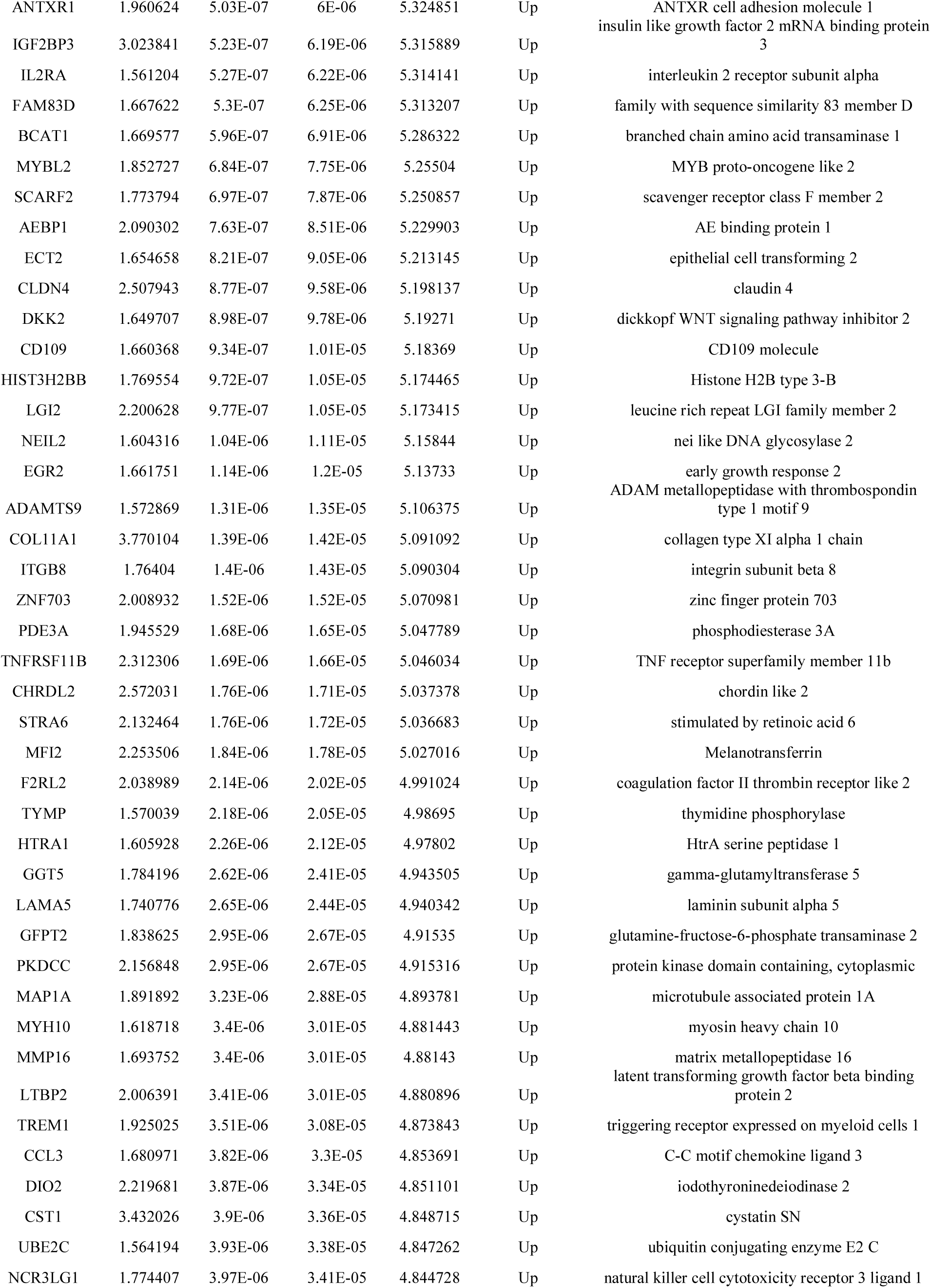
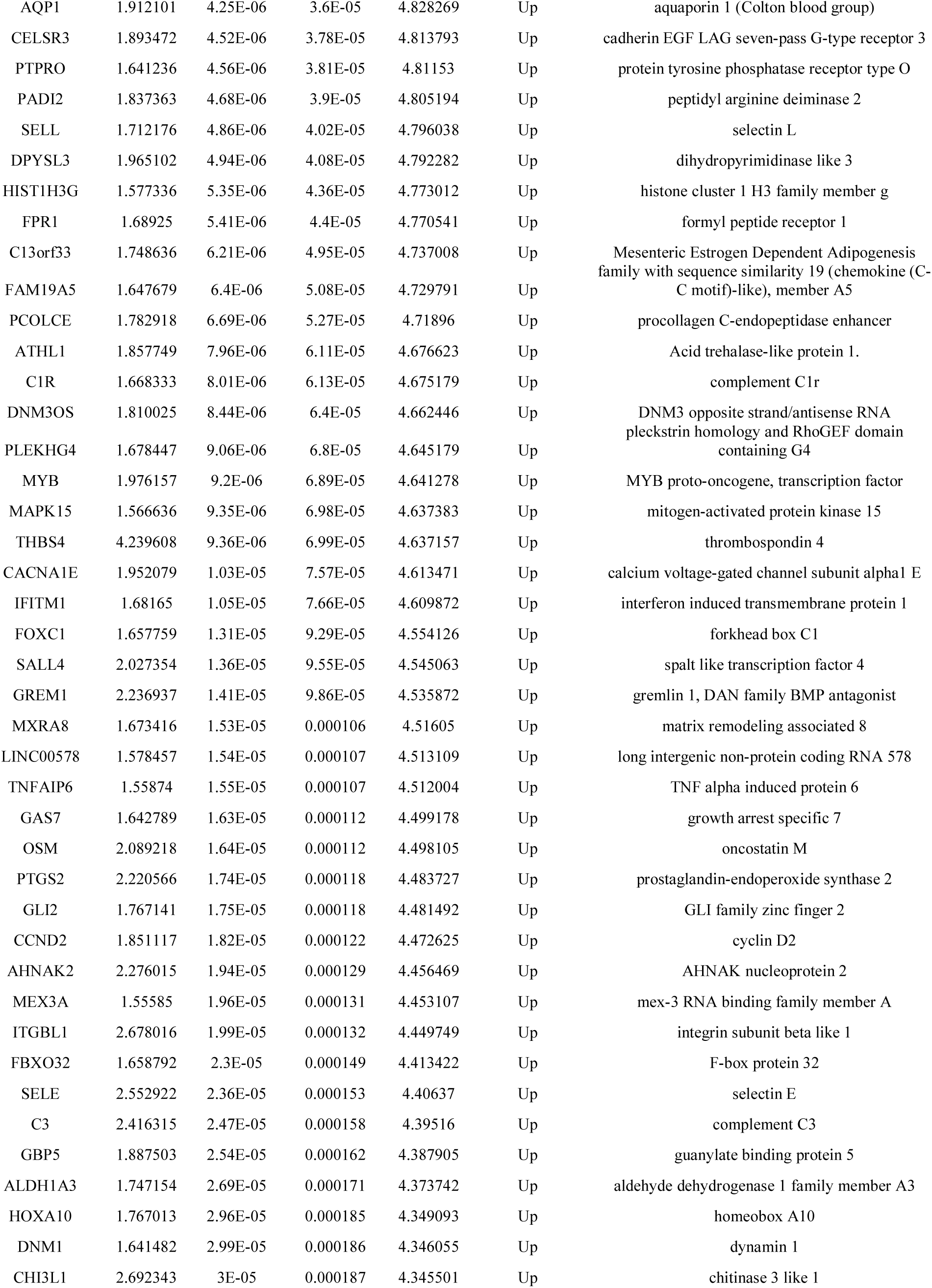
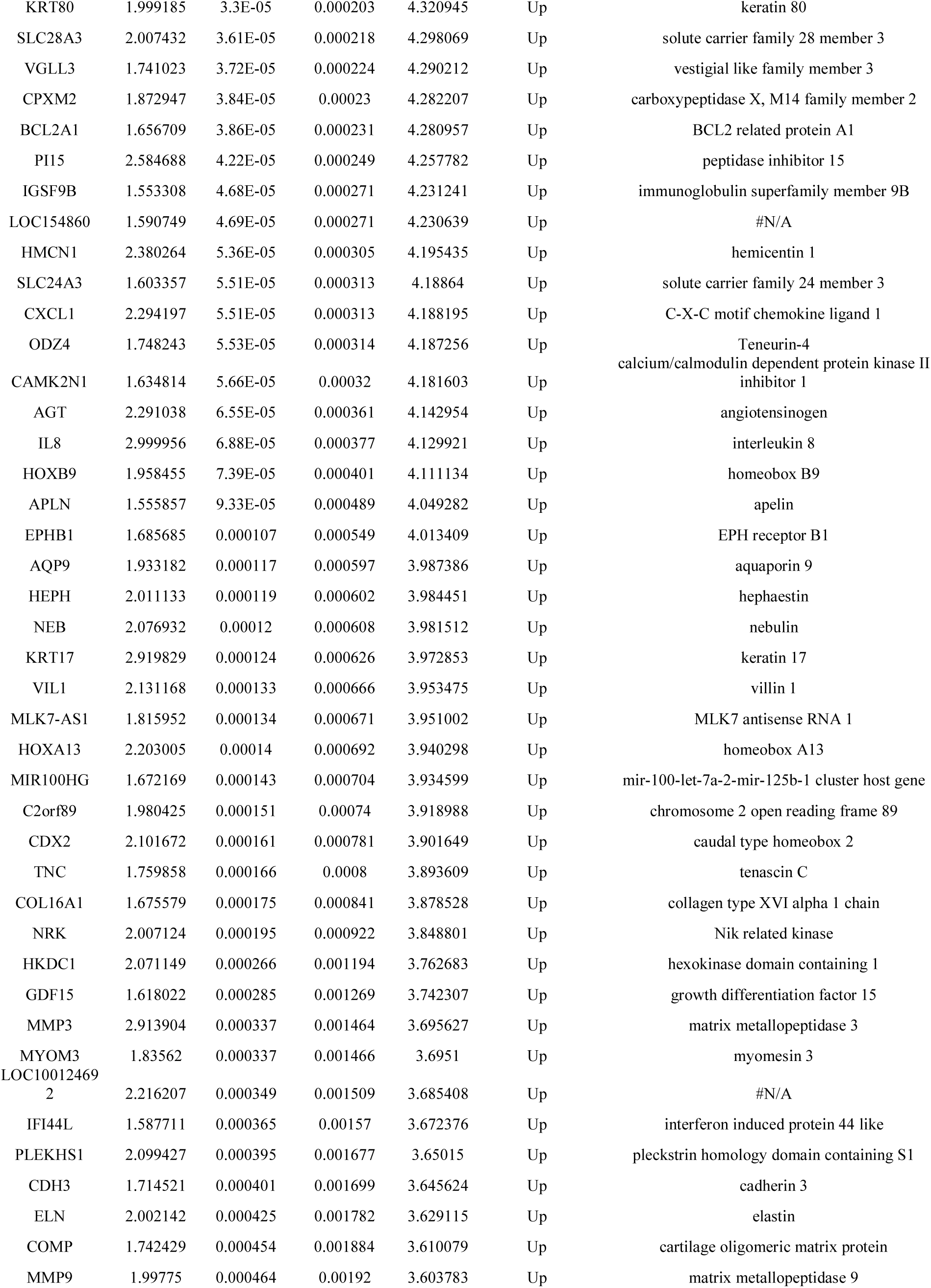
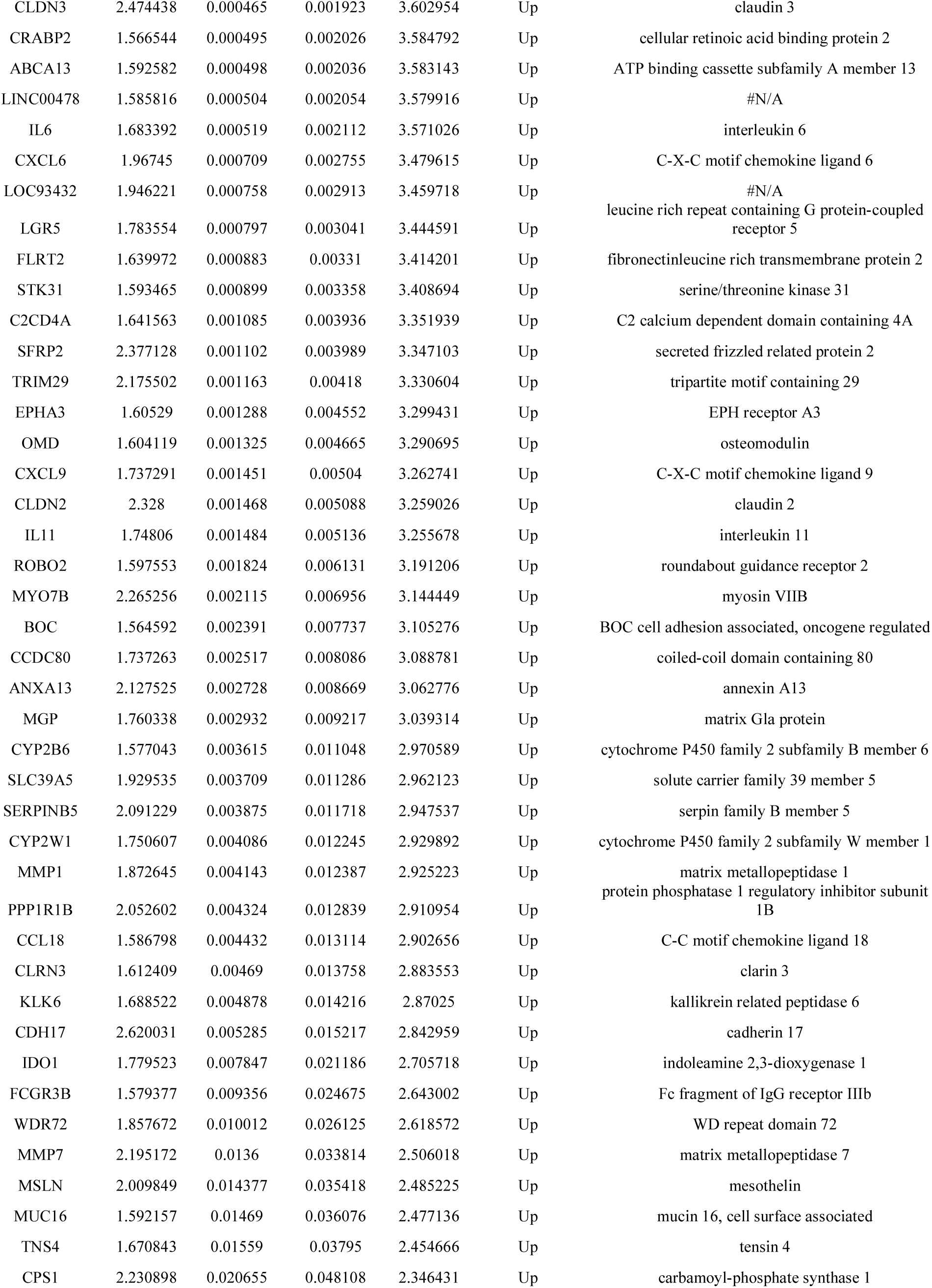
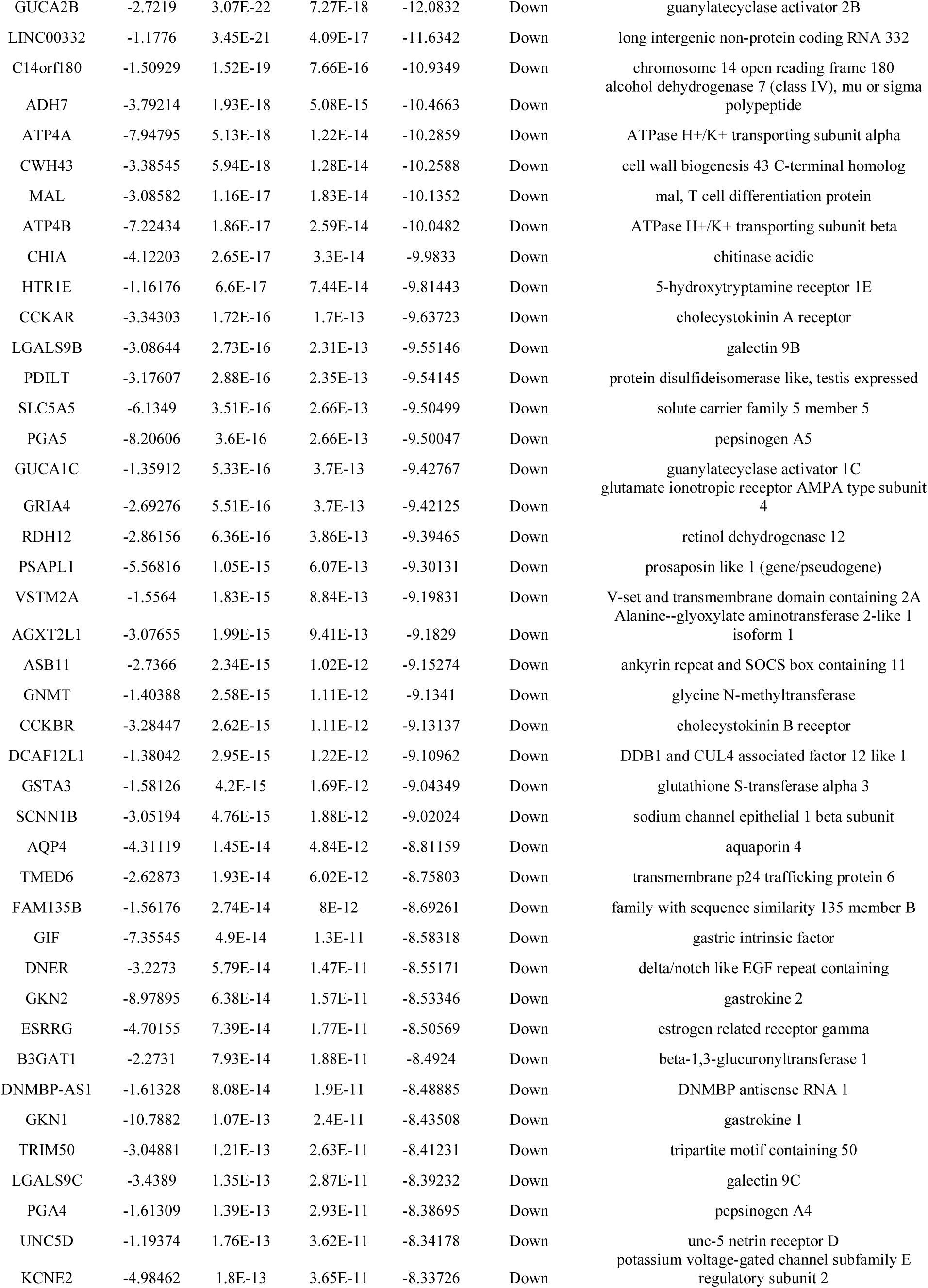
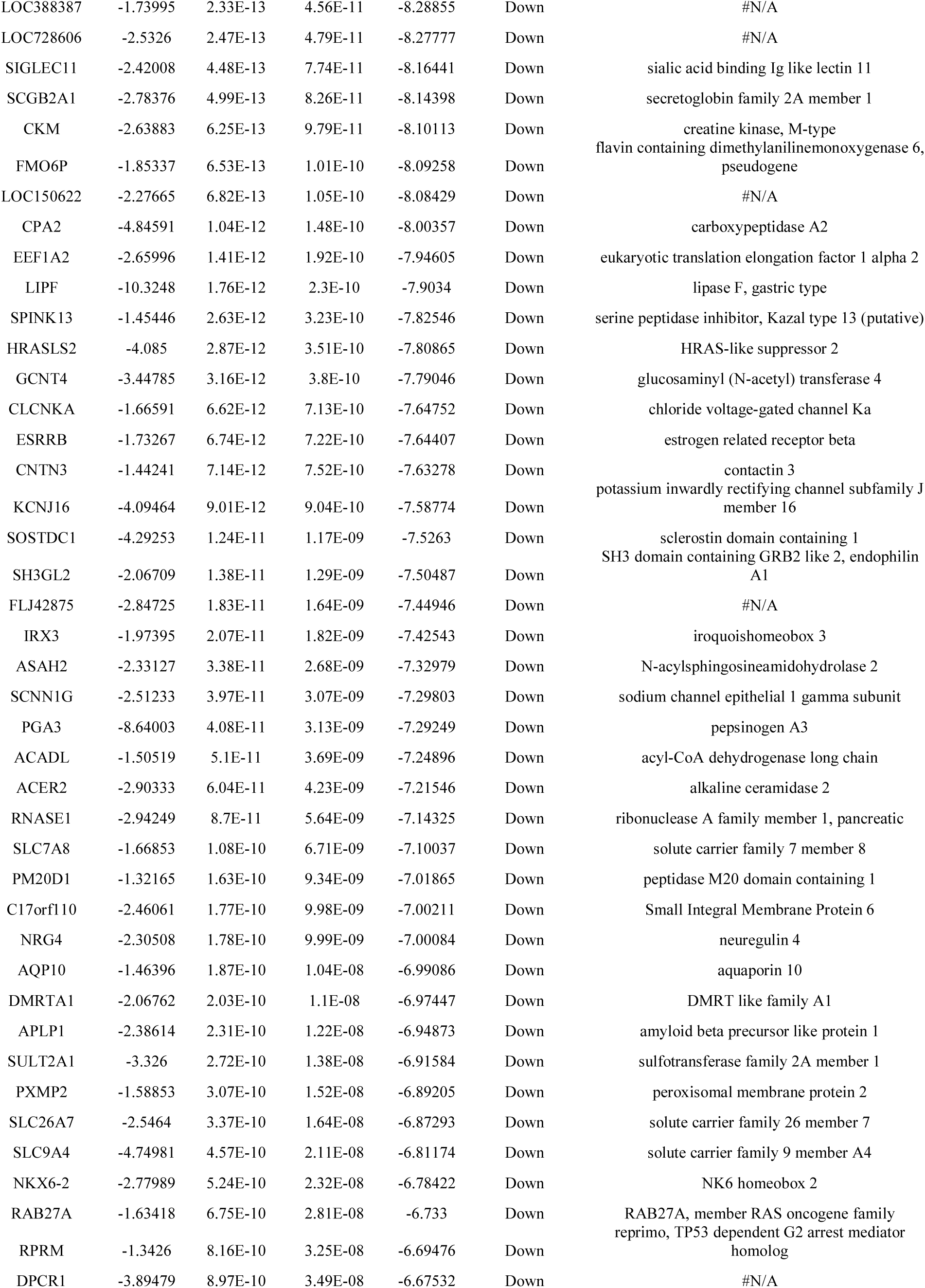
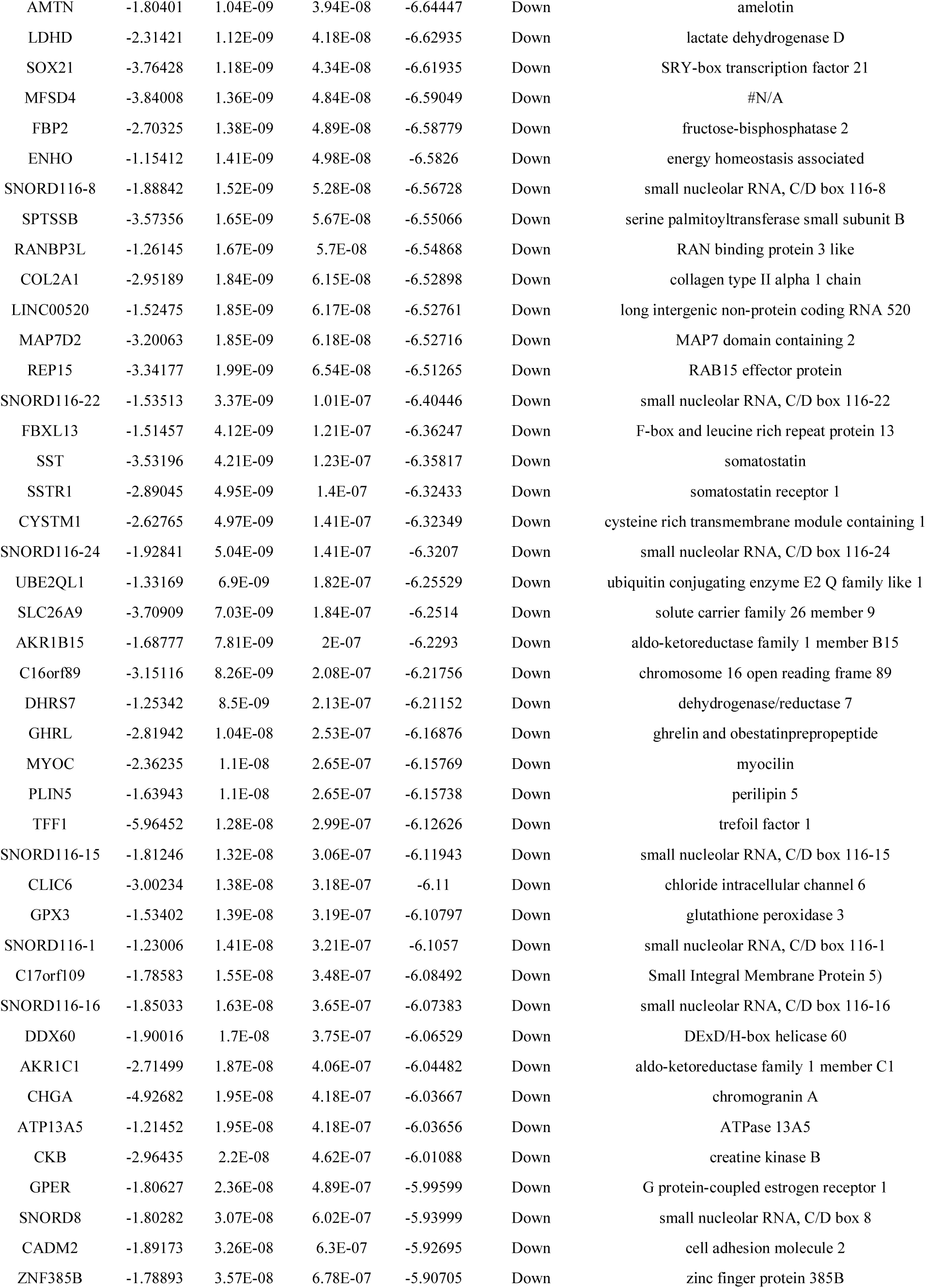
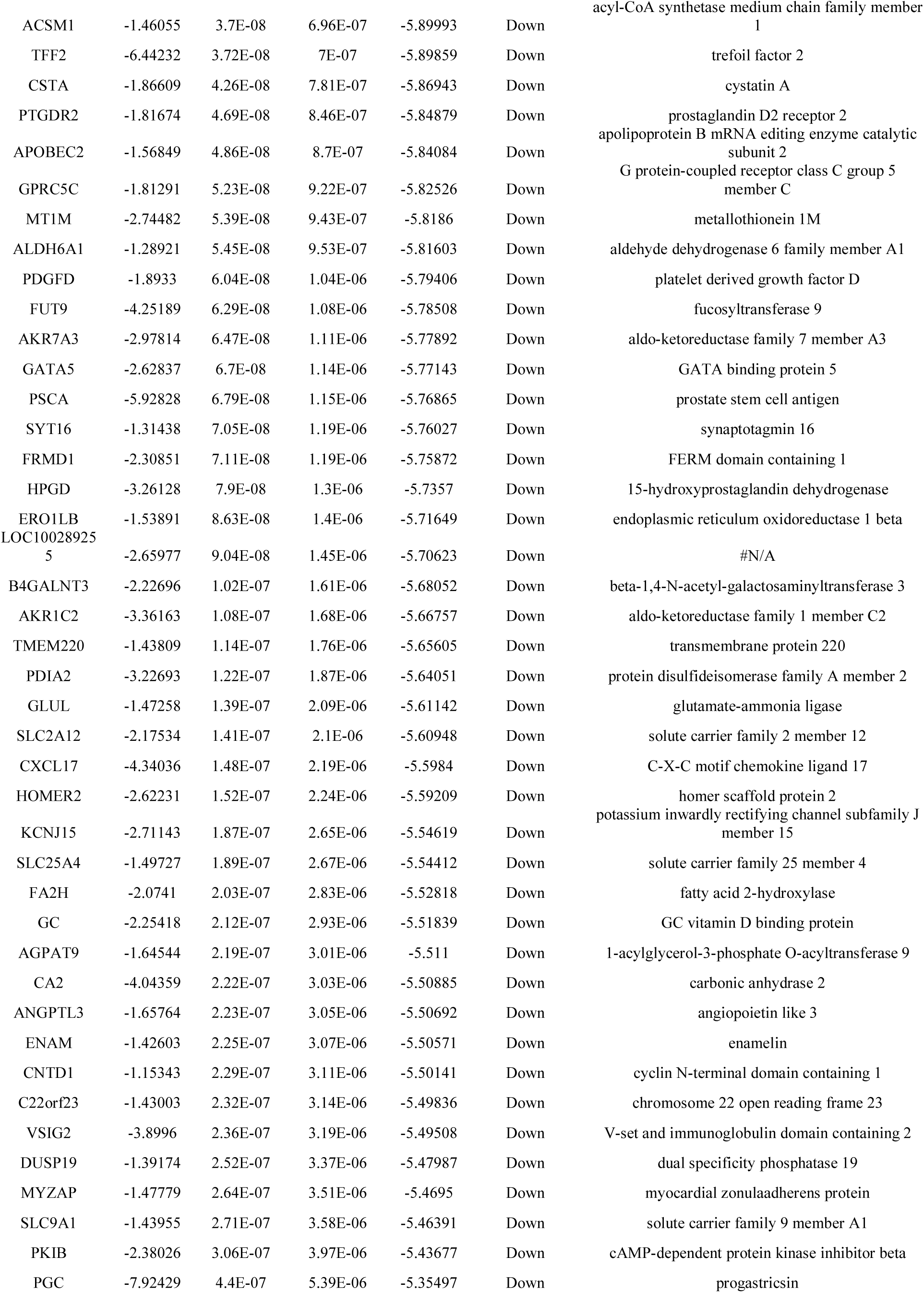
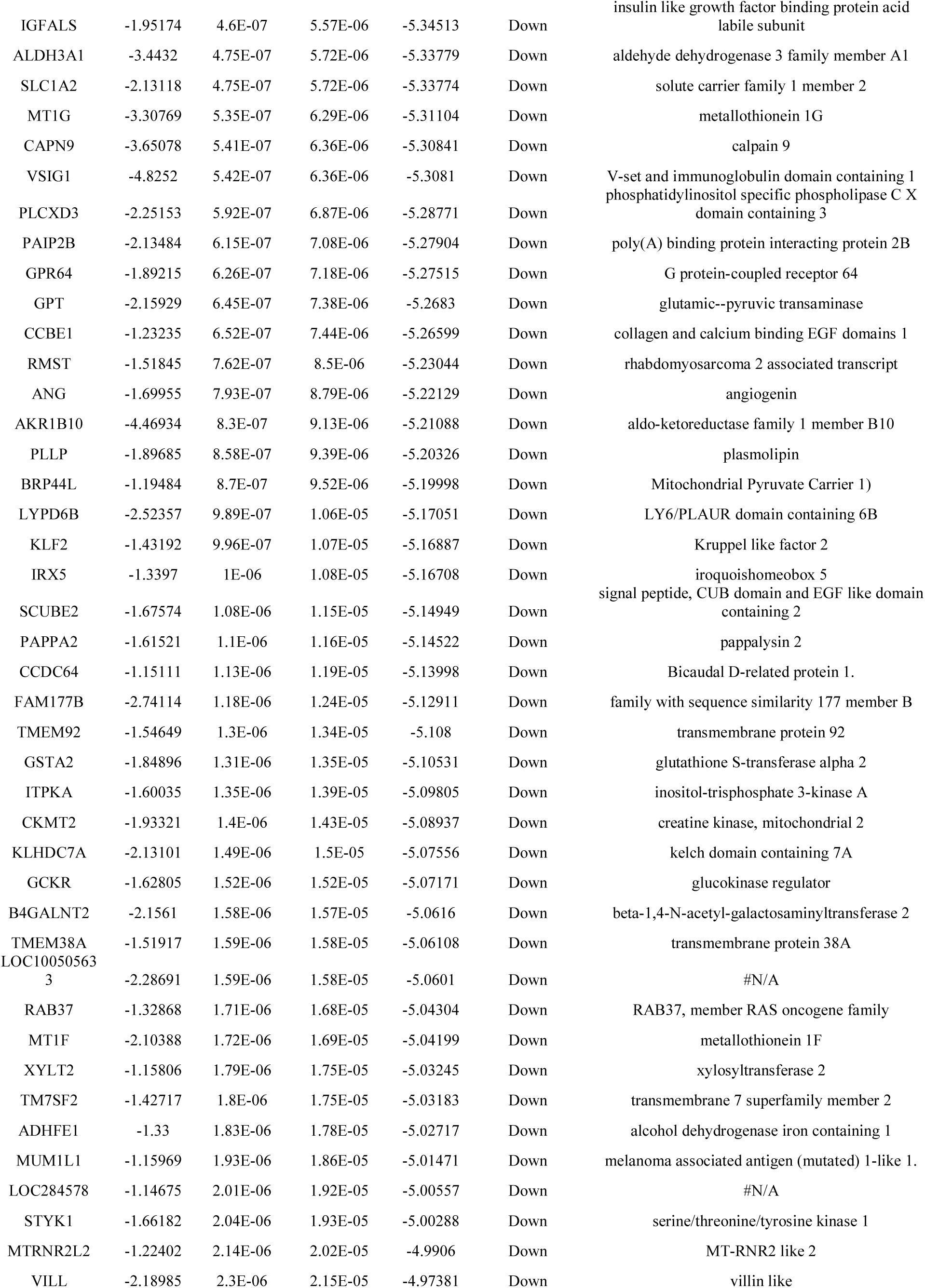
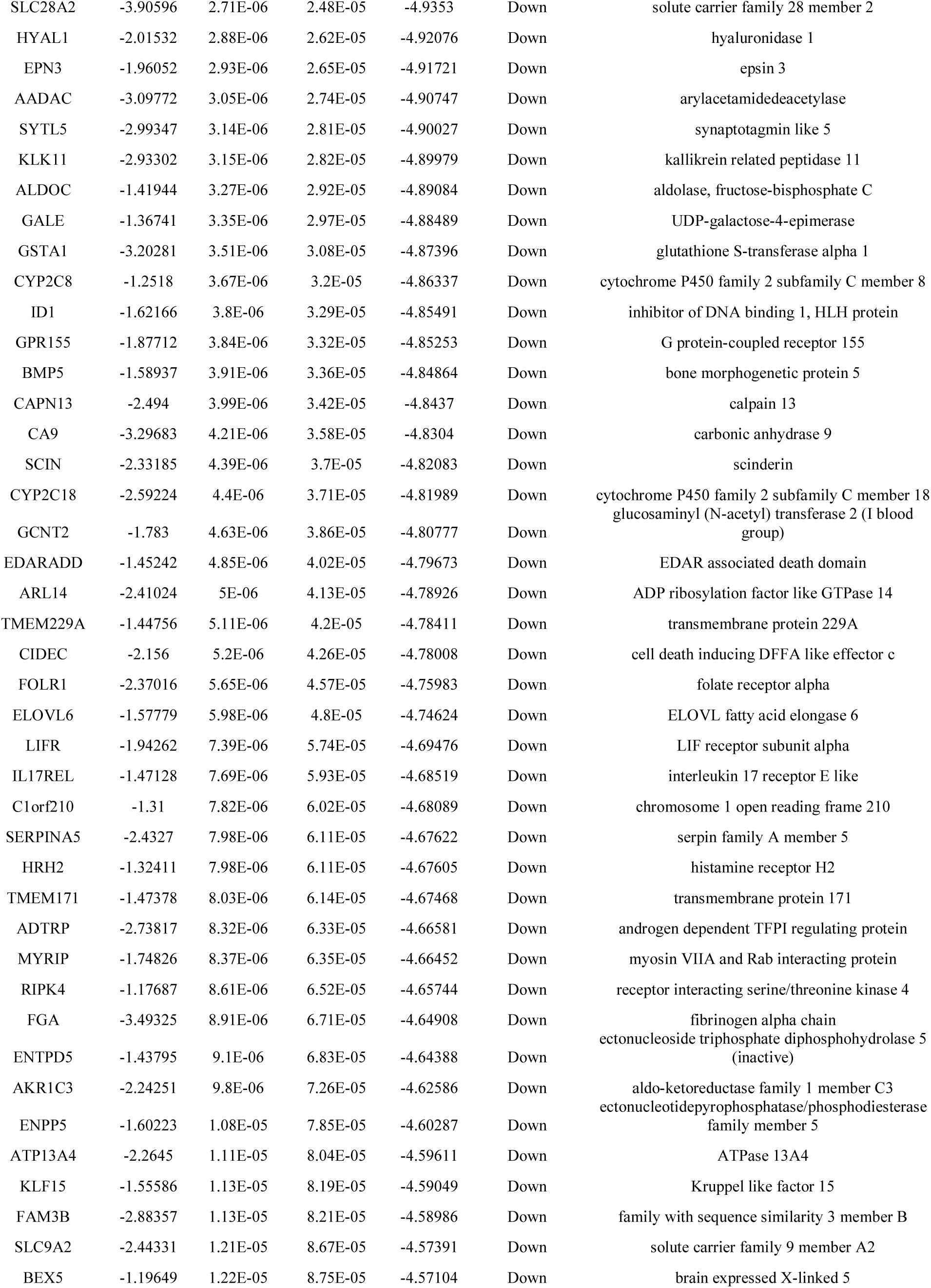
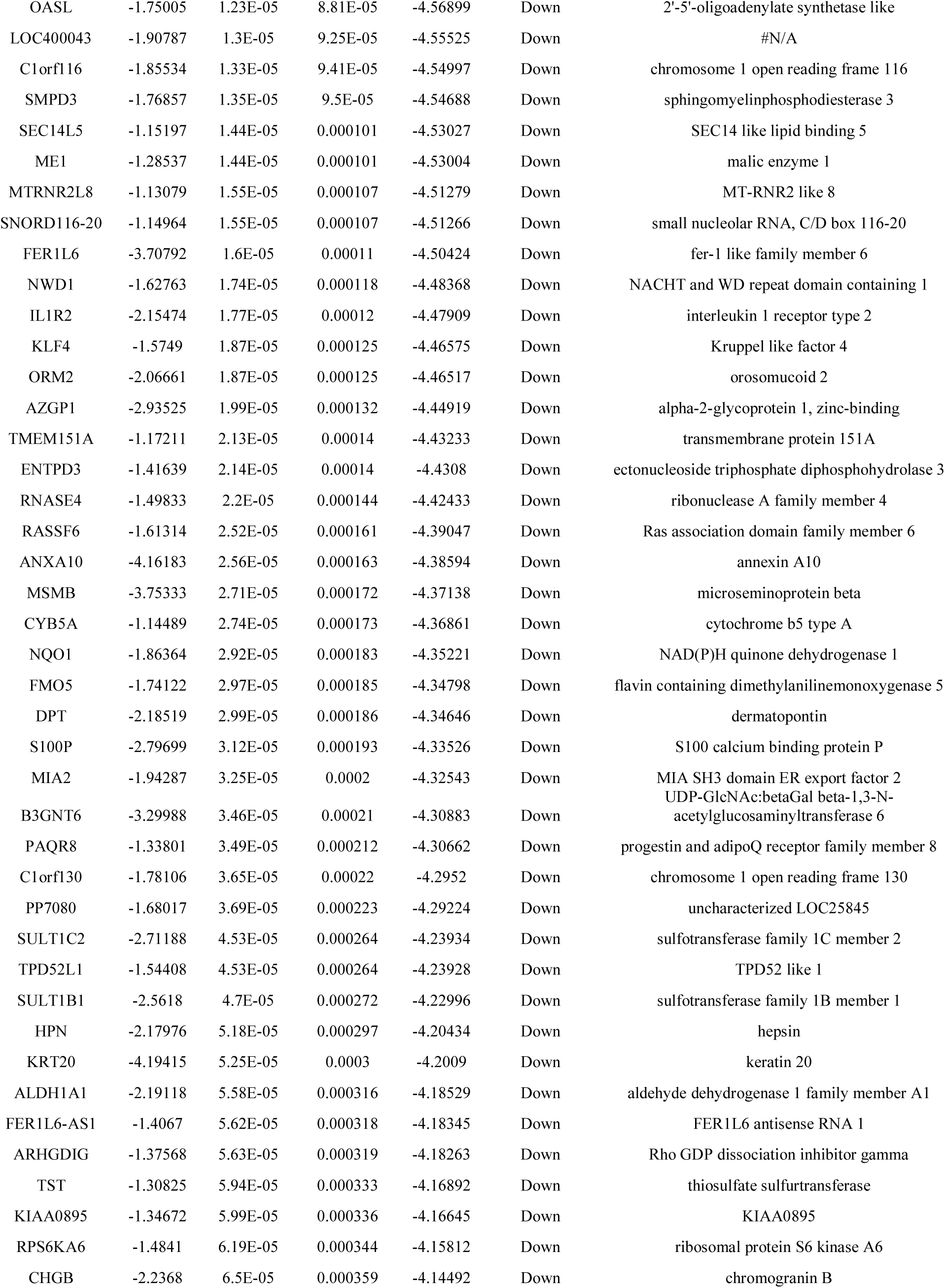
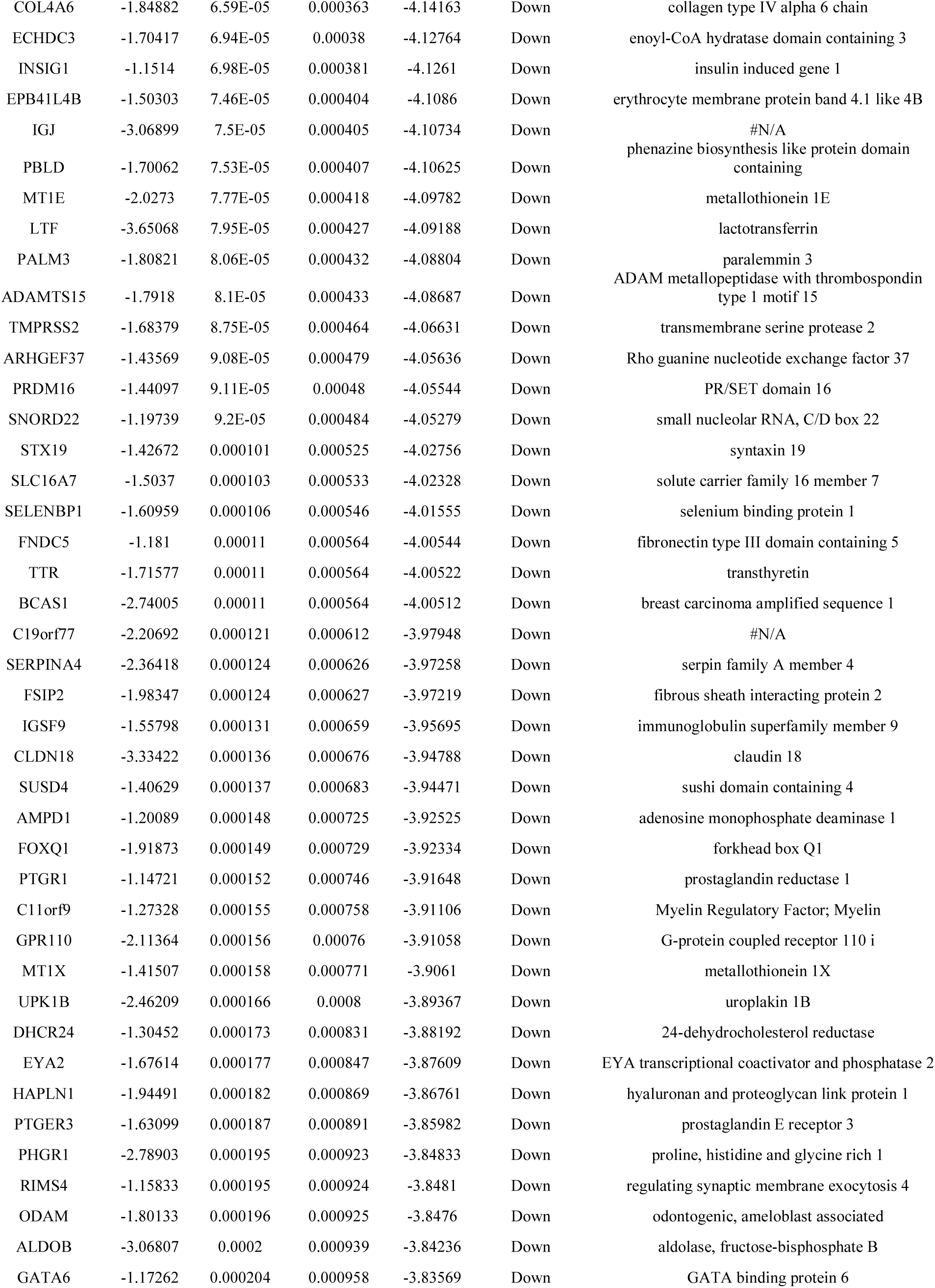
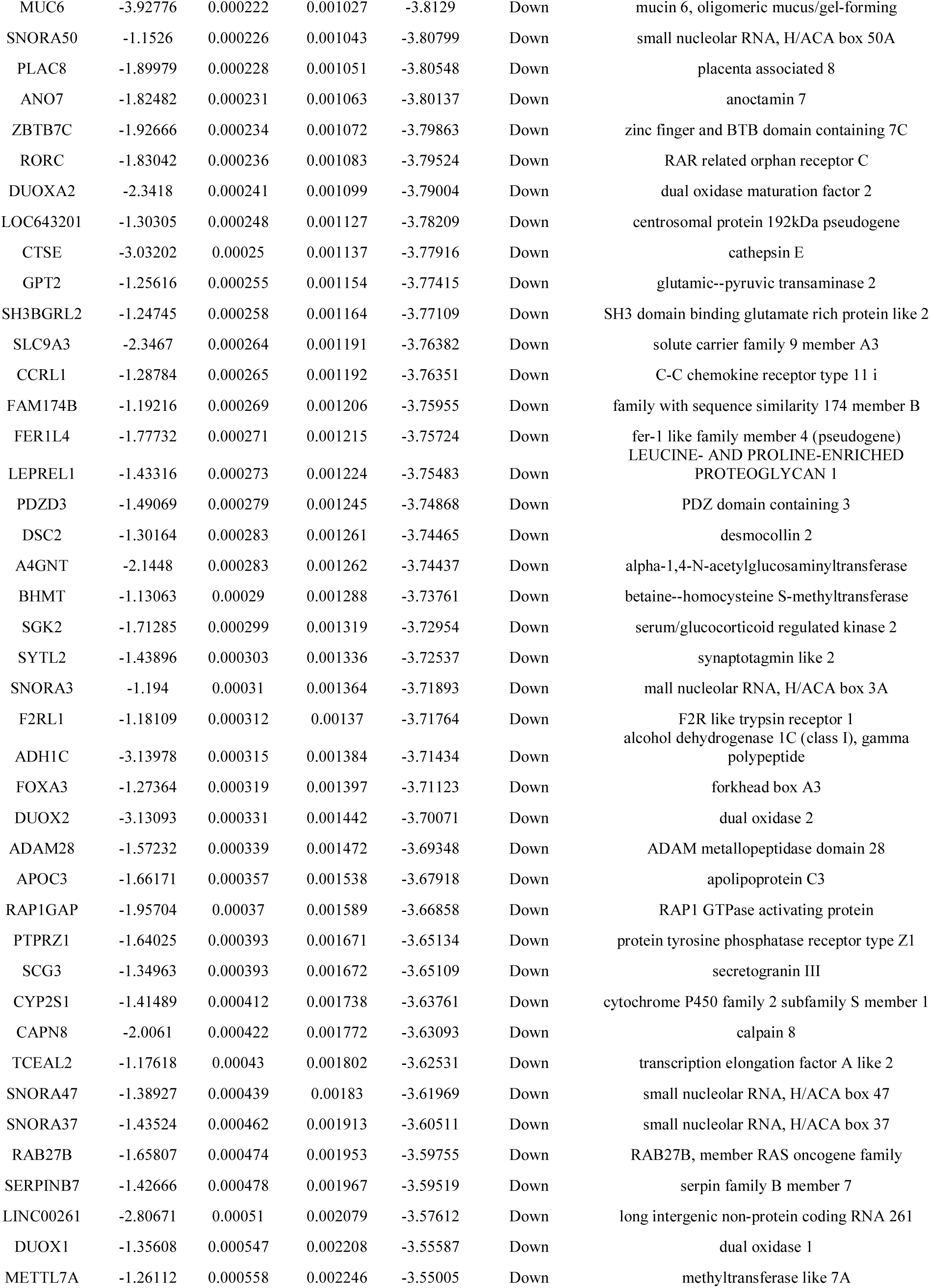
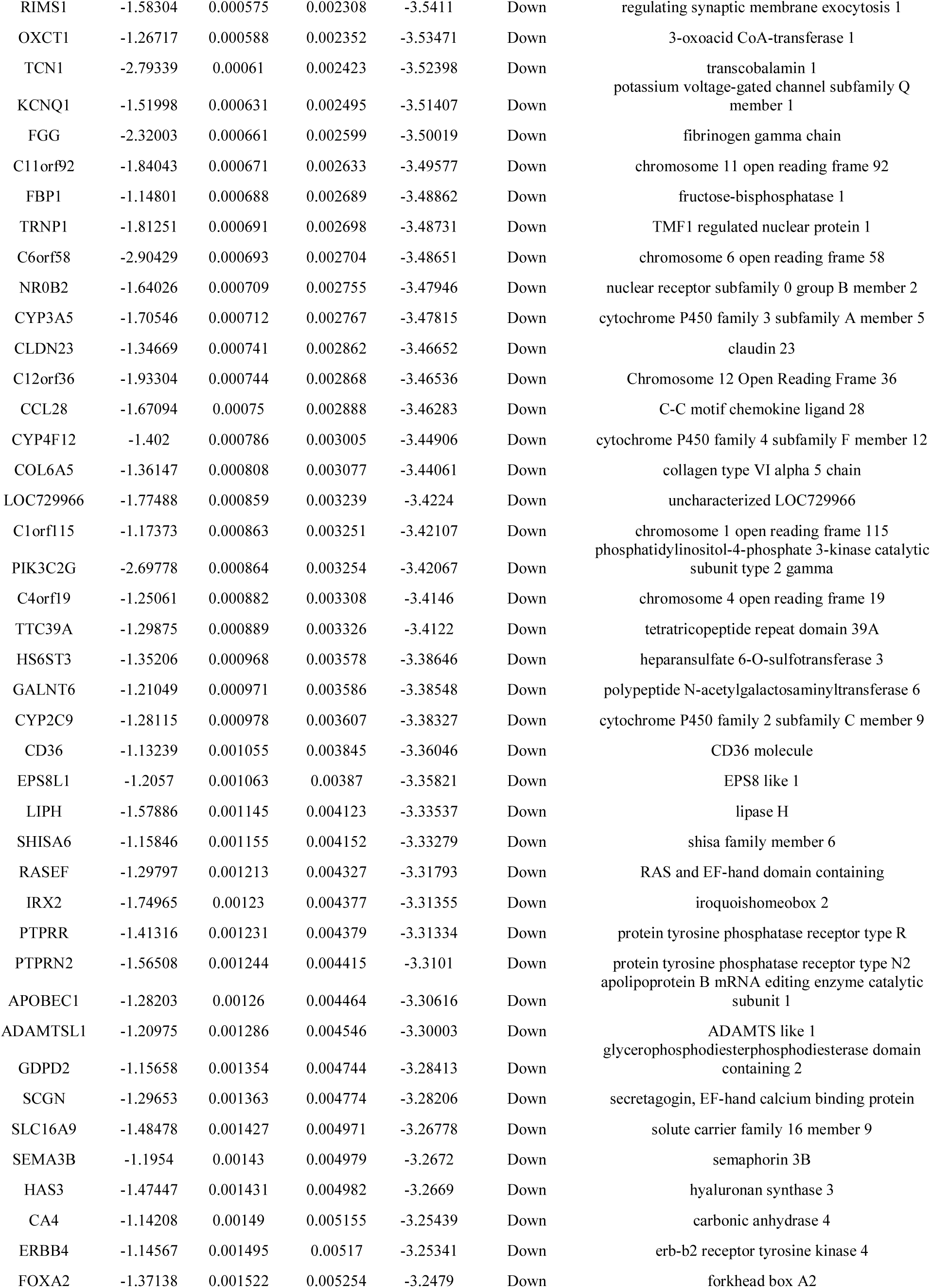
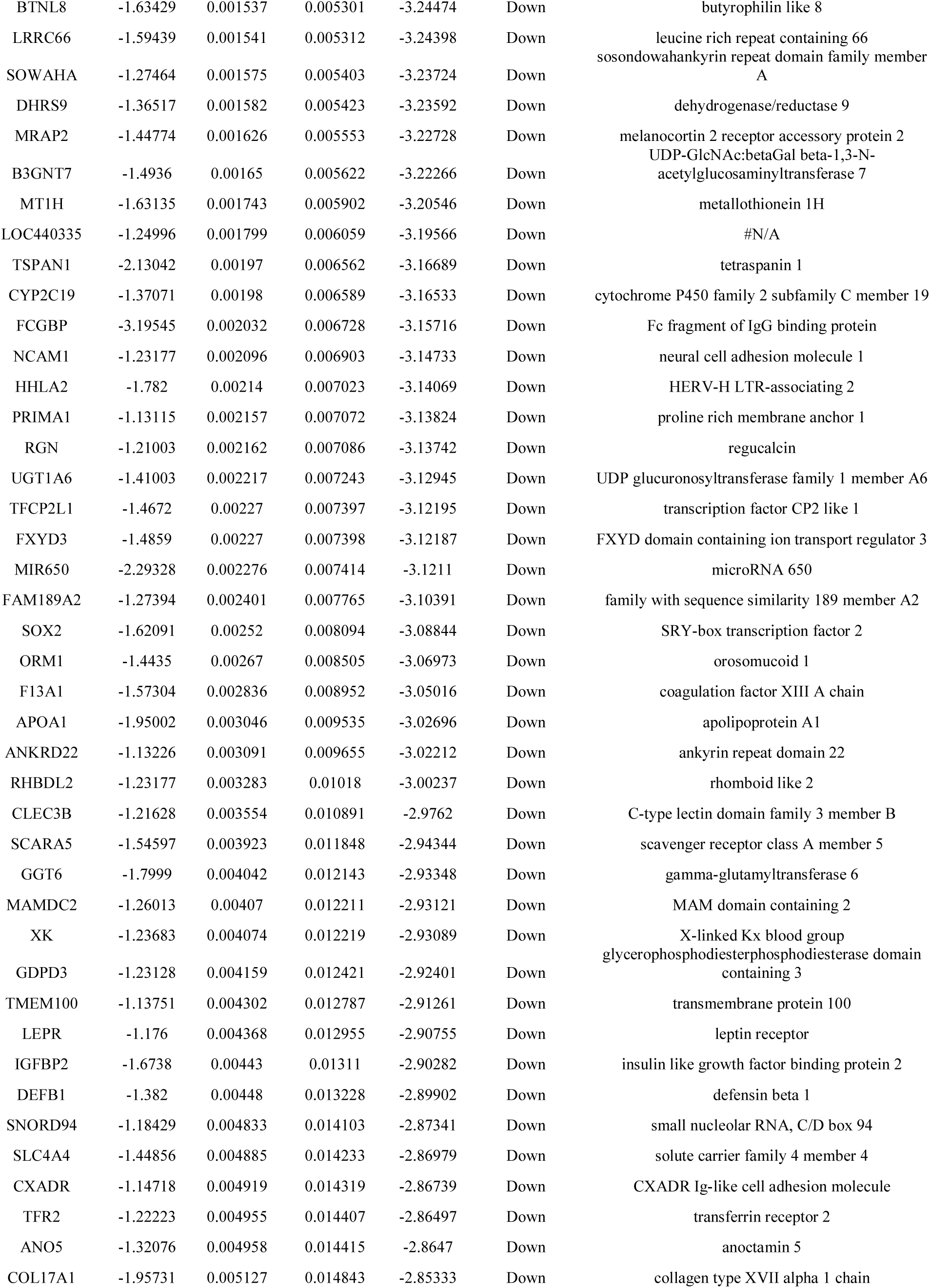
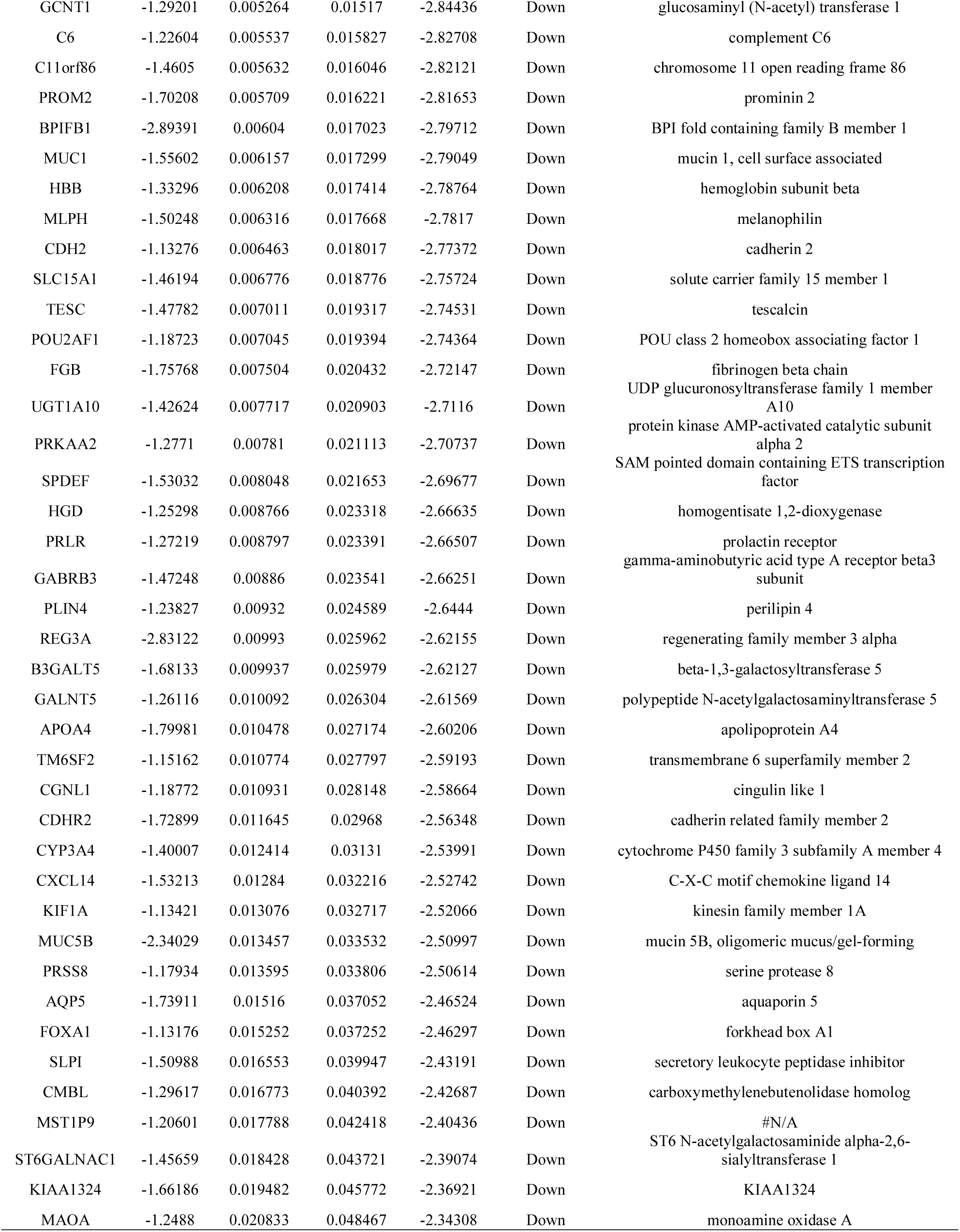
The statistical metrics for key differentially expressed genes (DEGs)

### Pathway enrichment analysis of DEGs

To analyze the biological importance of DEGs, pathway enrichment analyses were executed. Pathway enrichment analysis was predicted that up and down regulated genes were associated with several key physiological processes and are listed in Table 3 and Table 4. Up regulated genes were significantly enriched in pyrimidine deoxyribonucleosides degradation, aspirin triggered resolvin D biosynthesis, ECM-receptor interaction, protein digestion and absorption, syndecan-4-mediated signaling events, PLK1 signaling events, extracellular matrix organization, hemostasis, glutamate metabolism, pyrimidine metabolism, ensemble of genes encoding extracellular matrix and extracellular matrix-associated proteins, ensemble of genes encoding core extracellular matrix including ECM glycoproteins, collagens and proteoglycans, integrin signalling pathway, plasminogen activating cascade, altered lipoprotein metabolic, ammonia recycling and MNGIE (Mitochondrial Neurogastrointestinal Encephalopathy), while down regulated genes were significantly enriched in allopregnanolone biosynthesis, gluconeogenesis, drug metabolism - cytochrome P450, chemical carcinogenesis, FOXA2 and FOXA3 transcription factor networks, RhoA signaling pathway, biological oxidations, phase 1 - functionalization of compounds, carbon fixation, fatty acid metabolism, ensemble of genes encoding extracellular matrix and extracellular matrix-associated proteins, ensemble of genes encoding ECM-associated proteins including ECM-affilaited proteins, ECM regulators and secreted factors, 5-Hydroxytryptamine degredation, fructose galactose metabolism, lipoprotein metabolic, pirenzepine pathway and tyrosine metabolism.

**Table 3.**
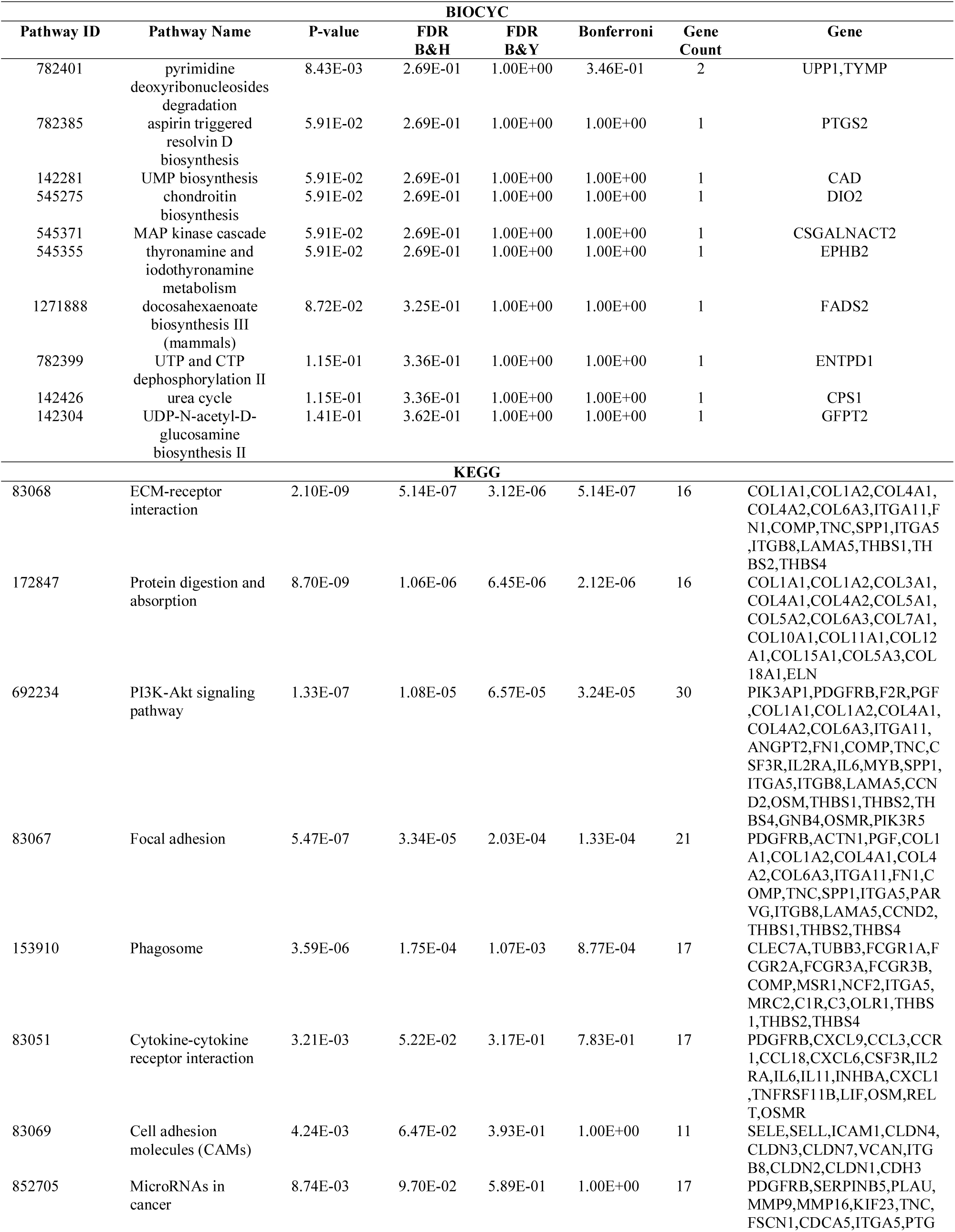
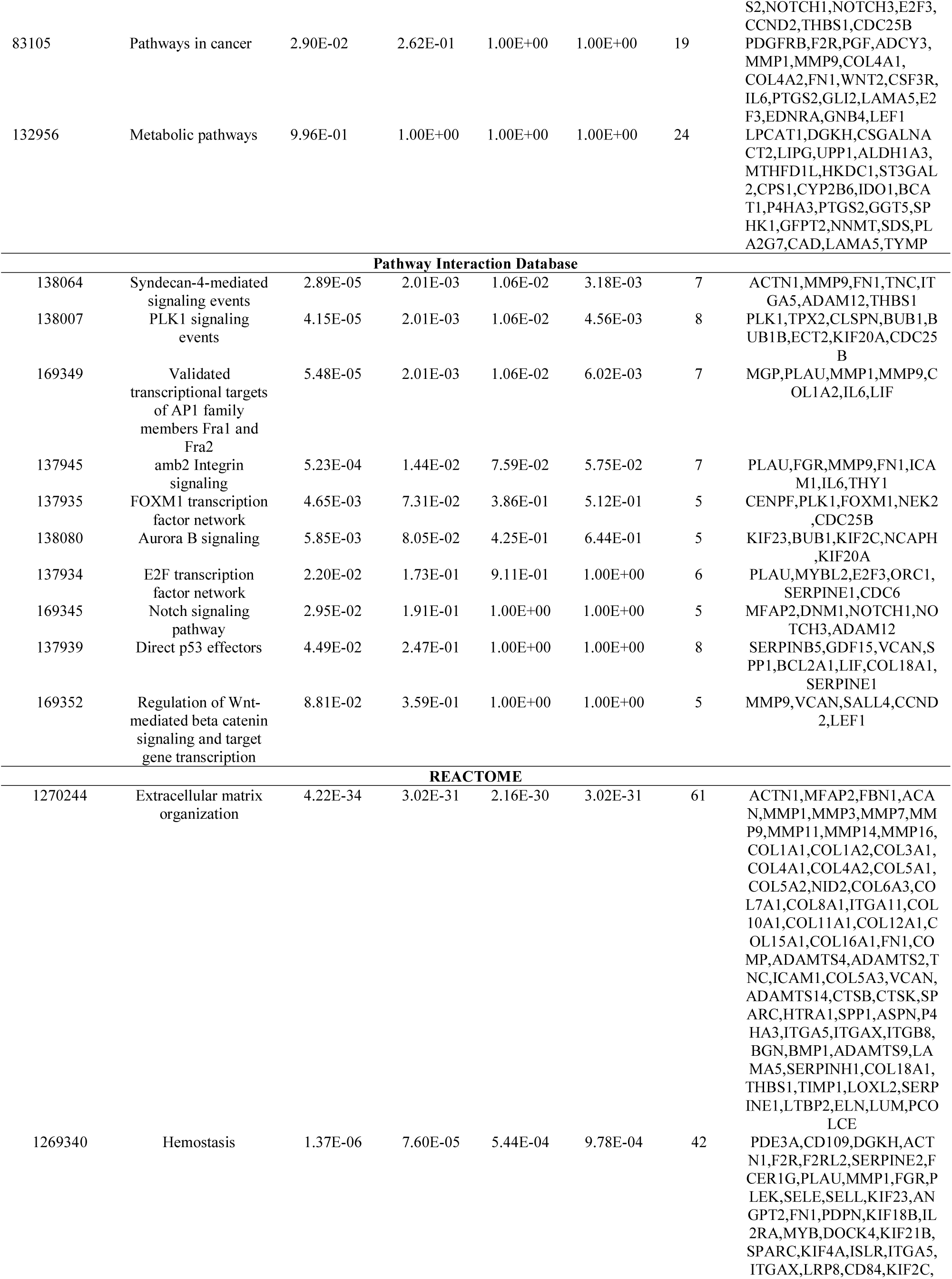
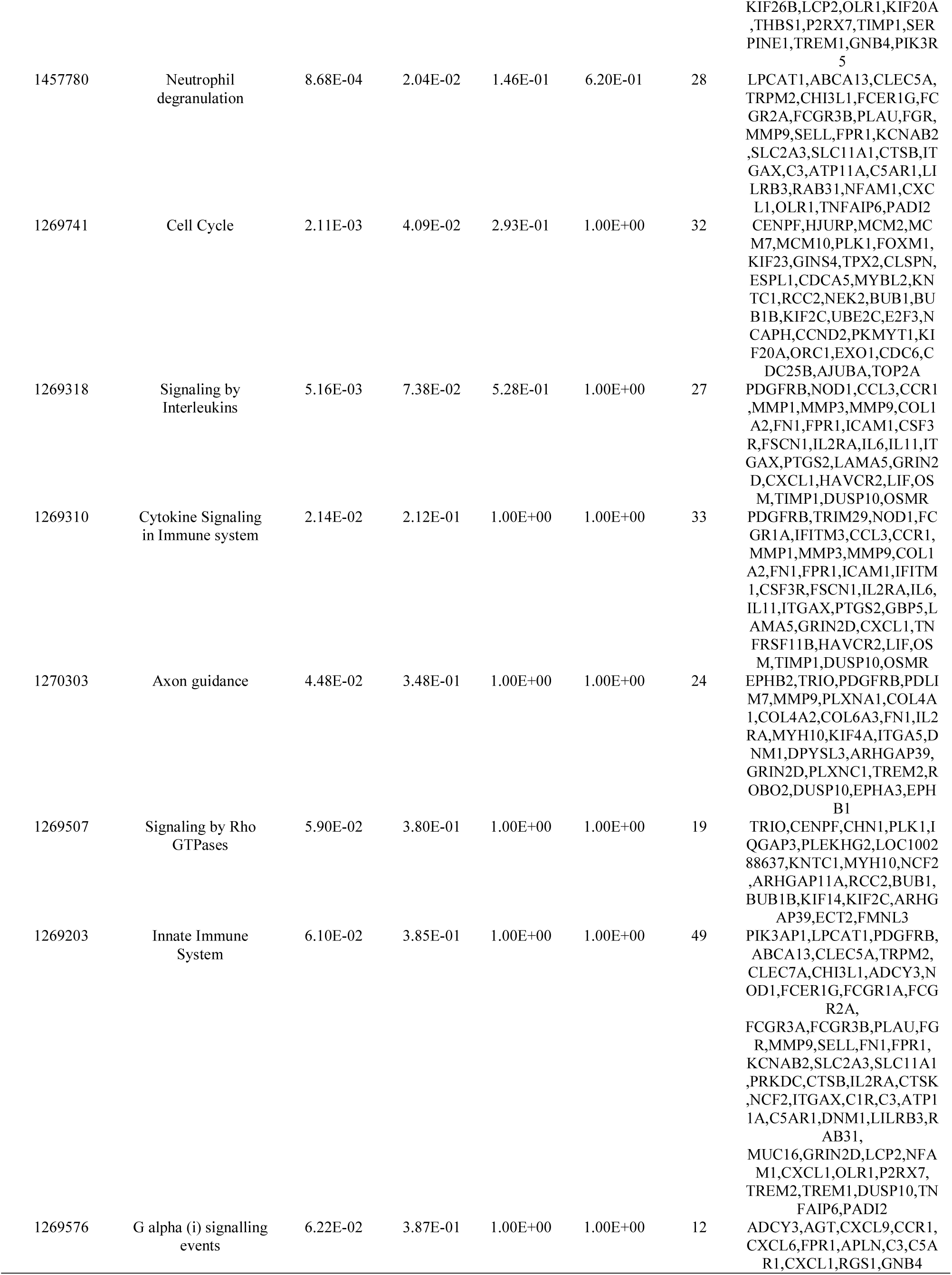
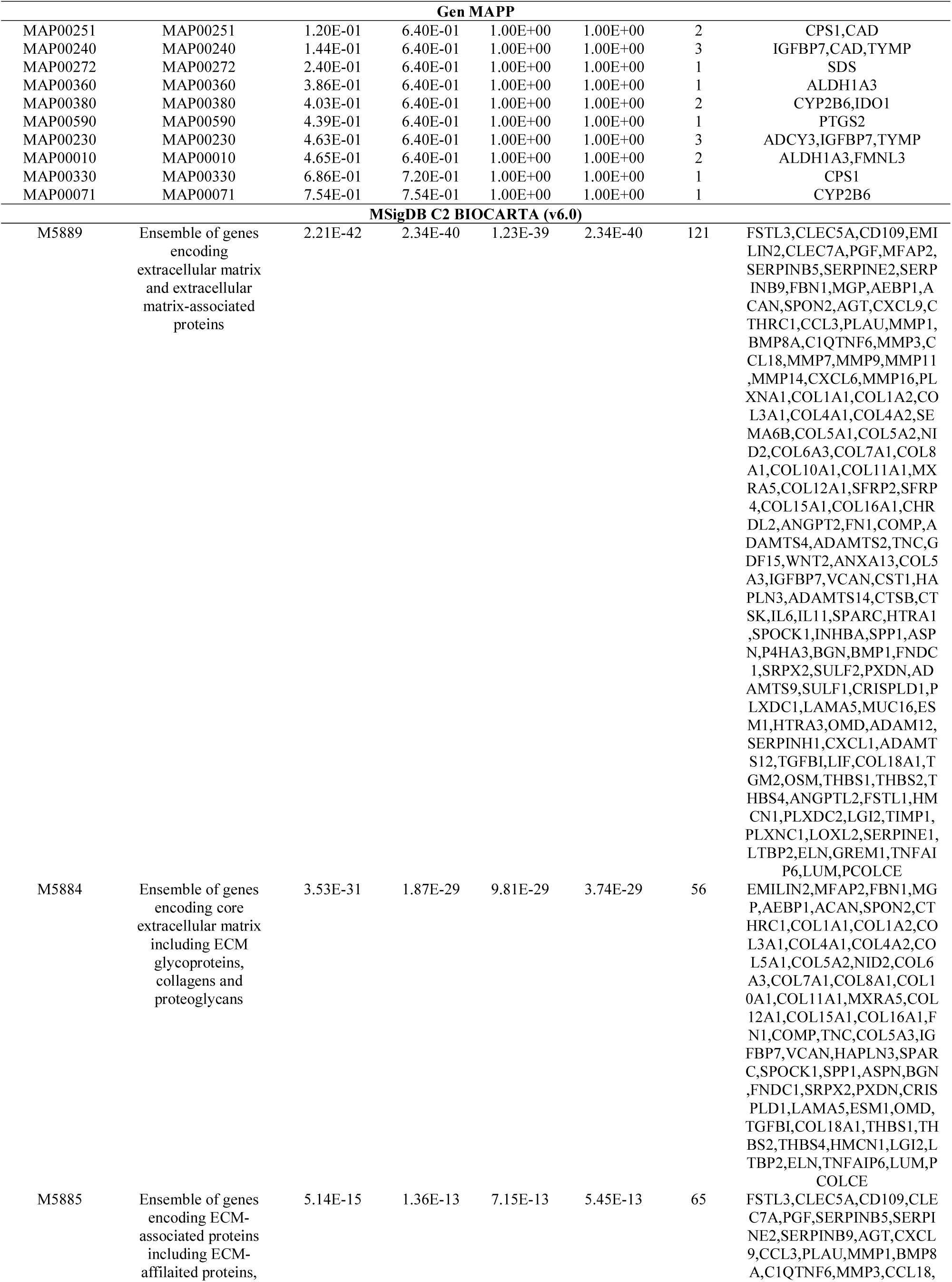
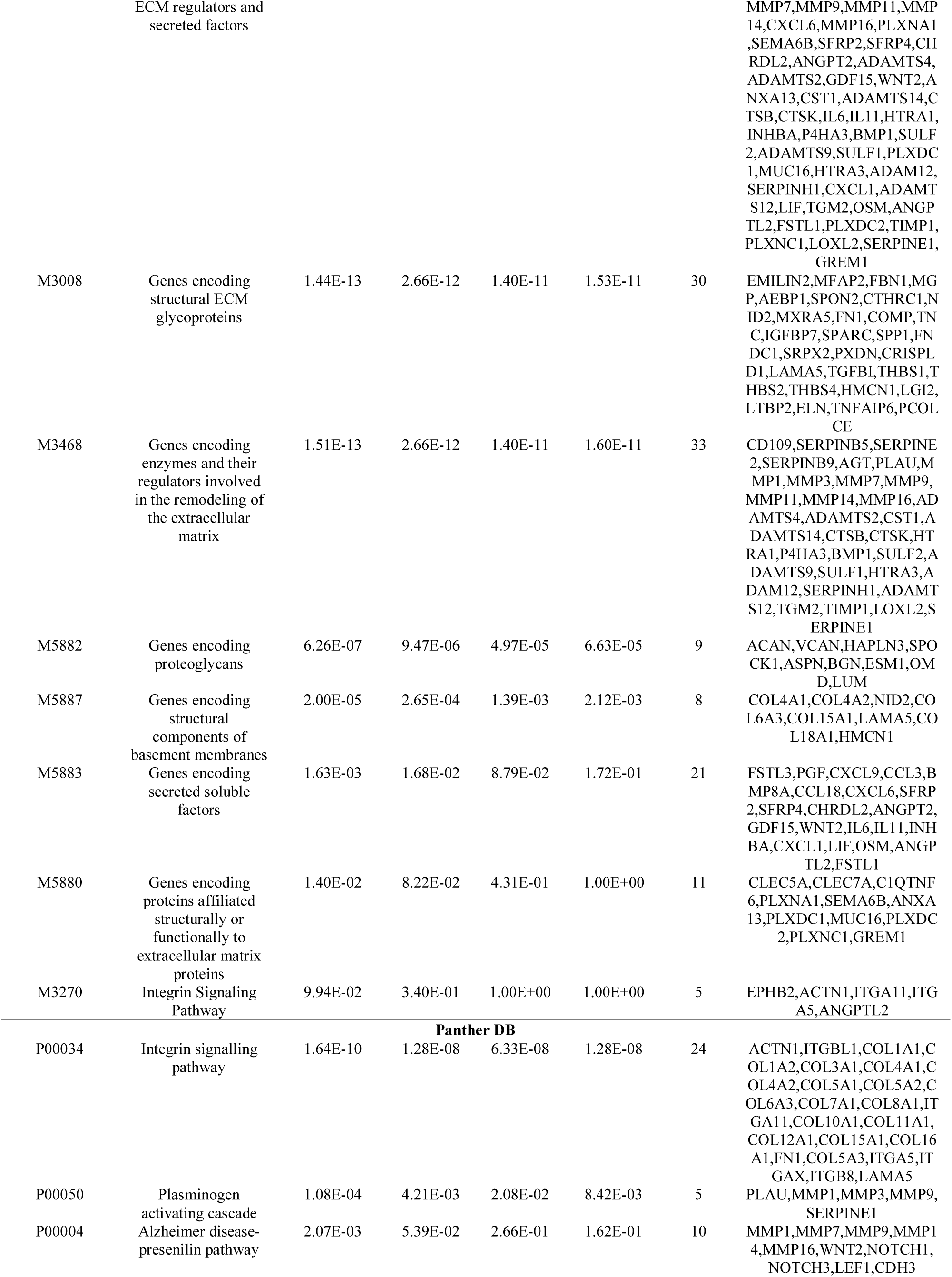
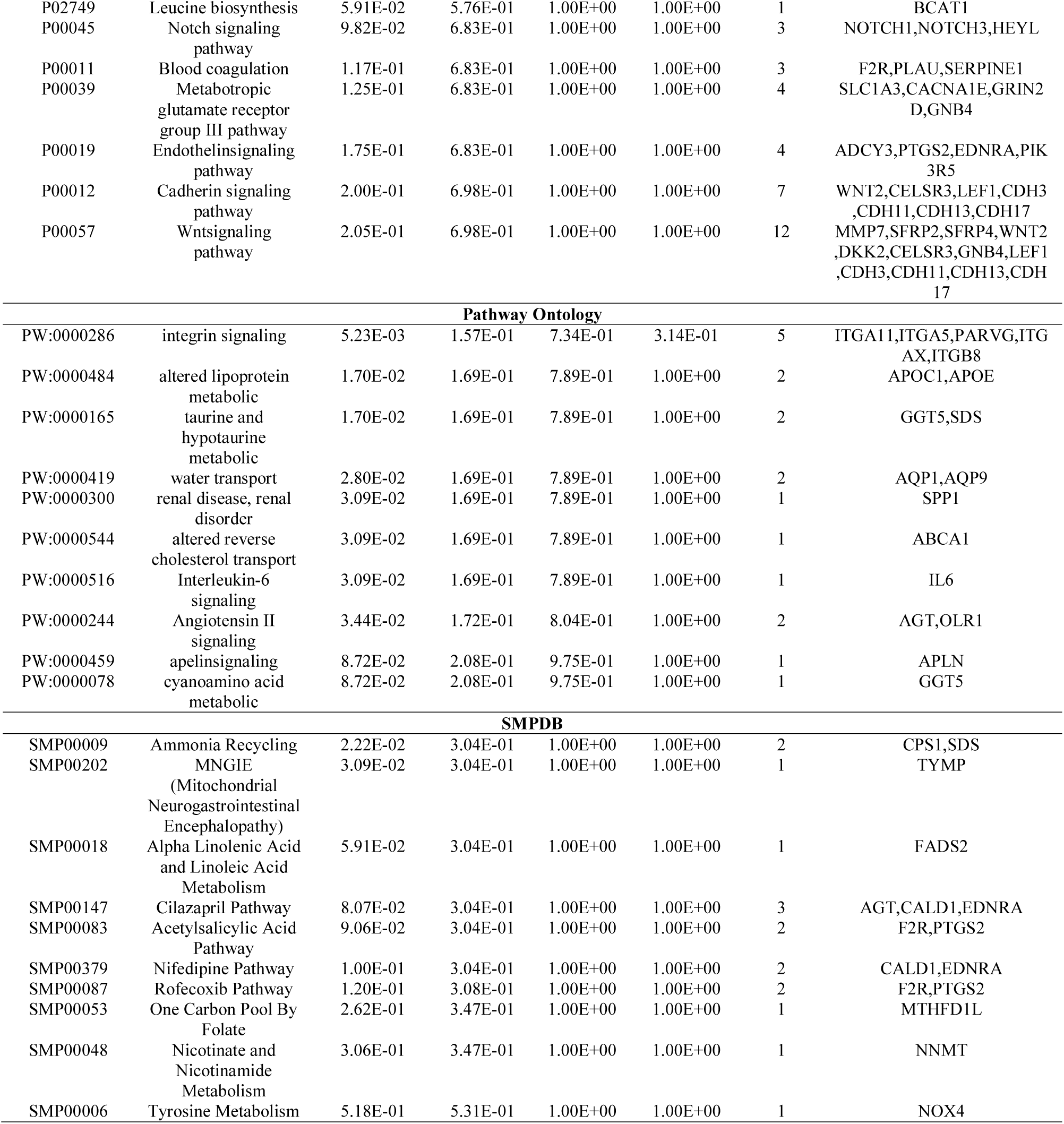
The enriched pathway terms of the up regulated differentially expressed genes

**Table 4.**
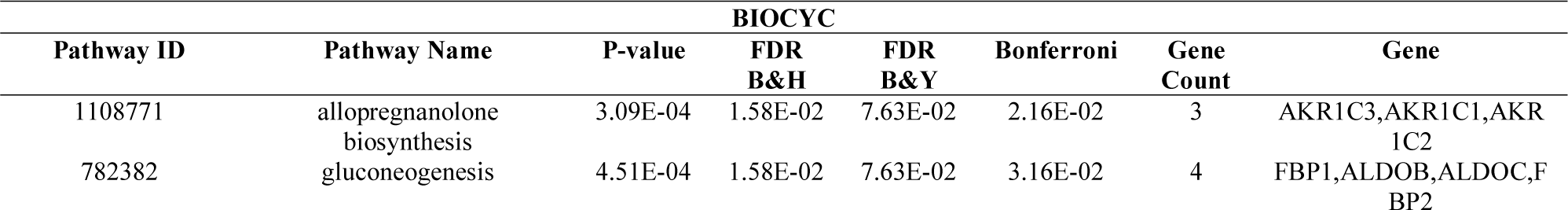
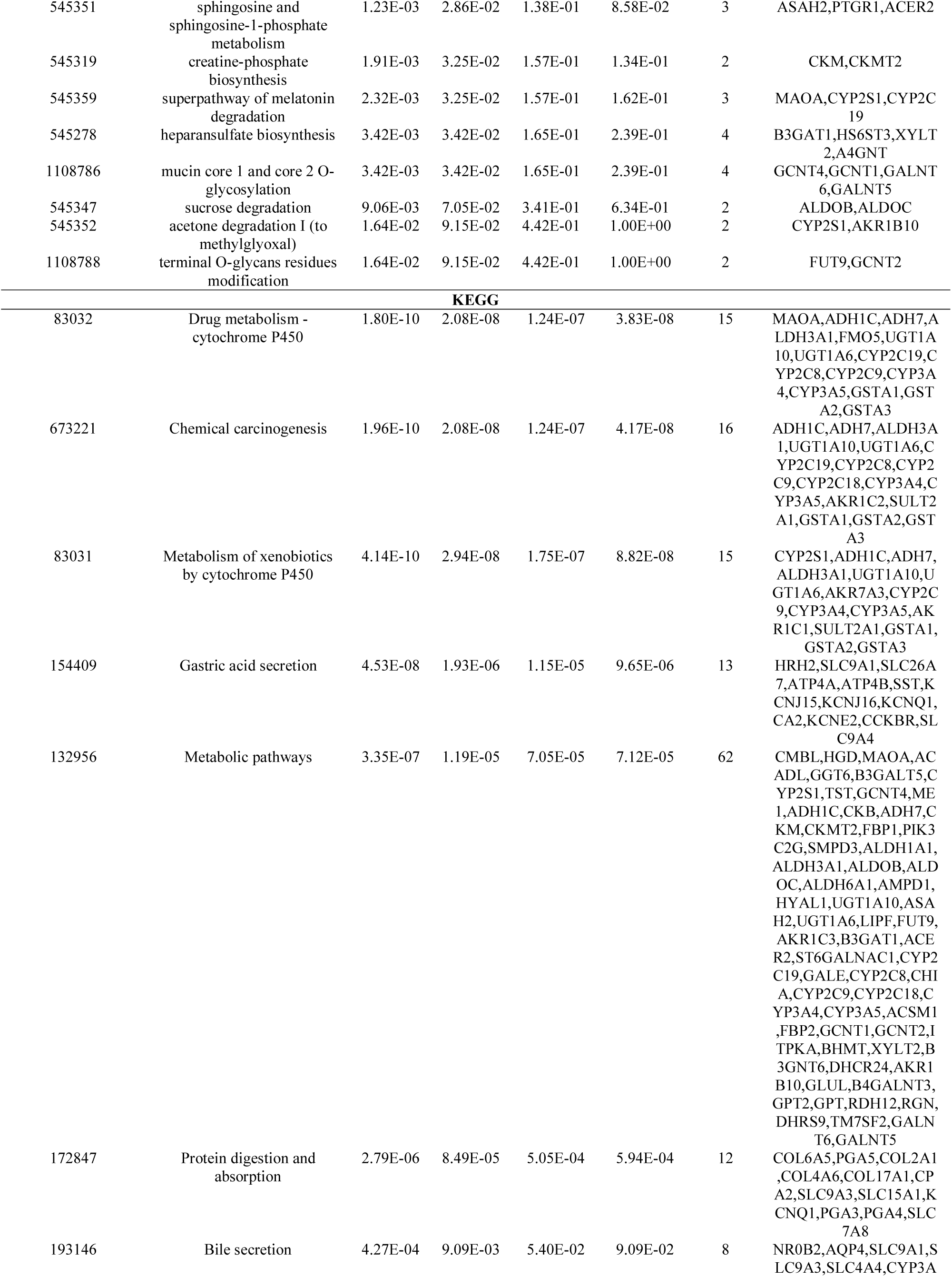
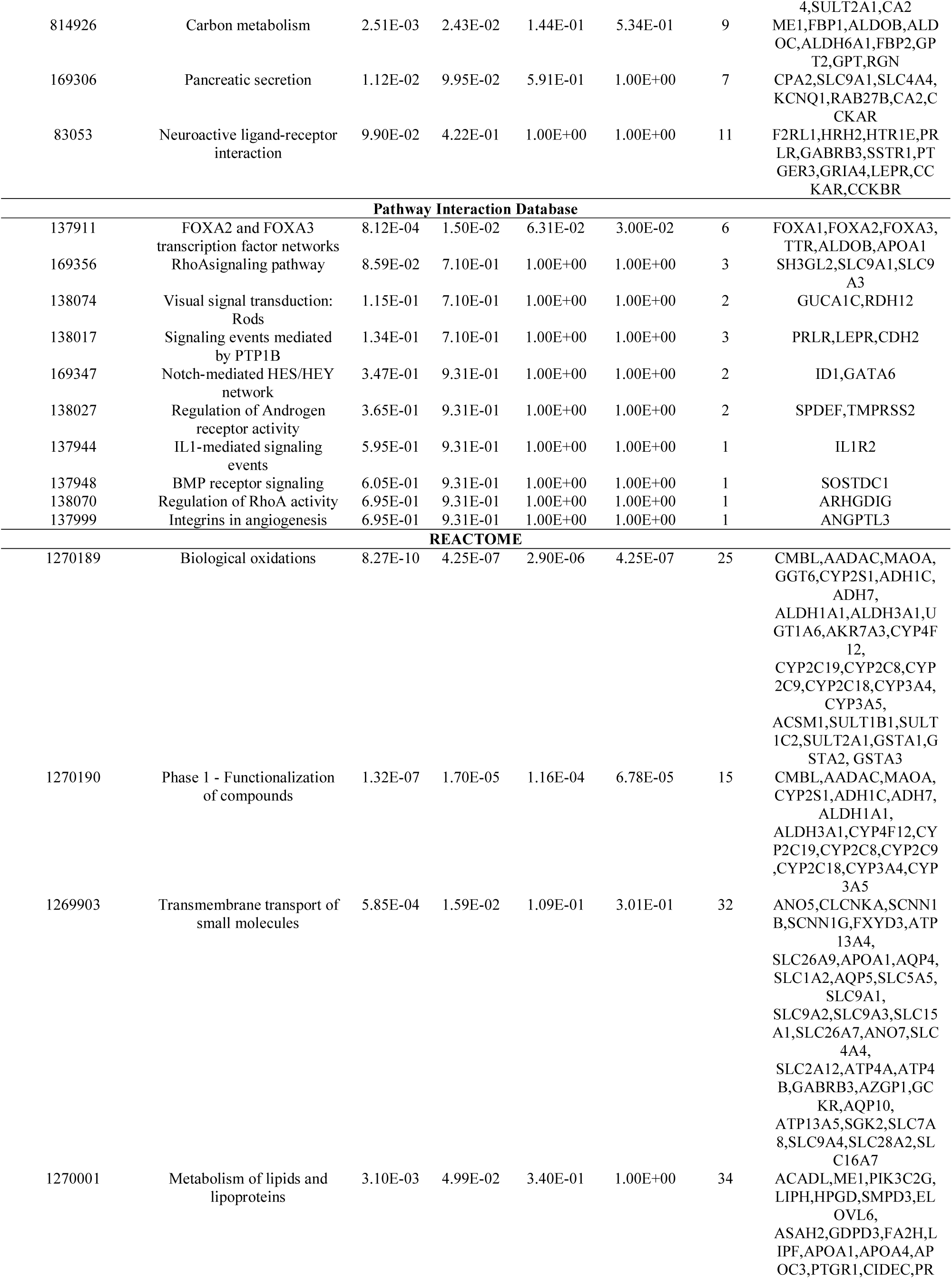
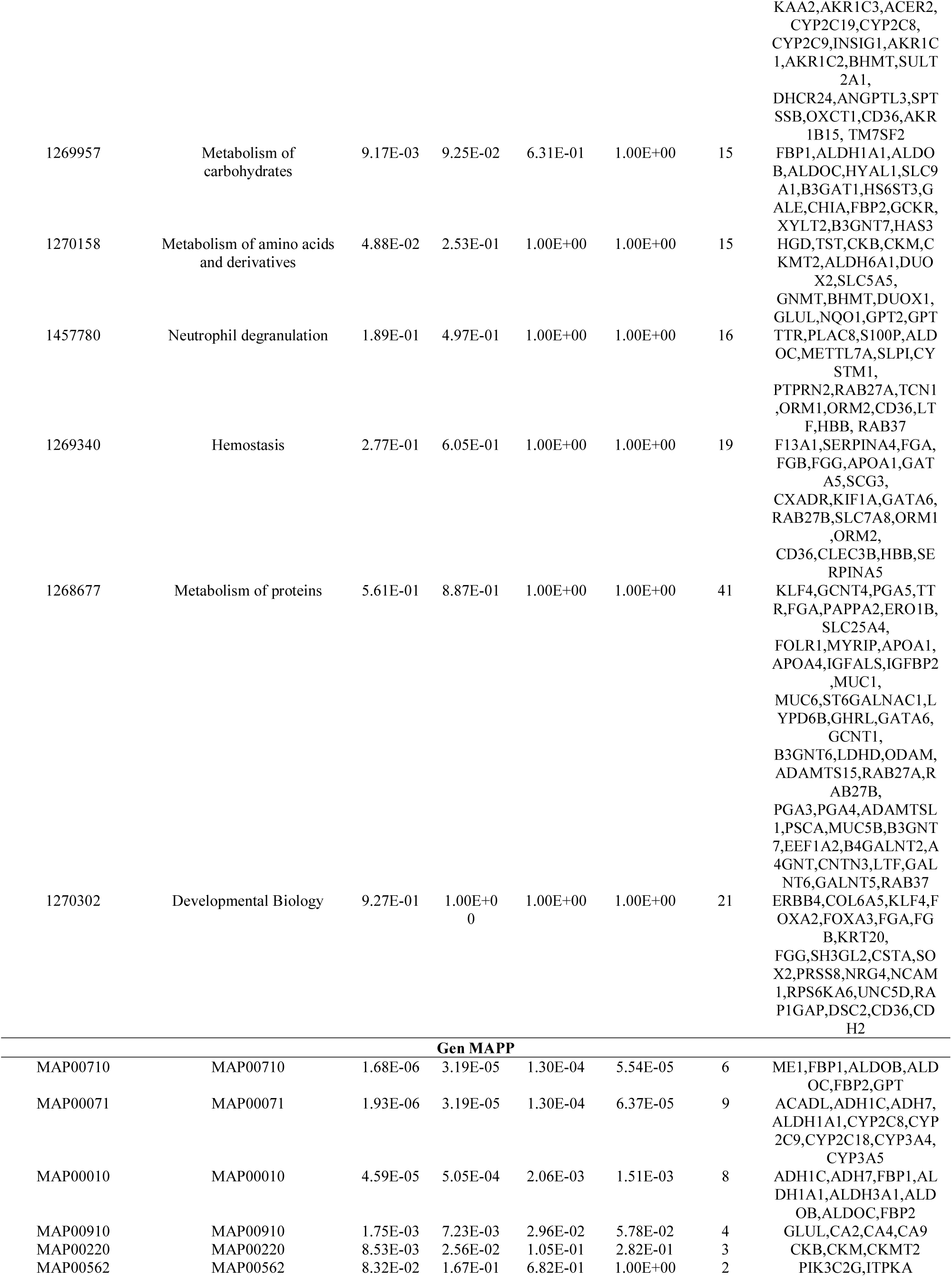
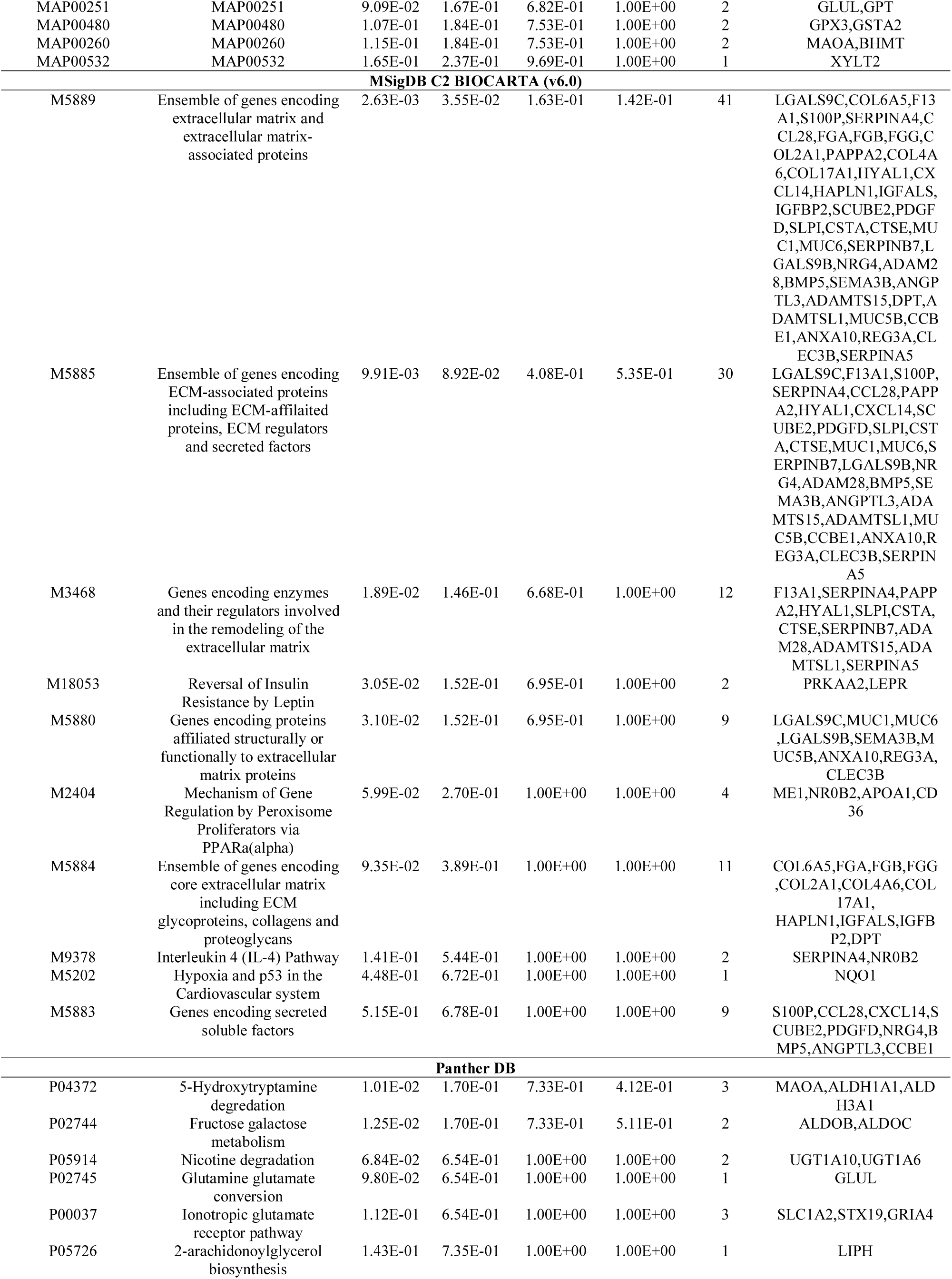
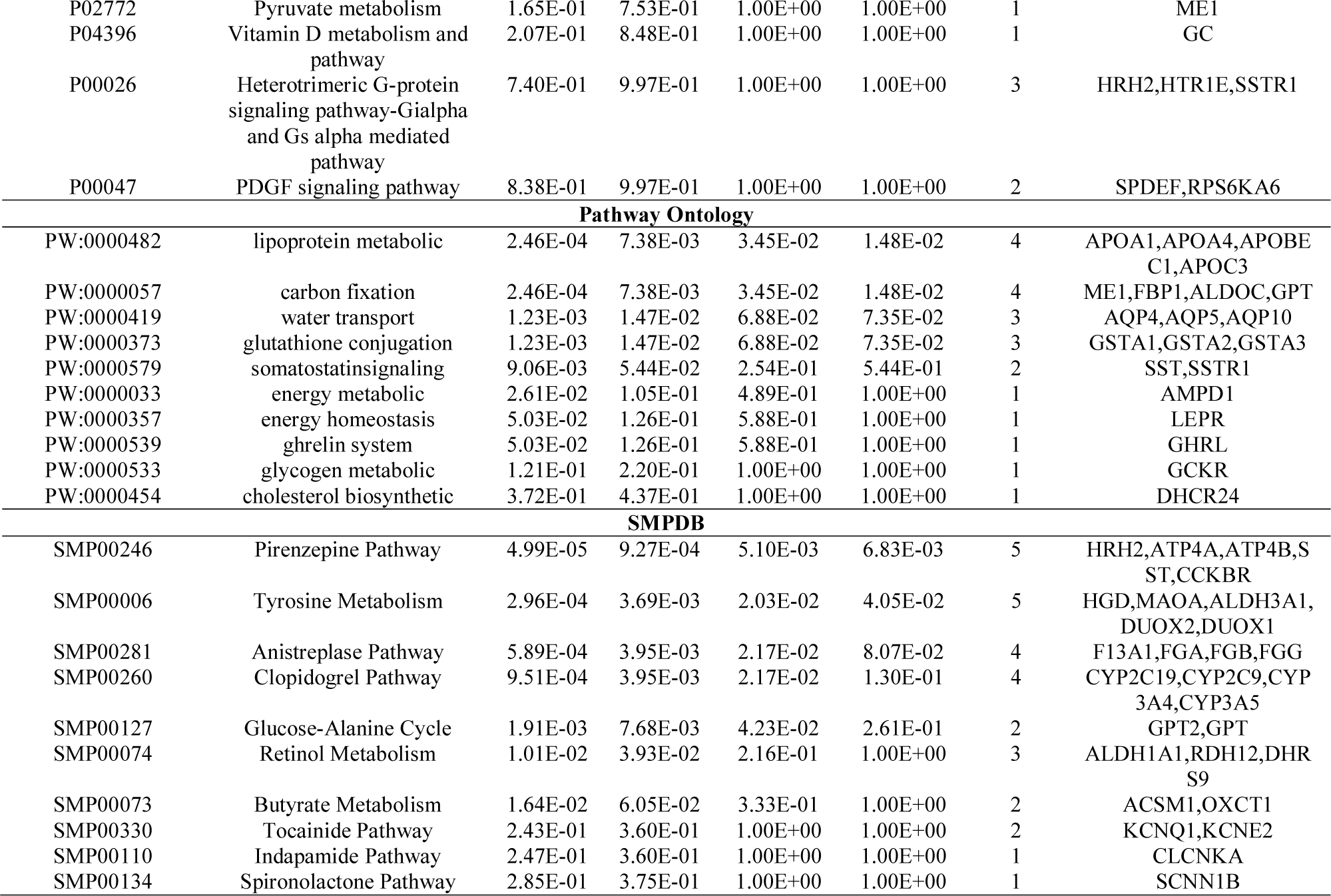
The enriched pathway terms of the down regulated differentially expressed genes

### Gene ontology (GO) enrichment analysis of DEGs

We uploaded all up and down regulated genes to the online software ToppGene to identify overrepresented GO categories. GO analysis results showed that up regulated genes were significantly enriched in BP, CC and MF, including extracellular structure organization, cell adhesion, extracellular matrix, collagen-containing extracellular matrix, extracellular matrix structural constituent and structural molecule activity, while down regulated genes were significantly enriched in BP, CC and MF, including digestion, organic acid metabolic process, apical part of cell, extracellular matrix, oxidoreductase activity and transition metal ion binding and are listed in Table 5 and Table 6.

**Table 5.**
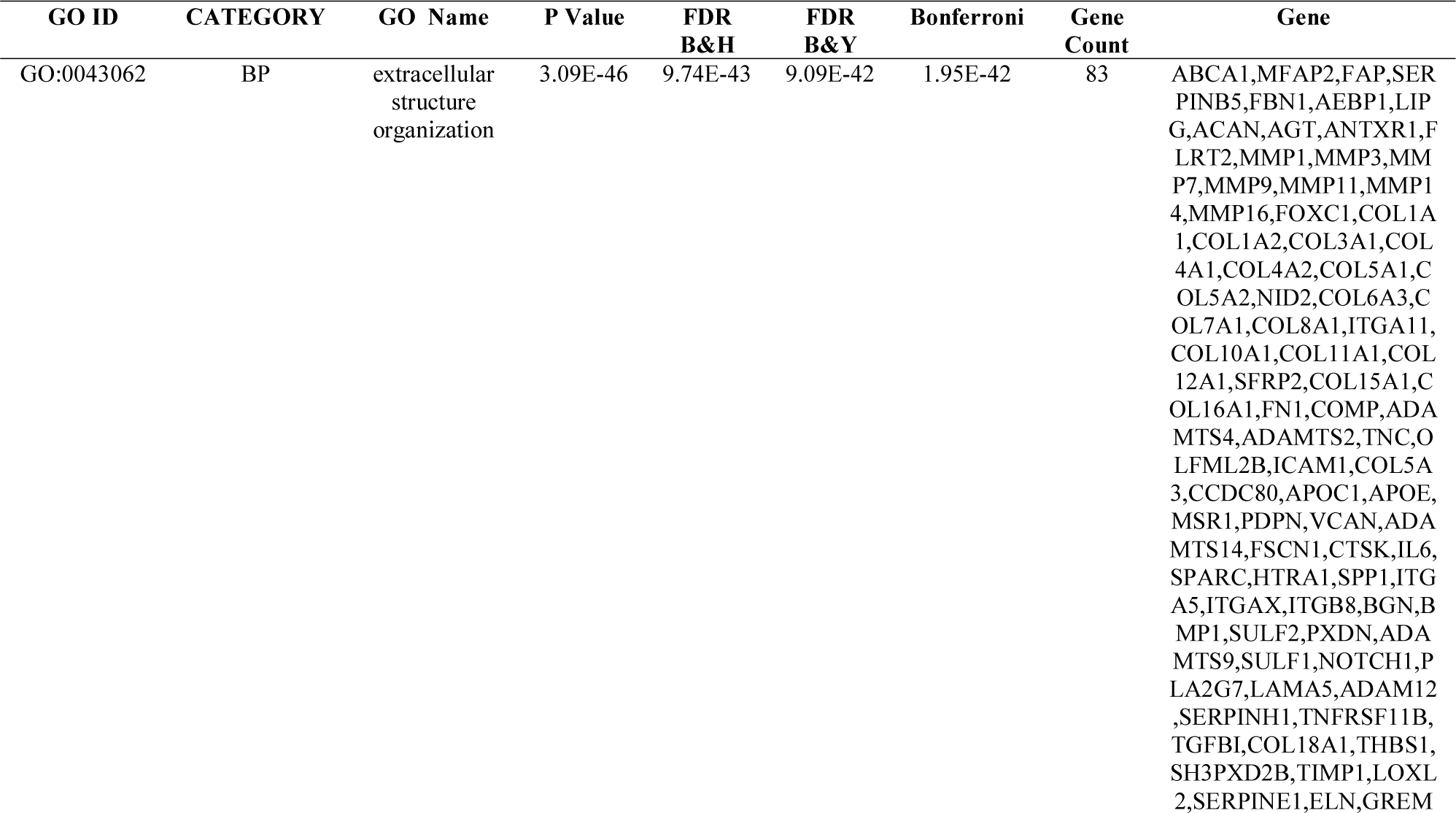
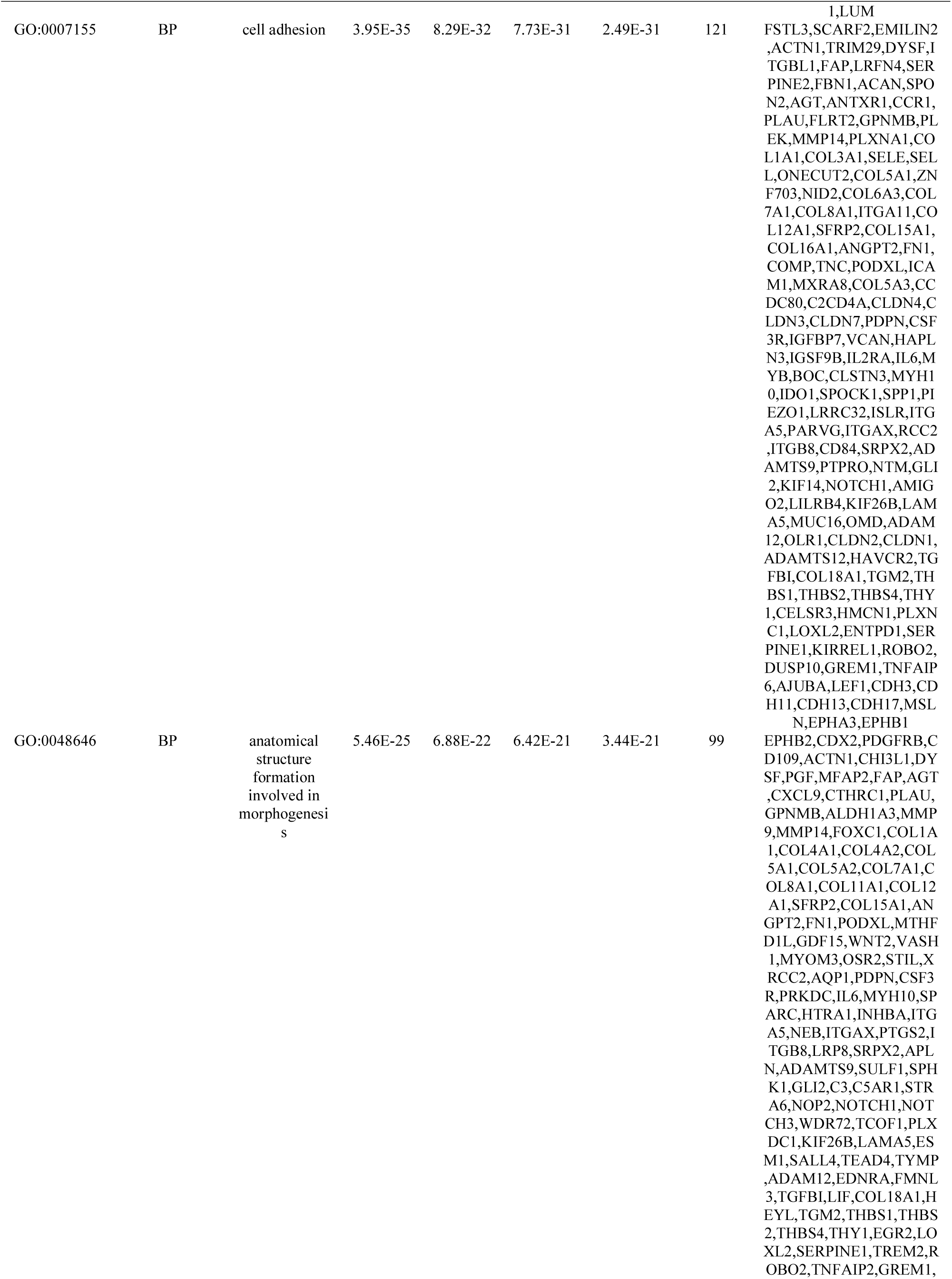
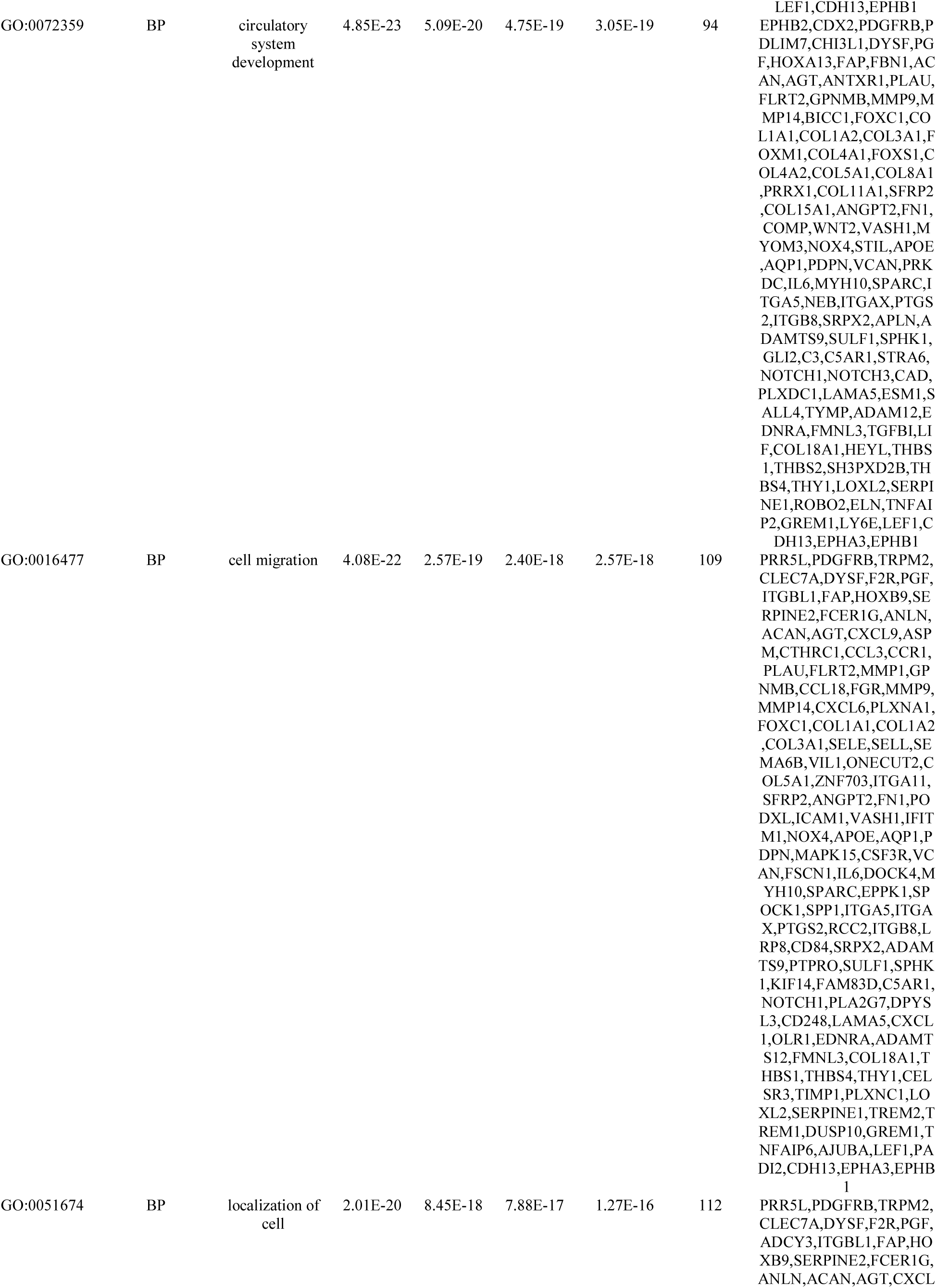
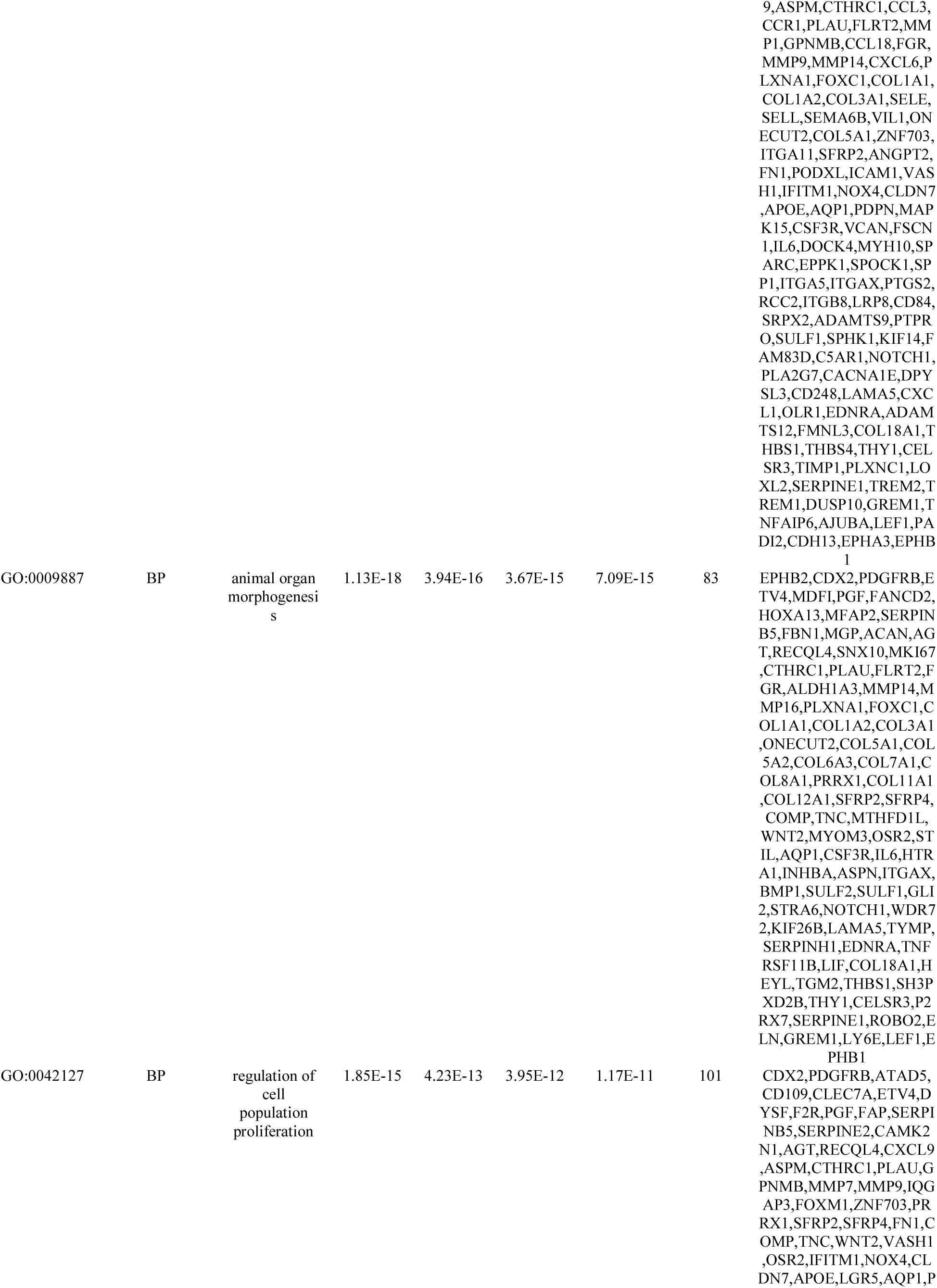
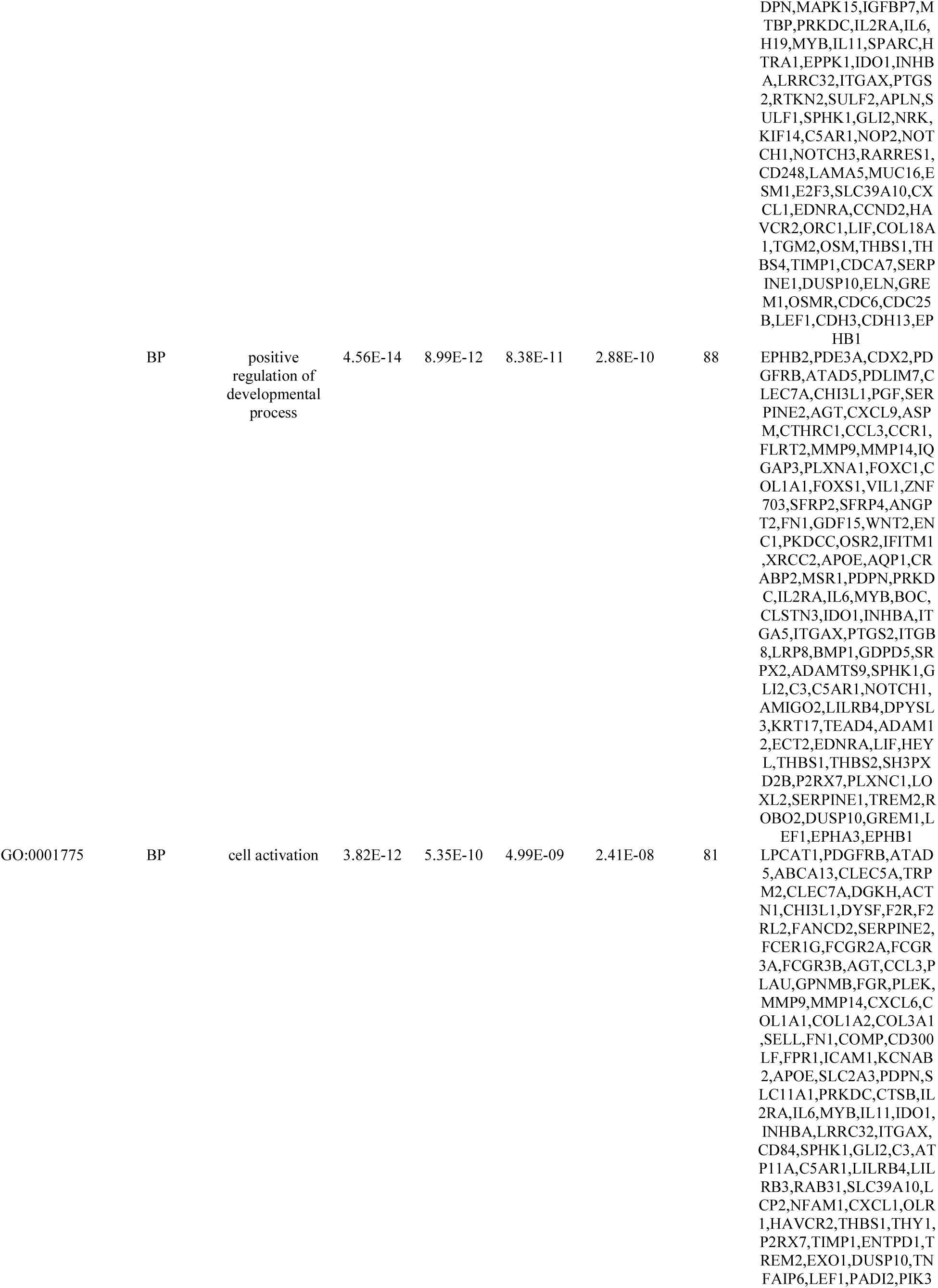
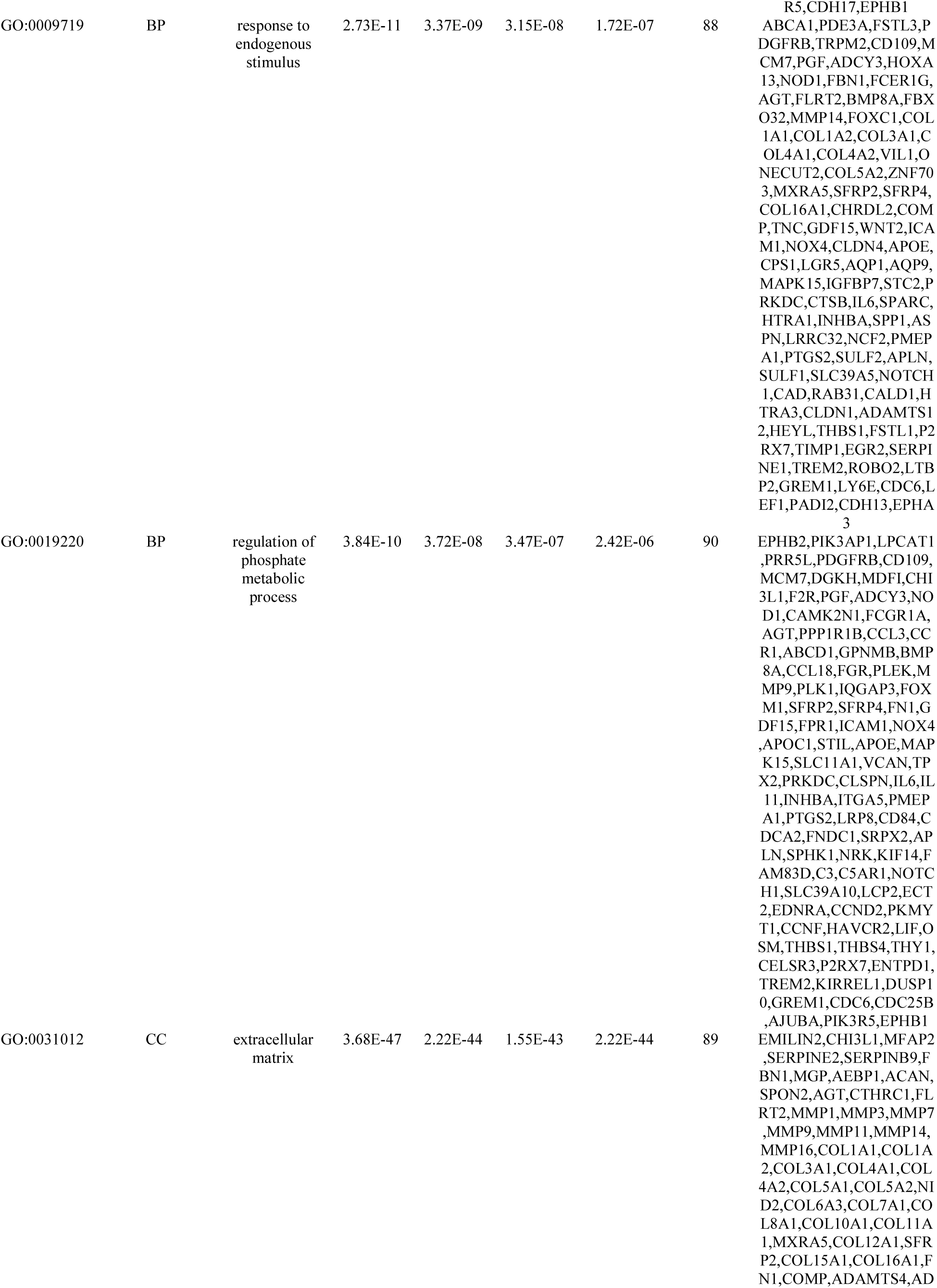
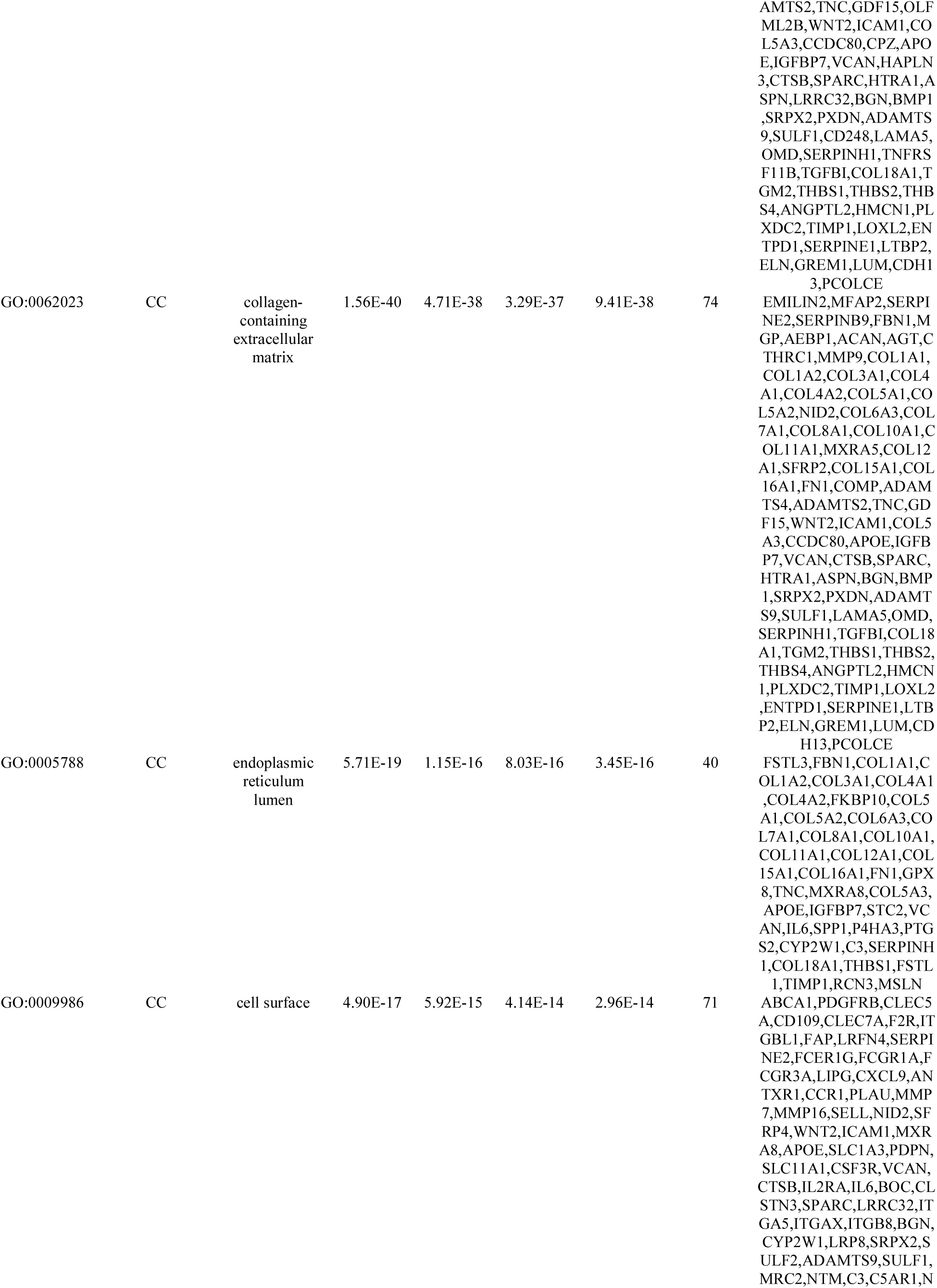
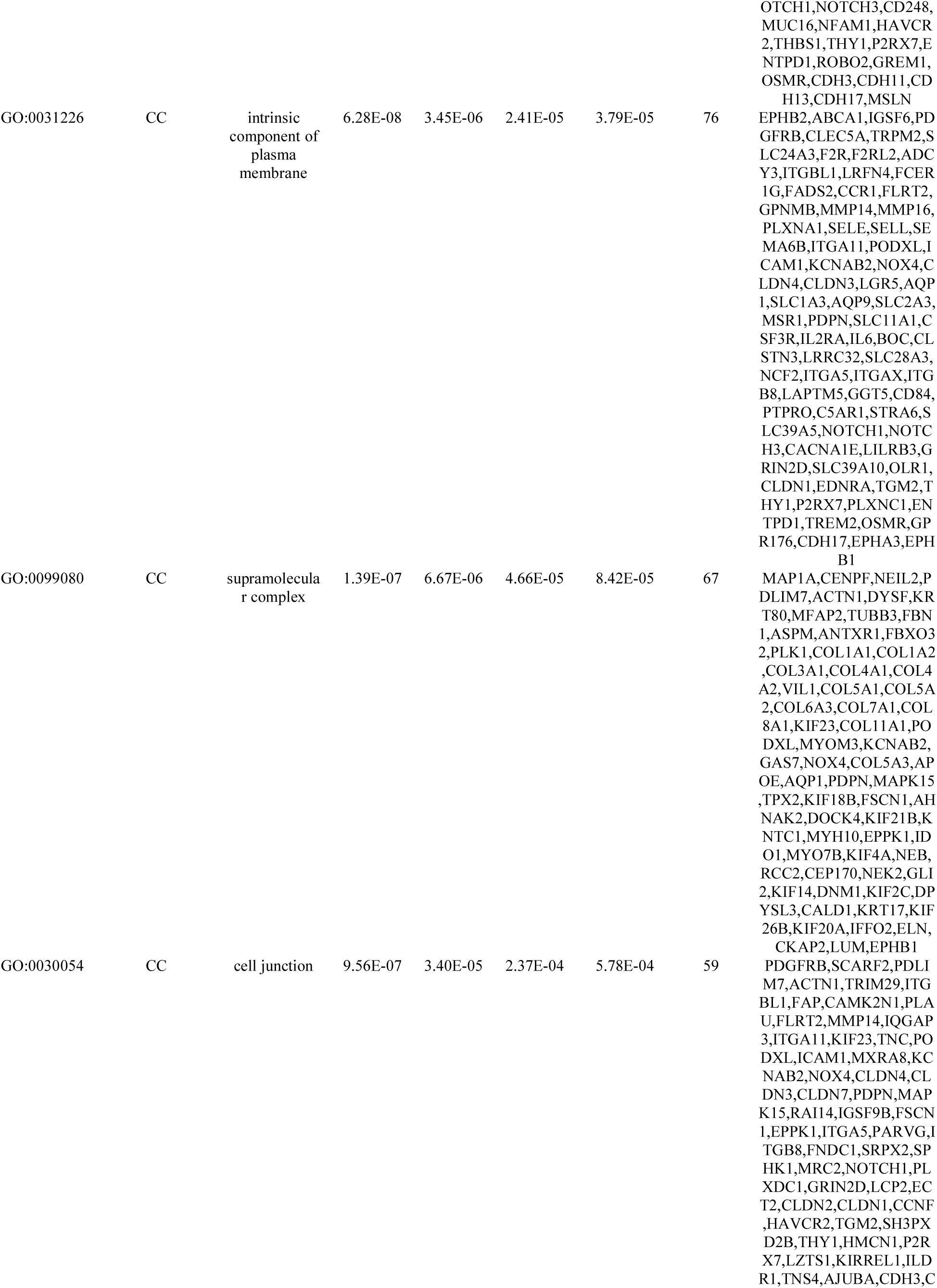
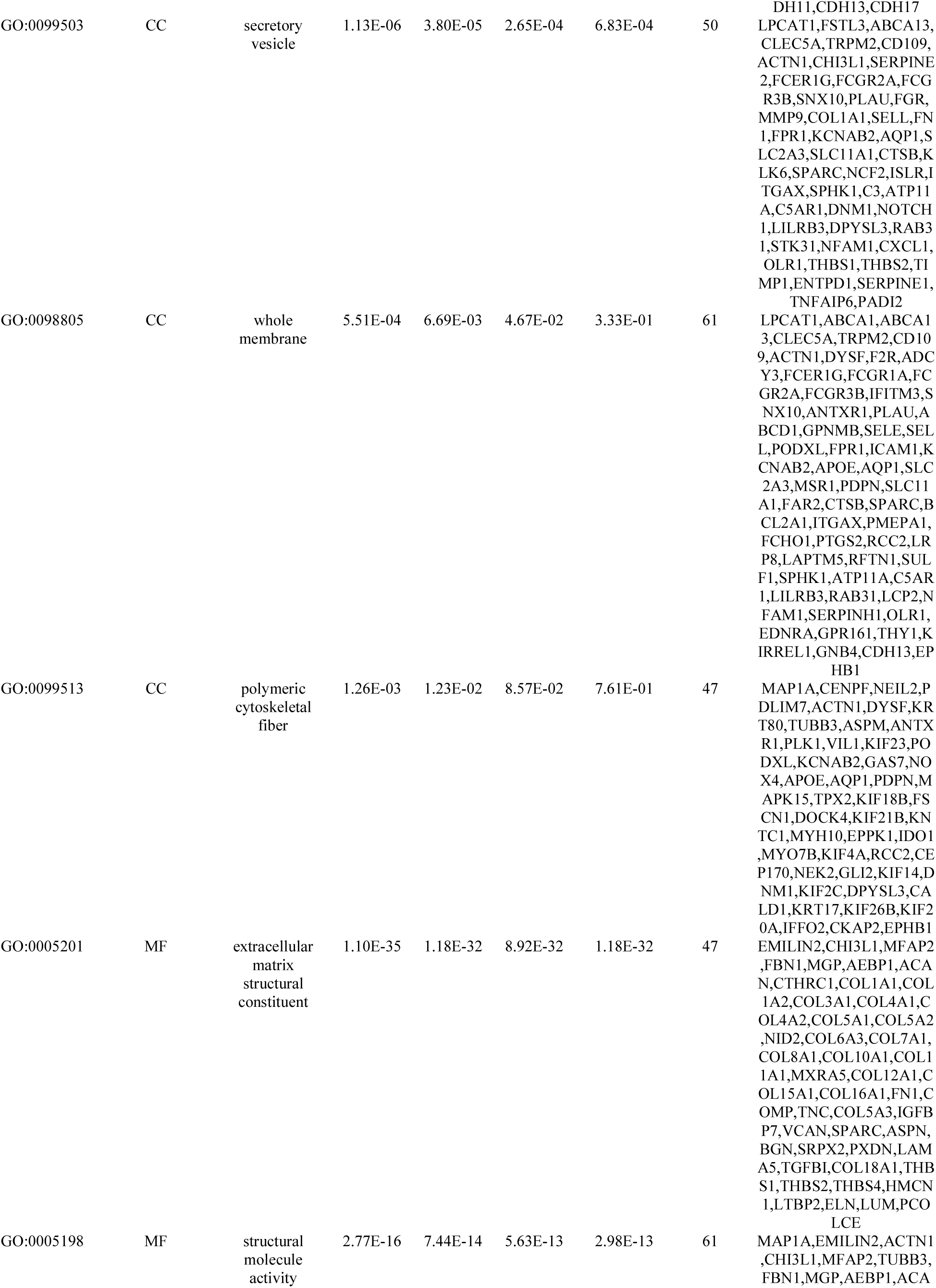
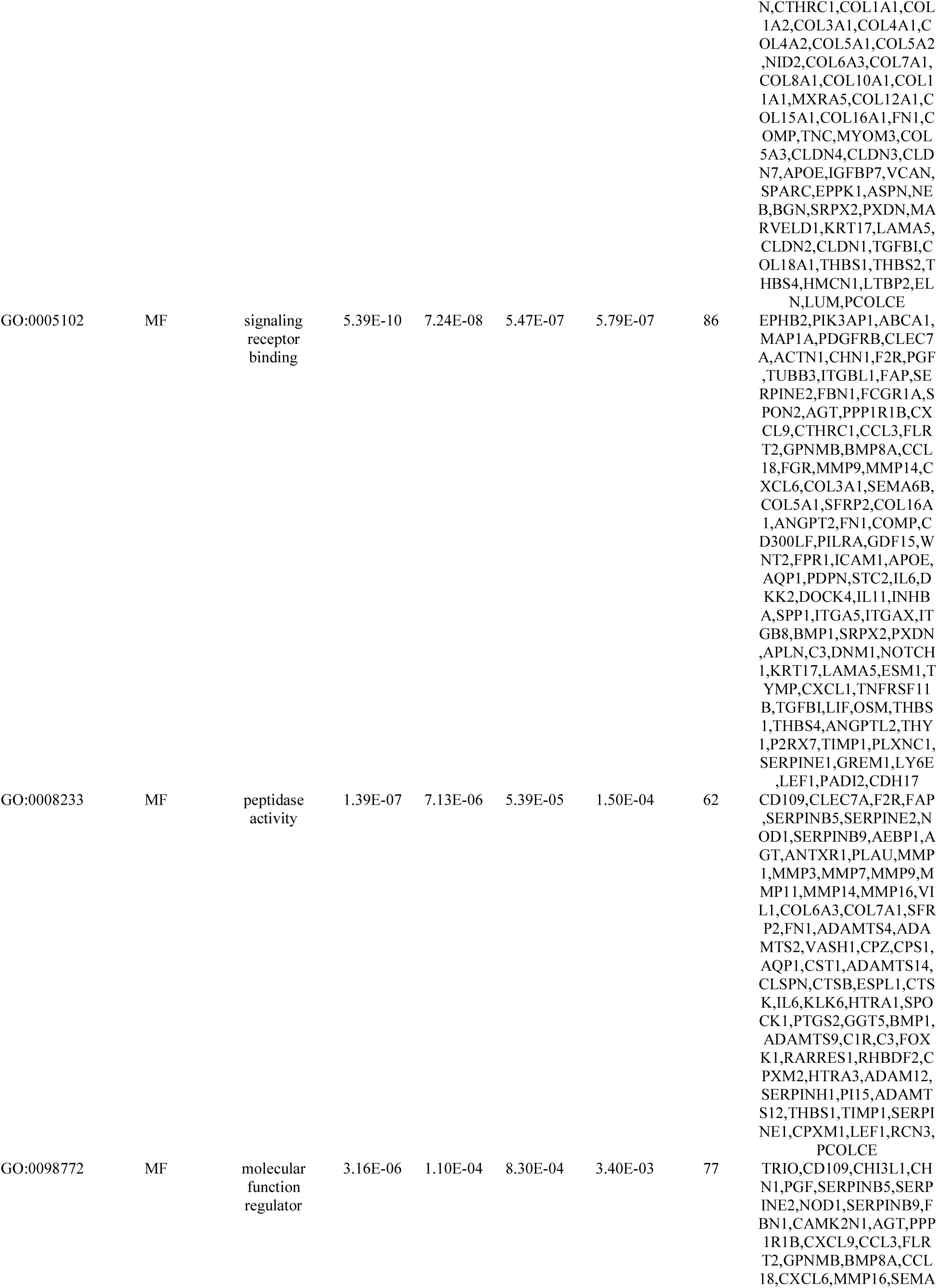
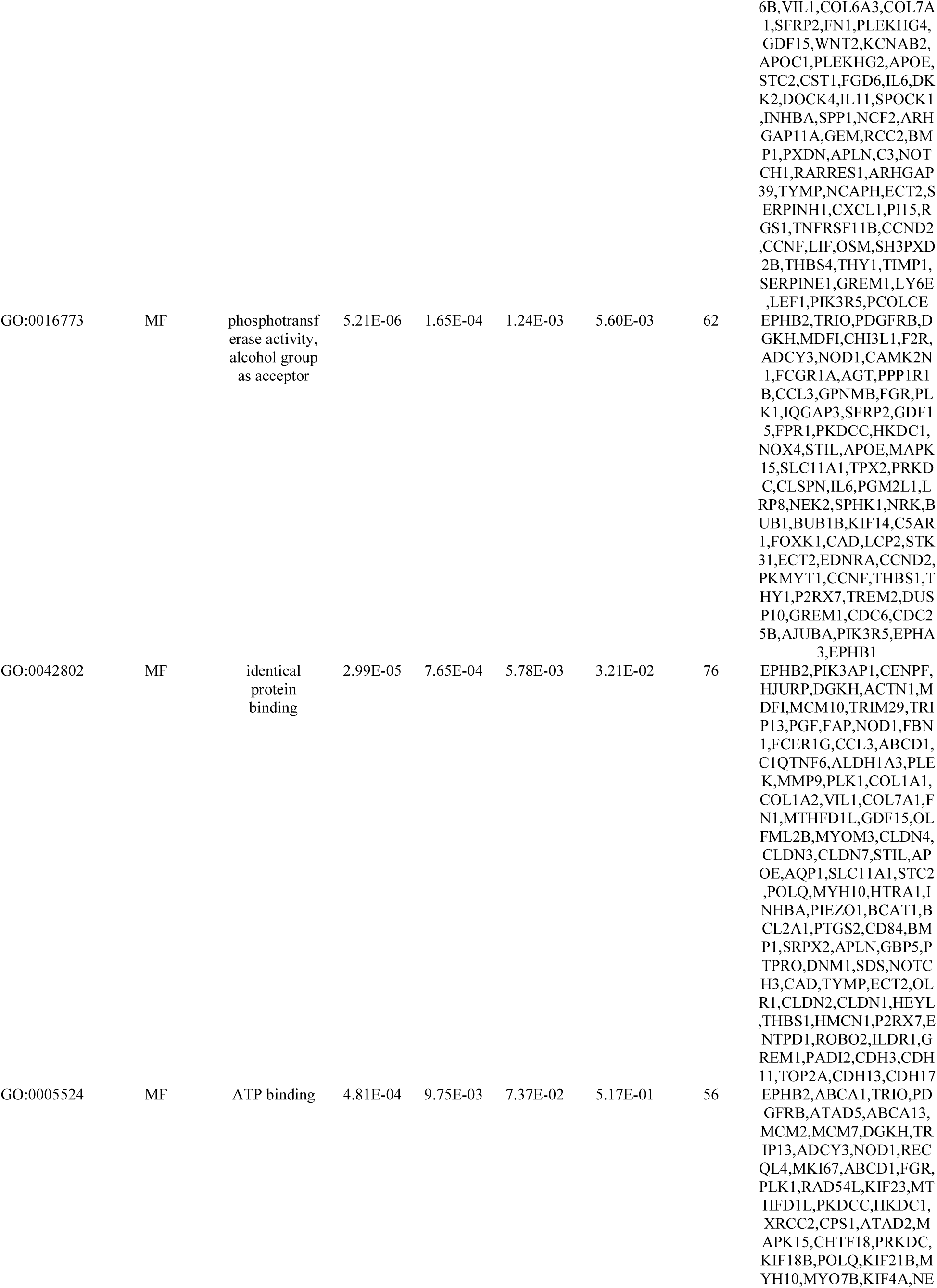
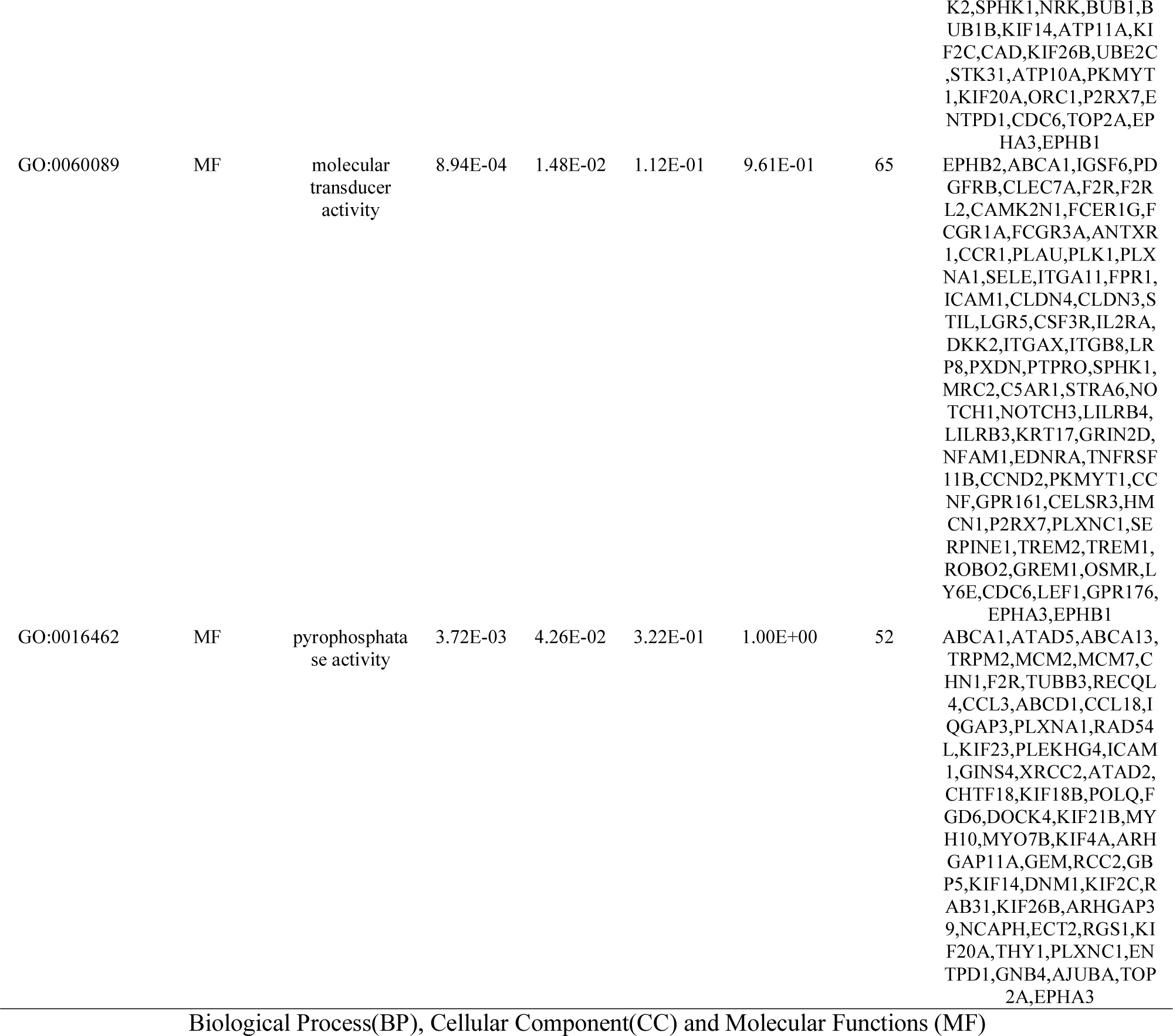
The enriched GO terms of the up regulated differentially expressed genes

**Table 6.**
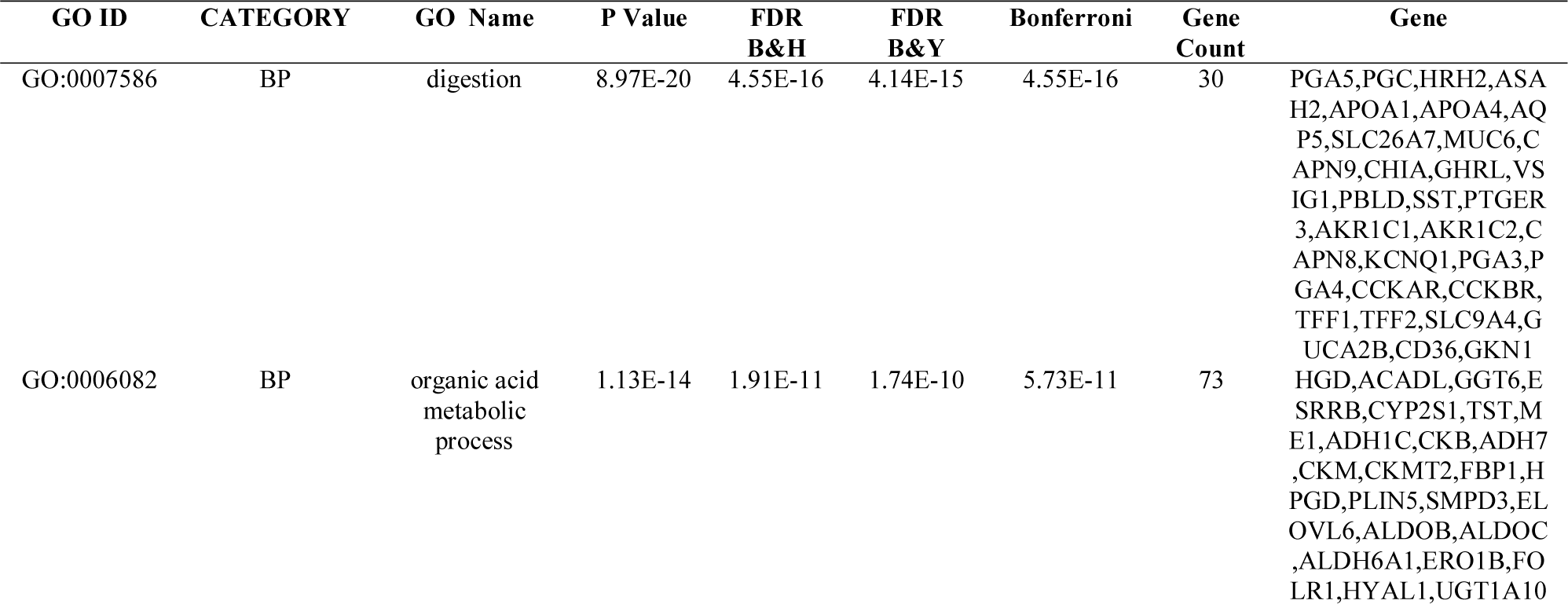
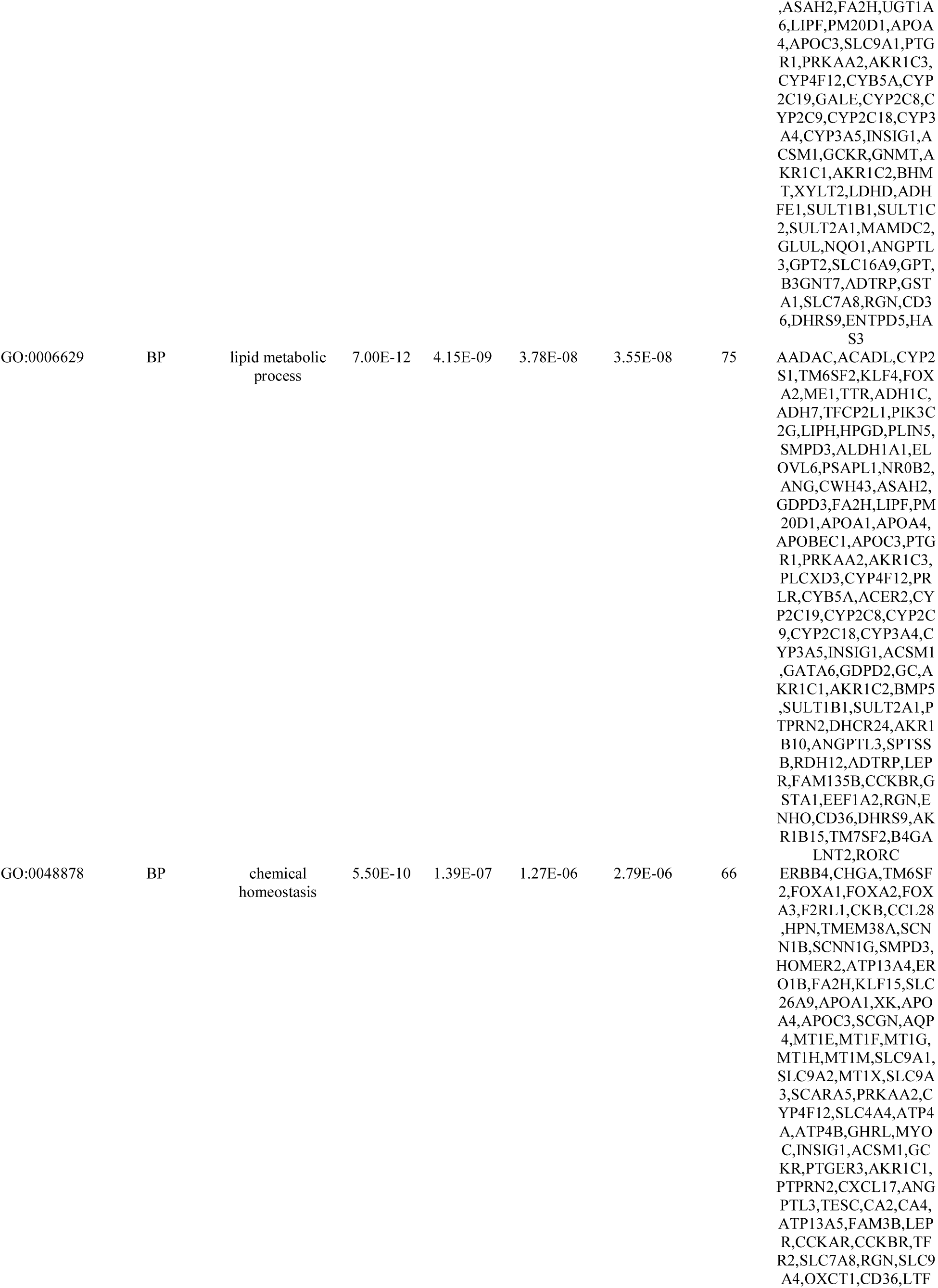
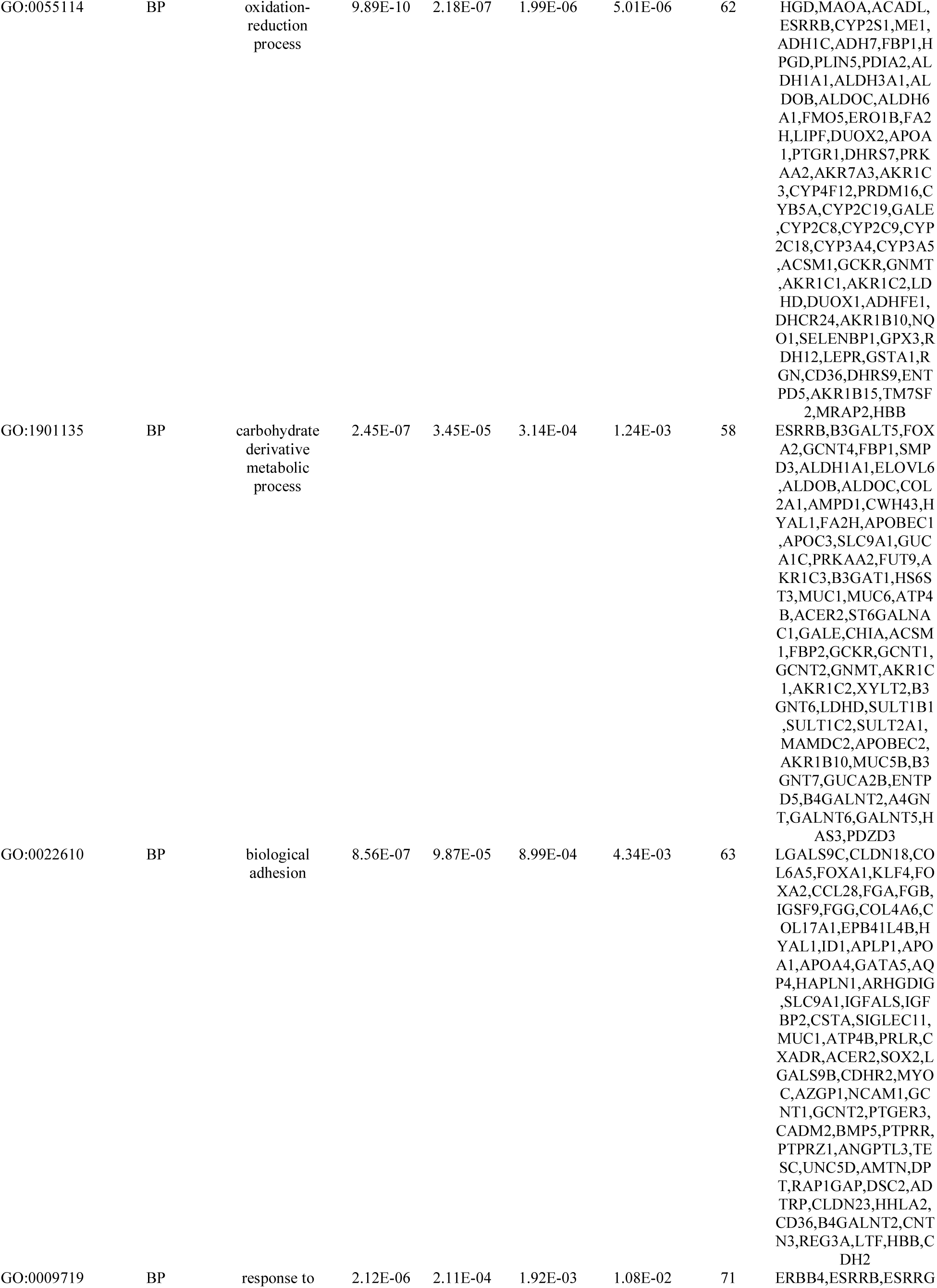
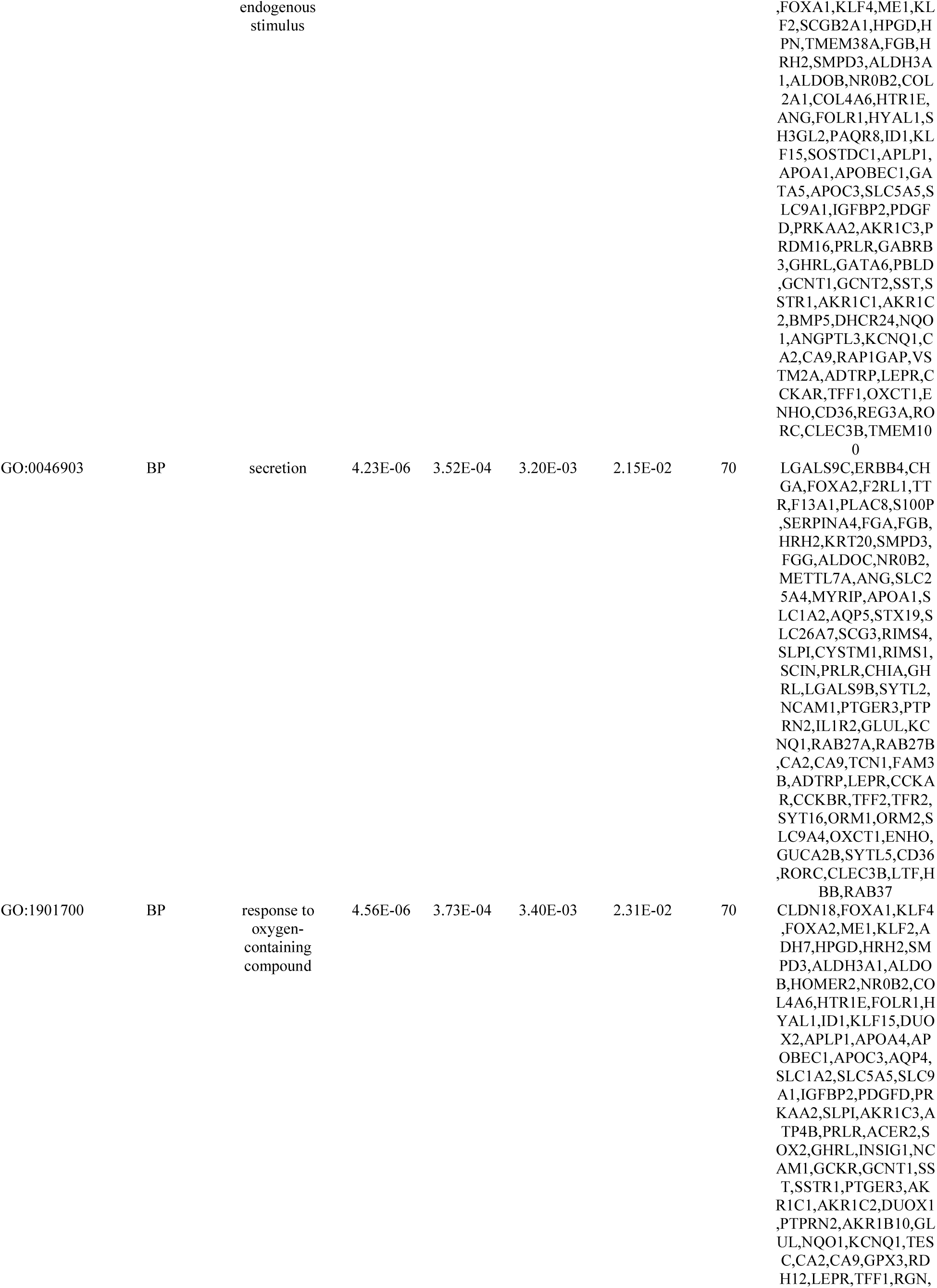
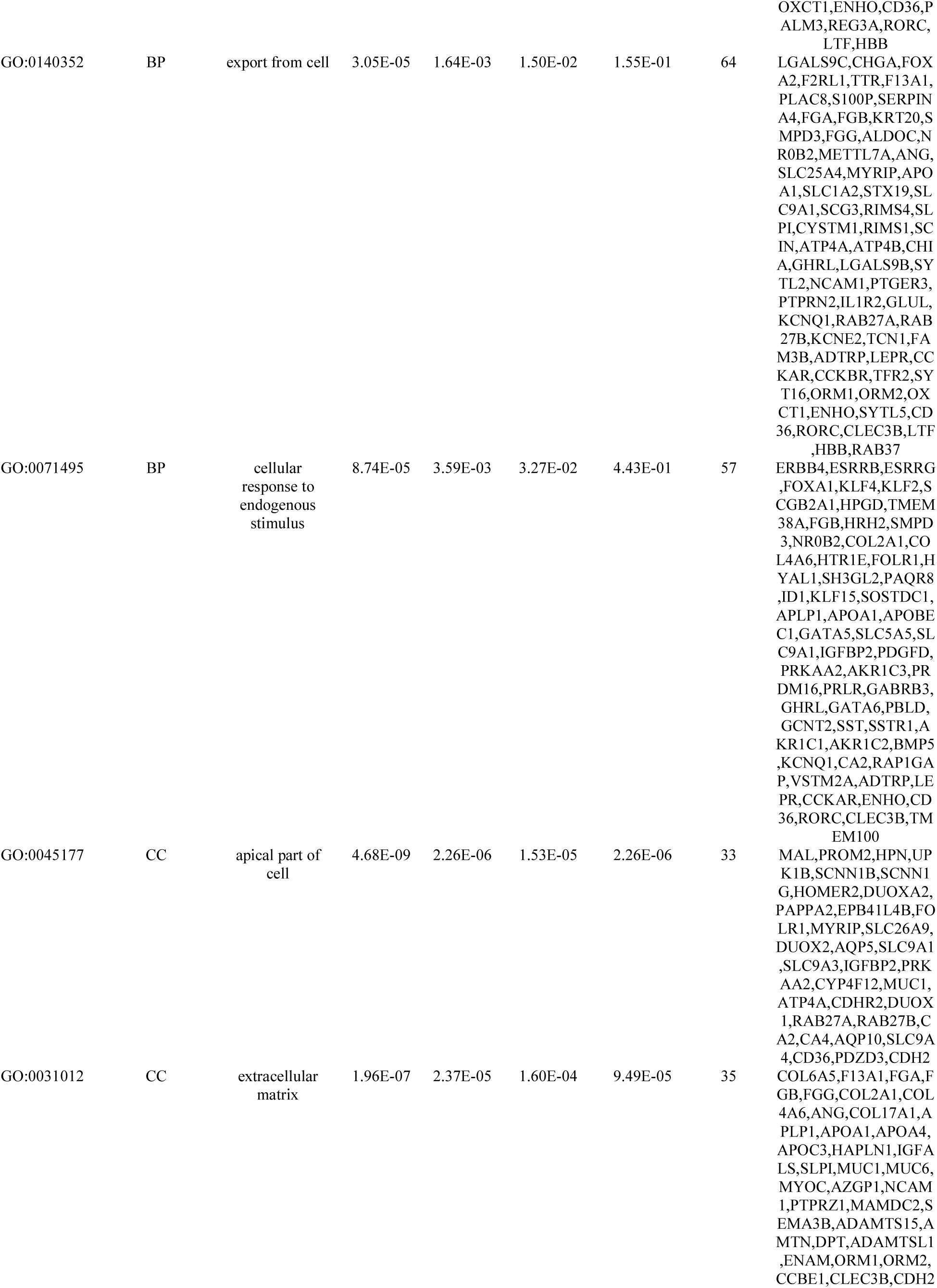
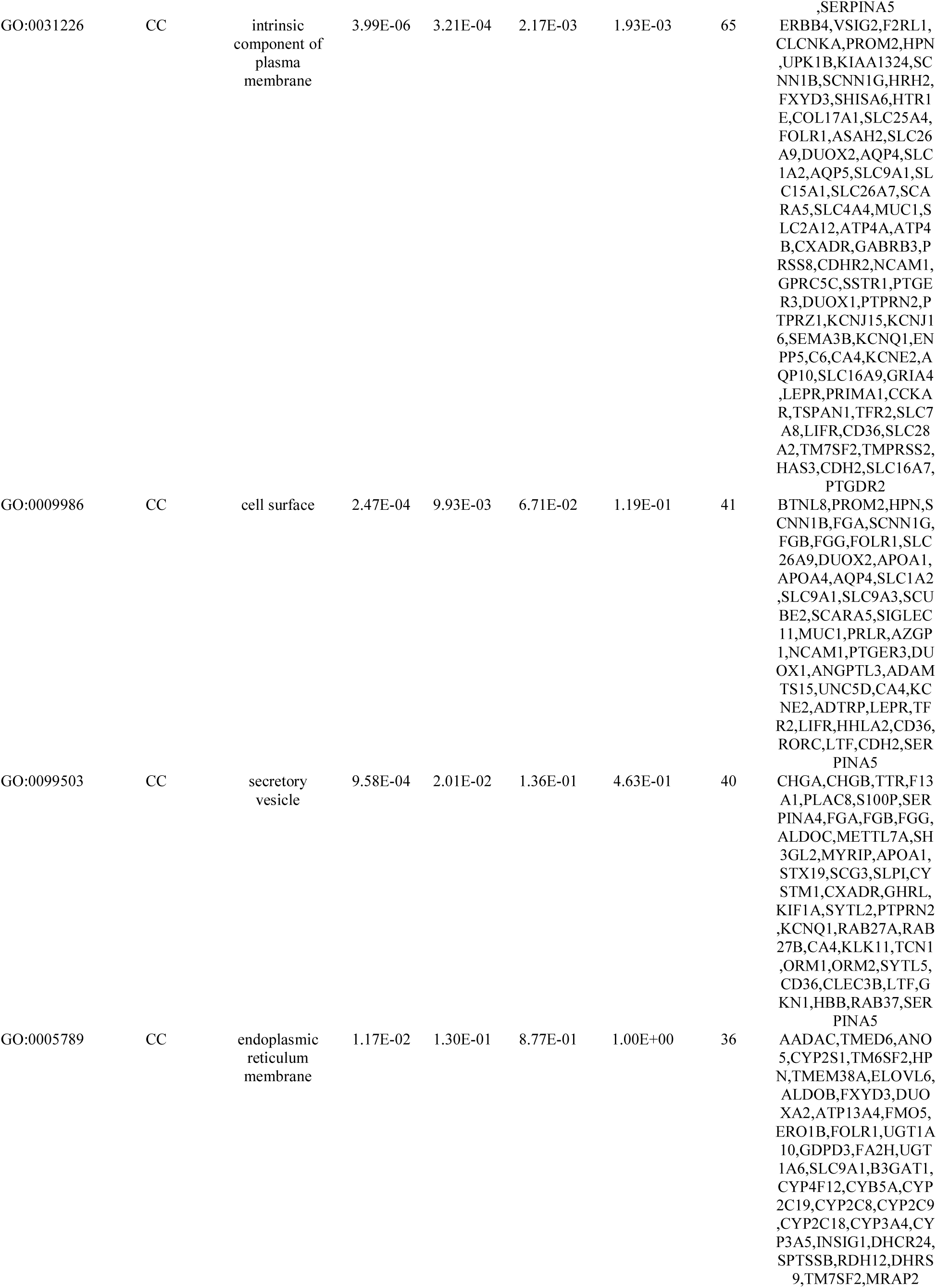
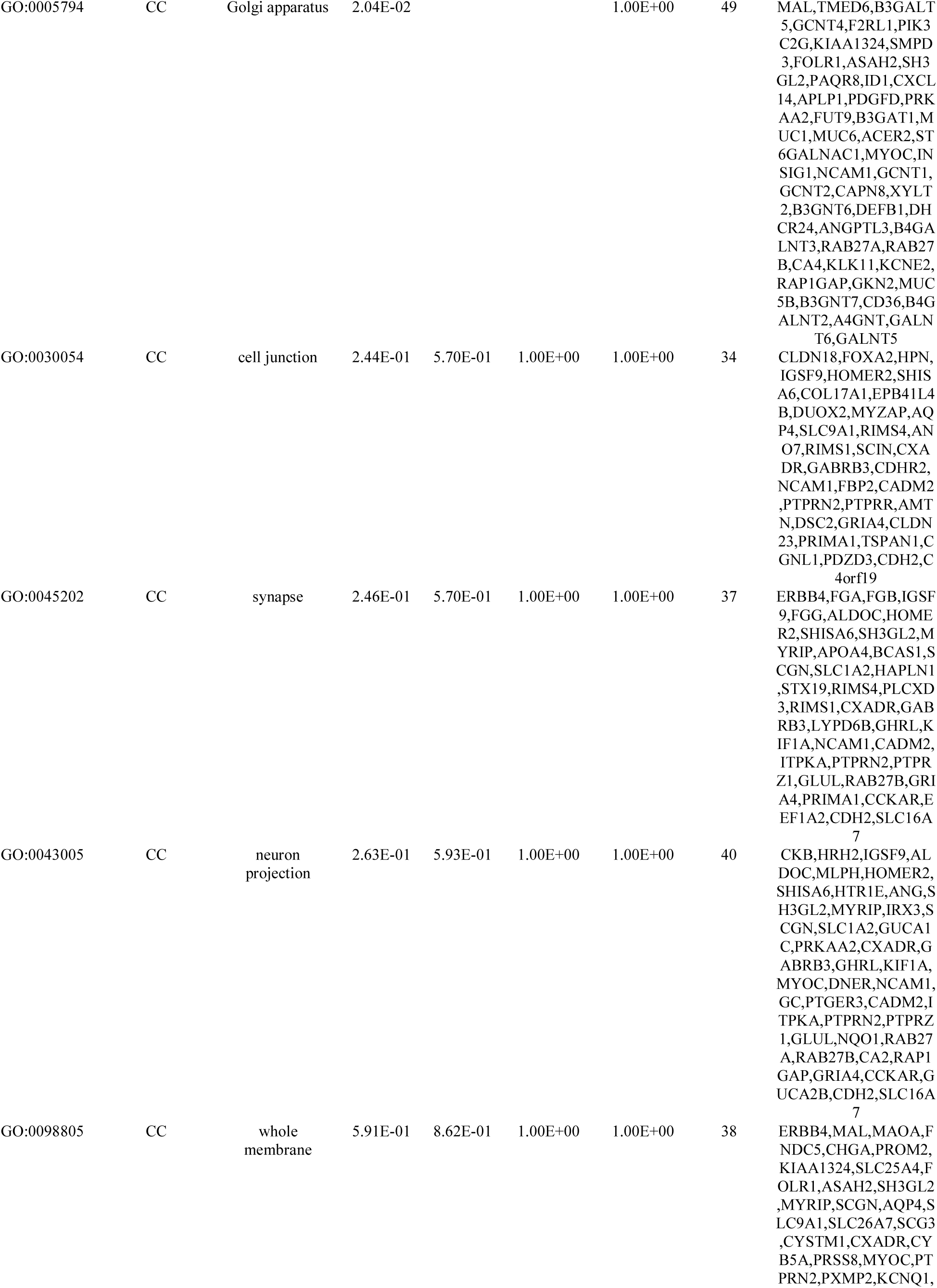
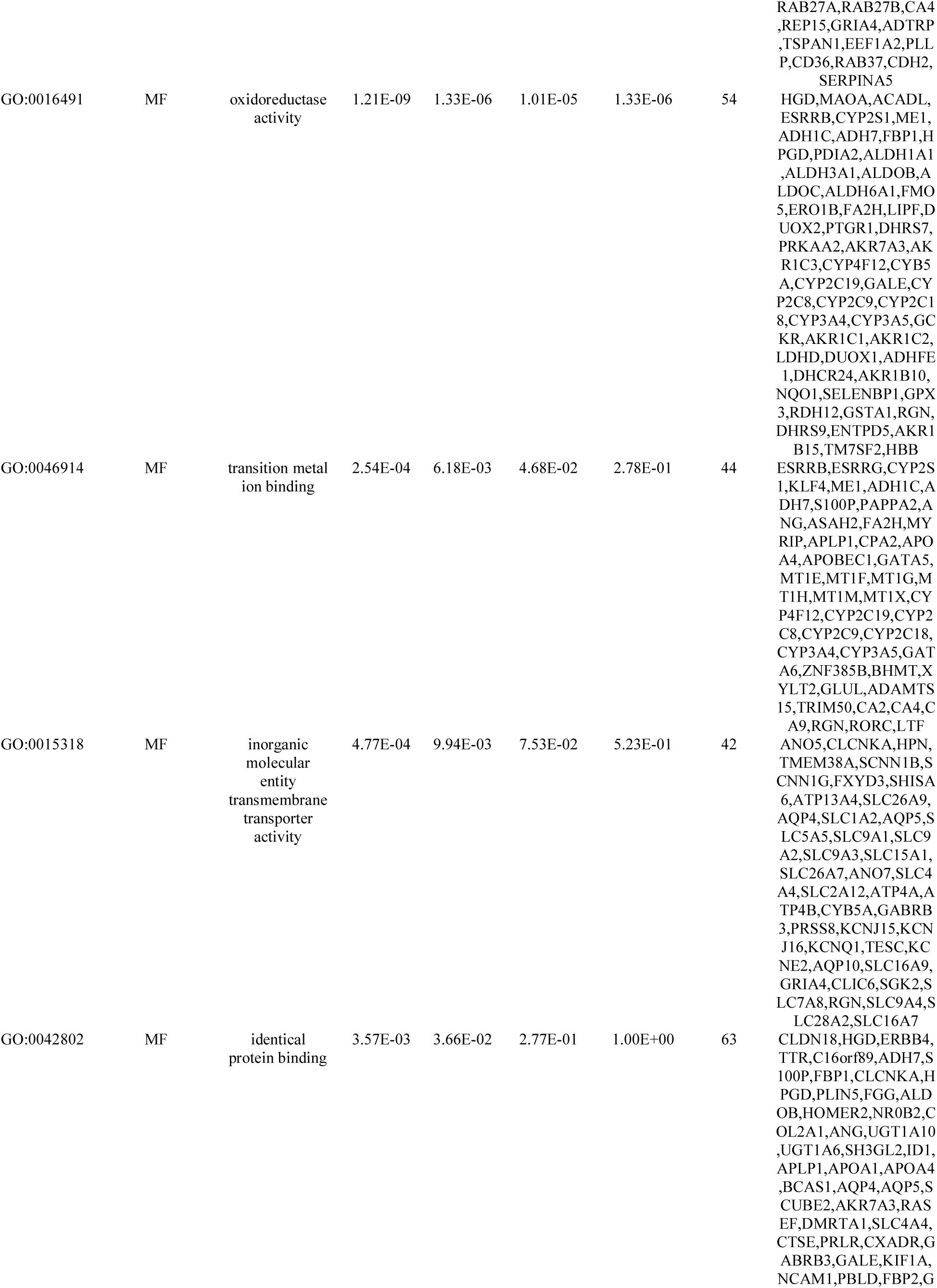
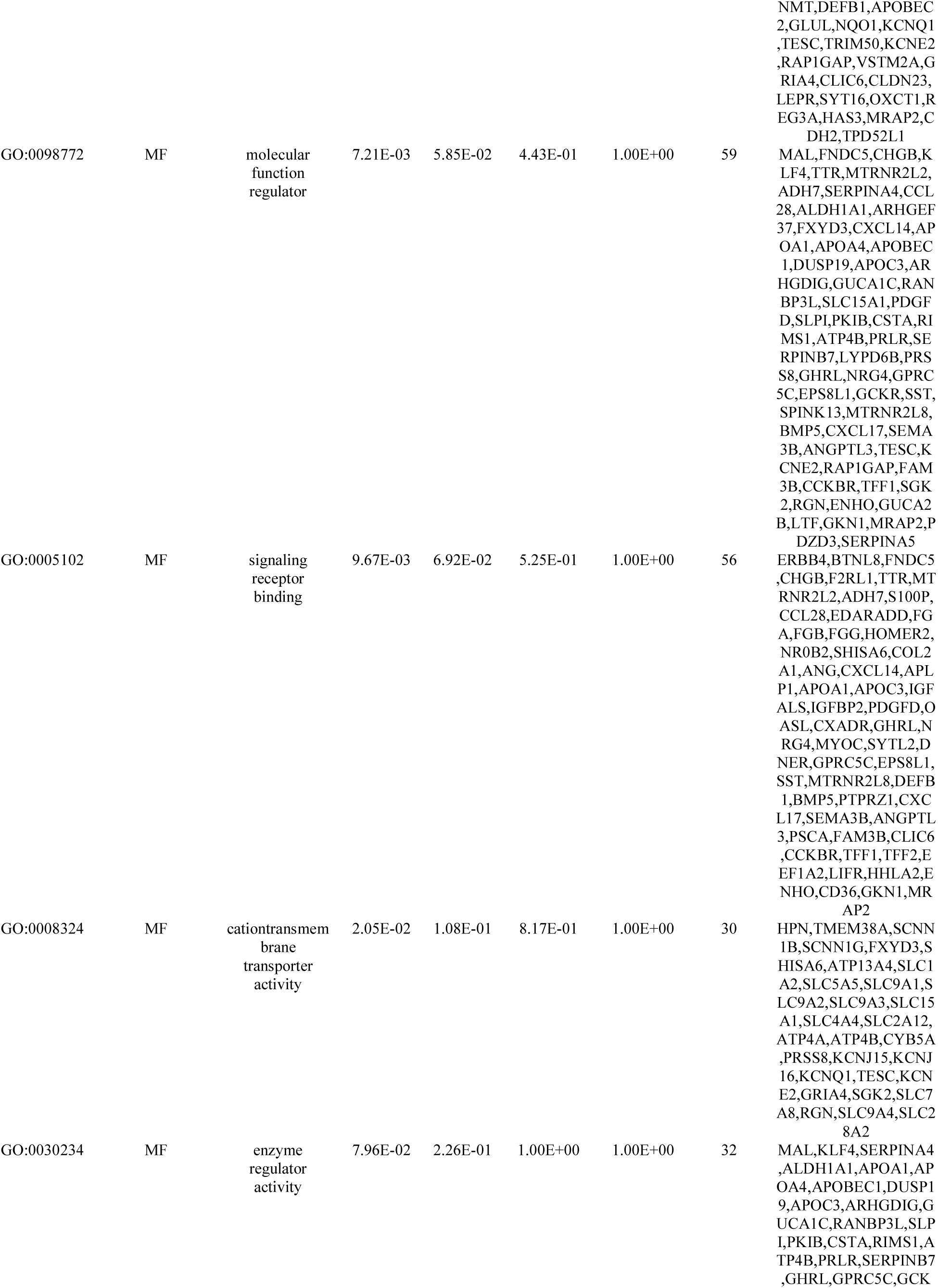
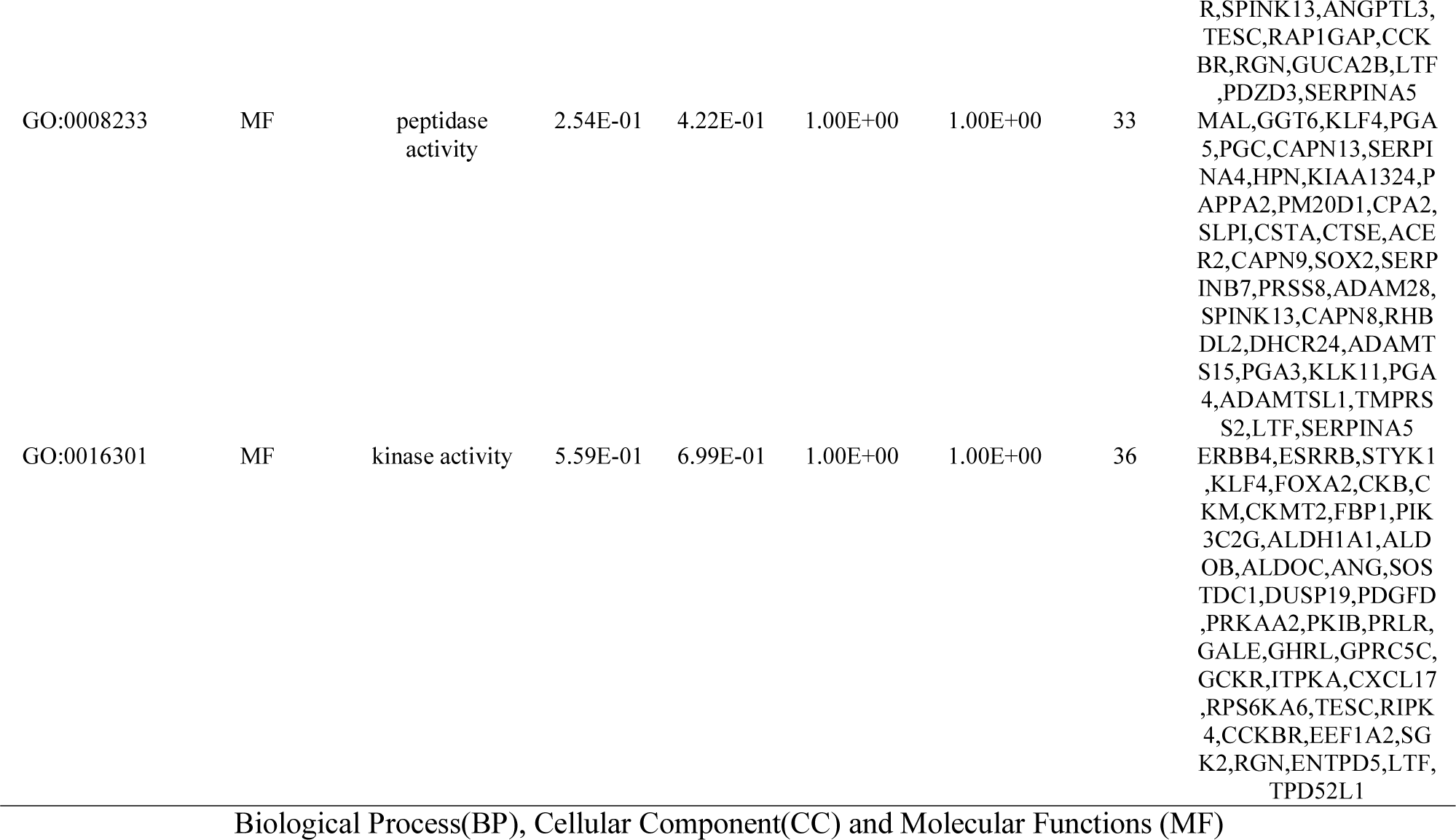
The enriched GO terms of the down regulated differentially expressed genes

### PPI network construction and module analysis

All the up and down regulated genes were entered into the HIPPIE database to obtain the interaction data. The protein-protein interaction (PPI) network of up regulated genes exhibited 3222 nodes and 4943 edges and is shown in Fig 5. The hub genes with highest node degree distribution, betweenness centrality, stress centrality, closeness centrality and lowest clustring coefficient were as follows: FN1, NOTCH1, PLK1, ANLN, MDFI, CPZ, PRKDC, TREM1, TREM2, CYP2W1, CST1, CCL3, PLXDC2, AMIGO2 and GPR161, and are listed in Table 7. The statistical scores of node degree distribution, betweenness centrality, stress centrality, closeness centrality and clustring coefficient are shown as scatter plot in Fig. 6A - 6E. The functional enrichment analysis showed that these hub genes were significantly involved in ECM-receptor interaction, microRNAs in cancer, PLK1 signaling events, cell migration, animal organ morphogenesis, extracellular matrix, phosphotransferase activity, alcohol group as acceptor, molecular transducer activity, axon guidance, endoplasmic reticulum lumen, peptidase activity, identical protein binding, ensemble of genes encoding extracellular matrix and extracellular matrix-associated proteins, cell adhesion and whole membrane. Similarly, protein-protein interaction (PPI) network of down regulated genes exhibited 5142 nodes and 8412 edges and is shown in Fig 7. The hub genes with highest node degree distribution, betweenness centrality, stress centrality, closeness centrality and lowest clustring coefficient were as follows: SOX2, RIPK4, FOXA1, EEF1A2, ANG, FGB, CNTD1, GGT6, PTGDR2, ADH1C, DCAF12L1, CAPN9 and TTC39A, and are listed in Table 7. The statistical scores of node degree distribution, betweenness centrality, stress centrality, closeness centrality and clustring coefficient are shown as scatter plot in Fig. 8A - 8E. The functional enrichment analysis showed that these hub genes were significantly involved in developmental biology, kinase activity, FOXA2 and FOXA3 transcription factor networks, metabolism of proteins, ensemble of genes encoding extracellular matrix and extracellular matrix-associated proteins, hemostasis, metabolic pathways, intrinsic component of plasma membrane, drug metabolism - cytochrome P450 and digestion.

**Fig. 5.**
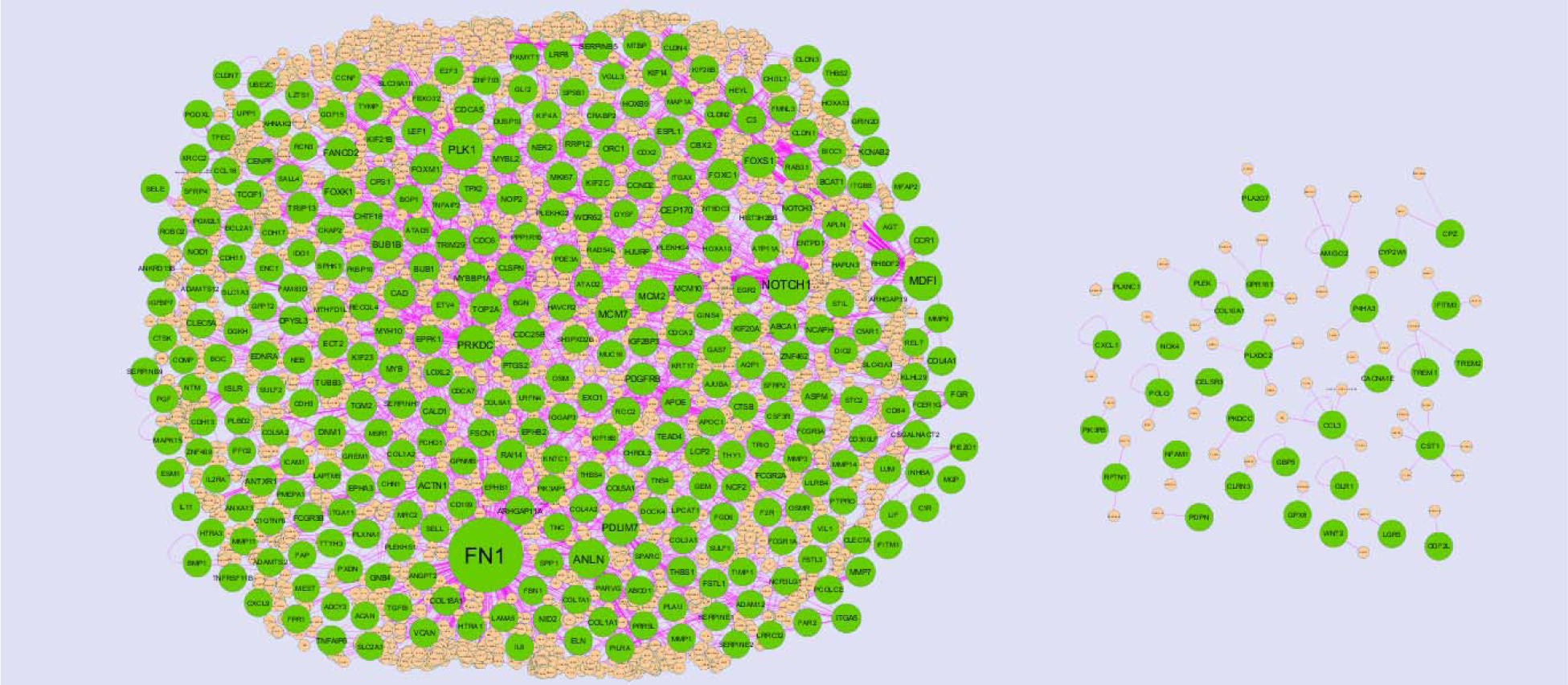
Protein–protein interaction network of up regulated genes. Green nodes denotes up regulated genes.

**Fig. 6.**
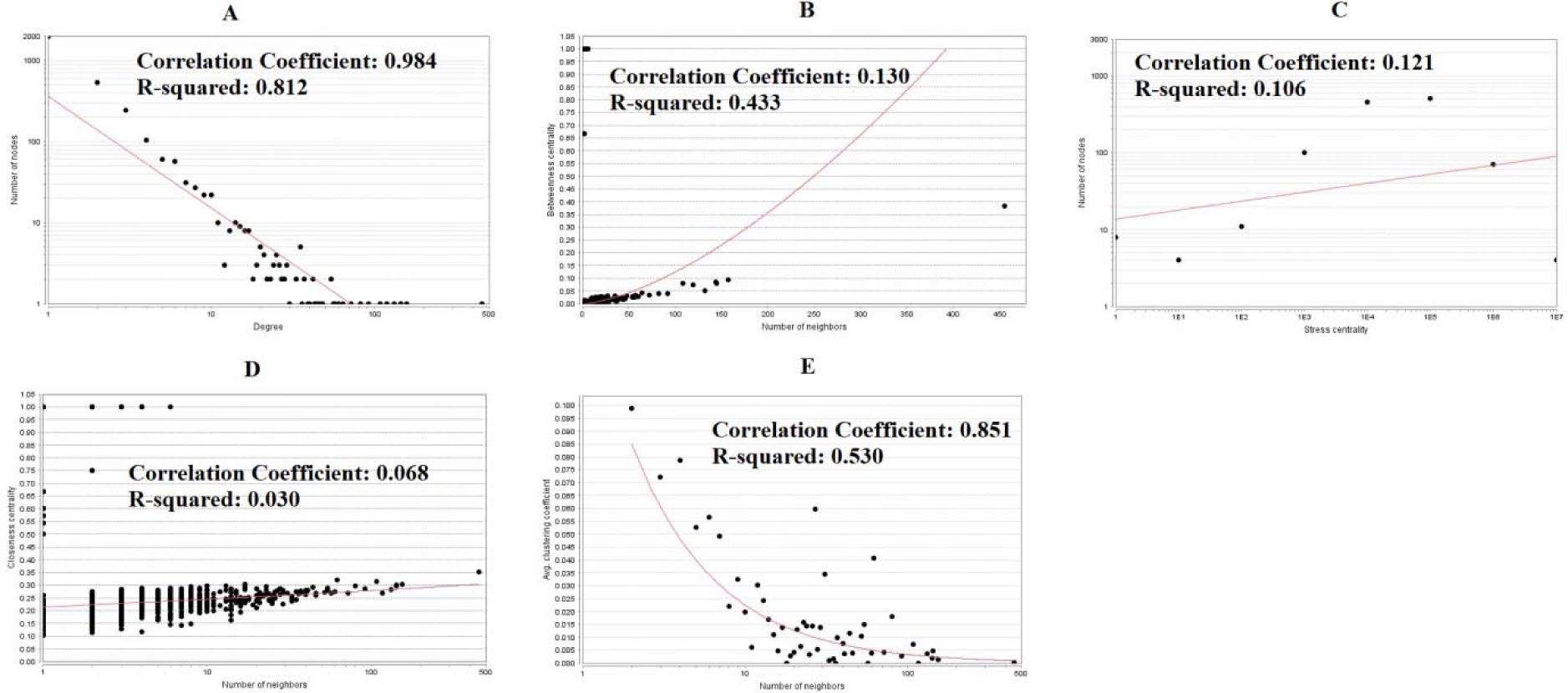
Scatter plot for up regulated genes. (A- Node degree; B- Betweenness centrality; C- Stress centrality; D- Closeness centrality; E- Clustering coefficient)

**Fig. 7.**
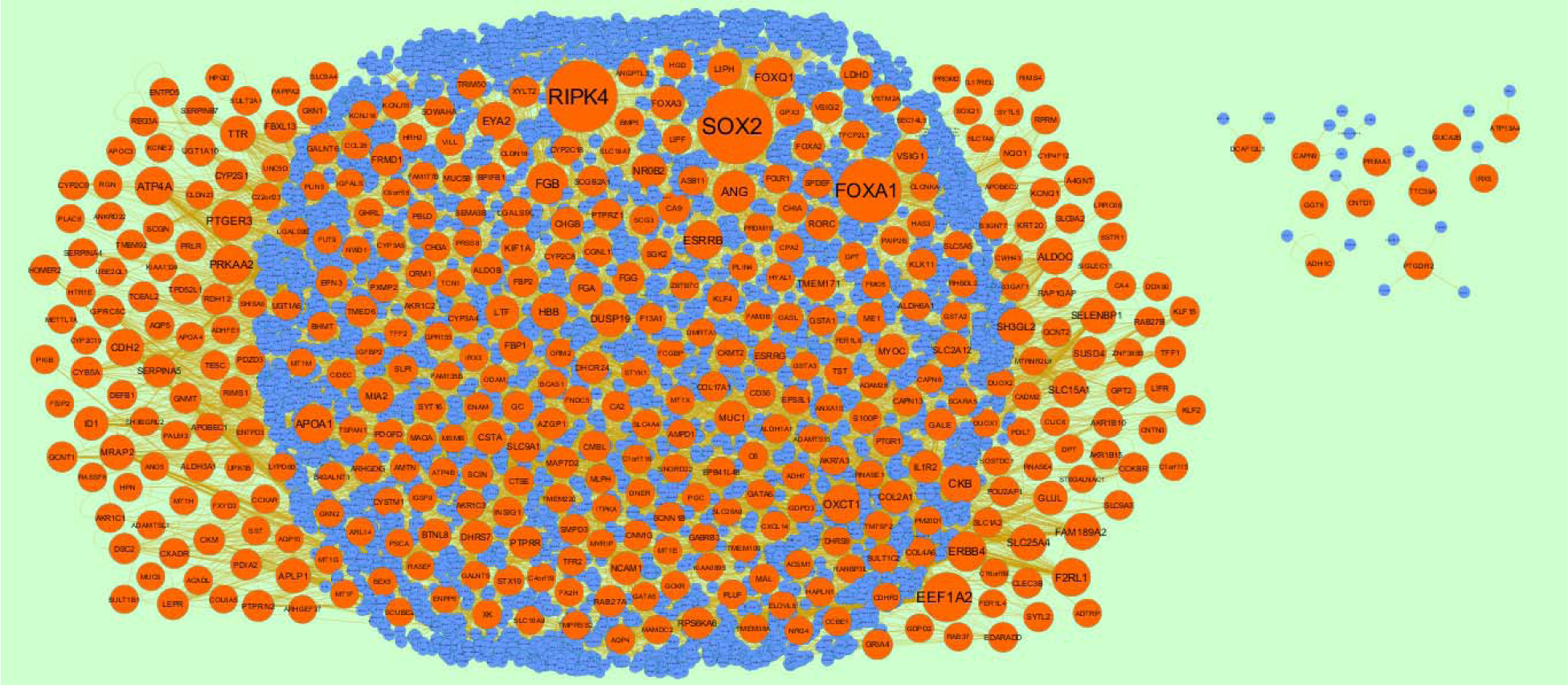
Protein–protein interaction network of down regulated genes. Red nodes denotes down regulated genes.

**Fig. 8.**
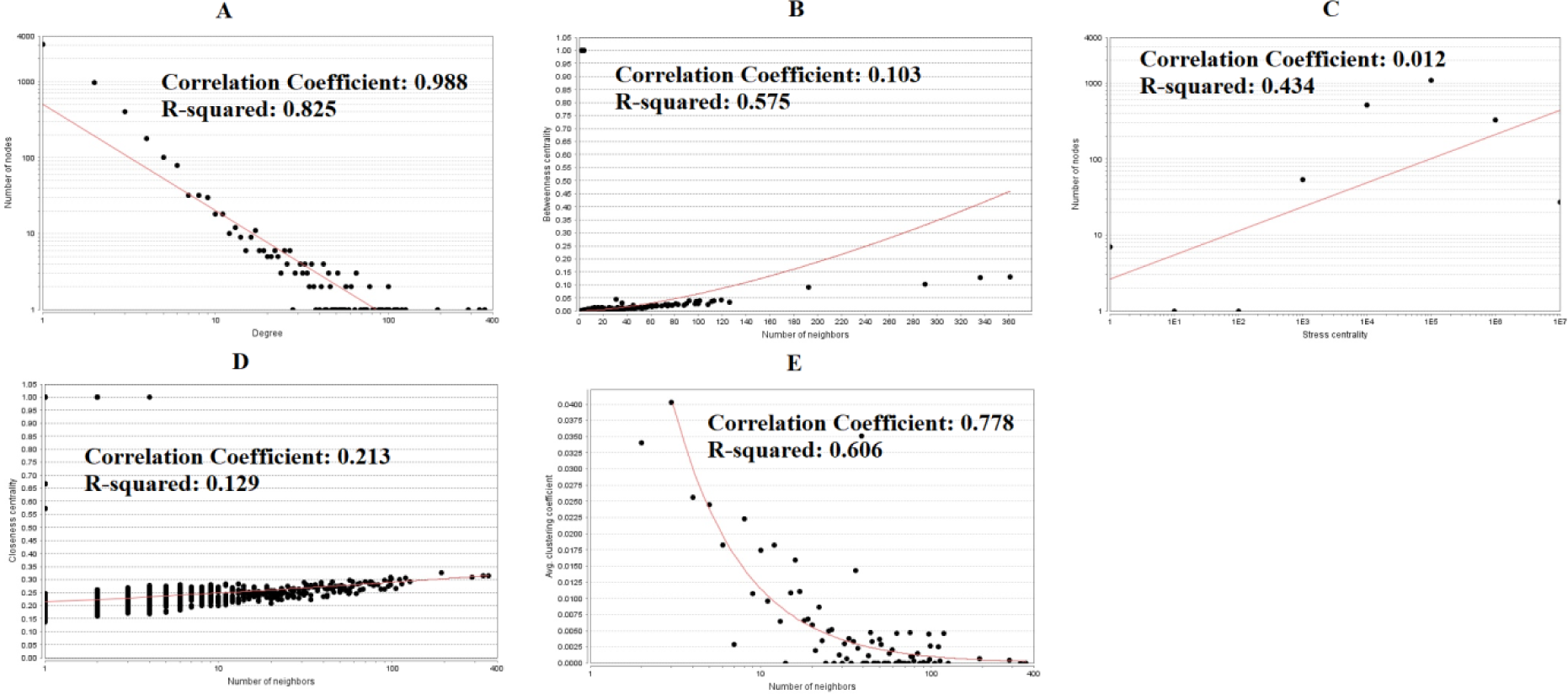
Scatter plot for down regulated genes. (A- Node degree; B- Betweenness centrality; C- Stress centrality; D- Closeness centrality; E- Clustering coefficient)

**Table 7.**
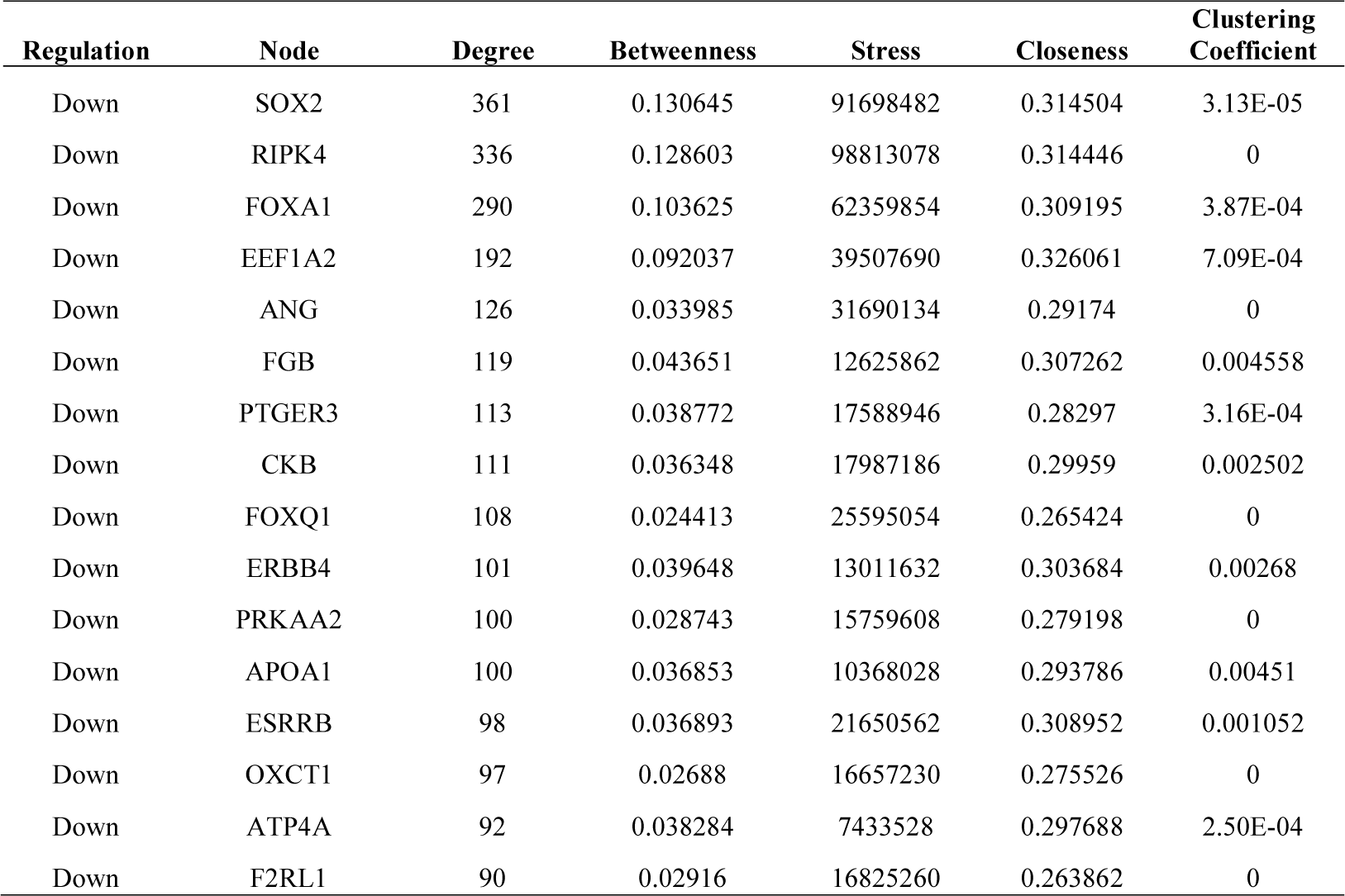
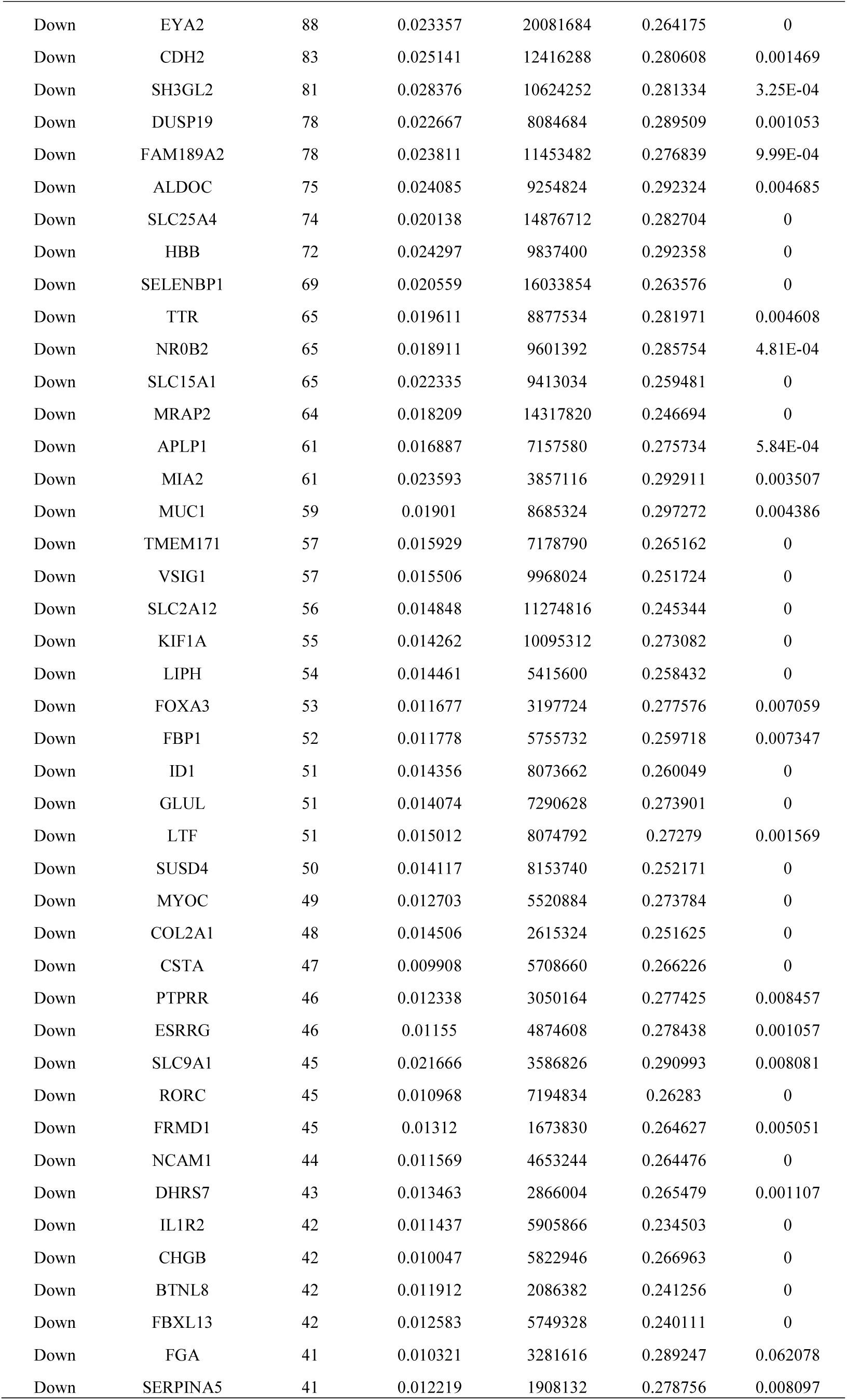
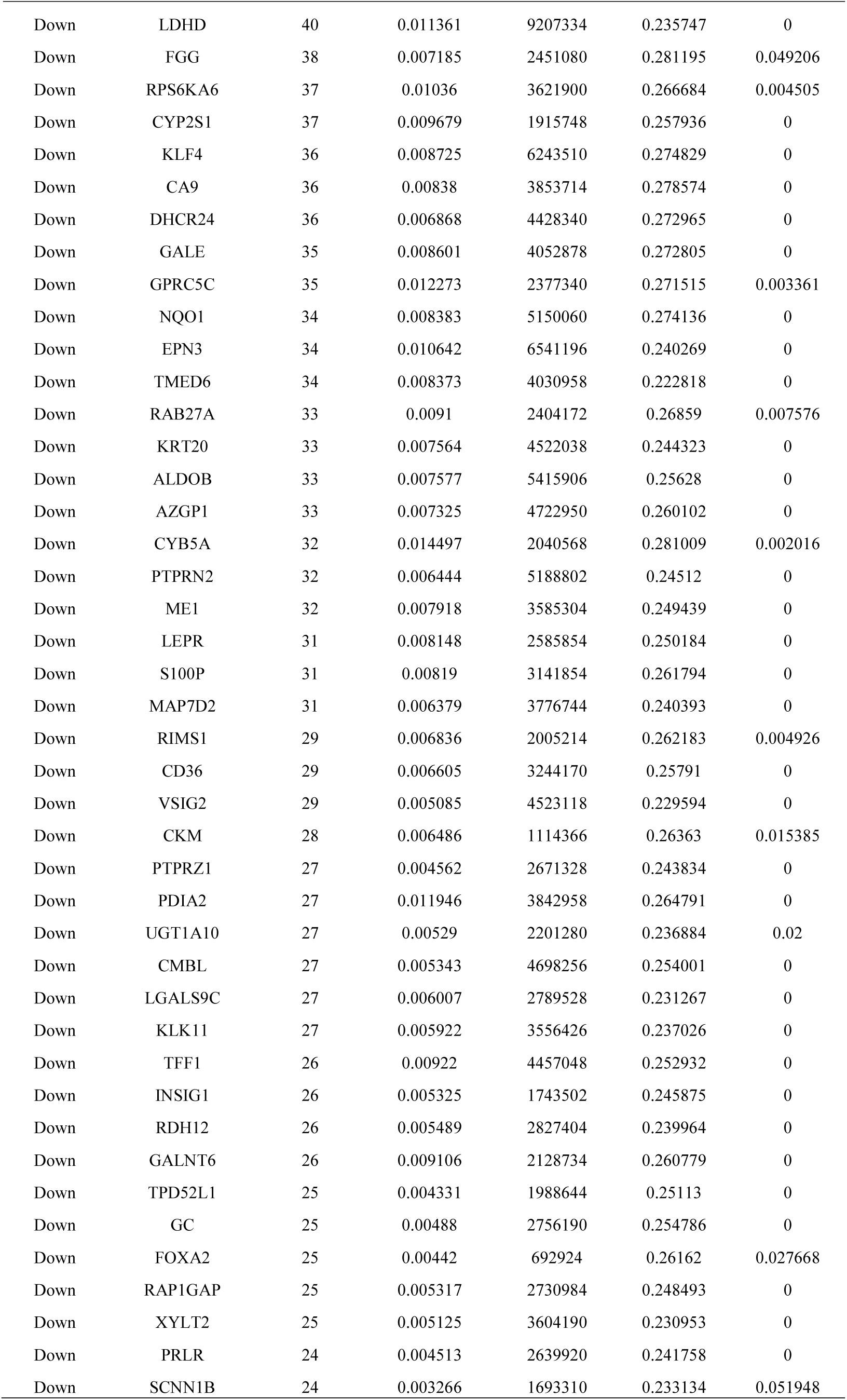
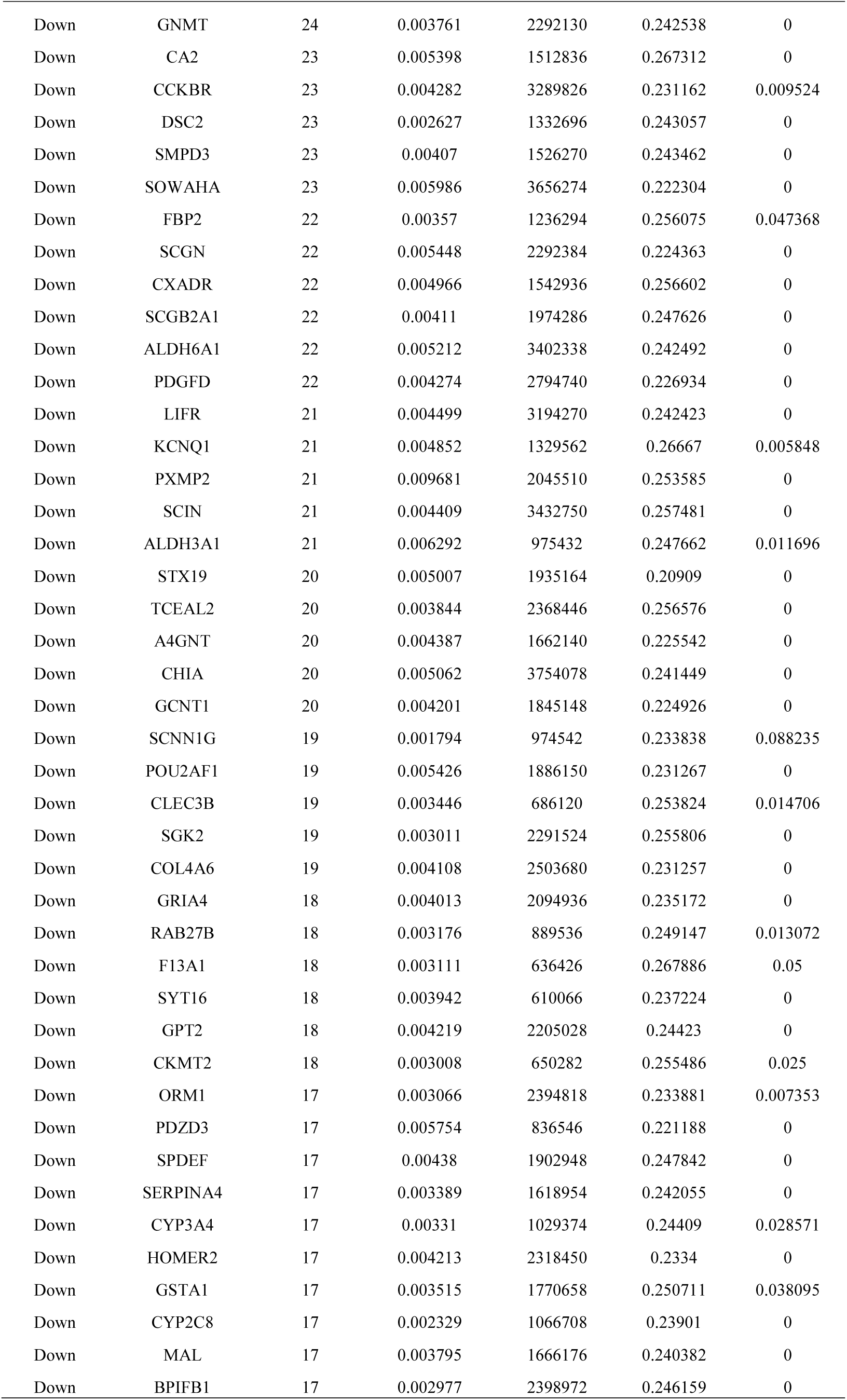
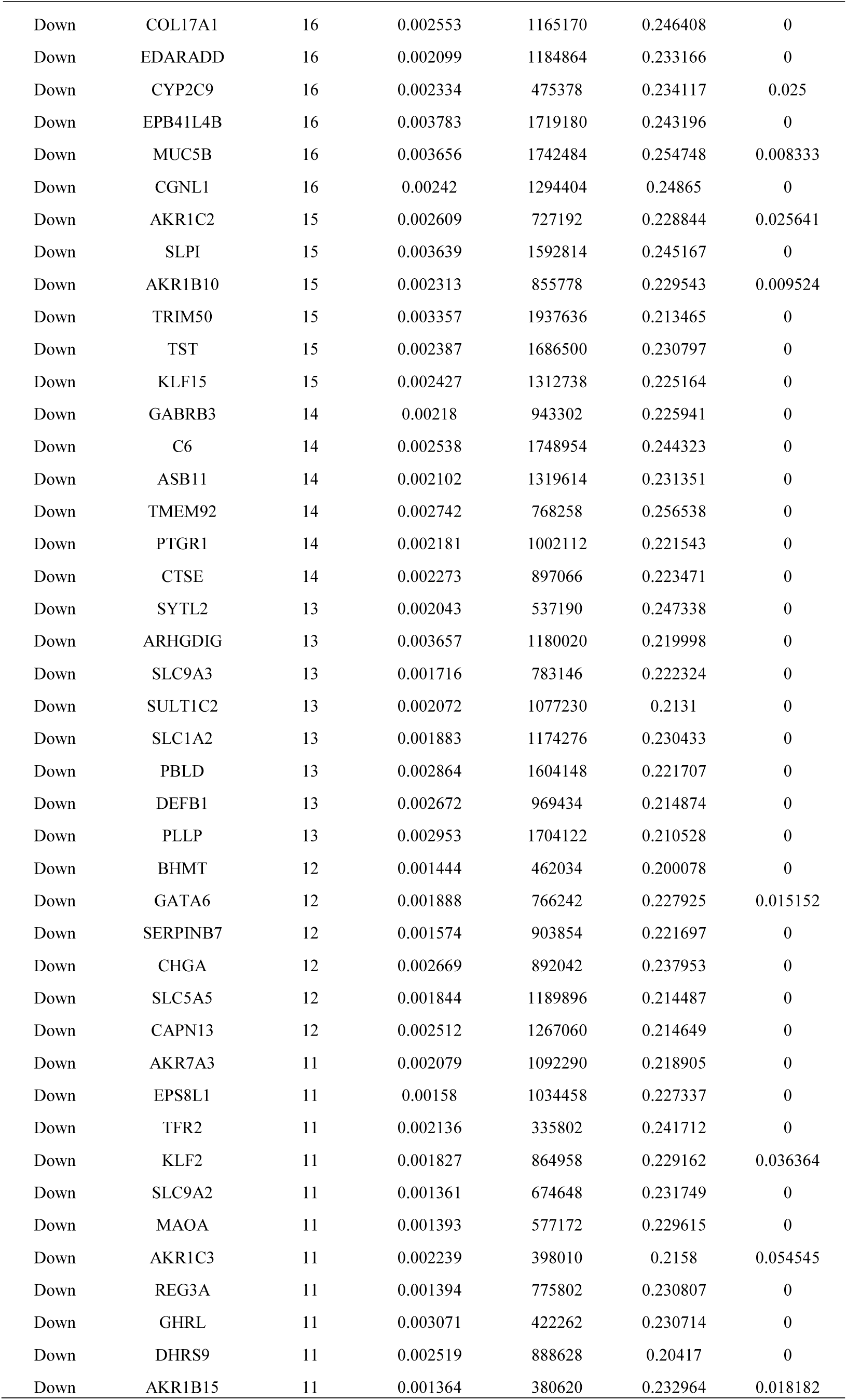
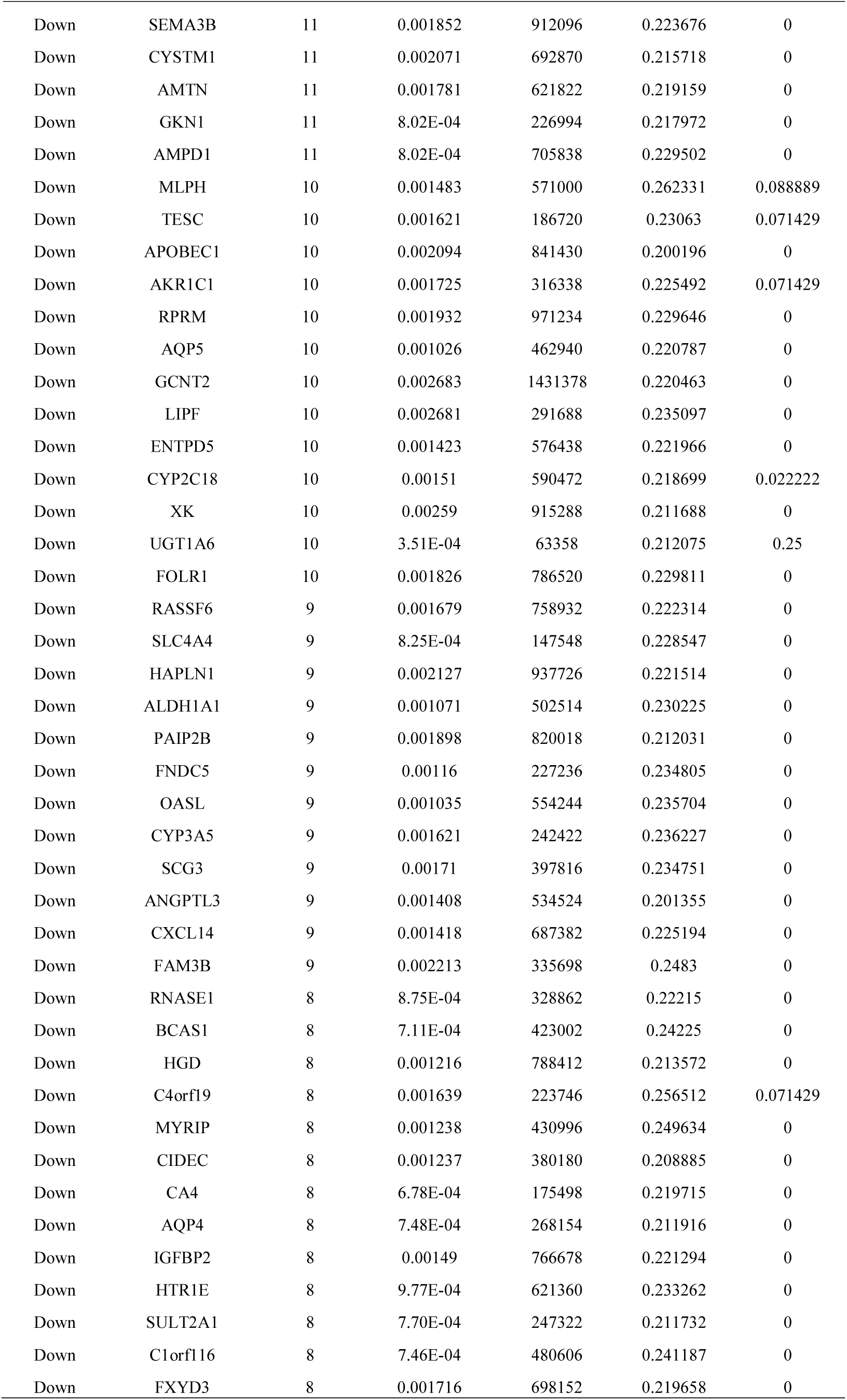
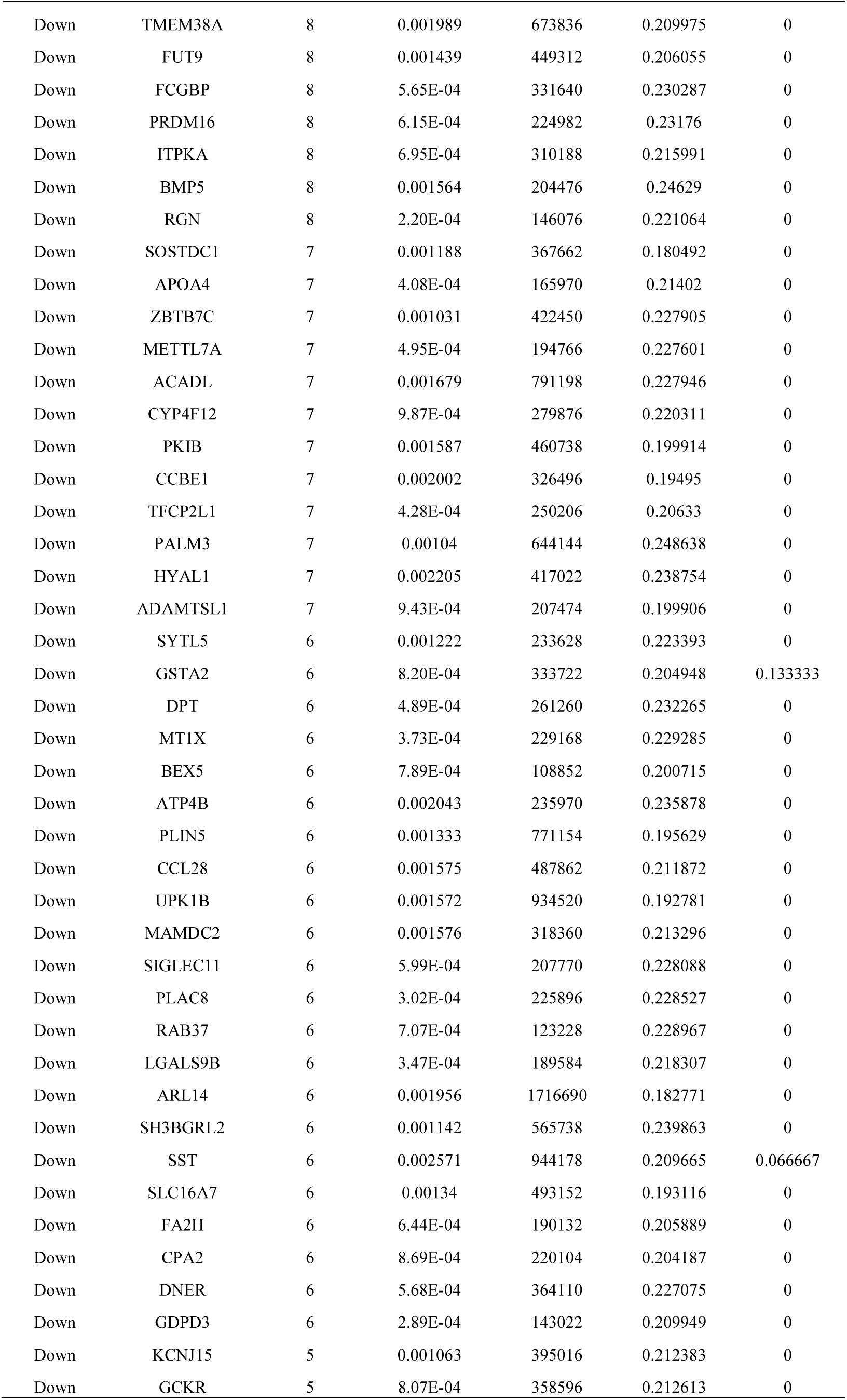
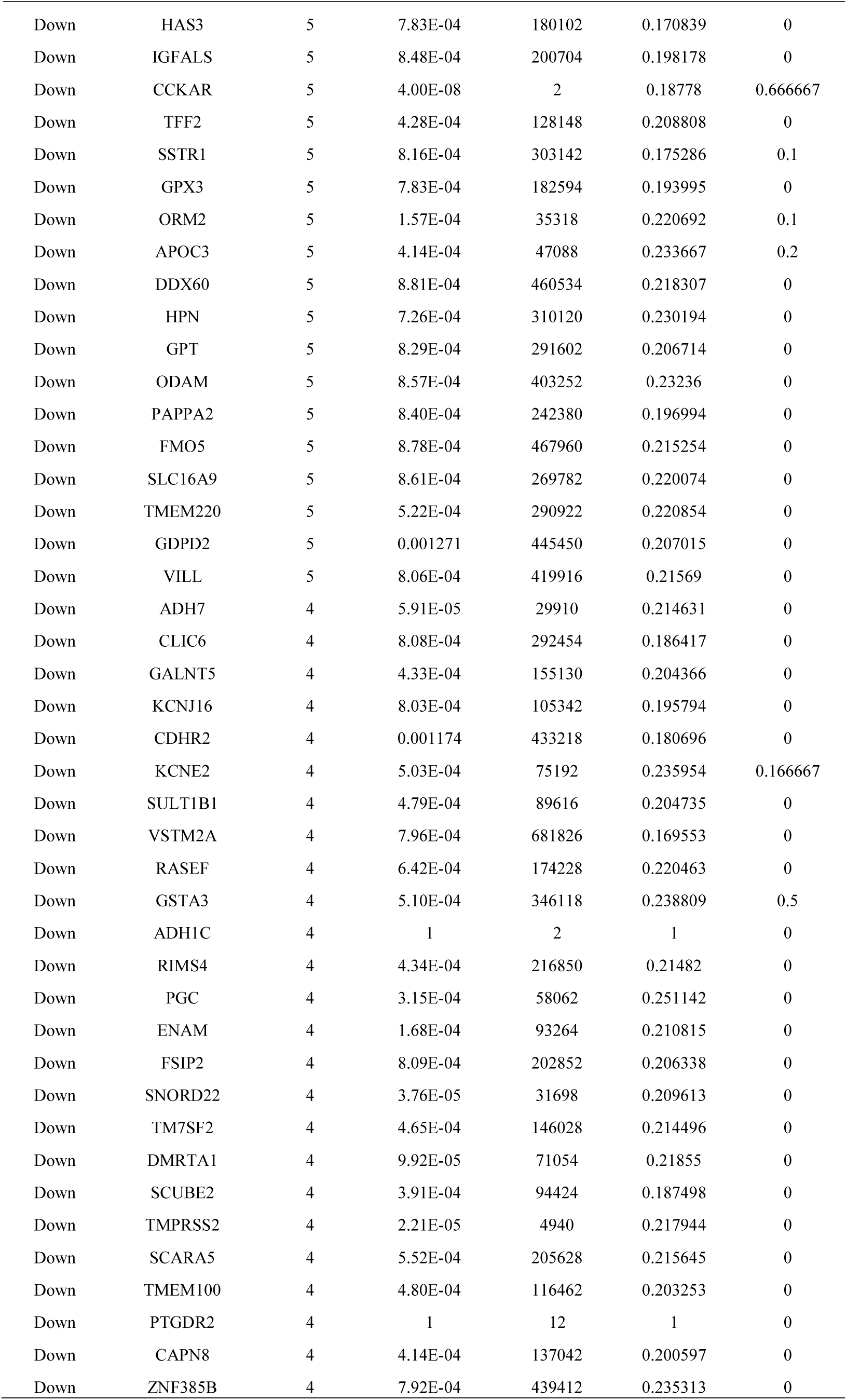
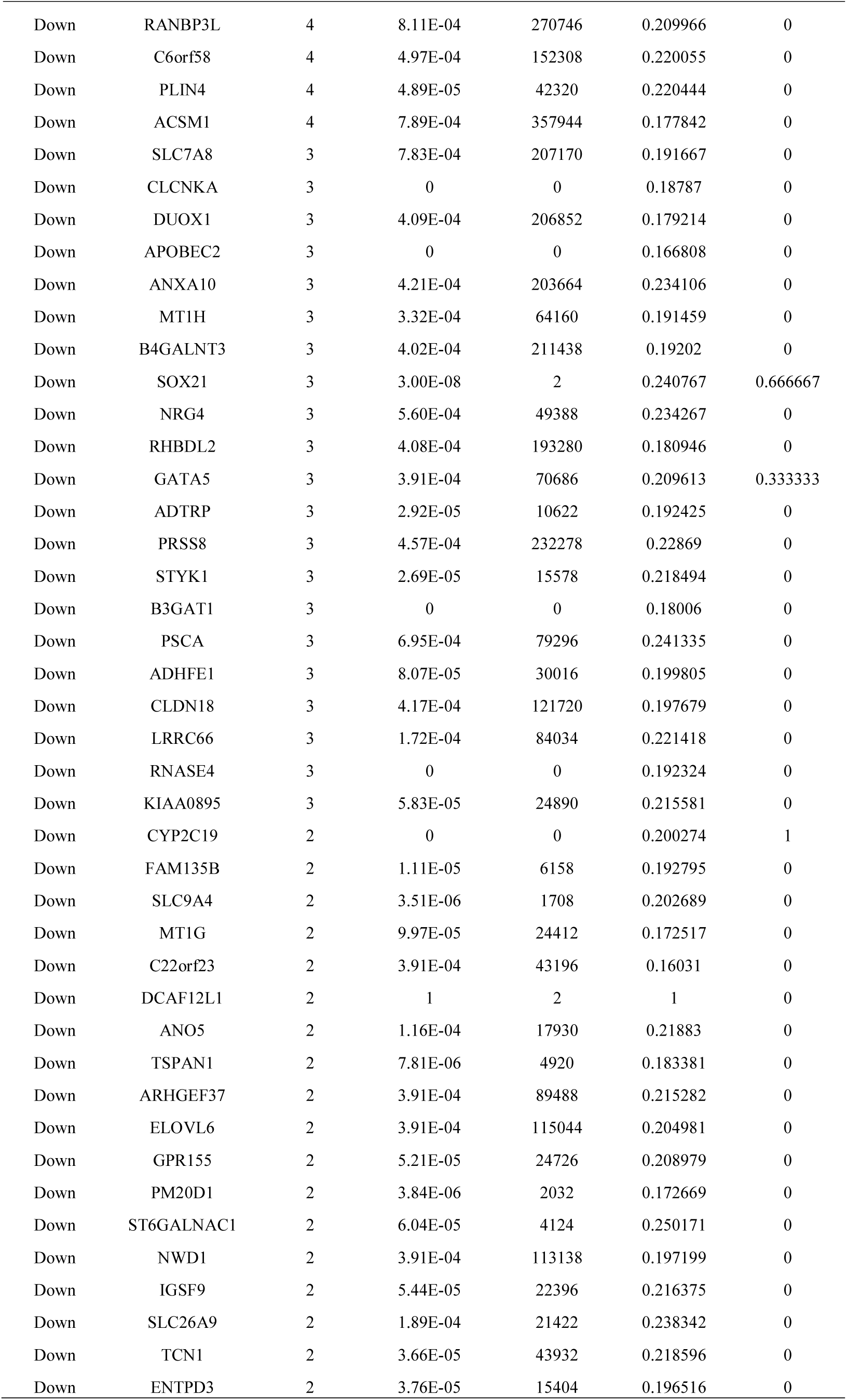
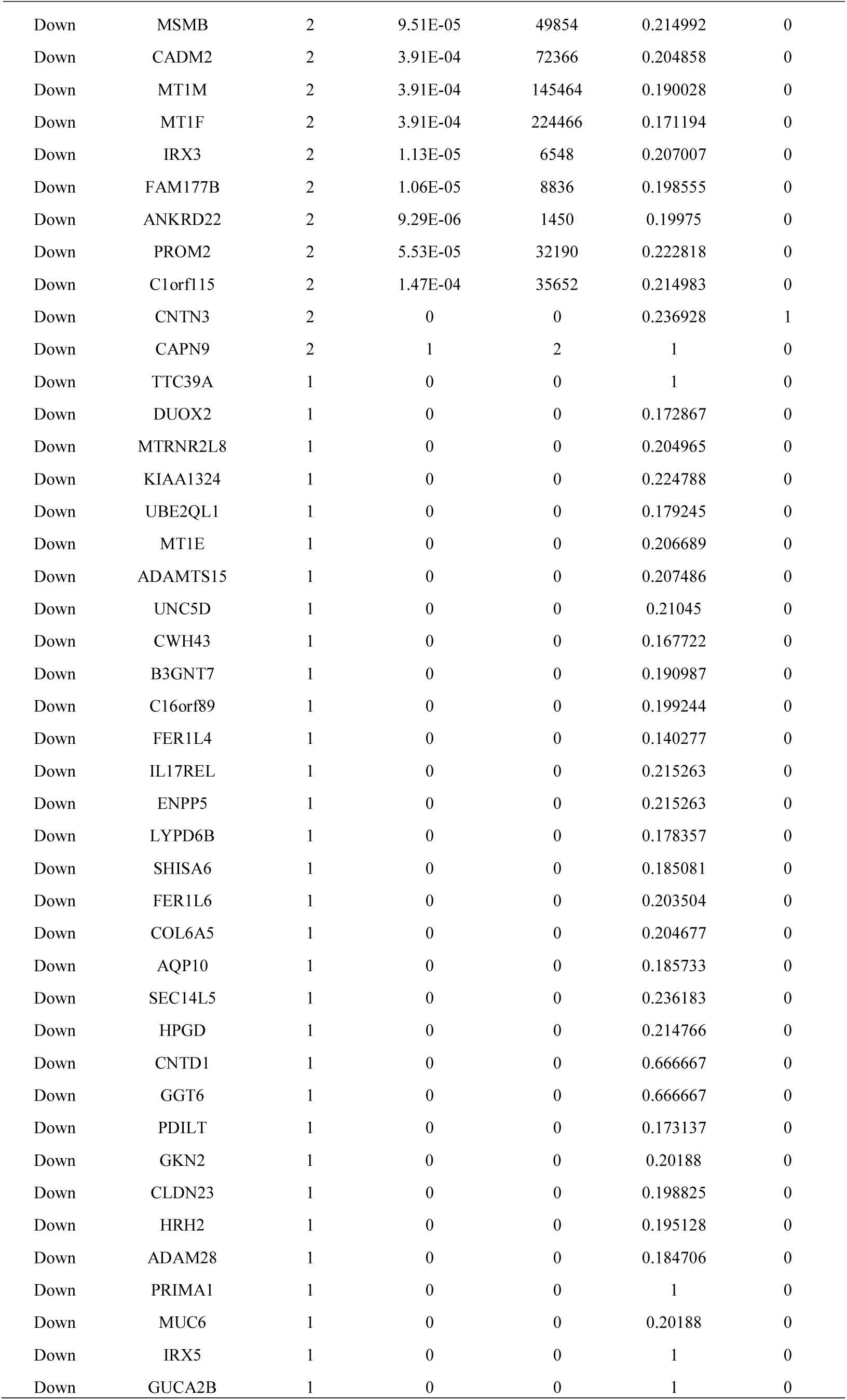
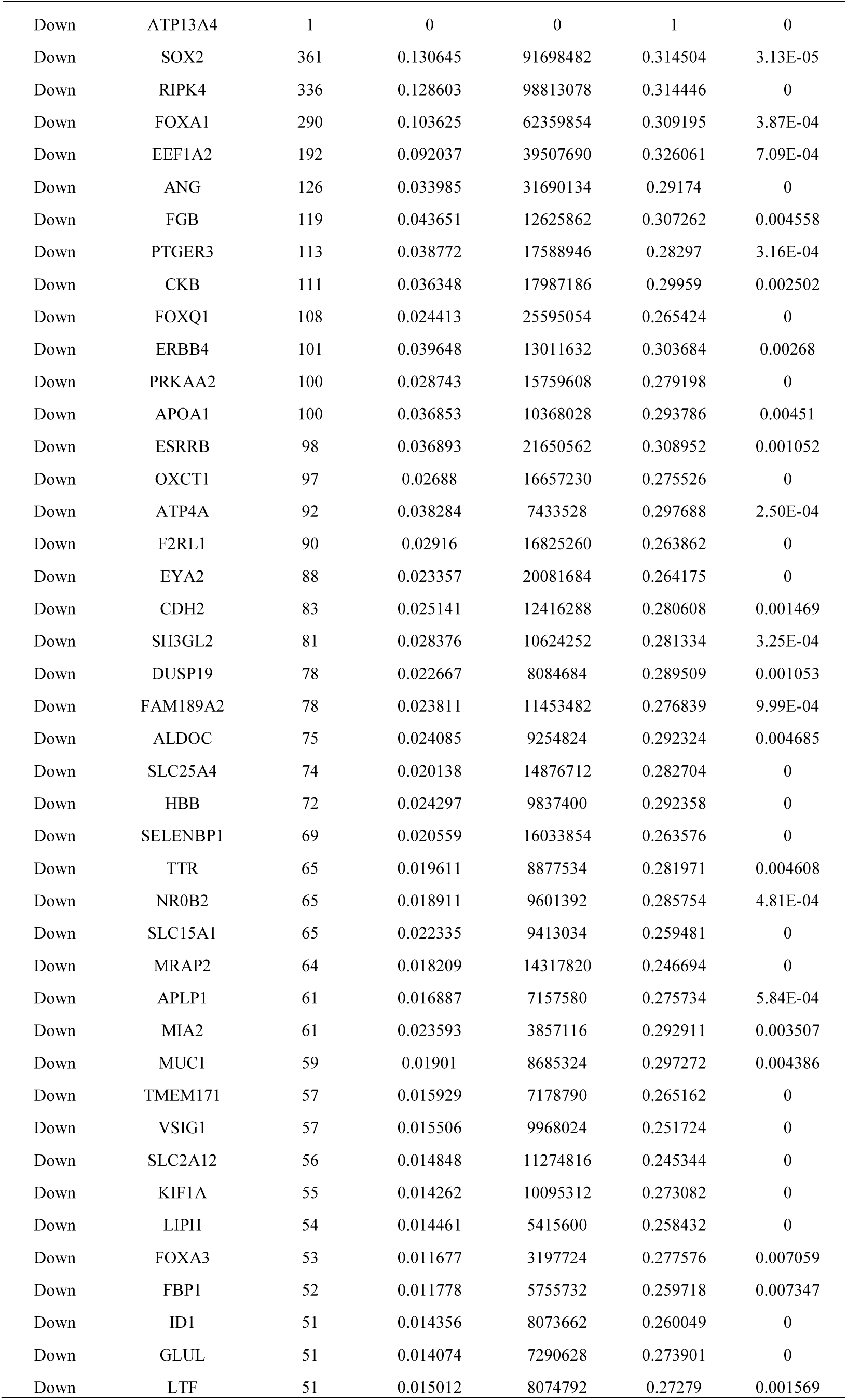
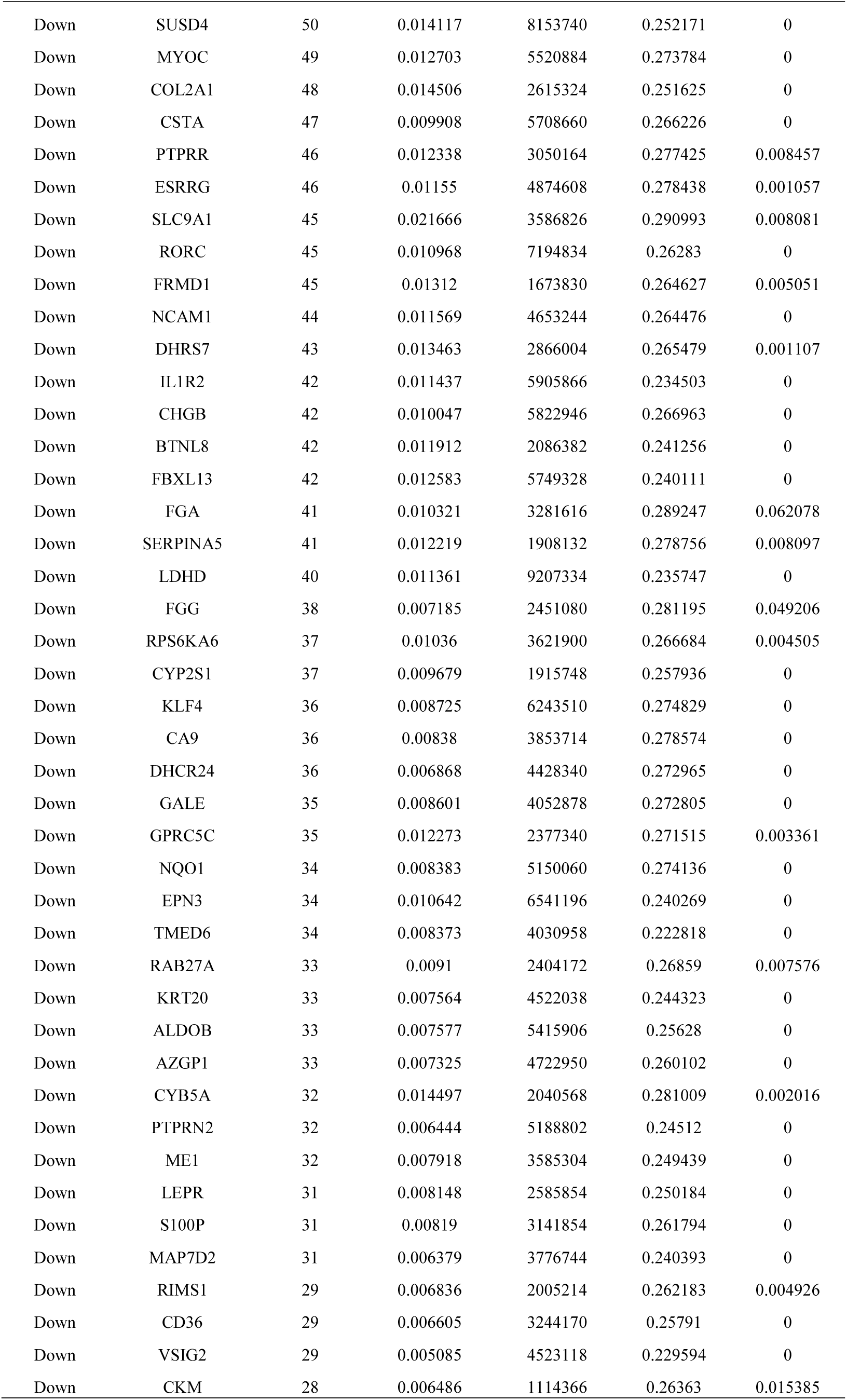
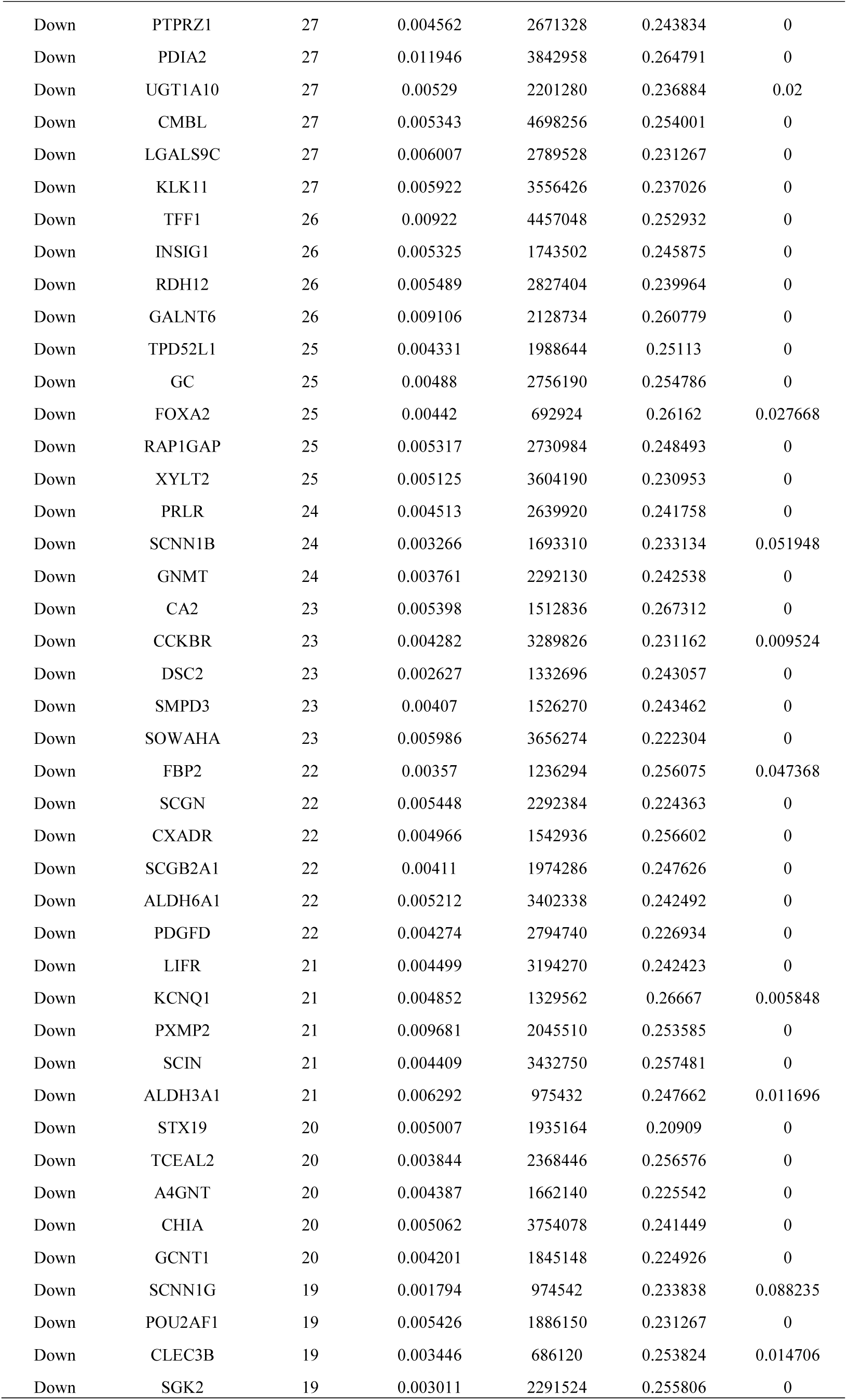
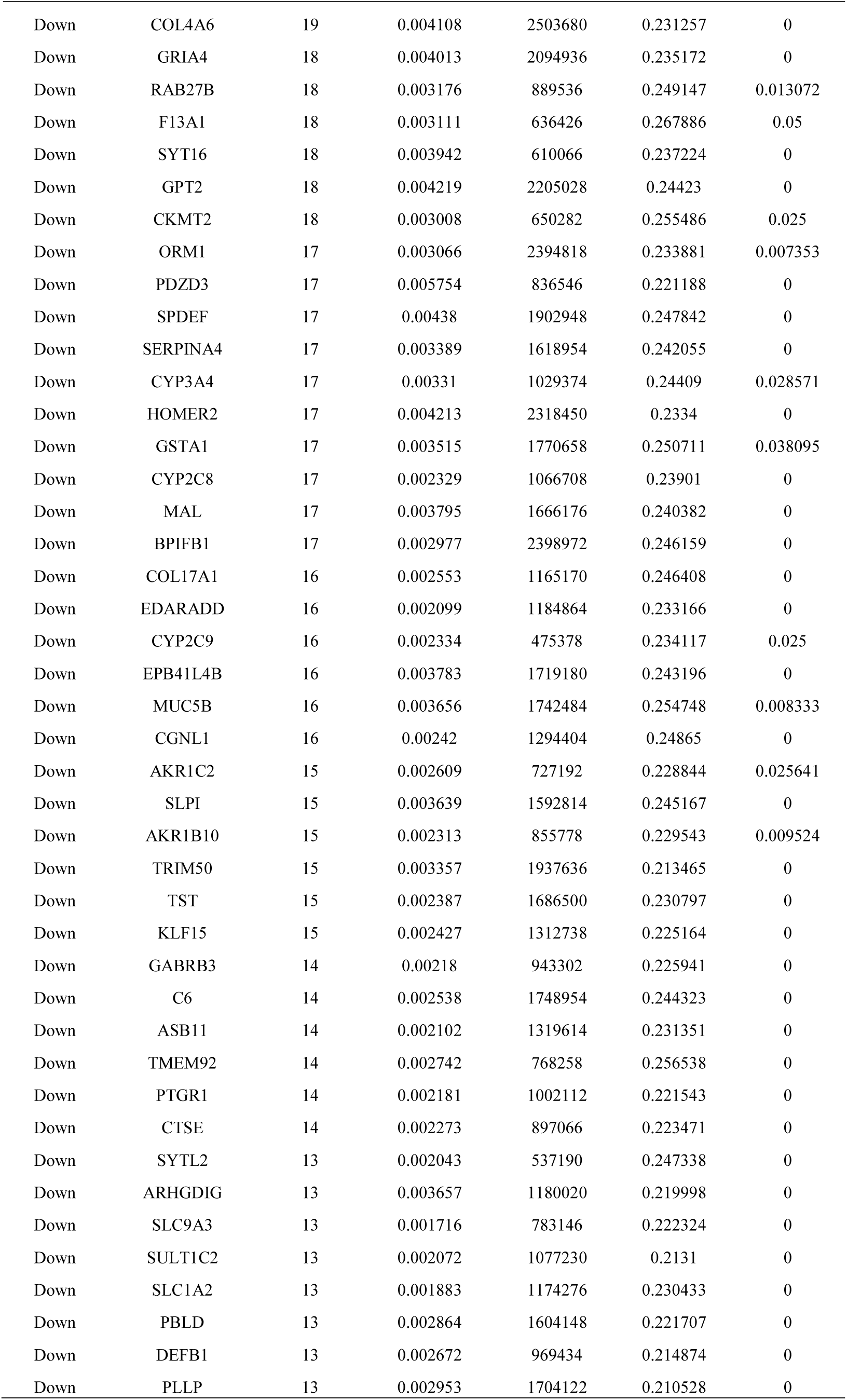
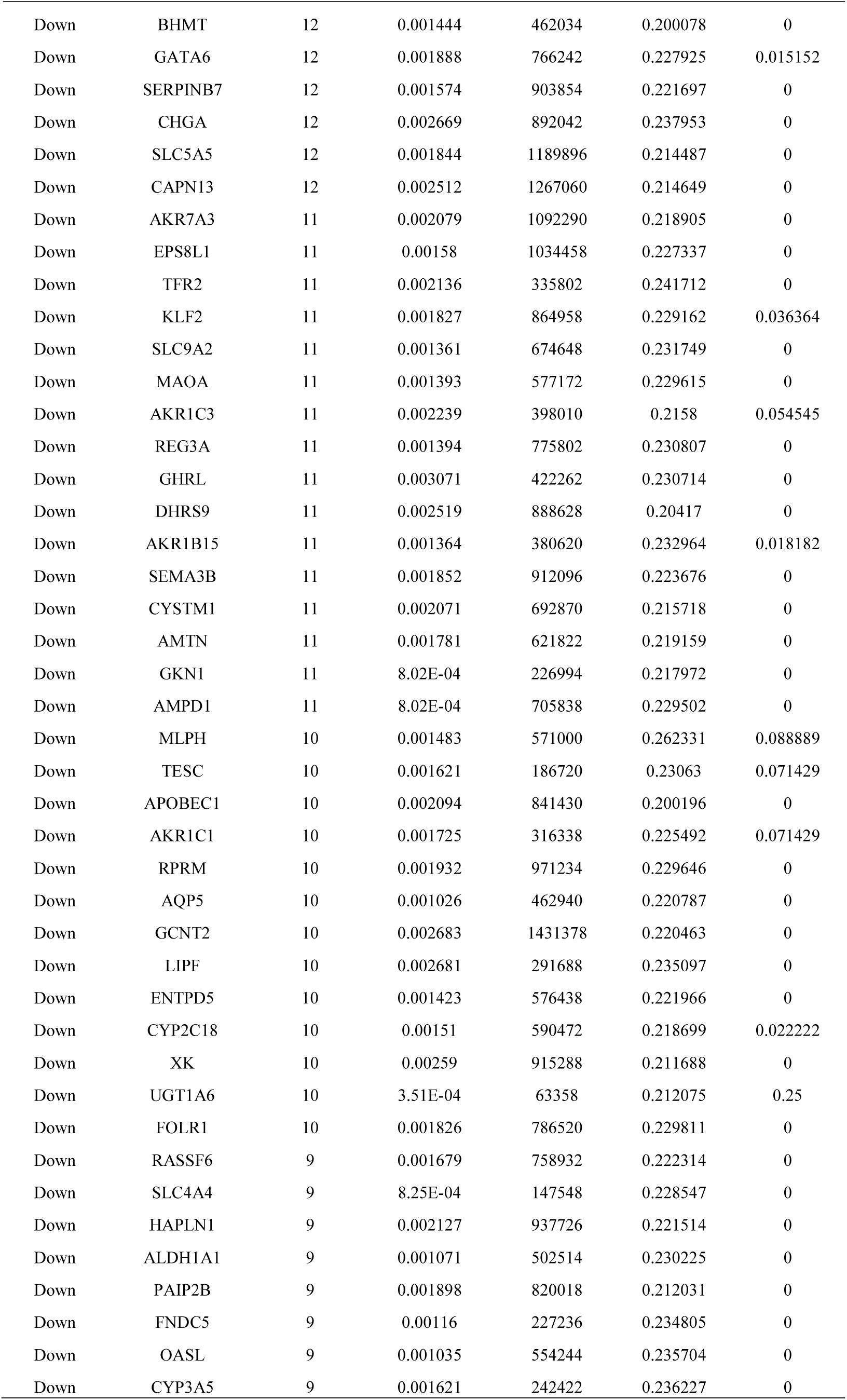
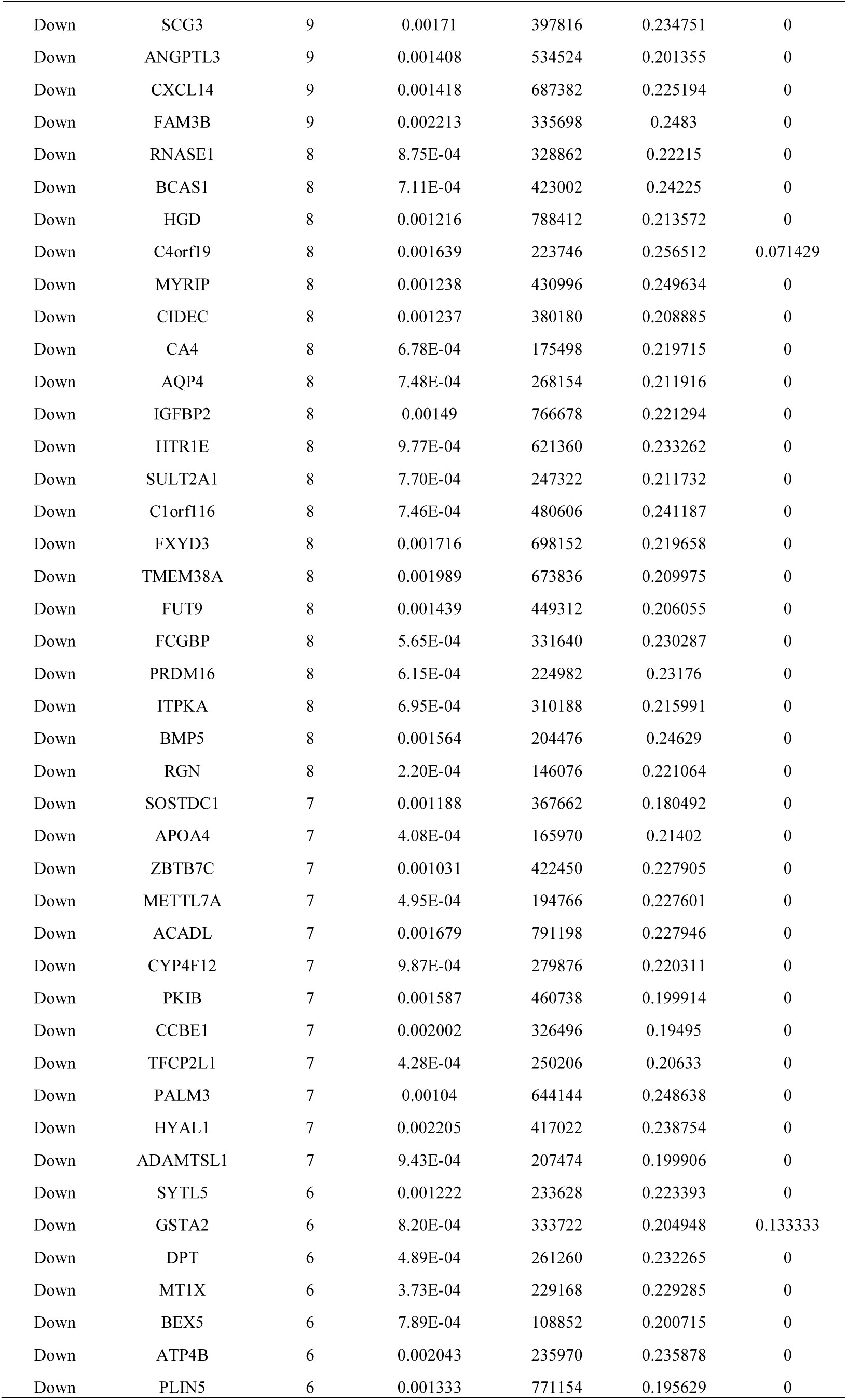
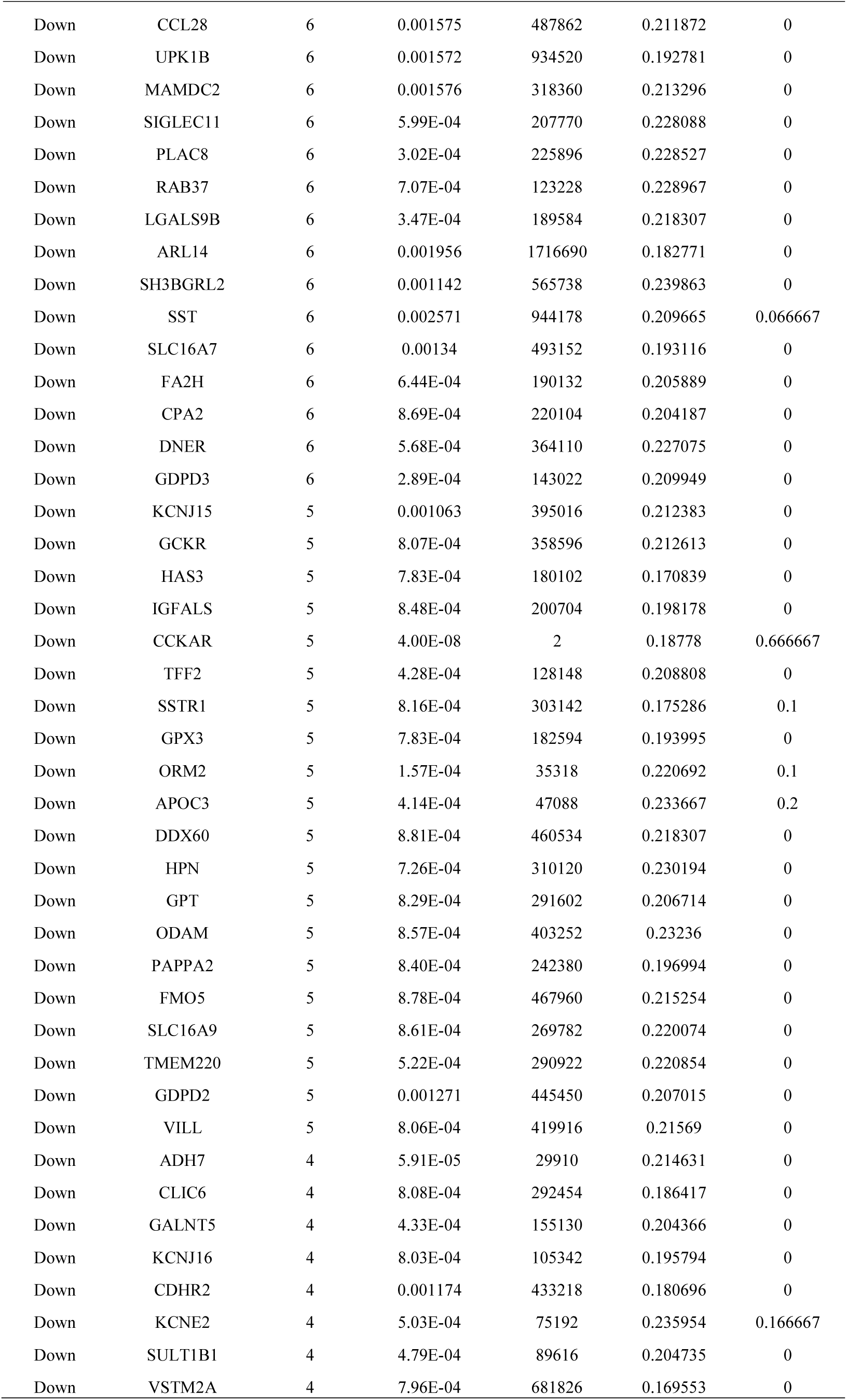
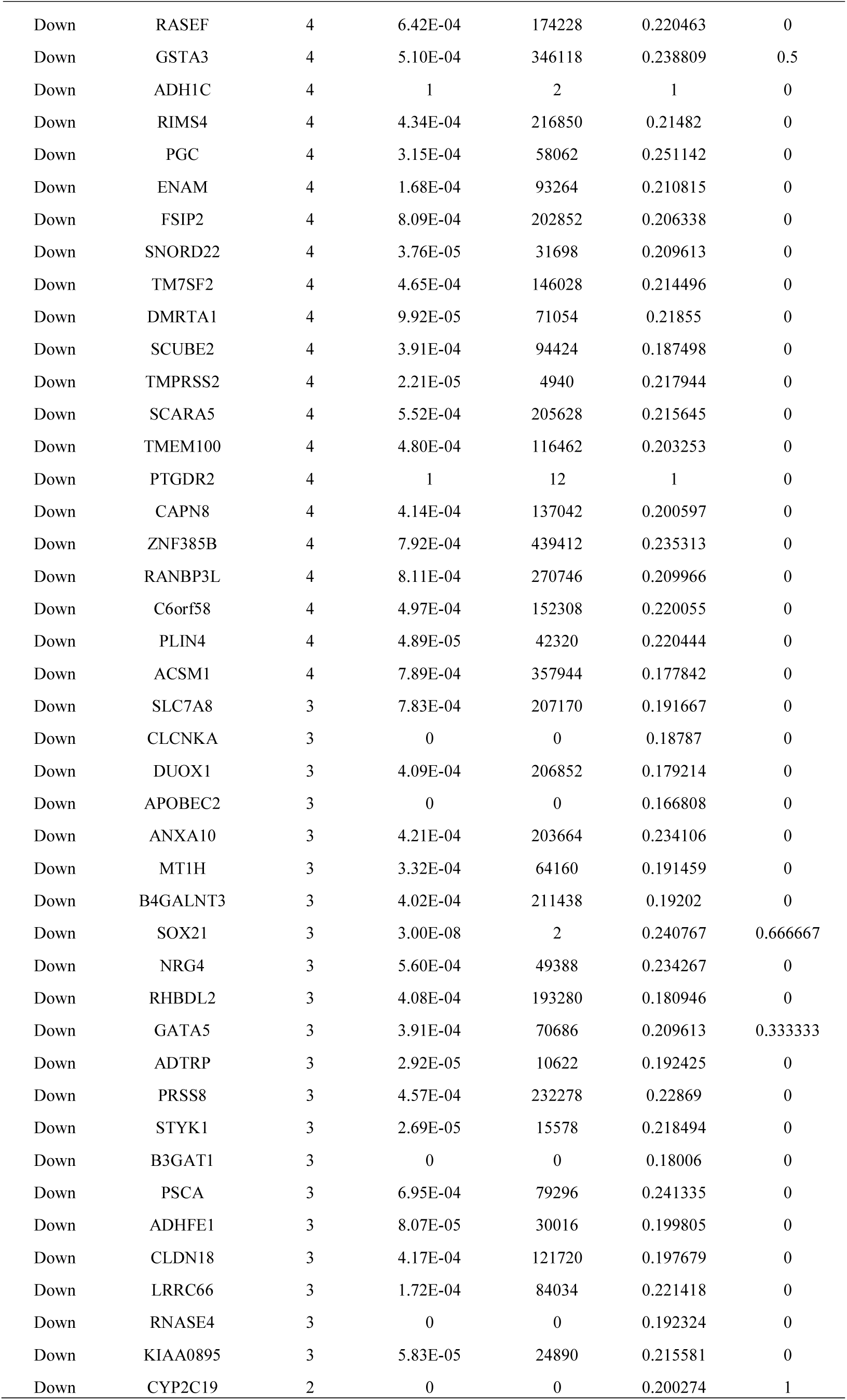
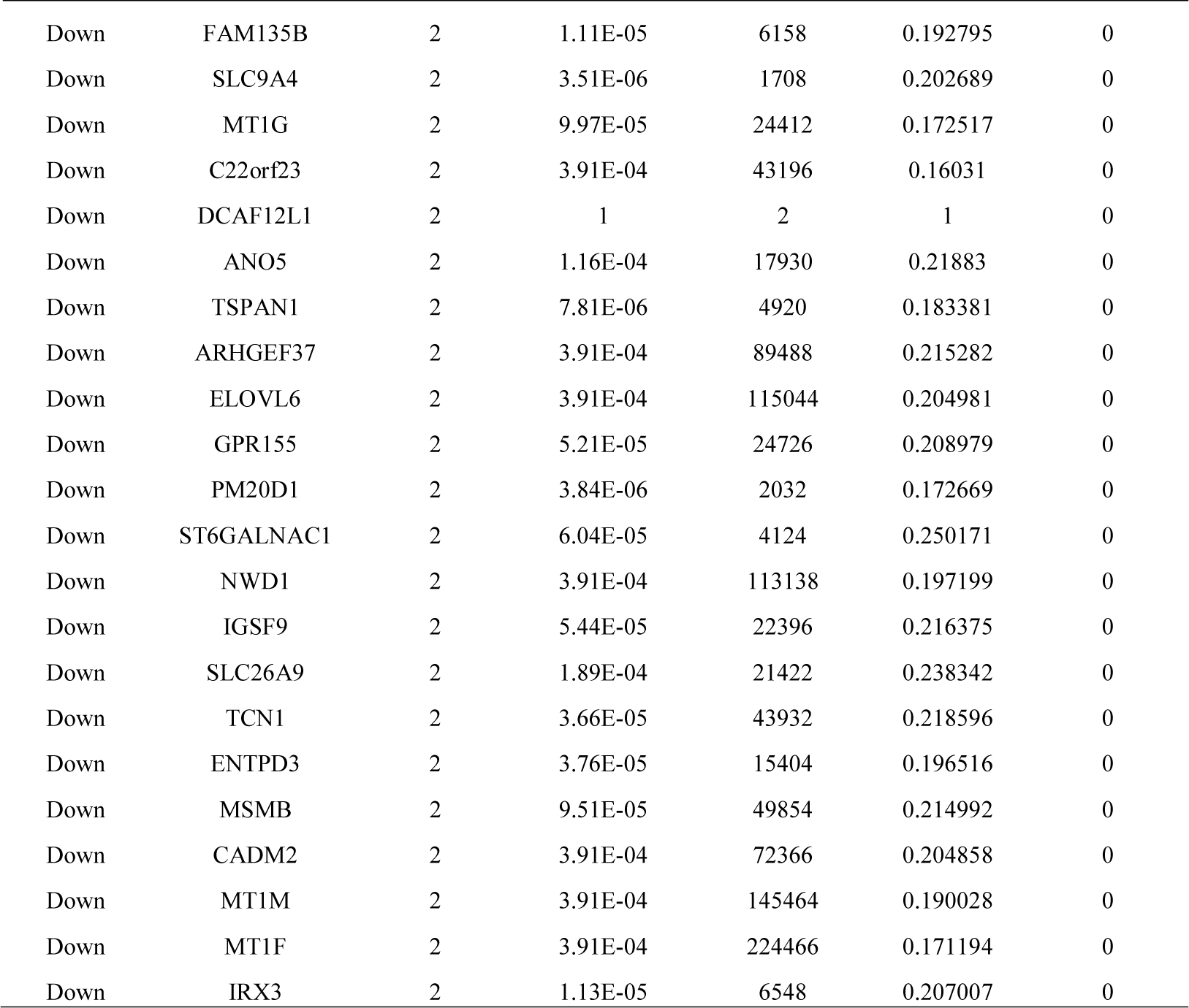
Topology table for up and down regulated genes

Furthermore, the PEWCC1 plugin was applied to investigate highly interconnected regions, known as modules, in the PPI network. Total 668 modules were isolated from PPI network of up regulated genes and four significant modules with the highest scores are shown in Fig. 9. Module 2, consisting of 156 nodes and 175 edges, was highly enriched in microRNAs in cancer, innate immune system, pathways in cancer and notch signaling pathway. Module 5, consisting of 133 nodes and 165 edges, was highly enriched in cell migration, microRNAs in cancer, signaling by Rho GTPases, focal adhesion and cell junction. Module 6, consisting of 118 nodes and 120 edges, was highly enriched in animal organ morphogenesis, G alpha (i) signalling events, water transport, notch signaling pathway and cell migration. Module 9, consisting of 81 nodes and 40 edges, was highly enriched in cell cycle, ATP binding, E2F transcription factor network, identical protein binding, regulation of cell population proliferation and pyrophosphatase activity. Similarly, total 1078 modules were isolated from PPI network of down regulated genes and four significant modules with the highest scores are shown in Fig. 10. Module 6, consisting of 120 nodes and 154 edges, was highly enriched in metabolism of proteins, hemostasis, developmental biology, ensemble of genes encoding extracellular matrix and extracellular matrix-associated proteins and ensemble of genes encoding core extracellular matrix including ECM glycoproteins, collagens and proteoglycans. Module 7, consisting of 114 nodes and 115 edges, was highly enriched in neuroactive ligand-receptor interaction, oxidation-reduction process, gastric acid secretion and metabolism of proteins. Module 10, consisting of 100 nodes and 116 edges, was highly enriched in developmental biology, molecular function regulator, ensemble of genes encoding extracellular matrix and extracellular matrix-associated proteins, metabolism of proteins and biological adhesion. Module 15, consisting of 91 nodes and 95 edges, was highly enriched in gastric acid secretion, nitrogen metabolism, FOXA2 and FOXA3 transcription factor networks, metabolism of proteins, transmembrane transport of small molecules, chemical homeostasis, apical part of cell, secretion, pirenzepine pathway, digestion, and protein digestion and absorption.

**Fig. 9.**
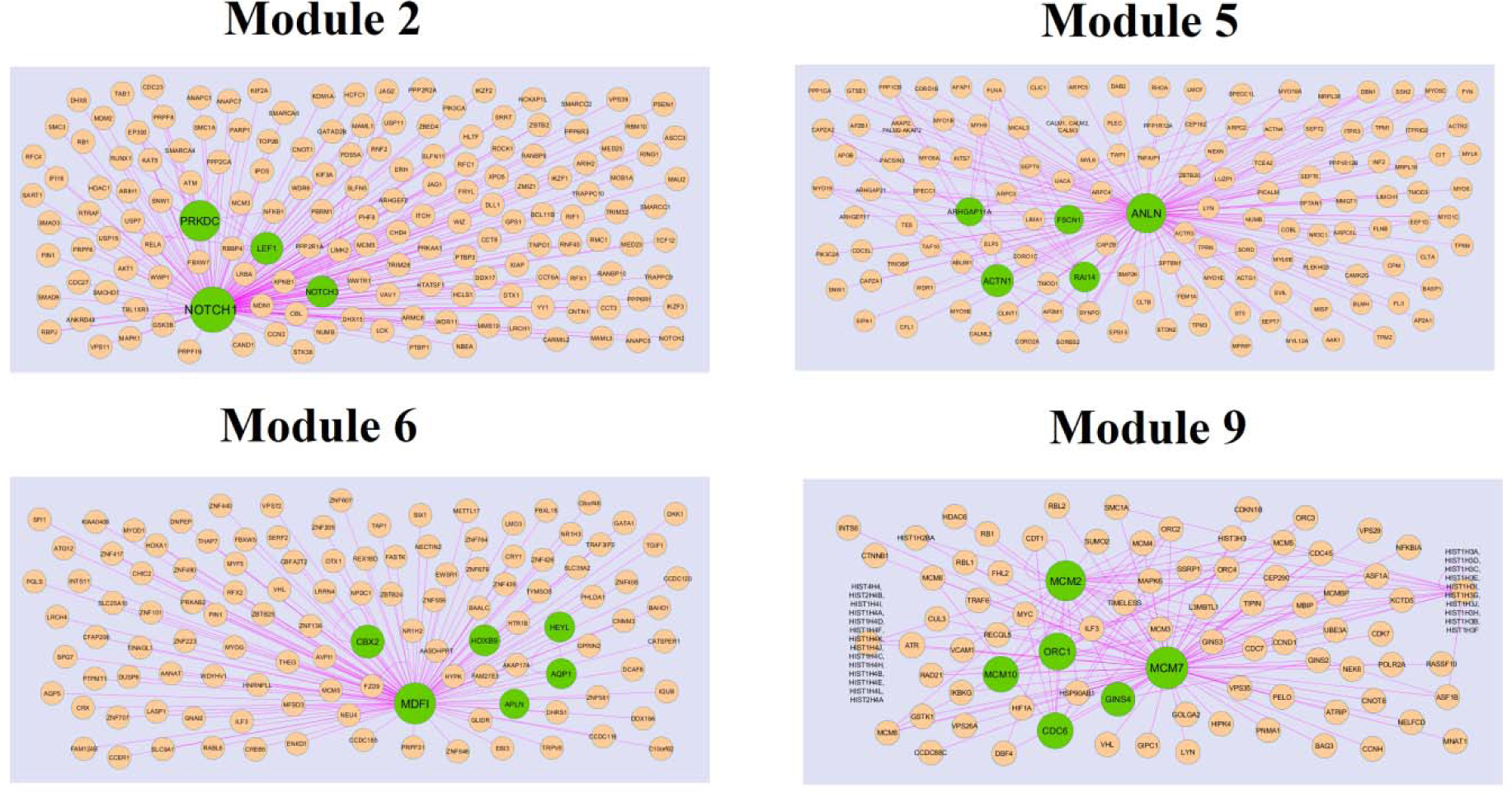
Modules in PPI network. The green nodes denote the up regulated genes

**Fig. 10.**
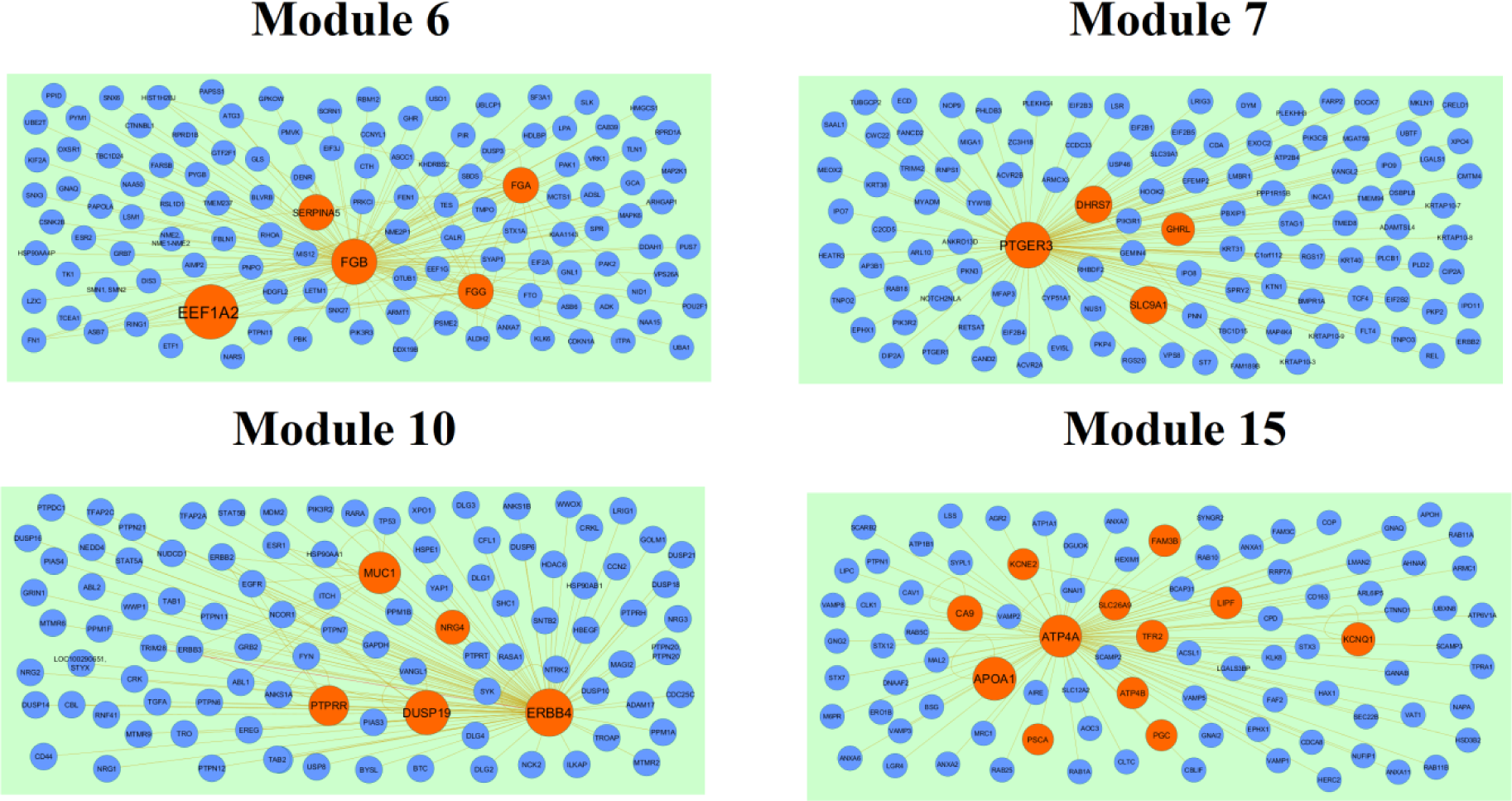
Modules in PPI network. The red nodes denote the down regulated genes

### Target genes - miRNA regulatory network construction

The target genes - miRNA regulatory network of up regulated genes was shown in Fig. 11, consisting of 2669 nodes and 10414 interactions. The nodes with high degree score (interaction with maximum number of miRNAs) can be regarded as essential network nodes. The top target genes with higher degree included FOXK1 interacts with 259 miRNAs (ex,hsa-mir-5693), CCND2 interacts with 179 miRNAs (ex,hsa-mir-3976), RCC2 interacts with 135 miRNAs (ex,hsa-mir-3911), CCDC80 interacts with 135 miRNAs (ex,hsa-mir-6083) and E2F3 interacts with 135 miRNAs (ex,hsa-mir-4271), and are listed in Table 7. The functional enrichment analysis showed that these target genes were significantly involved in peptidase activity, PI3K-Akt signaling pathway, cell cycle, extracellular structure organization and regulation of cell population proliferation. Similarly, target genes - miRNA regulatory network of down regulated genes was shown in Fig. 12, consisting of 2220 nodes and 5985 interactions. The top target genes with higher degree included GATA6 interacts with 207 miRNAs (ex, hsa-mir-4284), SULT1B1 interacts with 131 miRNAs (ex, hsa-mir-4643), GLUL interacts with 126 miRNAs (ex, hsa-mir-4459), ENPP5 interacts with 114 miRNAs (ex,hsa-mir-8082) and KLF2 interacts with 107 miRNAs (ex,hsa-mir-5193) and are listed in Table 7. The functional enrichment analysis showed that these target genes were significantly involved in notch-mediated HES/HEY network, biological oxidations, metabolic pathways, intrinsic component of plasma membrane and response to endogenous stimulus.

**Fig. 11.**
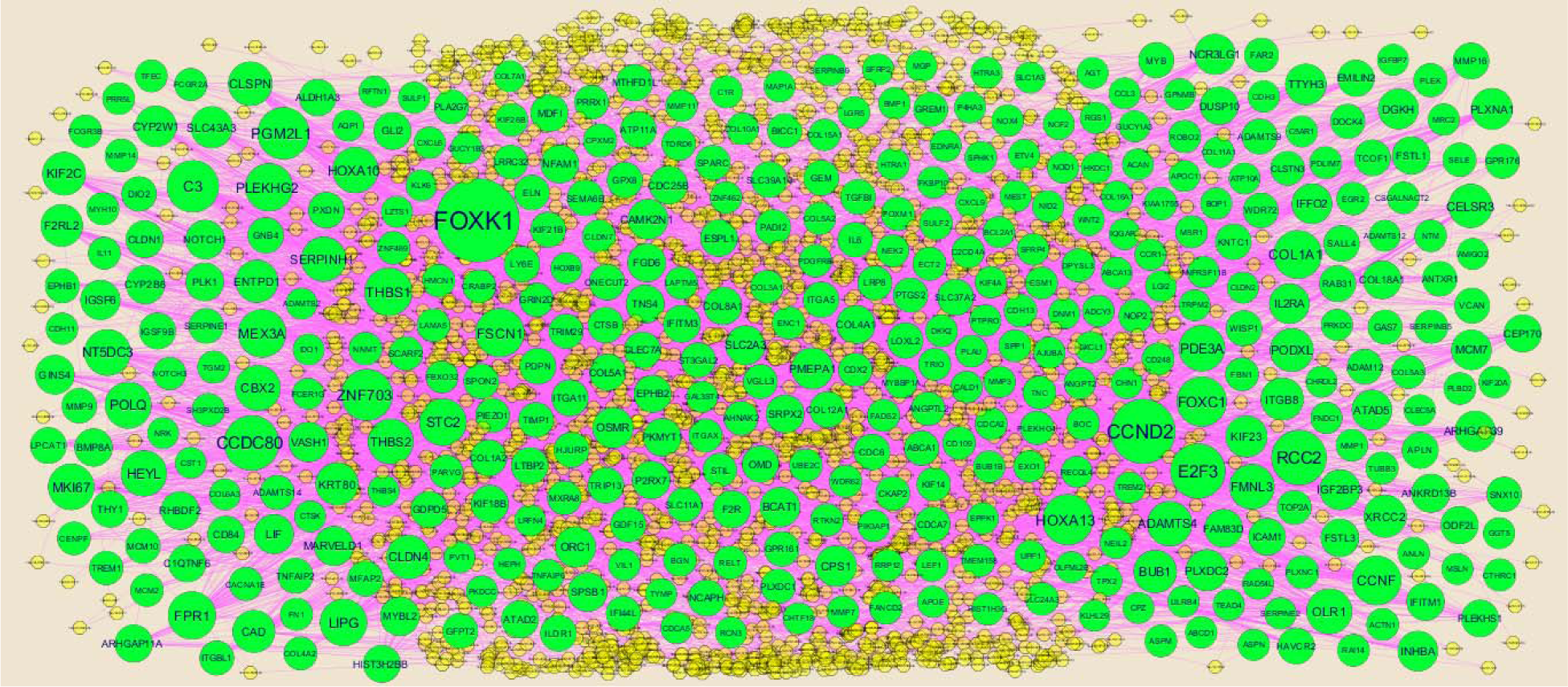
The network of up regulated genes and their related miRNAs. The green circles nodes are the up regulated genes, and yellow diamond nodes are the miRNAs

**Fig. 12.**
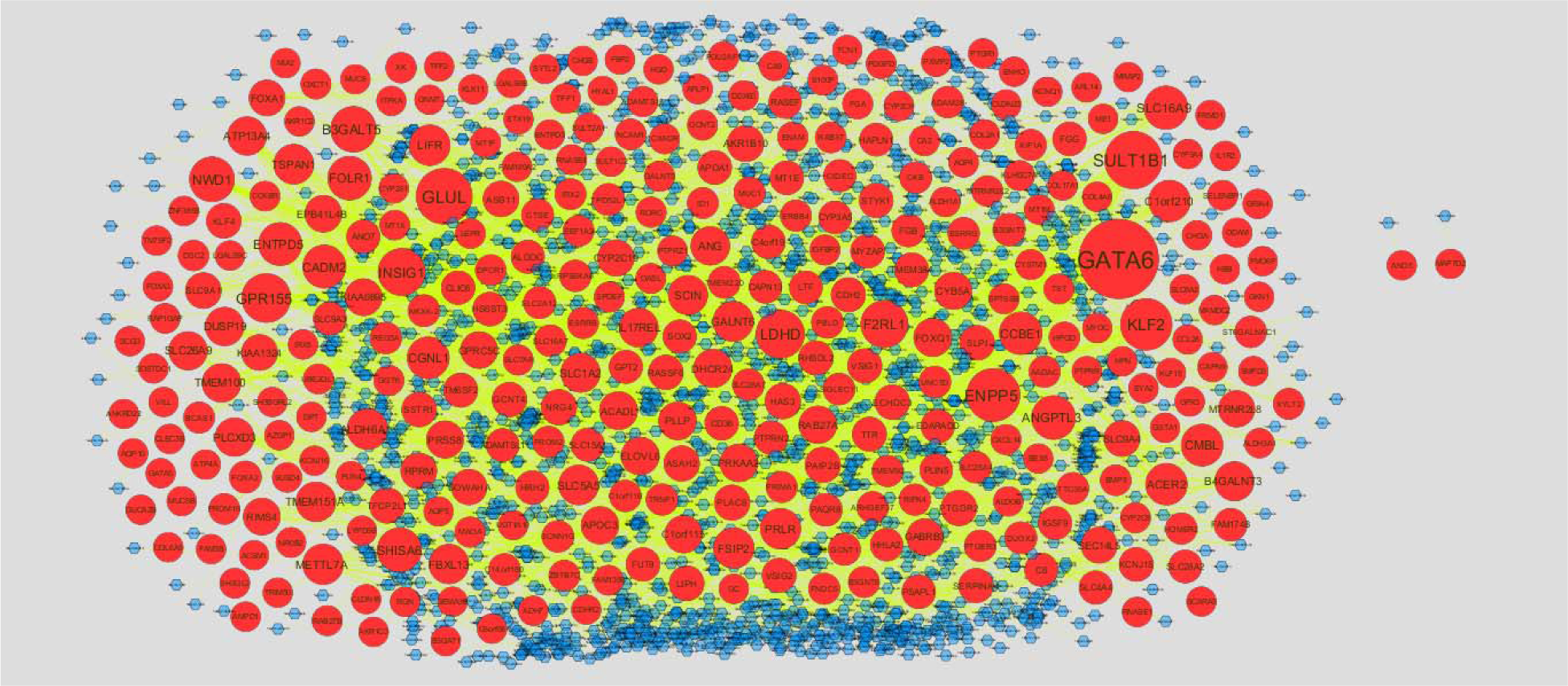
The network of down regulated genes and their related miRNAs. The red circles nodes are the down regulated genes, and blue diamond nodes are the miRNAs

### Target genes - TF regulatory network construction

The target genes - TF regulatory network of up regulated genes was shown in Fig. 13, consisting of 657 nodes and 10767 interactions. The nodes with high degree score (interaction with maximum number of TFs) can be regarded as essential network nodes. The top target genes with higher degree included ABCA13 interacts with 247 TFs (ex, SOX2), ACTN1 interacts with 215 TFs (ex, MYC), ABCD1 interacts with 202 TFs (ex, EGR1), ADAMTS14 interacts with 202 TFs (ex, SPI1) and ADAMTS4 interacts with 194 TFs (ex, NANOG) and are listed in Table 8. The functional enrichment analysis showed that these target genes were significantly involved in neutrophil degranulation, focal adhesion, regulation of phosphate metabolic process, peptidase activity and extracellular matrix organization. Similarly, target genes - TF regulatory network of down regulated genes was shown in Fig. 14, consisting of 639 nodes and 7623 interactions. The top target genes with higher degree included DNER interacts with 177 (ex, TP63), CKB interacts with 169 (ex, STAT3), PKIB interacts with 165 (ex, HNF4A), RAB27B interacts with 164 (ex, AR) and IRX3 interacts with 150 (ex, SUZ12) and are listed in Table 8. The functional enrichment analysis showed that these target genes were significantly involved in neuron projection, gastric acid secretion, molecular function regulator and pancreatic secretion.

**Fig. 13.**
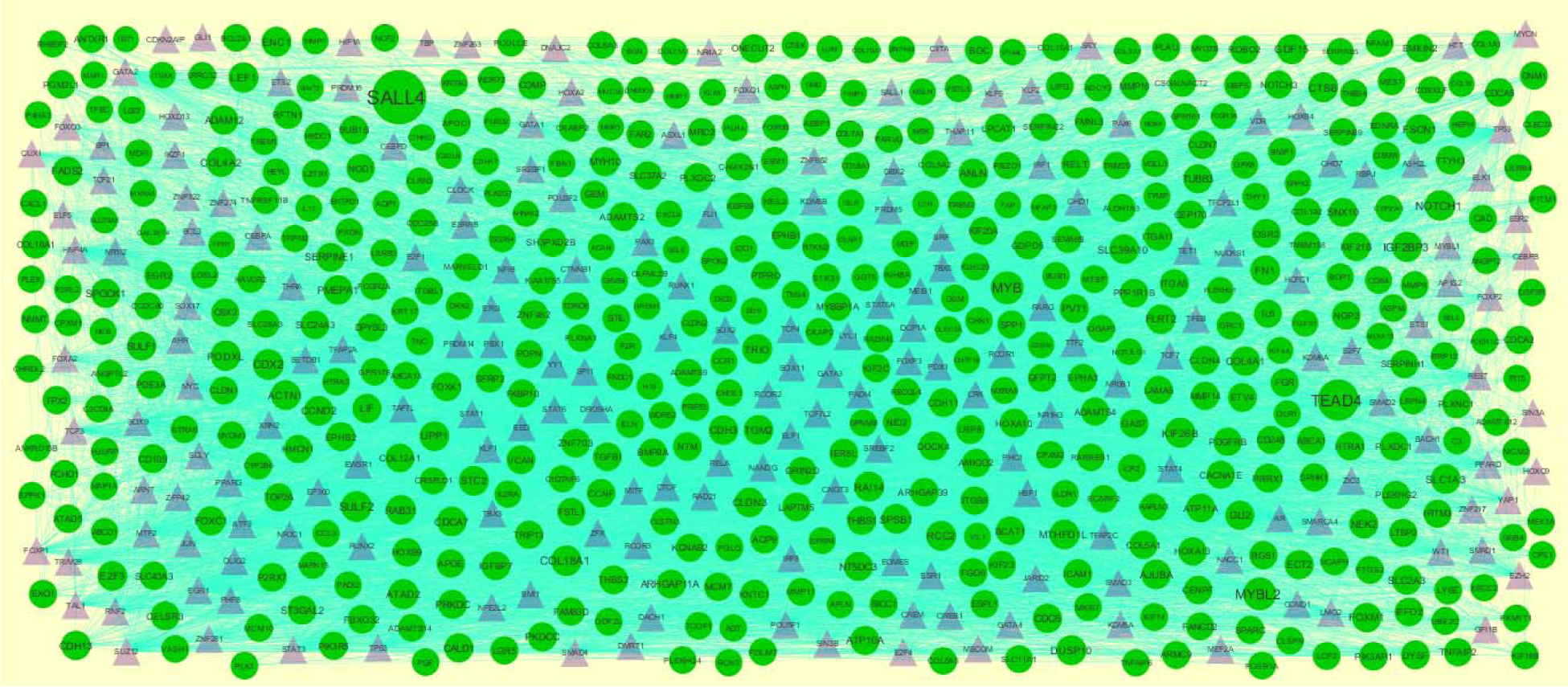
The network of up regulated genes and their related TFs. The green circles nodes are the up regulated genes, and purple triangle nodes are the TFs

**Fig. 14.**
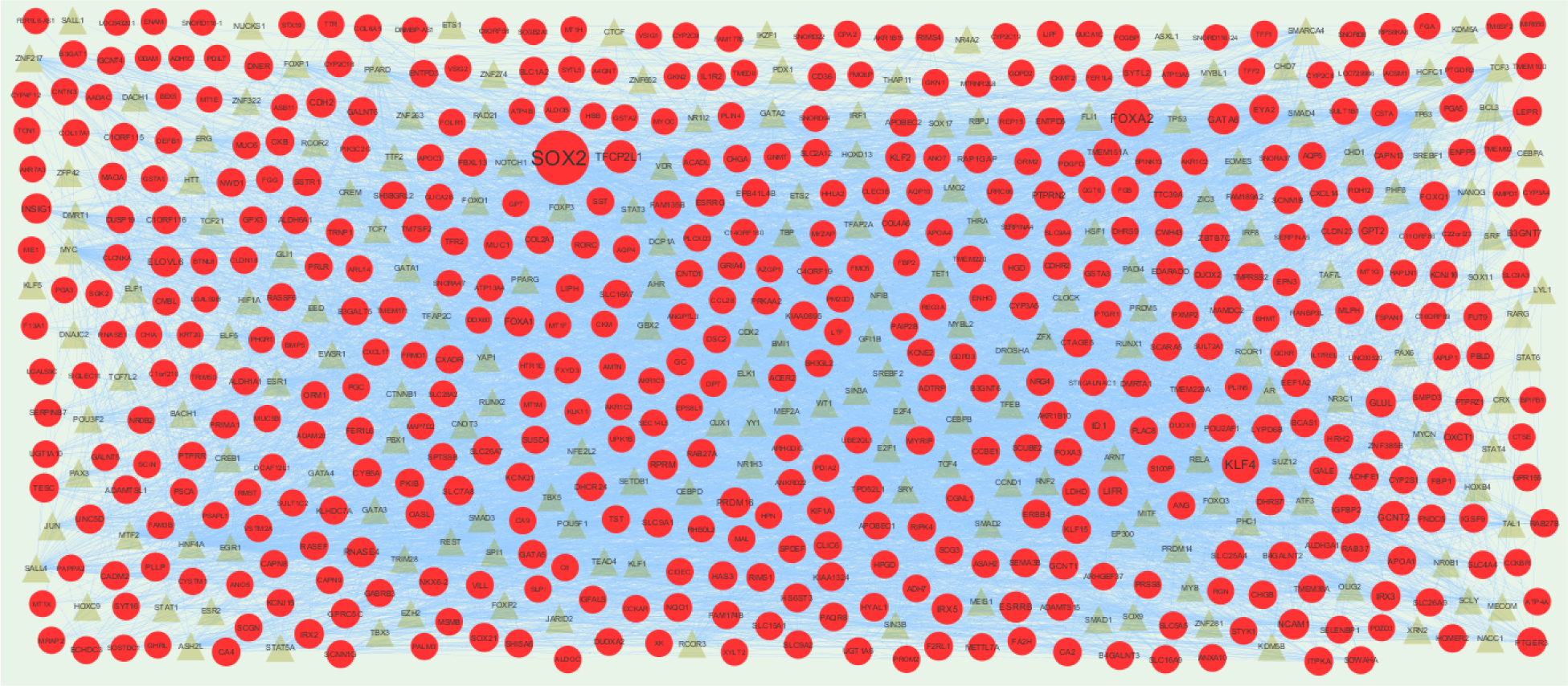
The network of down regulated genes and their related TFs. The green circles nodes are the down regulated genes, and yellow triangle nodes are the TFs.

**Table 8.**
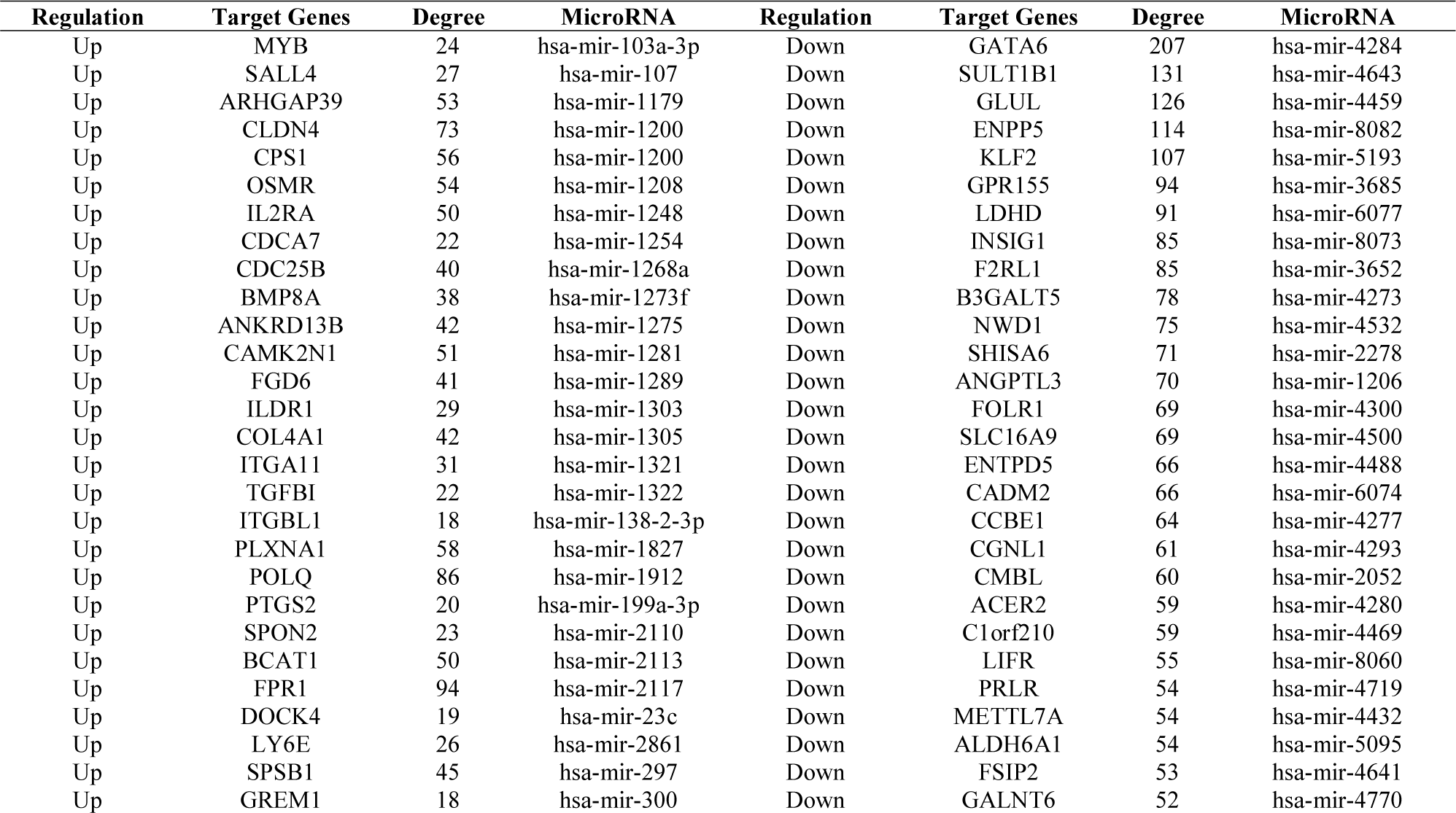
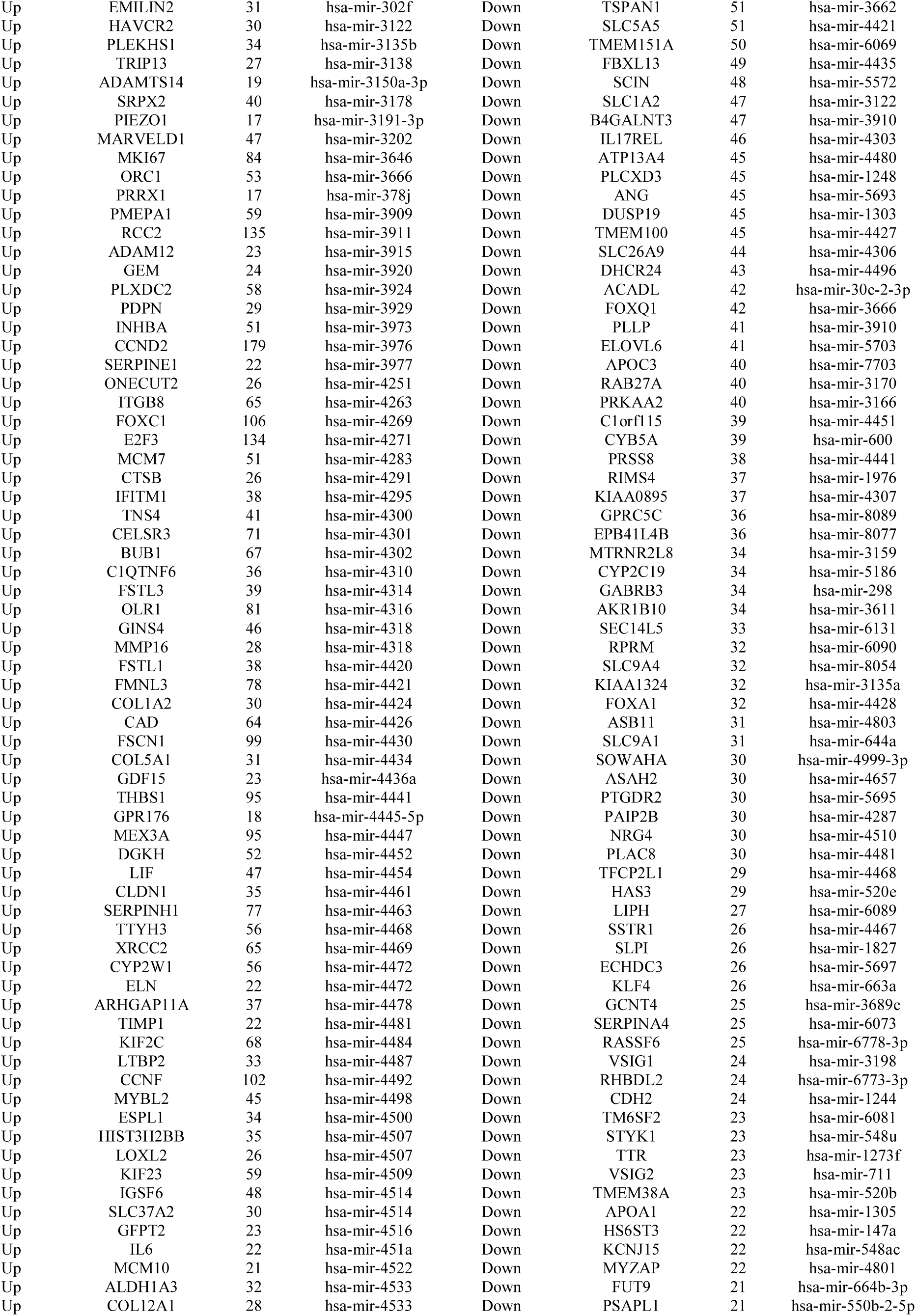
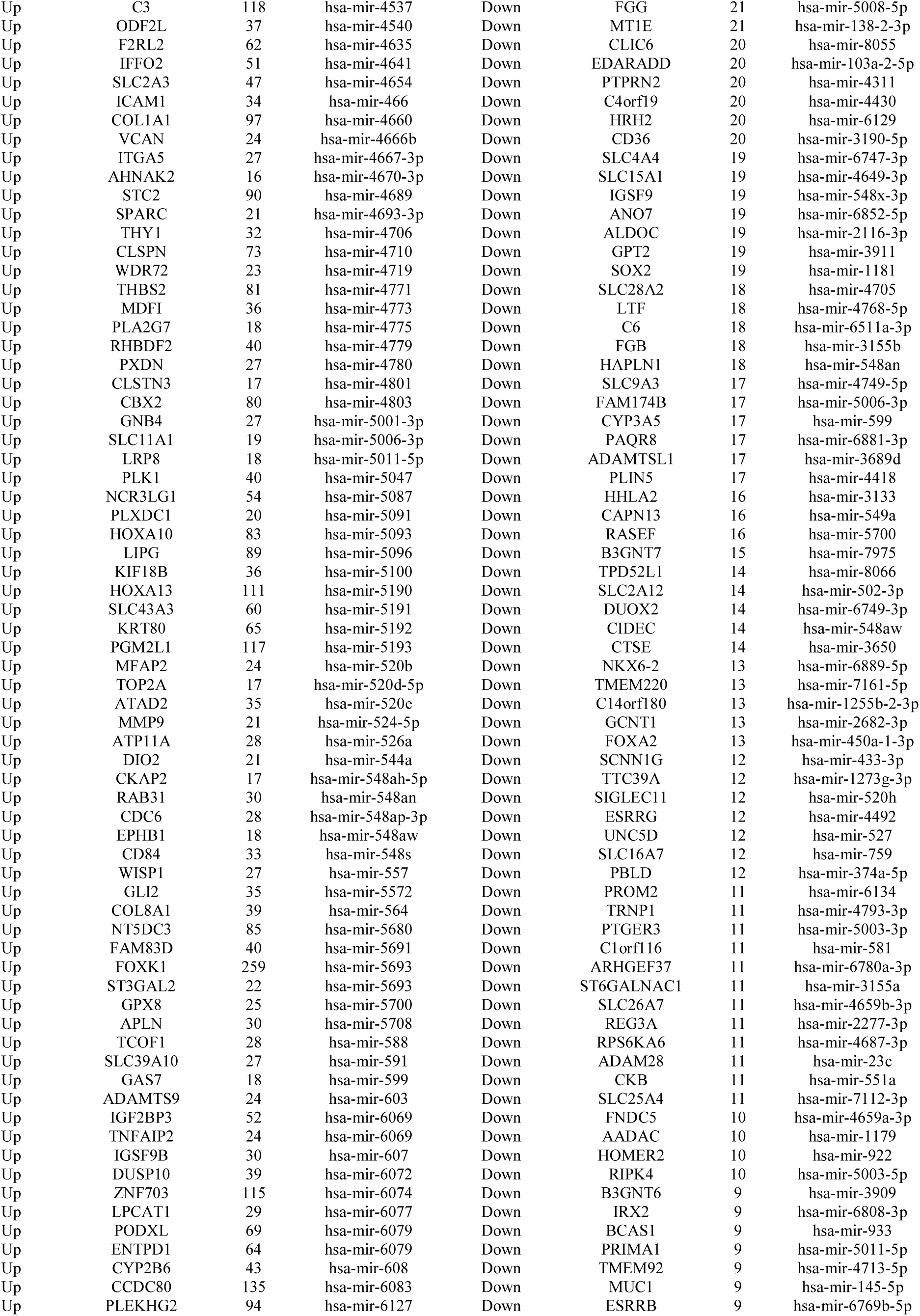
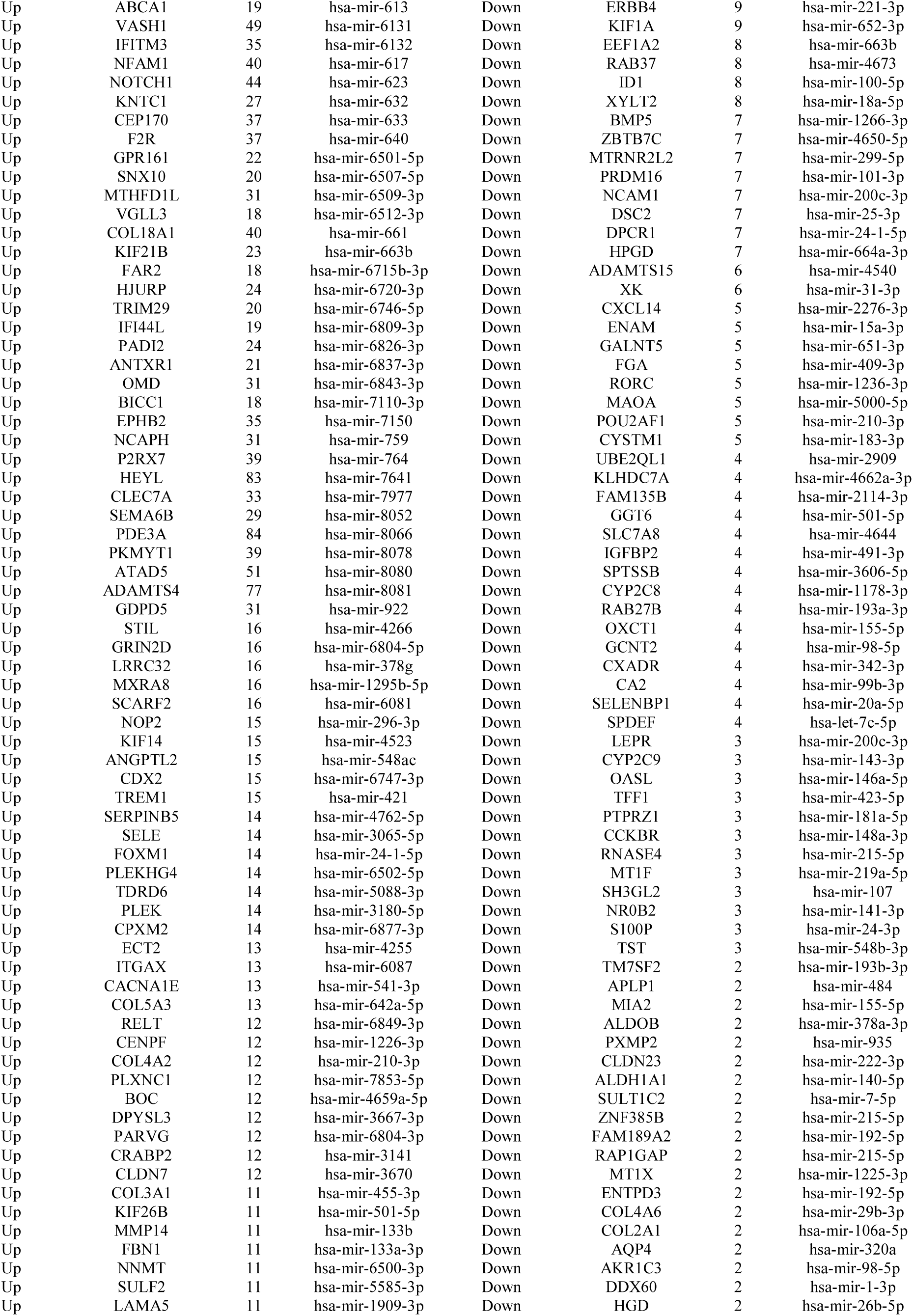
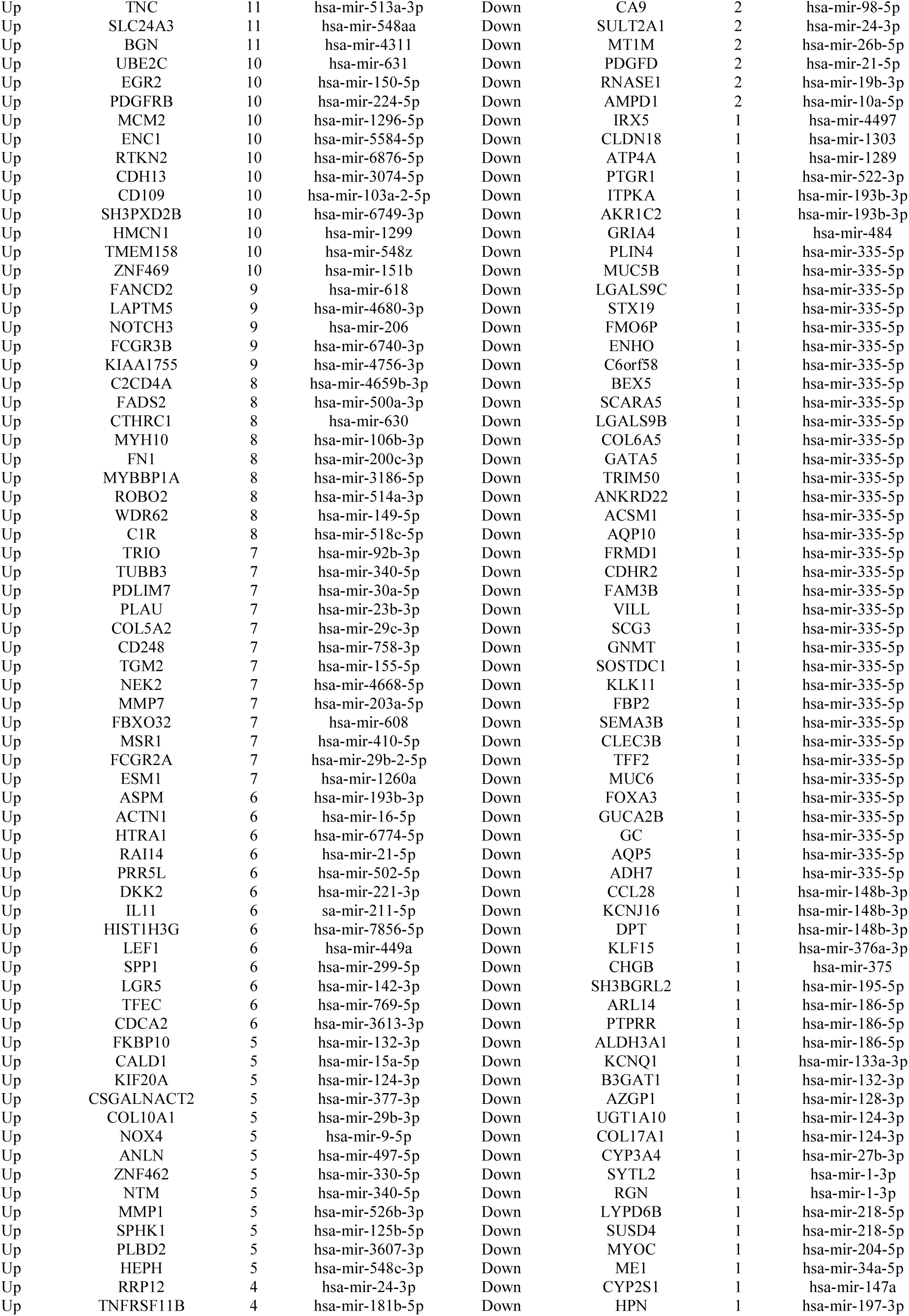
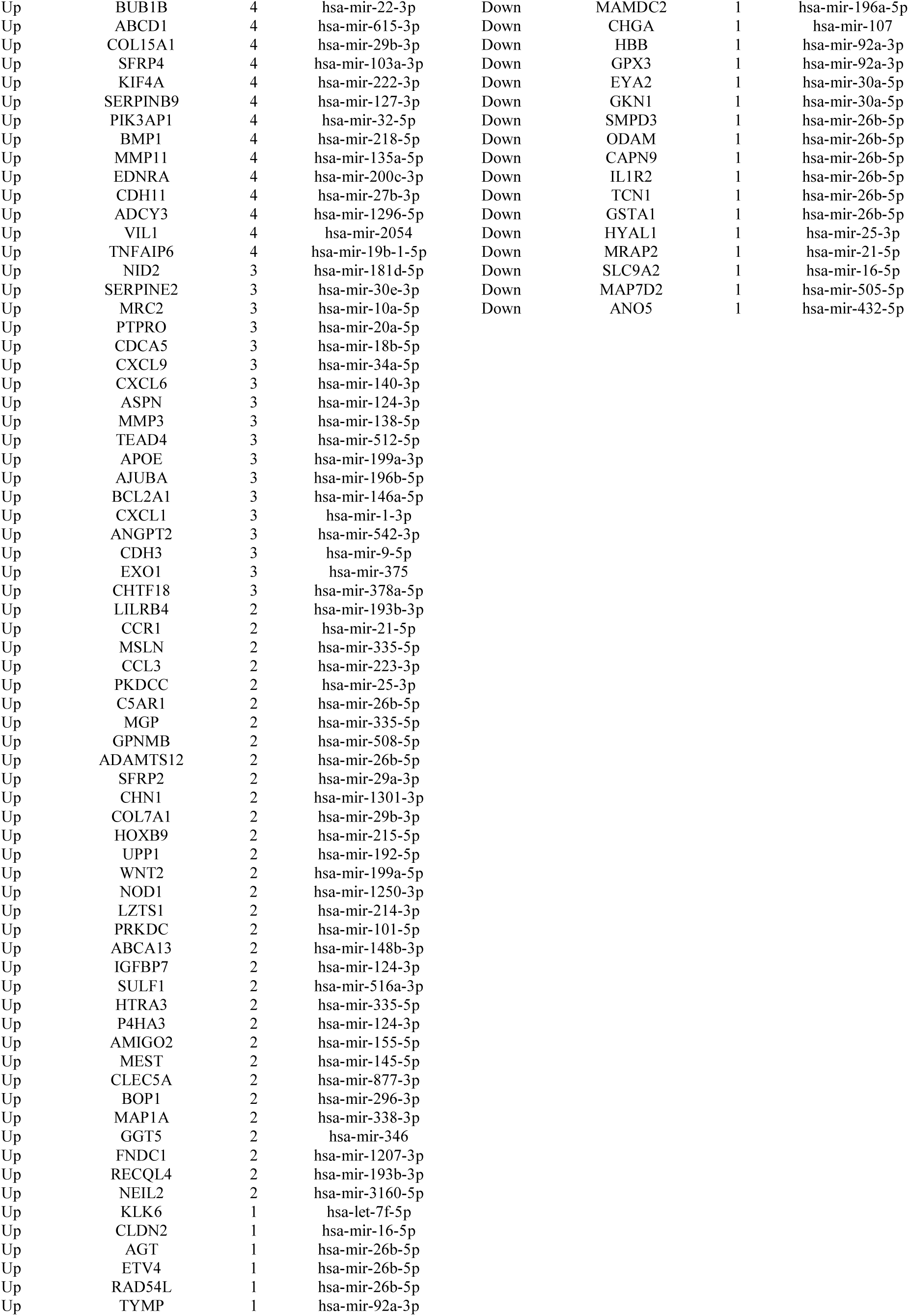
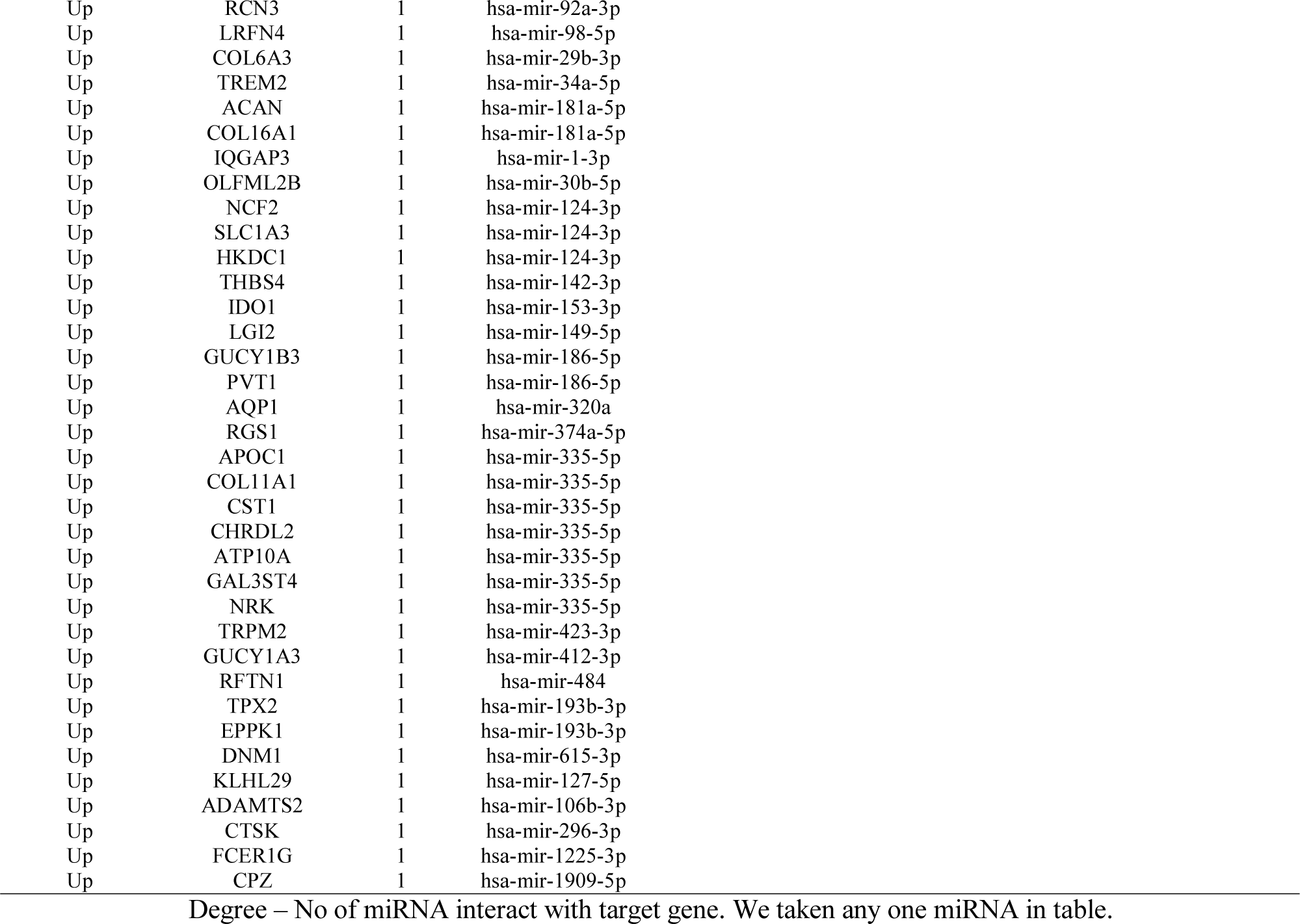
miRNA - target gene interaction table

**Table 9.**
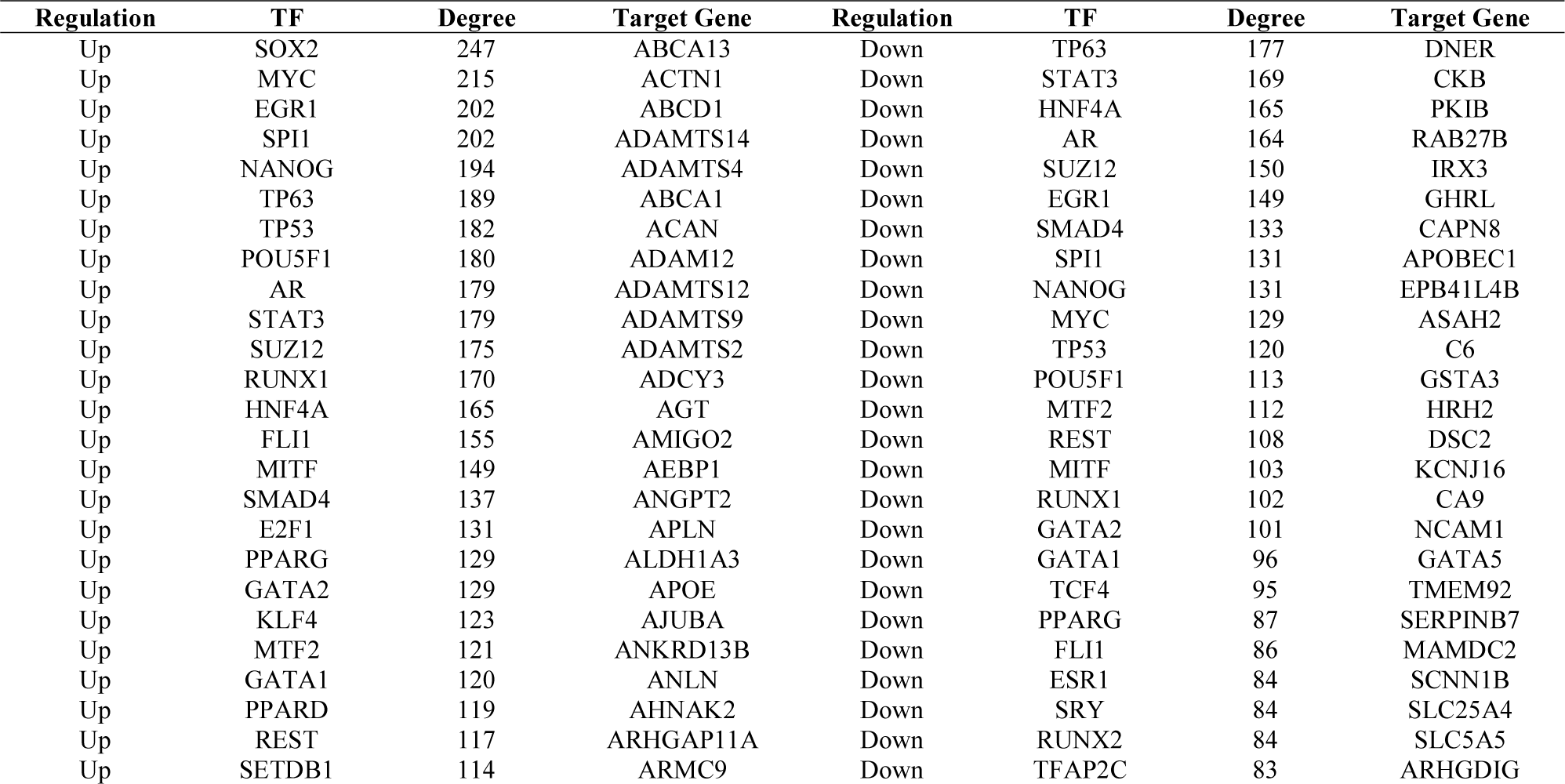
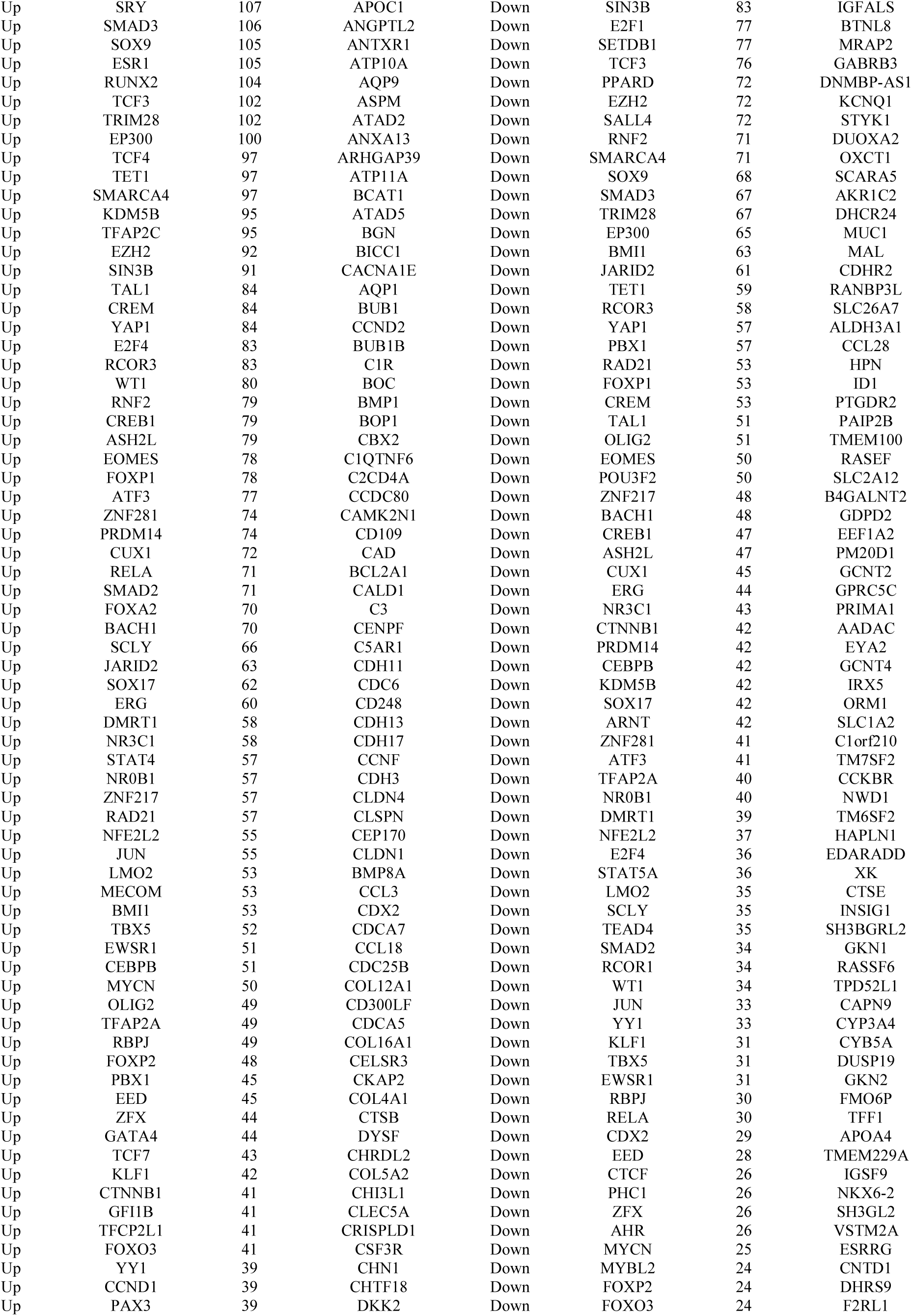
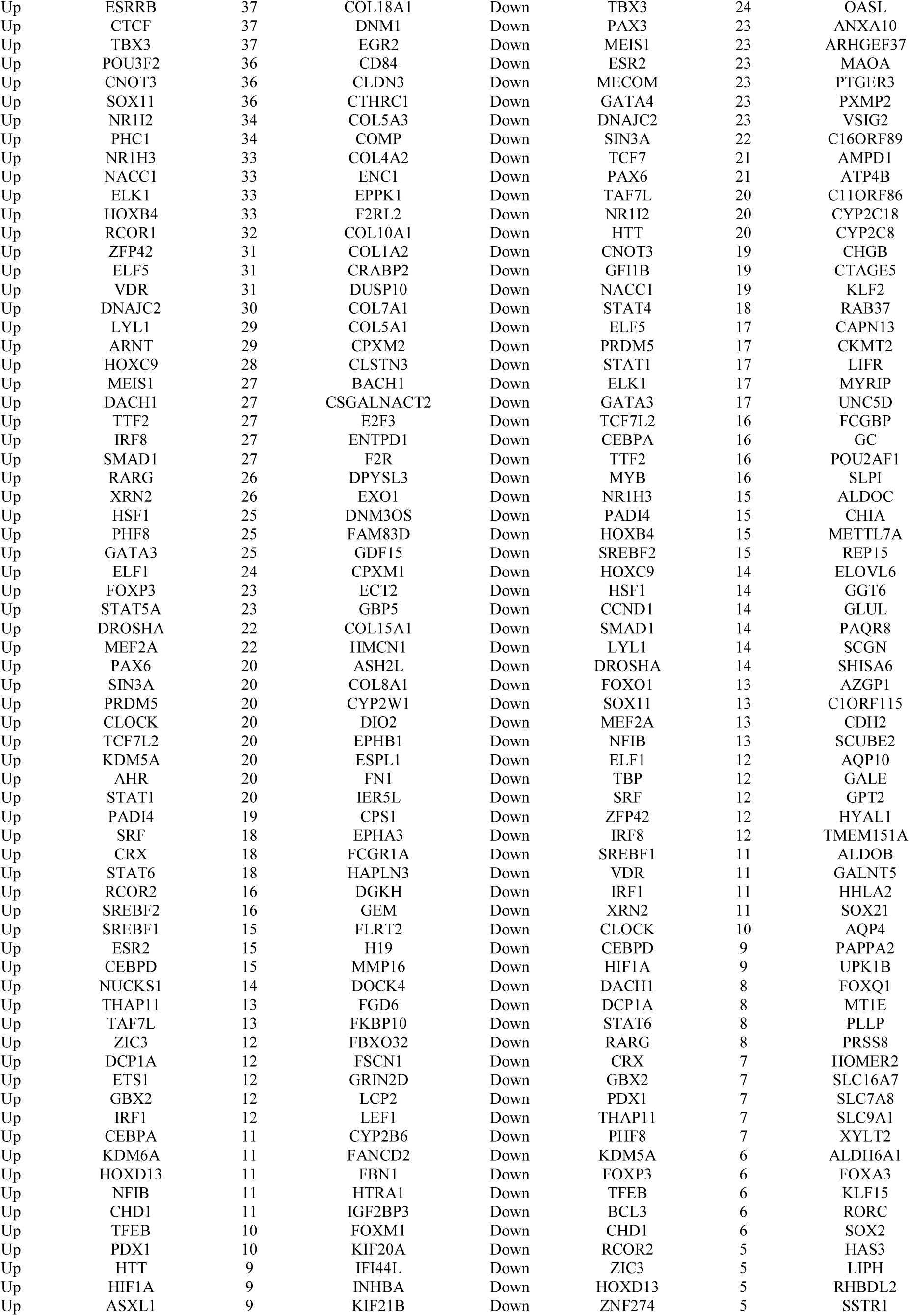
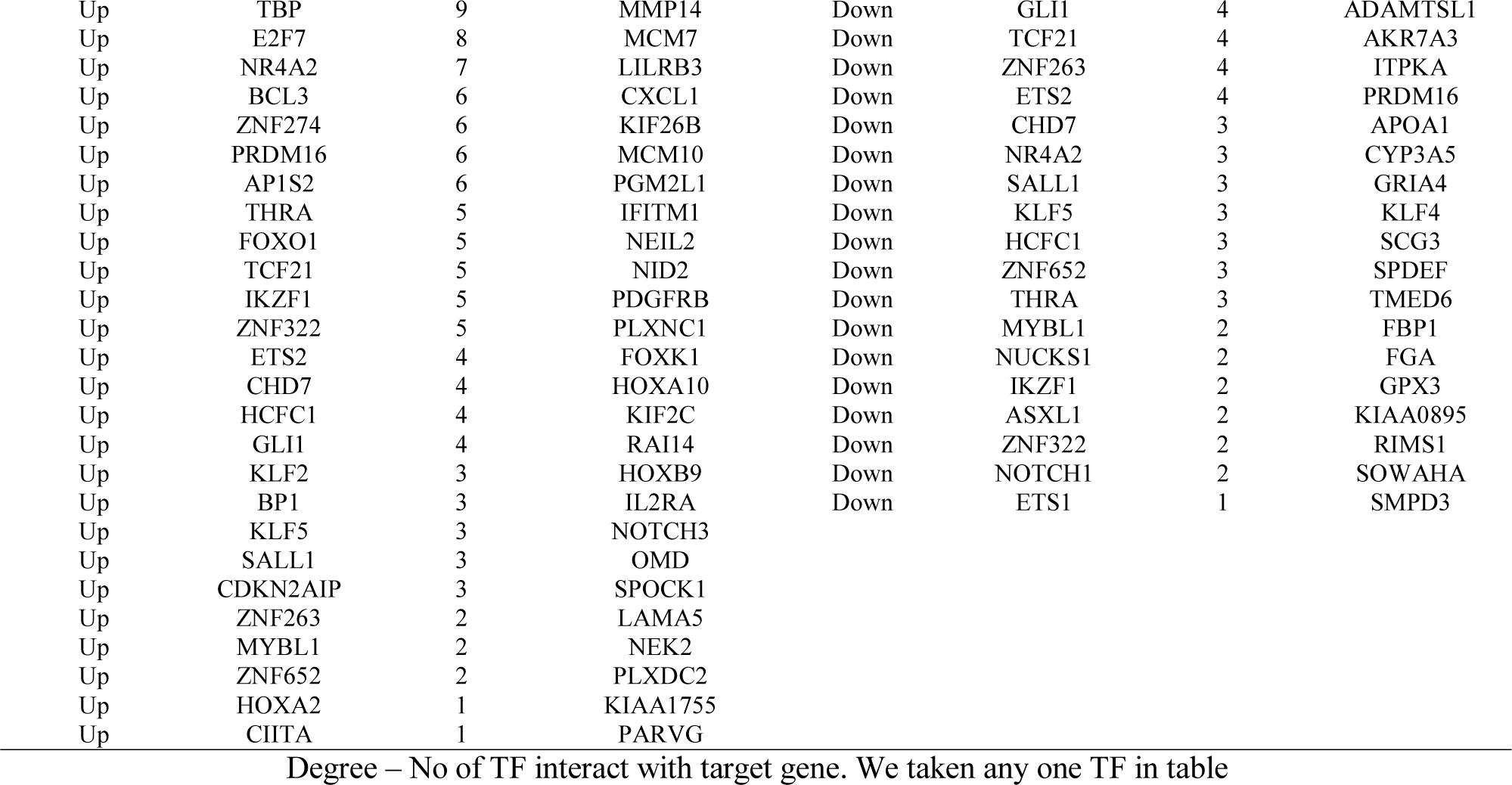
TF - target gene interaction table

### Validations of hub genes

Up regulated hub genes and down regulated hub genes were then validated in database TCGA to confirm the outcomes. We tried to analysis the relationship between hub genes and the survival in GC. The prognostic and diagnostics value of hub genes was determined by UALCAN database. Patients with high expression of FN1 (Fig. 15A), PLK1 (Fig. 15B), ANLN (Fig. 15C), MCM7 (Fig. 15D), MCM2 (Fig. 15E), EEF1A2 (Fig. 16A), PTGER3 (Fig. 16B), CKB (Fig. 16C), ERBB4 (Fig. 16D) and PRKAA2 (Fig. 16E) were associated with shorter overall survival. The UALCAN box plot analysis investigated the level of expression of the hub genes in 415 GC tissue samples and 34 normal tissue samples. The boxplot in Fig. 17A - Fig. 17E shows a considerable increase in the level of hub gene expression (FN1, PLK1, ANLN, MCM7 and MCM2) in the GC, while boxplot in Fig. 17F - Fig. 17J shows a considerable decrease in the level of hub gene expression (EEF1A2, PTGER3, CKB, ERBB4 and PRKAA2) in the GC. Furthermore, the UALCAN box plot analysis investigated the level of expression of the hub genes in individual GC stages (stage 1 (18 GC samples), stage 2 (123 GC samples), stage 3 (169 GC samples) and stage 4 (41 GC samples)) and 34 normal tissue samples. The boxplot in Fig. 18A - Fig. 18E shows a considerable increase in the level of hub gene expression (FN1, PLK1, ANLN, MCM7 and MCM2) in all four individual stages of GC, while boxplot in Fig. 18F - Fig. 18J shows a considerable decrease in the level of hub gene expression (EEF1A2, PTGER3, CKB, ERBB4 and PRKAA2) in all four individual stages of GC. Based on the results of the cBioPortal database, results of genomic modifications for these hub genes were depicted by using the cBioPortal. We found 6%, 5%, 3%, 8%, 4%, 5%, 1.4%, 0.7%, 13% and 2.5% of GC cases exhibited FN1, PLK1, ANLN, MCM7, MCM2, EEF1A2, PTGER3, CKB, ERBB4 and PRKAA2 hub genes modification including inframe mutation (unknown significance), missense mutation (putative driver), deep deletion, missense mutation (unknown significance), amplification and truncating mutation (unknown significance) and are shown in Fig. 19. Additionally, the expression and distribution of the hub genes (proteins) in GC samples and nontumor samples were measured through immunohistochemistry (IHC). IHC analysis of FN1, PLK1, ANLN, MCM7, MCM2, EEF1A2, PTGER3, CKB, ERBB4 and PRKAA2 in GC samples were found in the HPA database. The antibodies used were as follows: FN1 (CAB000126), PLK1 (HPA051638), ANLN (HPA050556), MCM7 (CAB016312), MCM2 (CAB000303), EEF1A2 (CAB034019), PTGER3 (HPA010689), CKB (HPA001254), ERBB4 (CAB000276) and PRKAA2 (HPA044540). Up regulated hub genes such as FN1, PLK1, ANLN, MCM7 and MCM2 were highly expressed in GC tissue but undetectable or expressed at low levels in normal tissue (Fig. 20A - Fig. 20E). Down regulated hub genes such as EEF1A2, PTGER3, CKB, ERBB4 and PRKAA2 were less expressed in GC tissue but detectable or expressed at high levels in normal tissue (Fig. 20F - Fig. 20J). The ROC curve analysis was accomplished to assess the diagnostic and prognostic values of hub genes. Our finding revealed that FN1 (AUC = 0.979), PLK1 (AUC = 0.921), ANLN (AUC = 0.946), MCM7 (AUC = 0.978), MCM2 (AUC = 0.961), EEF1A2 (AUC = 0.955), PTGER3 (AUC = 0.934), CKB (AUC = 0.869), ERBB4 (AUC = 0.769) and PRKAA2 (AUC = 0.761) had significant diagnostic and prognostic values for discriminating GC samples and normal controls (Fig. 21). Furthermore, RT PCR was used to verify the expression levels of the hub genes in primary GC and normal control tissue. In the GC tissues, FN1, PLK1, ANLN, MCM7 and MCM2 were significantly up regulated as compared with those in their normal control tissue, while in GC samples EEF1A2, PTGER3, CKB, ERBB4 and PRKAA2 were significantly down regulated as compared with those in their normal control tissue. These results indicated that, consistent with the TCGA results, most of these hub genes were dysregulated in GC samples compared with those in the normal control samples (Fig. 22). Finally, we investigated whether FN1, PLK1, ANLN, MCM7, MCM2, EEF1A2, PTGER3, CKB, ERBB4 and PRKAA2 expression were associated with tumor immune infiltration level by using the TIMER database. Interestingly, we found that the high expression of FN1, PLK1, ANLN, MCM7 and MCM2 were significantly negatively related to tumor purity, while low expression EEF1A2, PTGER3, CKB, ERBB4 and PRKAA2 were significantly positively related to tumor purity (Fig. 23).

**Fig. 15.**
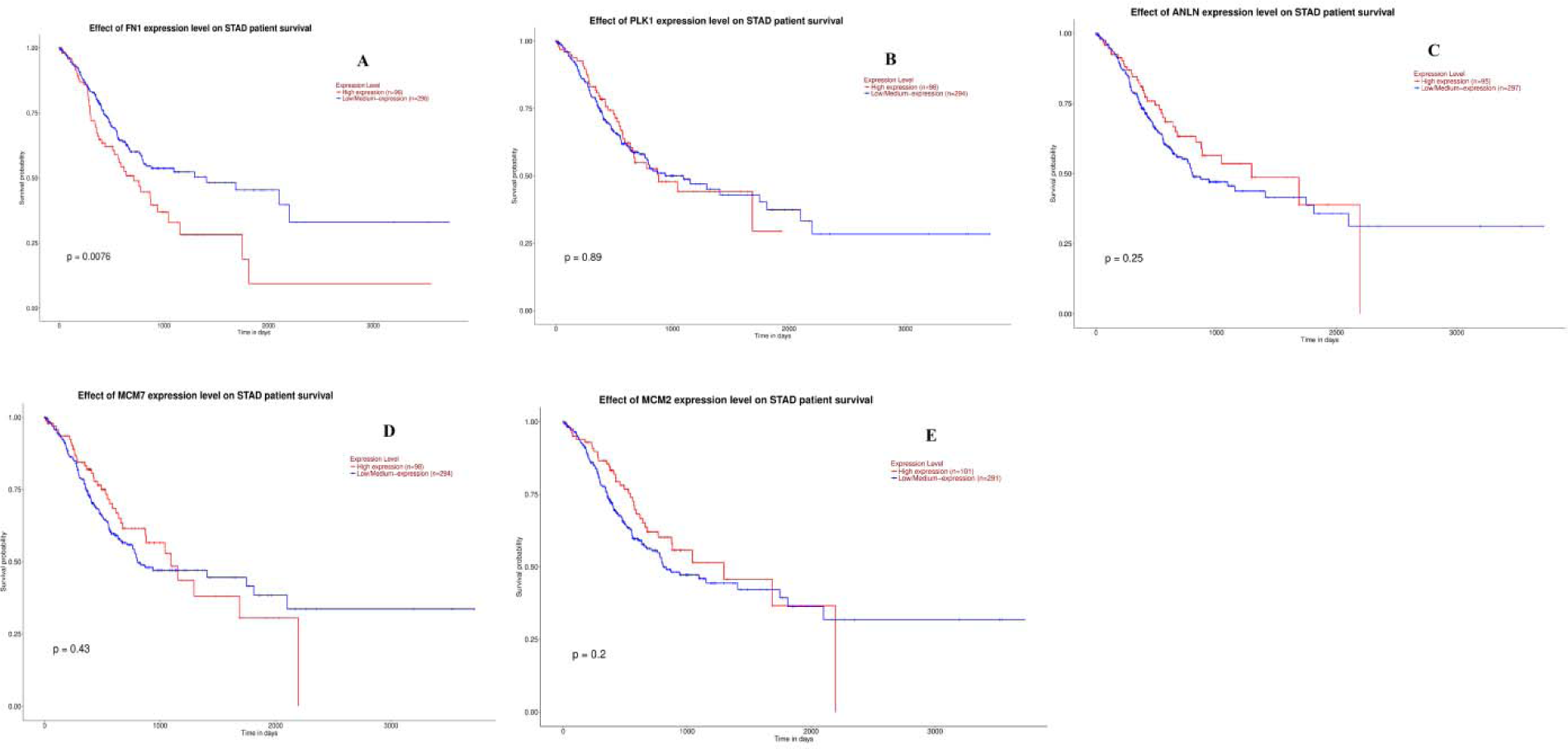
Survival analysis of up regulated hub genes. Survival analyses were performed using the UALCAN online platform. Red line denotes - high expression; Blue line denotes – low expression. A) FN1 B) PLK1 C) ANLN D) MCM7 E) MCM2

**Fig. 16.**
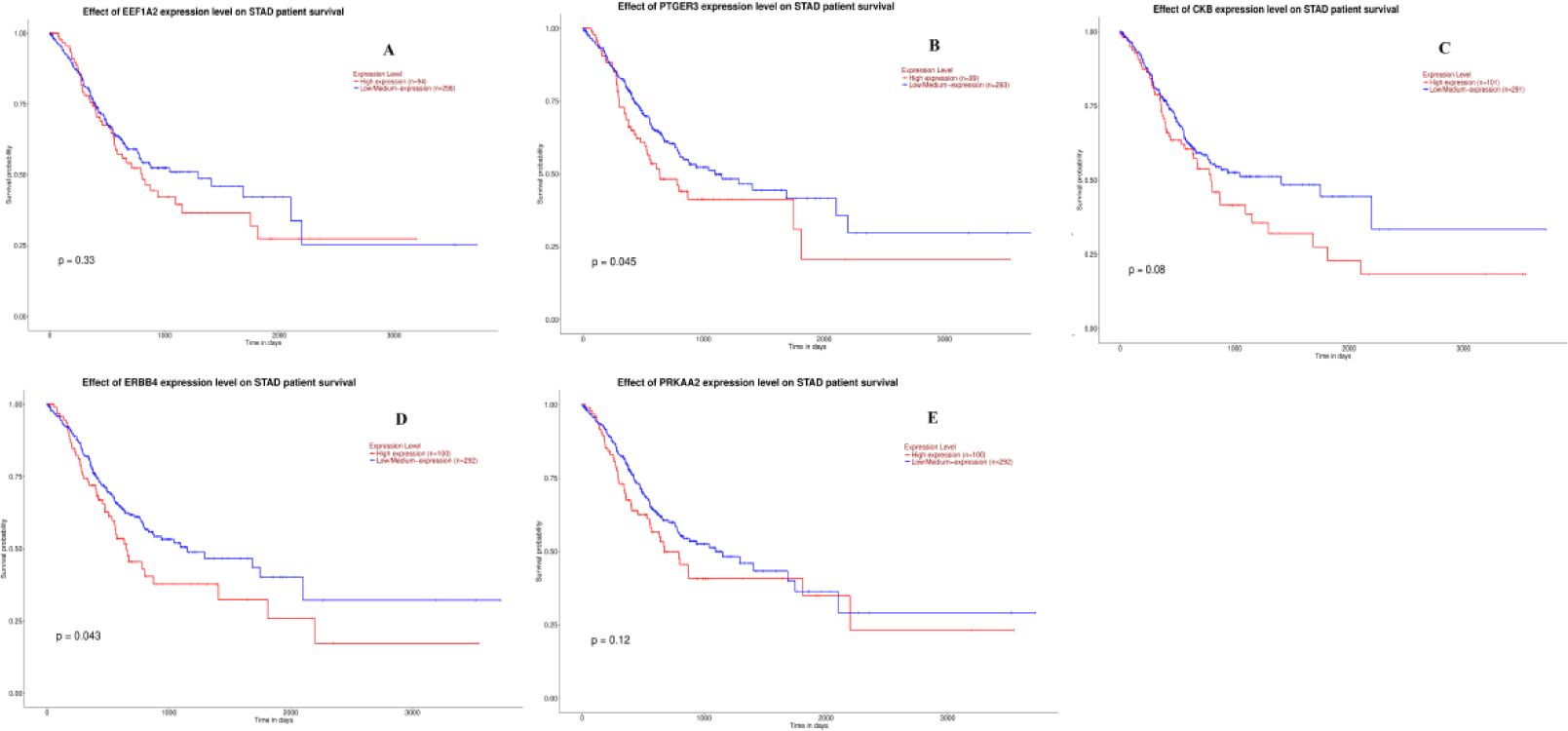
Survival analysis of down regulated hub genes. Survival analyses were performed using the UALCAN online platform. Red line denotes - high expression; Blue line denotes – low expression. A) EEF1A2 B) PTGER3 C) CKB D) ERBB4 E) PRKAA2

**Fig. 17.**
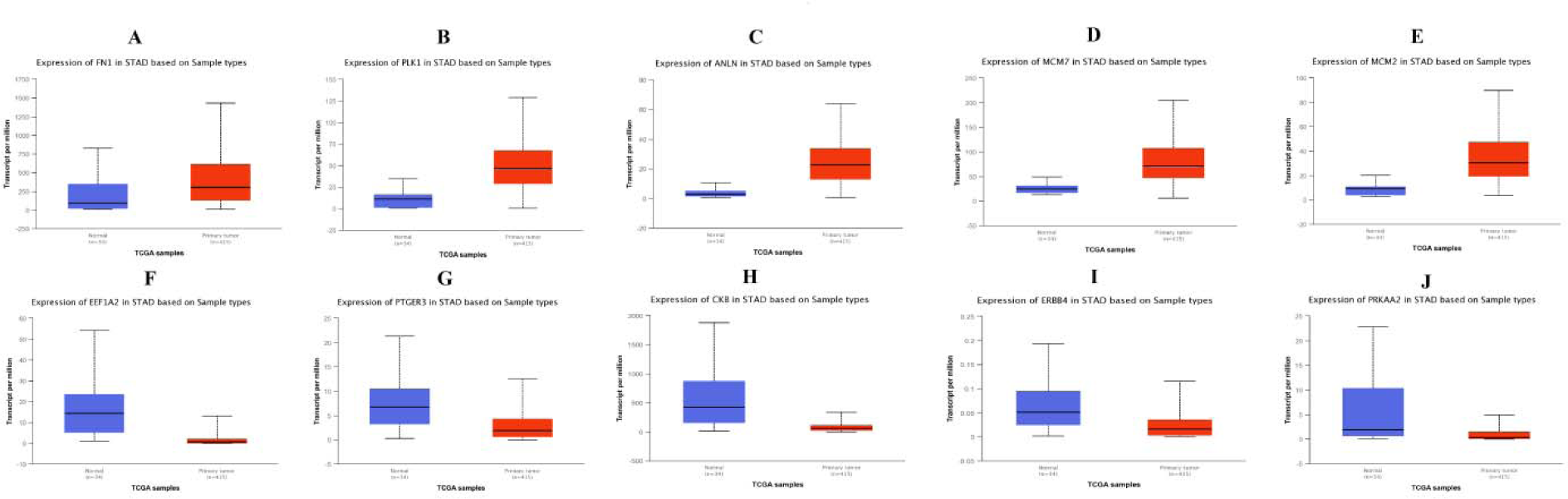
Box plots (expression analysis) hub genes (up and down regulated) were produced using the UALCAN platform. A) FN1 B) PLK1 C) ANLN D) MCM7 E) MCM2 F) EEF1A2 G) PTGER3 H) CKB I) ERBB4 J) PRKAA2

**Fig. 18.**
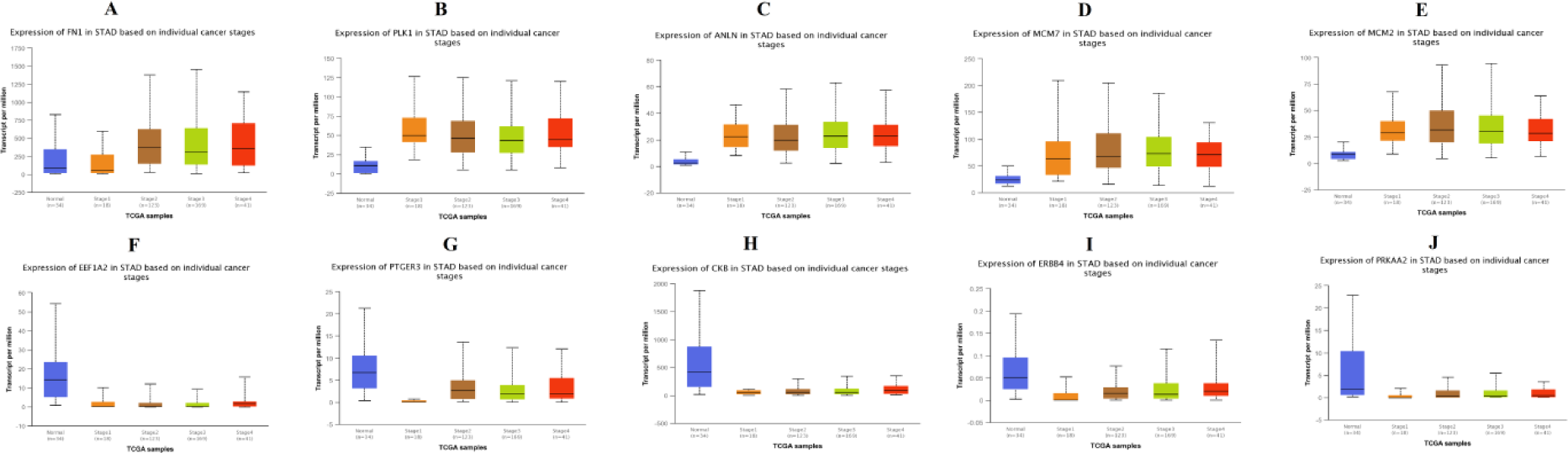
Box plots (stage analysis) of hub genes (up and down regulated) were produced using the UALCAN platform. A) FN1 B) PLK1 C) ANLN D) MCM7 E) MCM2 F) EEF1A2 G) PTGER3 H) CKB I) ERBB4 J) PRKAA2

**Fig. 19.**
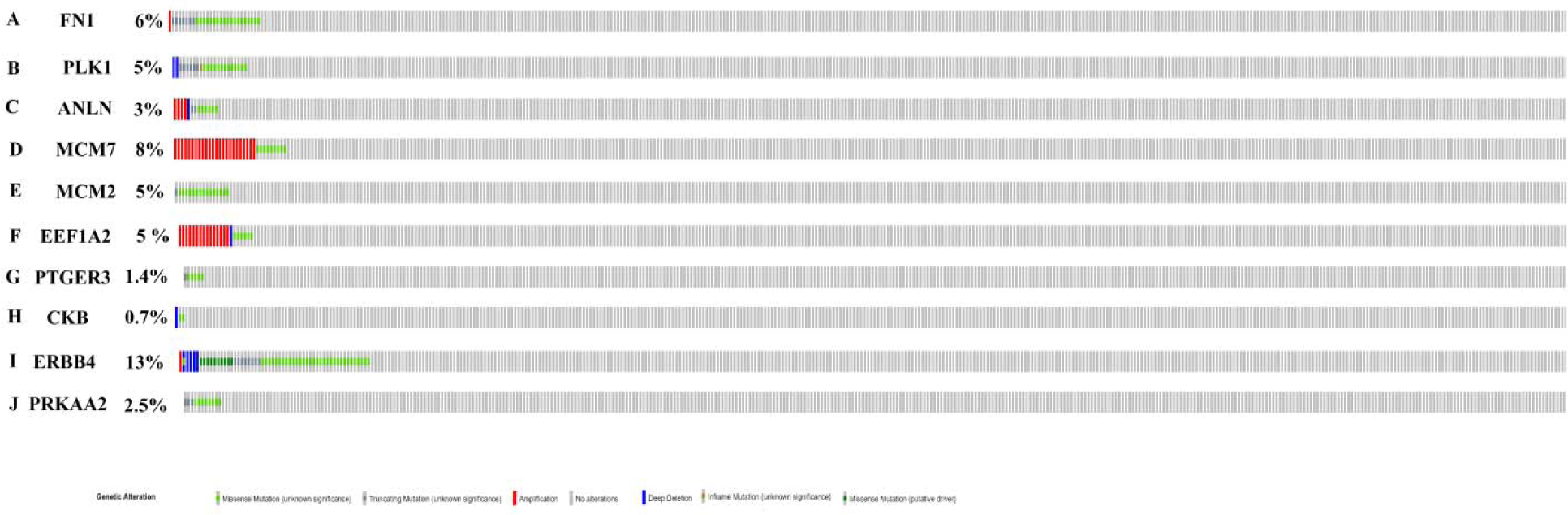
Mutation analyses of hub genes were produced using the CbioPortal online platform. A) FN1 B) PLK1 C) ANLN D) MCM7 E) MCM2 F) EEF1A2 G) PTGER3 H) CKB I) ERBB4 J) PRKAA2

**Fig. 20.**
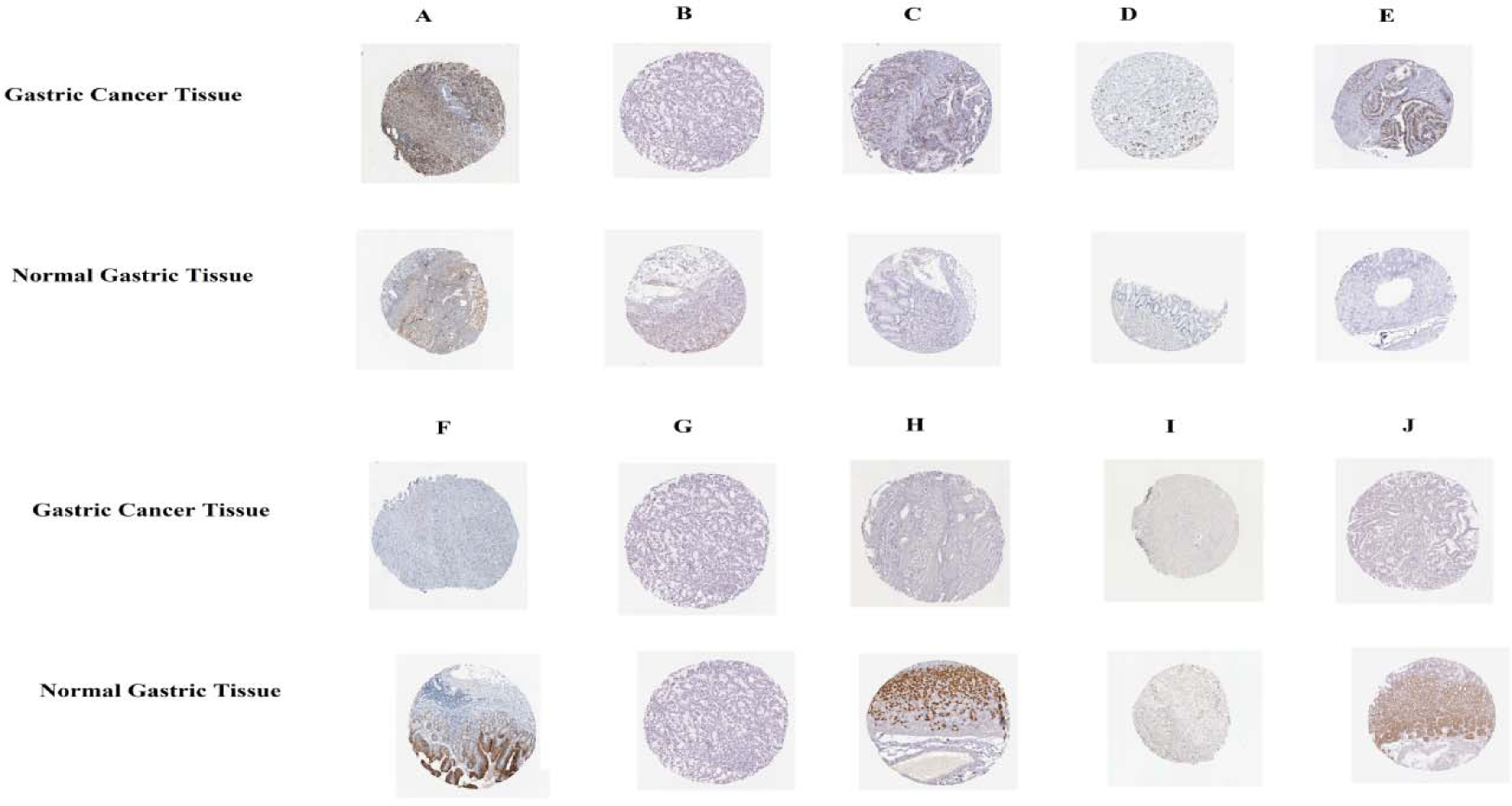
Immunohisto chemical (IHC) analyses of hub genes were produced using the human protein atlas (HPA) online platform. A) FN1 B) PLK1 C) ANLN D) MCM7 E) MCM2 F) EEF1A2 G) PTGER3 H) CKB I) ERBB4 J) PRKAA2

**Fig. 21.**
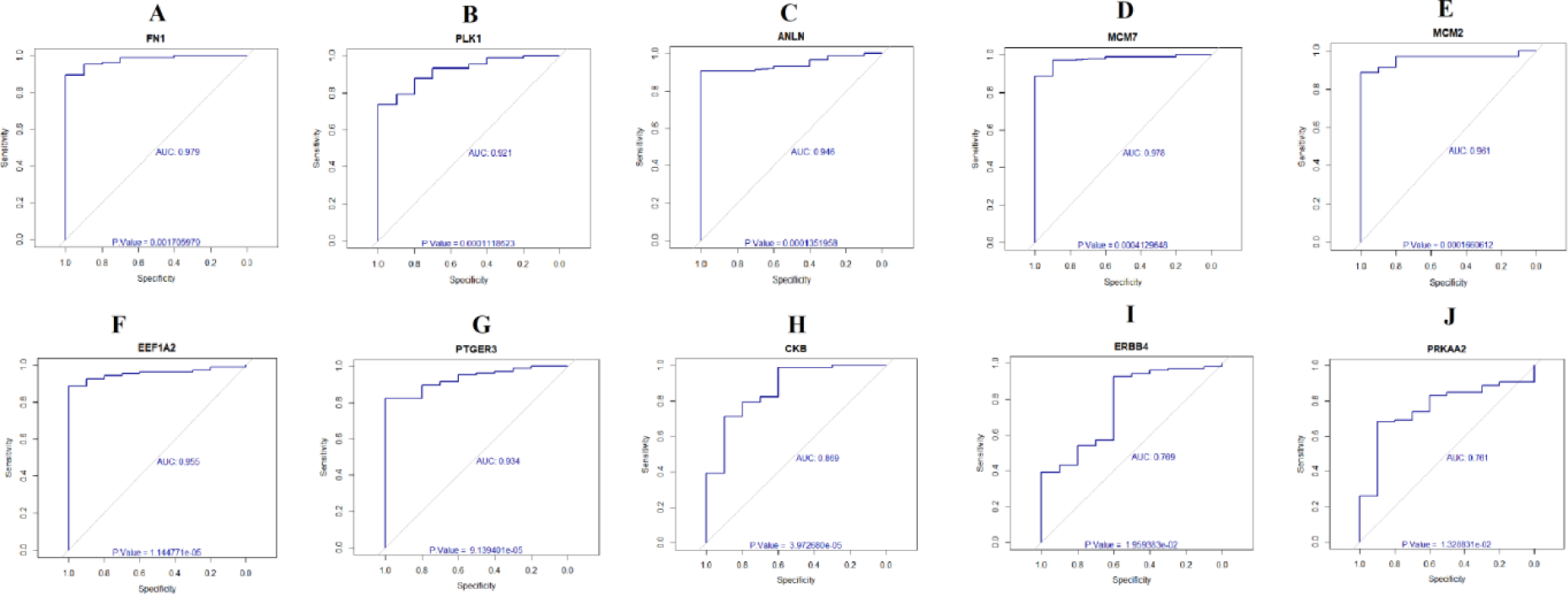
ROC curve validated the sensitivity, specificity of hub genes as a predictive biomarker for hepatoblastoma prognosis. A) FN1 B) PLK1 C) ANLN D) MCM7 E) MCM2 F) EEF1A2 G) PTGER3 H) CKB I) ERBB4 J) PRKAA2

**Fig. 22.**
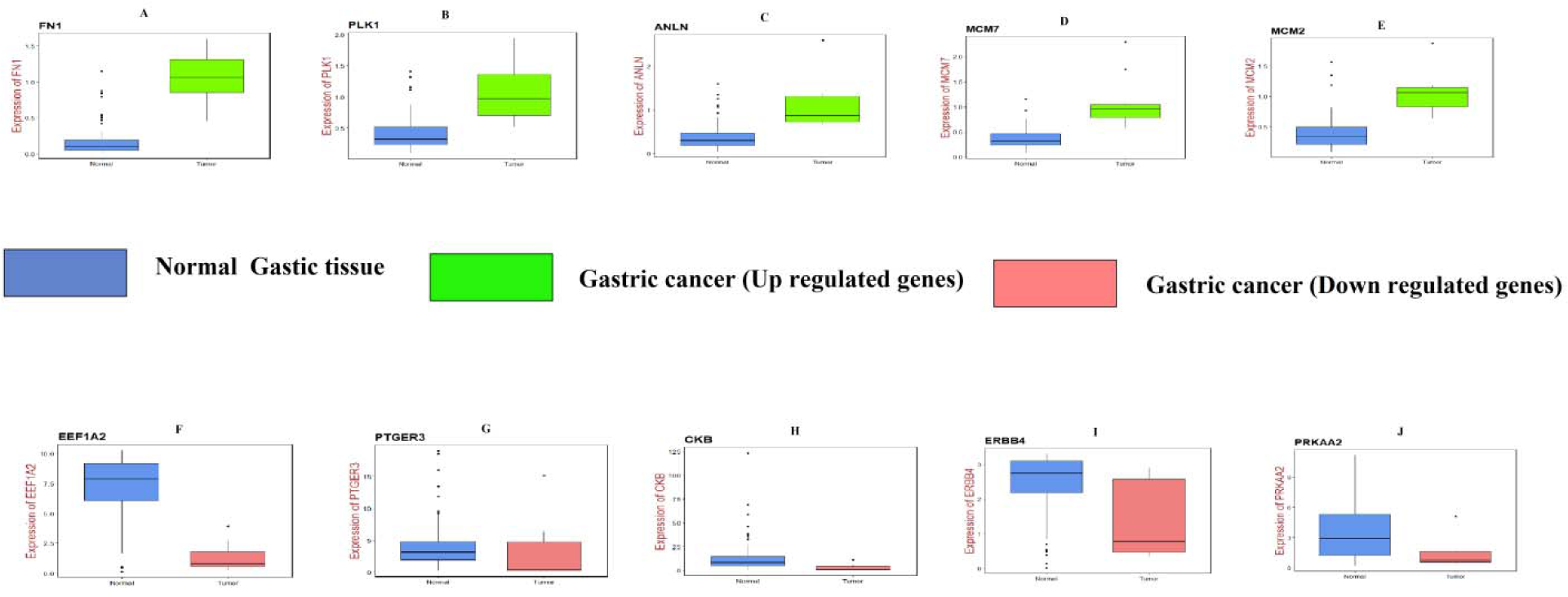
Validation of hub genes (up and down regulated) by RT- PCR. A) FN1 B) PLK1 C) ANLN D) MCM7 E) MCM2 F) EEF1A2 G) PTGER3 H) CKB I) ERBB4 J) PRKAA2

**Fig. 23.**
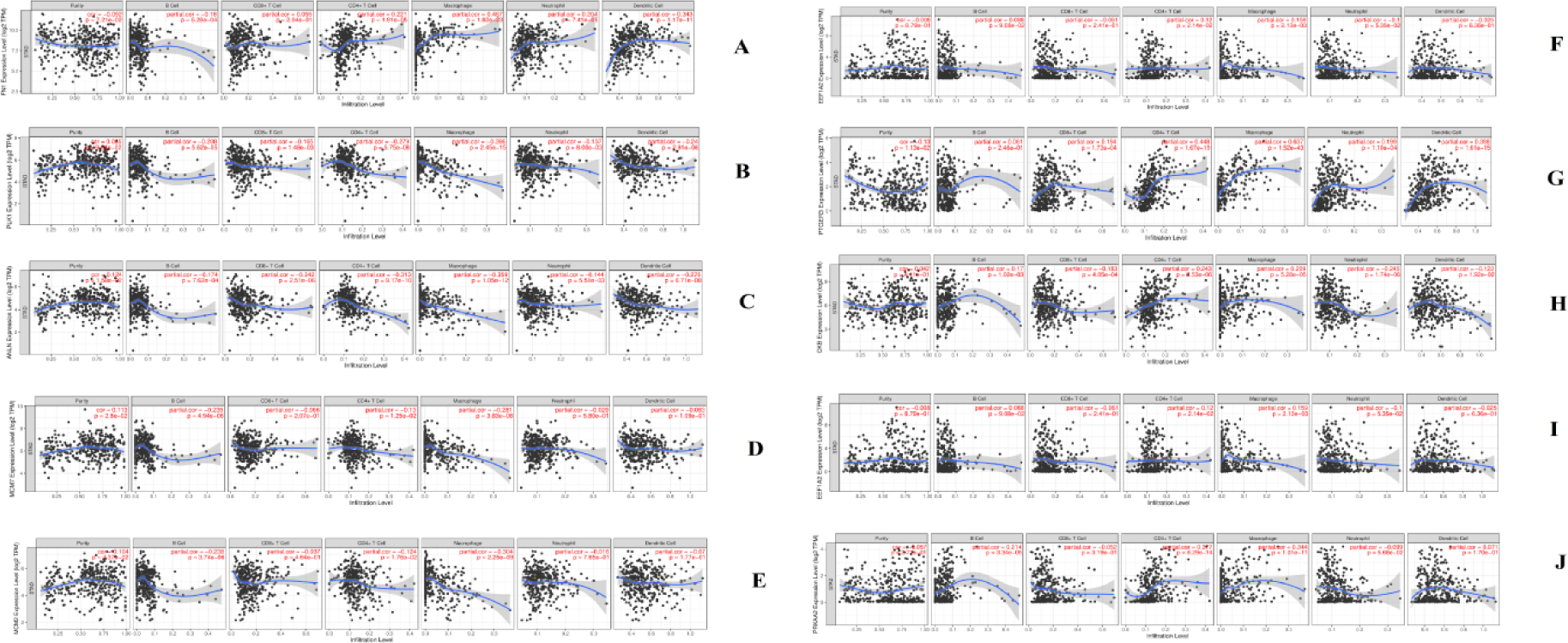
Scatter plot for immune infiltration for hub genes (up and down regulated). A) FN1 B) PLK1 C) ANLN D) MCM7 E) MCM2 F) EEF1A2 G) PTGER3 H) CKB I) ERBB4 J) PRKAA2

## Discussion

The frequency and fatality of GC were still on the increase in many countries. Gene alterations and abnormal expression have been exhibited in the advancement of GC. Understanding the molecular pathogenesis of GC is very essential for prognosis, diagnosis and treatment. With the advancement of microarray and high throughput RNA sequencing, the alternative expression levels of thousands of genes could be together examined. Consolidate and reanalyzing microarray data provide key data including hub genes, biological functions, and signaling pathways, which reveal novel clues for the prognosis, diagnosis and treatment of GC.

In this investigation, we extracted the RNA-seq data of GSE113255 from the GEO dataset with high-quality data and clinical characters. A total of 1008 DEGs were identified following RRA analysis, including 505 up and 503 down regulated genes. Genes such as COL4A1 [66], INHBA (inhibin subunit beta A) [67], RCC2 [68], THY1 [69], ADH7 [70], ATP4A [71], MAL (mal, T cell differentiation protein) [72] and ATP4B [73] were linked with development of GC. Polymorphic gene MSR1 was liable for advancement of prostate cancer [74], but this polymorphic gene may be involved in development of GC. Increases expression of CCKAR (cholecystokinin A receptor) was responsible for pathogenesis of gallbladder cancer [75], but over expression this gene may be important for advancement for GC.

To explore the possible role of up and down regulated genes in GC, we performed pathway enrichment analysis. Enriched up regulated genes such as TYMP (thymidine phosphorylase) [76], COL1A1 [77], COL1A2 [78], COL6A3 [79], FN1 [80], SPP1 [81], ITGA5 [82], THBS1 [83], THBS2 [78], THBS4 [84], MMP9 [85], ADAM12 [86], MFAP2 [87], FBN1 [88], MMP1 [89], MMP3 [90], MMP7 [91], MMP11 [92], MMP14 [93], MMP16 [94], NID2 [95], COL10A1 [96], COL11A1 [97], COL12A1 [98], ADAMTS2 [99], ICAM1 [100], VCAN (versican) [101], CTSK (cathepsin K) [102], HTRA1 [103], ASPN (asporin) [101], P4HA3 [104], BGN (biglycan) [105], BMP1 [106], ADAMTS9 [107], SERPINH1 [108], TIMP1 [109], LOXL2 [110], SERPINE1 [111], LTBP2 [112], LUM (lumican) [113], CLEC5A [114], EMILIN2 [115], SERPINB5 [116], SERPINE2 [117], MGP (matrix Gla protein) [118], AEBP1 [119], SPON2 [120], AGT (angiotensinogen) [121], CXCL9 [122] CTHRC1 [123], CCL3 [124], PLAU (plasminogen activator, urokinase) [125], C1QTNF6 [126], CCL18 [127], CXCL6 [128], SFRP2 [129], ANGPT2 [130], GDF15 [131], WNT2 [132], IGFBP7 [133], IL6 [134], IL11 [135], SPOCK1 [136], FNDC1 [137], SRPX2 [138], SULF2 [139], SULF1 [140], PLXDC1 [141], MUC16 [142], ESM1 [143], CXCL1 [144], TGFBI (transforming growth factor beta induced) [145], OSM (oncostatin M) [146], ANGPTL2 [147], FSTL1 [148], HMCN1 [149], PLXNC1 [150], GREM1 [151] and ITGBL1 [152] were involved in development of GC. Enriched up regulated genes such as COL4A2 [153], ITGA11 [154], LAMA5 [155], COL5A2 [156], ADAMTS4 [157], ADAMTS14 [158], ANXA13 [159], HTRA3 [160] and ADAMTS12 [161] were responsible for invasion of various types of cancer cells, but these genes may be associated with invasion of GC cells. Enriched up regulated genes such as COMP (cartilage oligomeric matrix protein) [162], TNC (tenascin C) [163], ITGB8 [164], FSTL3 [165] and PLXNA1 [166] were important for proliferation of many cancer cells types, but these genes may be liable for proliferation of GC cells. High expression of enriched genes such as COL3A1 [167], COL5A1 [168], COL7A1 [169], COL8A1 [170], CTSB (cathepsin B) [171], ELN (elastin) [172], CD109 [173], SERPINB9 [174], BMP8A [175], SEMA6B [176], MXRA5 [177], SFRP4 [178], CHRDL2 [179], CST1 [180], HAPLN3 [181], PXDN (peroxidasin) [182] and TGM2 [183] were liable for progression of many cancer types, but over expression of these genes may be responsible for pathogenesis of GC. Polymorphic gene COL18A1 was associated with progression of breast cancer [184], but this polymorphic gene may be linked with development of GC. Methylation inactivation of CPS1 was linked with development of hepatocellular carcinoma [185], but inactivation of this gene may be culpable for progression of GC. PGF (placental growth factor) was linked with angiogenesis in hepatocellular carcinoma [186], but this gene may be involved in angiogenesis in GC. Our investigation established that UPP1, ACTN1, ACAN (aggrecan), COL15A1, COL16A1, COL5A3, SPARC (secreted protein acidic and cysteine rich), ITGAX (integrin subunit alpha X), PCOLCE (procollagen C-endopeptidase enhancer), CAD (carbamoyl-phosphate synthetase 2, aspartate transcarbamylase, and dihydroorotase), CLEC7A, CRISPLD1, OMD (osteomodulin), LIF (LIF interleukin 6 family cytokine), PLXDC2, LGI2, TNFAIP6, PARVG (parvin gamma) and SDS (serine dehydratase) are up regulated in GC and has possible as a new diagnostic and prognostic biomarker, and therapeutic target. Enriched down regulated genes such as AKR1C3 [187], AKR1C2 [188], ADH1C [189], CYP2C19 [190], CYP3A4 [191], CYP3A5 [191], GSTA1 [192], GSTA3 [193], FOXA1 [194], FOXA2 [195], TTR (transthyretin) [196], ALDOB (aldolase, fructose-bisphosphate B) [197], APOA1 [198], ALDH1A1 [199], FBP2 [200], S100P [201], CCL28 [202], FGA (fibrinogen alpha chain) [193], FGB (fibrinogen beta chain) [203], FGG (fibrinogen gamma chain) [193], IGFBP2 [204], SCUBE2 [205], PDGFD (platelet derived growth factor D) [206], CTSE (cathepsin E) [207], MUC1 [208], MUC6 [209], SERPINB7 [210], NRG4 [211], SEMA3B [212], MUC5B [213], ANXA10 [214], REG3A [215], HRH2 [216], SST (somatostatin) [217] and CCKBR (cholecystokinin B receptor) [218] were associated with advancement of GC. Enriched down regulated genes such as AKR1C1 [219], MAOA (monoamine oxidase A) [220], ALDH3A1 [221], UGT1A10 [222], CYP2S1 [223], ME1 [224], FBP1 [225], GPT (glutamic--pyruvic transaminase) [226], COL2A1 [227], COL17A1 [228], HYAL1 [229], CSTA (cystatin A) [230], ADAM28 [231], BMP5 [232], ANGPTL3 [233], DPT (dermatopontin) [234] and APOBEC1 [235] were linked with invasion of many types of cancer cells, but these genes may be involved in invasion of GC cells. Enriched polymorphic genes such as UGT1A6 [236], CYP2C8 [237], CYP2C9 [238], GSTA2 [239], SULT2A1 [240], ADAMTSL1 [241], SERPINA5 [242] and APOC3 [243] were important for pathogenesis of many cancer types, but these polymorphic genes may be responsible for development of GC. Low expression of enriched genes such as AKR7A3 [244], ALDOC (aldolase, fructose-bisphosphate C) [245], SERPINA4 [246], ADAMTS15 [247], CCBE1 [248] and CLEC3B [249] were liable for progression of many cancer types, but decrease expression of these genes may be responsible for advancement of GC. Methylation inactivation of CXCL14 was linkes with progression of colorectal cancer [250], but loss of this gene may be key foe development of GC. Our investigation established that FMO5, FOXA3, CMBL (carboxymethylenebutenolidase homolog), AADAC (arylacetamidedeacetylase), GGT6, CYP4F12, CYP2C18, ACSM1, SULT1B1, SULT1C2, LGALS9C, COL6A5, F13A1, PAPPA2, COL4A6, HAPLN1, IGFALS (insulin like growth factor binding protein acid labile subunit), SLPI (secretory leukocyte peptidase inhibitor), LGALS9B and APOA4 are down regulated in GC and has possible as a new diagnostic and prognostic biomarker, and therapeutic target.

To explore the possible role of up and down regulated genes in GC, we performed GO enrichment analysis. Methylation inactivation of enriched up regulated genes such as ABCA1 [251], FLRT2 [252] and CDH13 [253] were liable for progression of many cancer types, but inactivation of these genes may be involved in growth of GC. Enriched up regulated genes such as FAP (fibroblast activation protein alpha) [254], ANTXR1 [255], FOXC1 [256], OLFML2B [257], APOC1 [258], APOE (apolipoprotein E) [259], PDPN (podoplanin) [260], FSCN1 [261], NOTCH1 [262], TNFRSF11B [263], CHI3L1 [264] and CD248 [265] were responsible for progression of GC. LIPG (lipase G, endothelial type) was liable for proliferation of breast cancer [266], but this gene may be liable for proliferation of GC cells. Enriched up regulated genes such as CCDC80 [267], PLA2G7 [268] and ENTPD1 [269] were associated with invasion of many types of cancer cells, but these genes may be linked with invasion of GC cells. Our investigation established that SH3PXD2B, CPZ (carboxypeptidase Z) and LRRC32 are up regulated in GC and has possible as a new diagnostic and prognostic biomarker, and therapeutic target. Enriched down regulated genes such as PGC (progastricsin) [270], AQP5 [271], CAPN9 [272], VSIG1 [273], KCNQ1 [274], TFF1 [275], TFF2 [276], CD36 [277], GKN1 [278], SCNN1B [279], FOLR1 [280], DUOX2 [281], RAB27A [282], HPGD (15-hydroxyprostaglandin dehydrogenase) [283], FA2H [284], LIPF (lipase F, gastric type) [285], GCKR (glucokinase regulator) [286], AKR1B10 [287], NQO1 [288], SELENBP1 [289] and GPX3 [290] were involved in development of GC. Enriched down regulated genes such as PBLD (phenazine biosynthesis like protein domain containing) [291], HPN (hepsin) [292], HOMER2 [293], DUOXA2 [294], SLC9A1 [295], DUOX1 [296], CA2 [297], CDH2 [298], ACADL (acyl-CoA dehydrogenase long chain) [299], PDIA2 [300], CYB5A [301], GALE (UDP-galactose-4-epimerase) [302], DHCR24 [303] and HBB (hemoglobin subunit beta) [304] were liable for invasion of many types of cancer cells, but these genes may be involved in invasion of GC cells. Low expression of enriched genes such as PTGER3 [305], RAB27B [306], ESRRB (estrogen related receptor beta) [307], ALDH6A1 [308], DHRS7 [309], LDHD (lactate dehydrogenase D) [310], DHRS9 [311] and ENTPD5 [312] were important for progression of many cancer types, but decrease expression of these genes may be responsible for advancement of GC. Enriched down regulated genes such as PRKAA2 [313], PTGR1 [314], ADHFE1 [315] and RGN (regucalcin) [316] were involved in proliferation of many types of cancer cells, but these genes may be associated with proliferation of GC cells. Our investigation established that PGA5, ASAH2, SLC26A7, CHIA (chitinase acidic), GHRL (ghrelin and obestatinprepropeptide), CAPN8, PGA3, PGA4, SLC9A4, GUCA2B, PROM2, UPK1B, SCNN1G, EPB41L4B, MYRIP (myosin VIIA and rab interacting protein), SLC26A9, SLC9A3, CDHR2, CA4, AQP10, PDZD3, HGD (homogentisate 1,2-dioxygenase), ERO1B, RDH12, AKR1B15 and TM7SF2 are down regulated in GC and has possible as a new diagnostic and prognostic biomarker, and therapeutic target.

The PPI network and module analysis found that DEGs were considered as up and down regulated hub genes in the network and modules. Up regulated hub genes such PLK1 [317], ANLN (anillin actin binding protein) [318], TREM2 [319], AMIGO2 [320], LEF1 [321], NOTCH3 [322], RAI14 [323], AQP1 [324], HOXB9 [325], MCM7 [326], MCM2 [327], CDC6 [328] and GINS4 [329] were liable for advancement of GC. Methylation inactivation of up regulated hub genes such as MDFI (MyoD family inhibitor) [330] and HEYL (hes related family bHLH transcription factor with YRPW motif like) [331] were involved in the progression of many cancer types, but inactivation of these genes may be key for advancement of GC. Increased expression of hub genes such as TREM1 [332], CYP2W1 [333], GPR161 [334], ARHGAP11A [335], APLN (apelin) [336], CBX2 [337], ORC1 [338] and MCM10 [339] were linked with development of many cancer types, but over expression of these genes may be liable for advancement of GC. Our investigation established that PRKDC (protein kinase, DNA-activated, catalytic subunitprotein kinase, DNA-activated, catalytic subunit) is up regulated in GC and has possible as a new diagnostic and prognostic biomarker, and therapeutic target. Down regulated hub genes such as SOX2 [340], EEF1A2 [341], ANG (angiogenin) [342], PTGDR2 [343], ERBB4 [344], KCNE2 [345], CA9 [346] and PSCA (prostate stem cell antigen) [347] were linked with pathogenesis of GC. Down regulated hub genes such as RIPK4 [348] and FAM3B [349] were involved in invasion of cells of many cancer types, but this gene may be associated with invasion of GC cells. Our investigation established that CNTD1, DCAF12L1, TTC39A, DUSP19, PTPRR (protein tyrosine phosphatase receptor type R) and TFR2 are down regulated in GC and has possible as a new diagnostic and prognostic biomarker, and therapeutic target.

The target genes - miRNA regulatory network analysis found that DEGs were considered as up and down regulated target genes in the network. Up regulated target genes such MYB (MYB proto-oncogene, transcription factor) [350] and SALL4 [351] were responsible for pathogenesis of GC. High expression of CLDN4 was key for progression of pancreatic cancer [352], but elevated expression this gene may be liable for progression of GC. Our investigation established that ARHGAP39 is up regulated in GC and has possible as a new diagnostic and prognostic biomarker, and therapeutic target. Down regulated target genes such as GATA6 [353] and KLF2 [354] were linked with development of GC. GLUL (glutamate-ammonia ligase) was involved in invasion of glioma cells [355], but this gene may be associated with invasion of GC cells. Our investigation established that ENPP5 is down regulated in GC and has possible as a new diagnostic and prognostic biomarker, and therapeutic target.

The target genes - TF regulatory network analysis found that DEGs were considered as up and down regulated target genes in the network. ABCA13 was responsible for advancement of GC [356]. Up regulated target geneABCD1 was associated with invasion of renal cancer cells [357], but this gene may be liable for invasion of GC cells. Down regulated target genes DNER (delta/notch like EGF repeat containing) [358] and IRX3 [359] were key for proliferation of many types of cancer cells, but these genes may be involved in proliferation of GC cells. CKB (creatine kinase B) was linked with progression of GC [360]. Down regulated target gene PKIB (cAMP-dependent protein kinase inhibitor beta) was important for invasion of breast cancer cells [361], but this gene may be linked with invasion of GC cells.

In our investigation, we screened FN1, PLK1, ANLN, MCM7, MCM2, EEF1A2, PTGER3, CKB, ERBB4 and PRKAA2 as important hub genes of GC by integrated bioinformatics analysis. Internal validation (survival analysis, expression analysis, stage analysis, mutation analysis, immunio infiltration analysis, immunio histochemical analysis and ROC analysis) and external validation (RT-PCR) of RNA and protein levels showed that they were higher or lower in cancerous than in normal control tissues. Over expression of the five hub genes and lower expression of the five hub genes were also associated with an advanced clinical stage and poor overall survival rate. Thus, the important hub genes explored in our investigation are likely to become a group of probale therapeutic targets of GC.

In conclusion, the current investigation identifird important hub genes linked with the various clinical stage and overall survival of GC patients. Our investigation attempts a profound perceptive of the molecular pathogensis linked with the adavancement of GC and implements probale biomarkers for early prognosis, diagnosis and individualized treatment of patients at various stages of GC.

## Acknowledgement

I thank Seon-Kyu Kim, Korea Research Institutue of Bioscience & Biotechnology, Personalized Genomic Medicine Research Center, Daejeon, South Korea, very much, the author who deposited their microarray dataset, GSE113255, into the public GEO database.

## Ethical approval

This article does not contain any studies with human participants or animals performed by any of the authors.

## Informed consent

No informed consent because this study does not contain human or animals participants.

## Author Contributions

Basavaraj Vastrad was associated with methodology and review and editing. Ali Chanabasayya Vastrad was performed software, supervision, formal analysis and validation.

## Availability of data and materials

The datasets supporting the conclusions of this article are available in the GEO (Gene Expression Omnibus) (https://www.ncbi.nlm.nih.gov/geo/) repository. [(GSE113255) (https://www.ncbi.nlm.nih.gov/geo/query/acc.cgi?acc=GSE113255]

## Consent for publication

Not applicable.

